# Integrated analysis of a compendium of RNA-Seq datasets for splicing factors

**DOI:** 10.1101/2020.03.24.006791

**Authors:** Peng Yu, Jin Li, Su-Ping Deng, Feiran Zhang, Petar N. Grozdanov, Eunice W. M. Chin, Sheree D. Martin, Laurent Vergnes, M. Saharul Islam, Deqiang Sun, Janine M. LaSalle, Sean L. McGee, Eyleen Goh, Clinton C. MacDonald, Peng Jin

**Affiliations:** West China Biomedical Big Data Center, West China School of Medicine (West China Hospital), Sichuan University, Chengdu, China; Medical Big Data Center, Sichuan University, Chengdu, China; Center for Epigenetics & Disease Prevention, Institute of Biosciences and Technology, College of Medicine, Texas A&M University, Houston, TX 77030, USA; Department of Electrical and Computer Engineering, Texas A&M University, College Station, Texas 77843, USA; Department of Human Genetics, Emory University School of Medicine, Atlanta, Georgia, USA; Department of Cell Biology & Biochemistry, Texas Tech University Health Sciences Center, Lubbock, Texas 79430, USA; Neuroscience Academic Clinical Programme, Duke-NUS Medical School, Singapore; Metabolic Reprogramming Laboratory, Metabolic Research Unit, School of Medicine and Centre for Molecular and Medical Research, Deakin University, Geelong, Victoria, Australia.; Department of Human Genetics, David Geffen School of Medicine, University of California-Los Angeles, Los Angeles, CA, USA; Department of Medical Microbiology and Immunology, Genome Center, and MIND Institute, University of California Davis, CA, USA

## Abstract

A vast amount of public RNA-sequencing datasets have been generated and used widely to study transcriptome mechanisms. These data offer precious opportunity for advancing biological research in transcriptome studies such as alternative splicing. We report the first large-scale integrated analysis of RNA-Seq data of splicing factors for systematically identifying key factors in diseases and biological processes. We analyzed 1,321 RNA-Seq libraries of various mouse tissues and cell lines, comprising more than 6.6 TB sequences from 75 independent studies that experimentally manipulated 56 splicing factors. Using these data, RNA splicing signatures and gene expression signatures were computed, and signature comparison analysis identified a list of key splicing factors in Rett syndrome and cold-induced thermogenesis. We show that cold-induced RNA-binding proteins rescue the neurite outgrowth defects in Rett syndrome using neuronal morphology analysis, and we also reveal that SRSF1 and PTBP1 are required for energy expenditure in adipocytes using metabolic flux analysis. Our study provides an integrated analysis for identifying key factors in diseases and biological processes and highlights the importance of public data resources for identifying hypotheses for experimental testing.

## Introduction

High-throughput expression profiling has been used to identify transcriptional changes associated with many diseases and biological processes (BPs). However, the mechanism underlying the associated changes remains mostly unclear. To study the underlying mechanisms, a large amount of high-throughput transcriptomic data have been generated for various upstream factors such as splicing factors (SFs). SFs are proteins regulating pre-mRNA splicing in various BPs and diseases including cancers^1, 2^. Reanalyzing available public data renders an efficient approach to uncover upstream factors in BPs and diseases.

Given the large scale of high-throughput expression profiling data that are publicly available, any method that can utilize these data to identify upstream factors of transcription in diseases and BPs will be of great value. High-throughput expression profiling has become routine, and much of the resulting data are available from online repositories, such as Gene Expression Omnibus (GEO)^3^. Up to the second quarter of 2019, GEO hosted more than 112,000 data series comprising more than 3,000,000 samples (**Figure S1**). As a popular method for transcriptome analysis, RNA-sequencing (RNA-Seq)^4^ has enabled genome-wide analyses of RNA molecules at a high sequencing depth with high accuracy. It has been used successfully on many mouse models^5, 6^, and thousands of RNA-Seq datasets have been generated and released to the public. This massive amount of biological data brings great opportunity for generating prominent biological hypotheses^7, 8^. However, these data were produced for diverse purposes and are not friendly to large-scale data integration. Therefore, substantial work is needed to build well-organized resources using these data to enable efficient and extensive integrated analysis. Here, we developed an integrated analysis to reveal upstream factors of post-transcriptional changes and transcriptional changes in diseases and BPs using these public RNA-Seq data.

We focused on datasets related to splicing factors (SFs), as approximately 95% of human multi-exonic genes are alternatively spliced^9^. We previously curated the metadata of a comprehensive and accurate list of mouse RNA-Seq data with perturbed SFs, which are hosted on our SFMetaDB^10, 11^. Using these metadata, the corresponding RNA-Seq data were used to compute alternative splicing changes related to perturbed SFs, represented in RNA splicing signatures. Because SFs may also mediate gene expression^12^, gene expression changes also were calculated to generate gene expression signatures. These signatures were used to determine the biological relevance of SFs to a disease or a BP using signature comparison^13^. Highly relevant SFs were considered key factors in the disease or BP.

A number of signature comparison approaches have been introduced to infer relations among various datasets. For example, connectivity map (CMAP) has been used to measure the connectivity of gene expression signatures between disease datasets and compound-treated datasets in drug repositioning^14^. Compared to signature comparison methods, datasets themselves are more critical for meaningful biological inference. In our present study, we combined the works of public dataset collection and signature comparisons. The public RNA-Seq datasets in SFMetaDB serve as a variable resource for generating splicing and gene expression signatures. Using these signatures, new evaluation may provide additional biological insights that would not be possible when analyzing these datasets alone.

To demonstrate the effectiveness and generalizability of our approach, we applied it to Rett syndrome (RTT)^15^ and cold-induced thermogenesis (CIT)^16^. Among the key SFs identified in RTT (e.g., cold-induced RNA-binding protein [CIRBP], SF3B1, PTBP1, PTBP2, and RBM3), *Cirbp* knockdown partially rescued the neurite outgrowth defects according to neuronal morphology analysis. In CIT, previous *in vitro* experiments supported several key SFs identified, such as CELF1, PRMT5, HNRNPU, and PQBP1. In addition, NOVA1 and NOVA2 identified by our analysis had been shown to suppress adipose tissue thermogenesis activation via *in vivo* experiments^17^. Here, we also show SRSF1 and PTBP1 to regulate energy expenditure in adipocytes using Seahorse metabolic flux analysis.

In summary, our systematic integration of disorganized and unstructured RNA-Seq datasets along with generated signatures provides a novel approach for identifying the most promising hypotheses for experimental testing. These novel hypotheses will form the basis for new experiments leading to the elucidation of detailed regulatory mechanisms at a molecular level.

## Results

### Generation of a signature database using a comprehensive collection of mouse RNA-Seq datasets with perturbed SFs

A signature database was constructed using a comprehensive collection of mouse RNA-Seq dataset metadata deposited in SFMetaDB, with each dataset having at least one SF perturbed. A group of 75 datasets was used to generate the signature database targeting 56 SFs (some SFs are perturbed in multiple datasets). Specifically analyzed in our workflow were more than 6.6-TB sequences from 1,321 RNA-Seq libraries from various mouse tissues and cell lines.

RNA-Seq datasets in SFMetaDB have various types of SF manipulation (**Figure 1a**). Specifically, most SFs in SFMetaDB have been knocked-out (60%), knocked-down (28.75%), overexpressed, knocked-in, and others (e.g., point mutation) in fewer datasets. Besides various types of manipulation of SFs, datasets in SFMetaDB also span over many tissues and cell lines (**Figure 1b**), of which the central nervous system‒related tissue/cell types are the most frequent, such as frontal cortex, neural stem cells, and neural progenitor cells. In addition, embryonic tissues and cell lines are another prominent source for studying SF perturbation.

**Figure 1.**
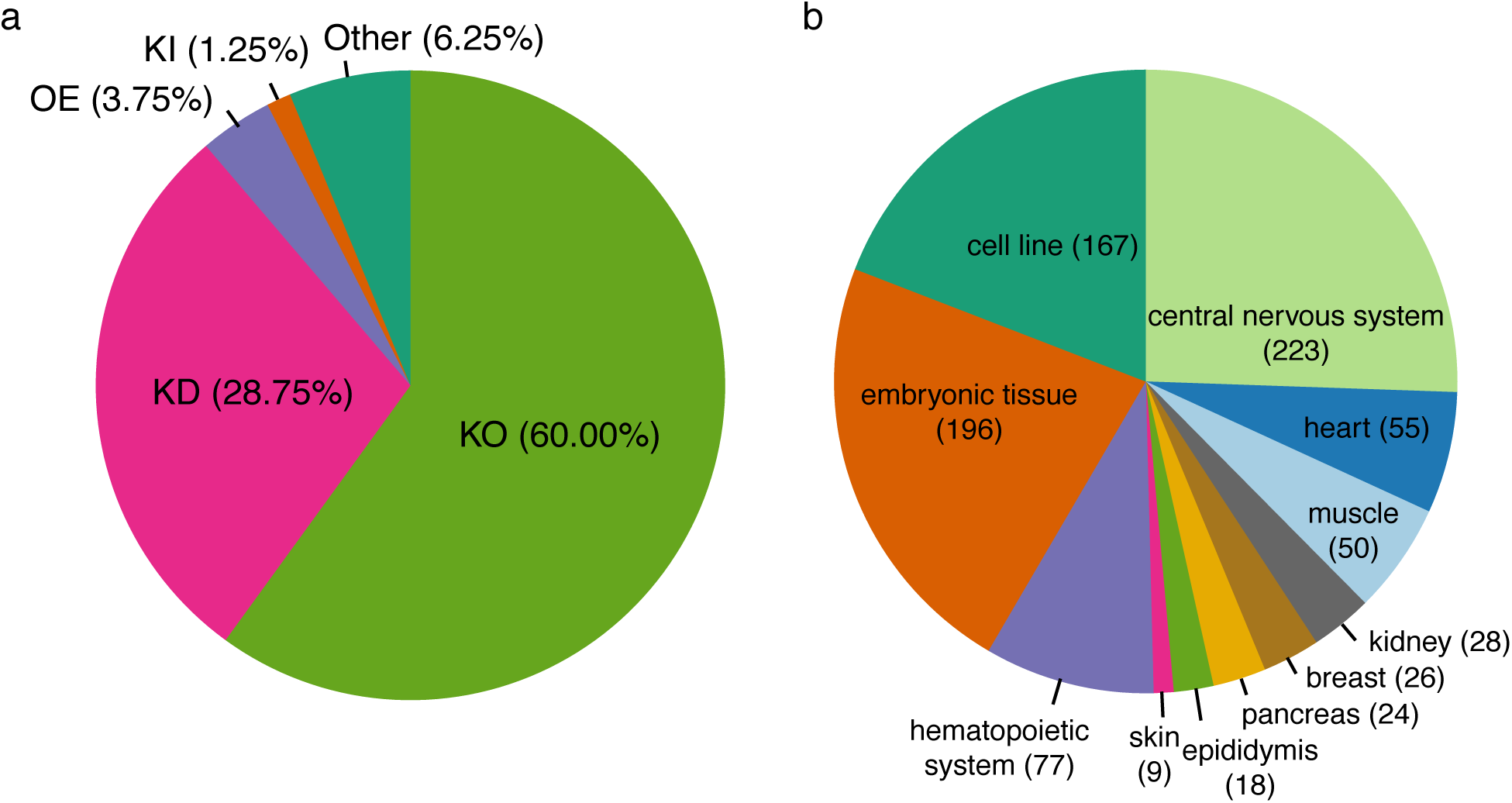
Meta-information of RNA-Seq datasets analyzed in the signature database. RNA-Seq datasets analyzed in our signature database include various perturbation and tissue types. (a) The pie chart shows the percentage of RNA-Seq datasets with perturbed SFs, including knockout (KO), knockdown (KD), overexpression (OE), knockin (KI), and other types (e.g., point mutation). (b) The pie chart depicts the number of RNA-Seq libraries for various tissues or cell lines.

To generate splicing and gene expression signatures for SFs, differential alternative splicing (DAS) and differentially expressed gene (DEG) analyses (see **Methods** section) were performed on the experimental comparisons of SF perturbation datasets. DAS events and DEGs formed splicing signatures and gene expression signatures for SFs. Among generated signatures, circular Manhattan overview plots show genome-wide splicing and gene expression changes regulated by SFs (**Data S1** and Figure 2).

**Figure 2.**
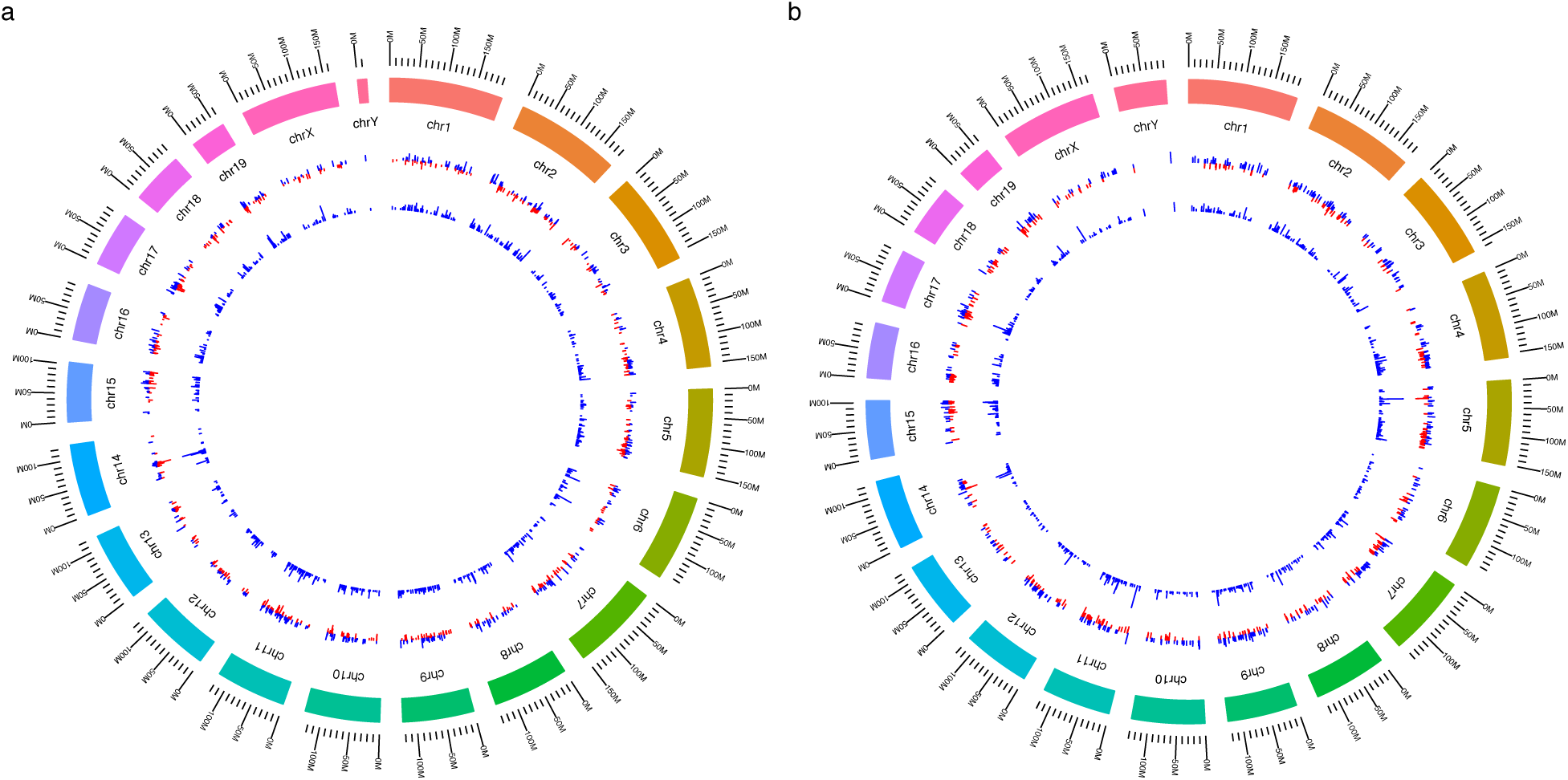
Genome-wide splicing and gene expression changes regulated by PRMT5. To evaluate splicing and gene expression changes regulated by SFs, circular Manhattan plots were generated across the whole genome (**Data S1**). This figure depicts the changes regulated by PRMT5 using the comparison in GSE63800. (a) Splicing changes are identified by ΔΨ > 0.05 and *q* < 0.05. Magenta or golden bars represent ΔΨs, and blue bars mean −log_10_(*q*-­-value). (b) Gene expression changes are identified by |log_2_(fold change)| > 0.5 and *q* < 0.05. Magenta or golden bars represent log_2_(fold change), and blue bars mean −log_10_(*q*-­-value).

### DAS events and DEGs of the datasets curated for SFMetaDB

To explore the entirety of our generated signature database, we examined the DAS events and the DEGs of the RNA-Seq datasets curated for SFMetaDB. Our DAS analysis identified large-scale splicing changes (**Figure S2a**), with exon skipping (ES) being the most common event type (**Figure S2b**), which is consistent with previous studies^18^. In addition, we also identified a large number of DEGs (**Figure S2c**). The normalized numbers of DAS events and DEGs correlated significantly (Pearson correlation coefficient *r* = 0.66, *p*-value = 2.29×10^*+^) in the selected comparisons (**Figure S2d**) (see **Methods** section), supporting the existence of potential crosstalk between the splicing process and the transcription process^19^.

### Identification of key factors in RTT

To demonstrate the effectiveness of our integrated analysis, it first was used to identify SFs in RTT, which is a severe neurological disorder^20^ without a cure. Because *Mecp2-*null mice feature RTT-like phenotypes^21^, our signature database and RNA-Seq data from *Mecp2*-deficient mice were integrated into our workflow to identify several key factors in RTT at the splicing and gene expression levels, respectively.

A DAS analysis was performed on the RNA-Seq data from dentate gyrus of six-week *Mecp2^-/y^* mice (see **Methods** section)^22^. Under |ΔΨ| > 0.05 and *q* < 0.05, 526 DAS events were identified in *Mecp2* knockout mice (**Table S1** and **Figure S3a**). The heatmap of percent-spliced-in (PSI, Ψ) values of ES events demonstrated large splicing changes in *Mecp2* knockout mice (**Figure S3b**). These large-scale splicing changes facilitated the downstream splicing signature comparison analysis in *Mecp2* knockout mice to elucidate key SFs that may regulate the splicing changes in RTT.

To discover key factors in RTT, a splicing signature comparison analysis was performed between the splicing signatures of the *Mecp2* knockout mice and each of the splicing signatures of the SF perturbation datasets (see **Methods** section). Out of 56 SFs, 7 SFs were identified as the potential key SFs that may regulate the splicing changes in *Mecp2* knockout mice (i.e., CIRBP, DDX5, METTL3, PRMT5, PTBP1, PTBP2, and SF3B1) (**Table S2**).

Among the identified SFs, CIRBP ranked highly (**Table S2**), indicating its potential role in modulating a significant number of splicing changes. We conducted a loss-of-function analysis to validate the role of *Cirbp* in the *Mecp2* knockout mice. The expression of *Cirbp* was increased significantly in *Mecp2* knockout mice according to our DEG analysis using RNA-Seq data (*q*-value= 1.27×10^*56^and log (fold change) = 1.064). This was confirmed experimentally using qRT-PCR (**Figure S4a**). A northern blot analysis of *Cirbp* also had shown that its expression level was up-regulated in RTT whole-brain samples^23^. Therefore, a knockdown of *Cirbp* was used to check whether it would rescue the neuronal morphology changes caused by lack of *Mecp2*. Here, the knockdown of *Cirbp* by shRNAs was efficient, as confirmed by the qRT-PCR assays (**Figure S4b)**. We analyzed the neuronal morphology of primary hippocampal neurons isolated from embryonic stage 18 (E18) rats, where replicates of neurons were examined from three groups of neurons, namely *Mecp2* knockdown, *Cirbp,* and *Mecp2* double knockdown, and the control (see **Methods** section)^24–27^. The representative neuronal images depict the neuron morphology for three groups of neurons (**Figure 3a**). Specifically, the branch numbers and the neurite lengths were decreased in *Mecp2* knockdown cells compared to the controls, but were partially rescued by the additional *Cirbp* knockdown (**Figure 3b-c**). These results suggest that the *Cirbp* knockdown can rescue the neurite outgrowth defects caused by *Mecp2* silencing.

**Figure 3.**
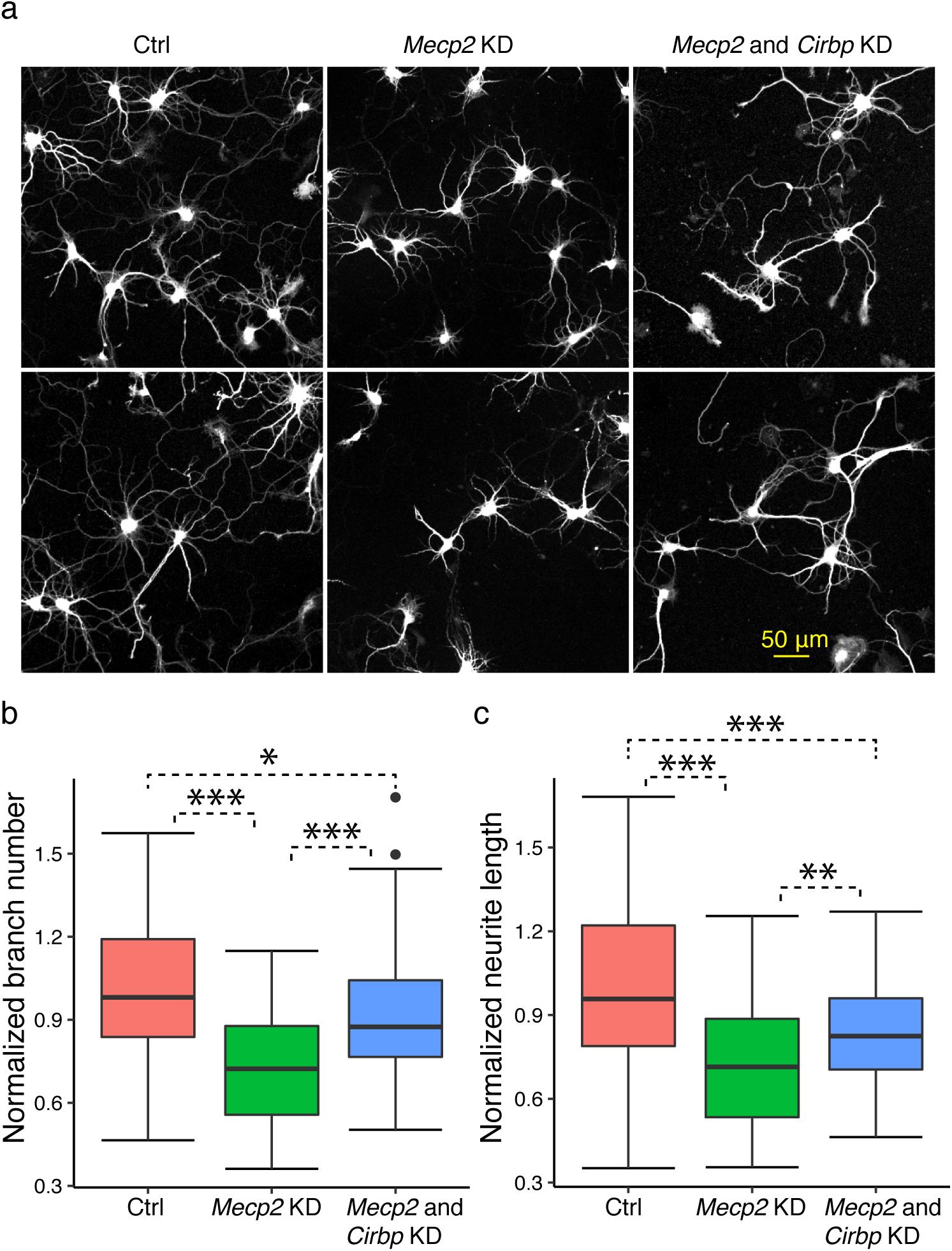
Neuronal morphology analysis on the role of *Cirbp* in RTT. (a) A neuronal morphology analysis was performed on the hippocampal neurons of *Mecp2* knockdown (KD), *Mecp2*-*Cirbp* double KD, and control. *Mecp2* KD neurons have less neurite outgrowth compared to normal neurons, yet *Mecp2*-*Cirbp* double KD neurons have more neurite outgrowth. (b) The normalized branch numbers are shown for control (Ctrl), *Mecp2* KD, and *Mecp2* and *Cirbp* KD neurons. ANOVA was used to test the changes of branch numbers among the three groups of neurons. *Mecp2* KD significantly reduced the branch numbers. The figure also depicts the significantly increased branch numbers in *Mecp2* and *Cirbp* KD neurons compared to *Mecp2* KD neurons but decreased branch numbers in *Mecp2* and *Cirbp* KD neurons compared to Ctrl. (c) The normalized neurite lengths were shown for Ctrl, *Mecp2* KD, and *Mecp2* and *Cirbp* KD neurons. ANOVA was used to test the changes of neurite lengths between the three groups of cells. *Mecp2* KD significantly reduced the neurite lengths. The figure also depicts the significantly increased neurite lengths in *Mecp2* and *Cirbp* KD neurons compared to *Mecp2* KD alone, but also significantly decreased neurite lengths in *Mecp2* and *Cirbp* KD neurons compared to Ctrl neurons. (ANOVA test. *: *p-*value < 0.05, **: *p-*value < 0.01, ***: *p-*value < 0.001. *n =* 47 to 61 neurons in each group).

To confirm the splicing changes in *Mecp2* knockout mice, the reverse-transcription polymerase chain reaction (RT-PCR) technique was performed on selected DAS events in *Mecp2* knockout mice (see **Methods** section)^28^. The effectiveness of our DAS analysis in RTT has been demonstrated in previous work^15^. Here, we specifically confirmed the potential effect of CIRBPs in this study by validating a subset of RTT DAS events that are also changed by CIRBP knockdown. A total of 11 predicted DAS events were tested by RT-PCR (see **Methods** section), and 8 events were differentially alternatively spliced in *Mecp2* knockout mice (**Figure S5**). These RT-PCR results confirmed the potential splicing regulatory contribution of CIRBP in RTT.

SFs may regulate gene expression alterations in various diseases and BPs. For example, *Celf1* promotes expression of *Cebpb* via interacting with *Eif2s1* and *Eif2s2* in proliferating livers and in tumor cells^29^. Therefore, we examined the potential role of SFs in regulating gene expression changes in RTT. A DEG analysis was performed on the *Mecp2* knockout mice to facilitate the key factors that regulate gene expression changes in RTT. Under log_2_(fold change) > 0.2 and *q* < 0.05, 579 genes were differentially expressed in *Mecp2* knockout mice (**Table S3**). The corresponding heatmap showed large expression changes in *Mecp2* knockout mice compared to wild-type mice (**Figure S6**).

To elucidate the key factors responsible for the expression changes in RTT, a gene expression signature comparison analysis was performed using the gene expression signatures of *Mecp2* knockout mice compared to the gene expression signatures derived from the SF perturbation datasets (see **Methods** section). The up-regulated genes in *Mecp2* knockout mice were compared to the up-/down-regulated genes in the SF perturbation datasets, and one SF was potentially responsible for the expression changes in *Mecp2* knockout mice, i.e. RBM3 (**Table S4**).

### Identification of key factors in cold-induced thermogenesis

To demonstrate the utility of our integrated analysis further, key factors of cold-induced thermogenesis (CIT) in adipose tissue were identified. CIT in adipose tissue can increase resting energy expenditure by approximately 10%^30^. If not compensated by changes in food intake, small changes in resting energy expenditure can have long-term effects on body weight. Therefore, activating CIT in adipose tissue is an attractive strategy to combat obesity. Although much is known about adipose commitment and differentiation^31^, the transcriptional mechanisms that ensure the readiness of mature adipose tissue to carry out adaptive thermogenesis remain unknown, including interactions between SFs and thermogenesis^32^. Thus, to improve our understanding of this complexity further, we combined the RNA-Seq data from SF perturbations and from adipose tissues under cold exposure to identify key factors relevant to CIT at the splicing and gene expression levels.

A DAS analysis was performed on RNA-Seq data from brown adipose tissue (BAT) and subcutaneous white adipose tissue (sWAT) from cold-exposed mice (see **Methods** section). Under |ΔΨ| 0.05 and *q* < 0.05, the DAS analysis revealed large-scale alternative splicing events in both BAT and sWAT upon cold exposure (**Figure 4a** and **Figure S7a**). Specifically, 760 and 1,481 alternative splicing events were identified in BAT and sWAT, respectively (**Table S5**). The heatmaps of PSI values demonstrated the large splicing changes of ES events in BAT and sWAT upon cold exposure (**Figure 4b** and **Figure S7b**).

**Figure 4.**
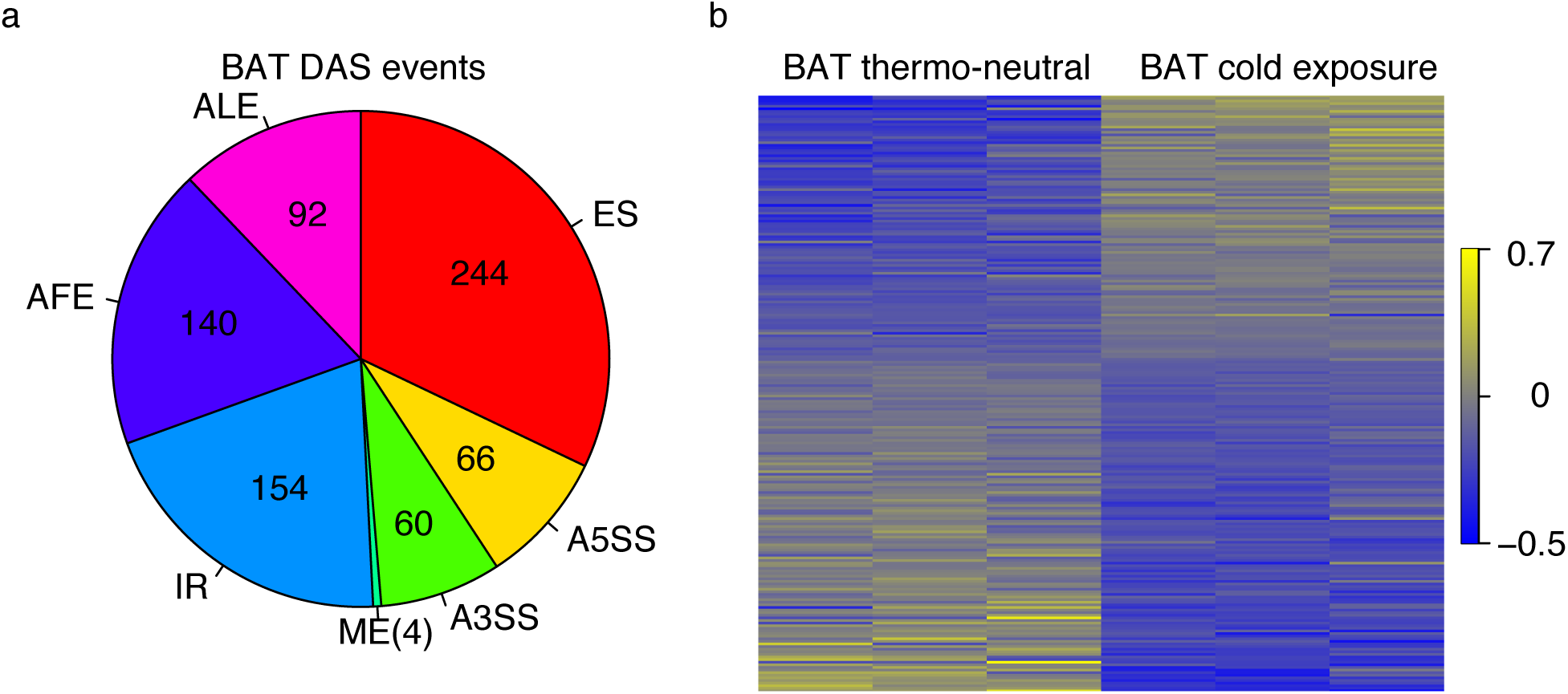
DAS events of BAT. (a) DAS analysis identified seven DAS event types, i.e. exon skipping (ES), alternative 5’ splice sites (A5SS), alternative 3’ splice sites (A3SS), mutually exclusive (ME) exons, intron retention (IR), alternative first exons (AFEs), and alternative last exons (ALEs). The pie chart depicts the number of DAS events of the seven splicing event types in BAT. (b) The heatmaps show the PSI values (scaled by standard deviation) for the differential alternative ES events in BAT. Yellow: high PSI value; blue: low PSI value.

To discover key factors in CIT, a splicing signature comparison analysis then was performed on the signatures of BAT and sWAT derived from cold-exposed mice compared to the curated SF perturbation datasets (see **Methods** section). Out of the full SF perturbation datasets that related to a total of 56 SFs, 2 SFs and 6 SFs were shown to be potentially responsible for the splicing changes in BAT and sWAT upon cold exposure, respectively. From these data, NOVA1 and PRMT5 were associated with splicing changes in BAT (**Table S6**). In addition, MAGOH, PRMT5, PTBP1, RBFOX2, RBM8A, and U2AF1 were linked with alternative splicing events in sWAT (**Table S6**). These SFs potentially regulate the splicing changes in adipose tissue that are critical for the activation of adipose tissue thermogenesis upon cold exposure.

In addition to the DAS analysis, a DEG analysis was performed on the RNA-Seq data of BAT and sWAT from cold-exposed mice to help identify key factors of gene expression in adipose tissue CIT. Under log_2_(fold change) > 1.0 and *q* < 0.05, a total of 1,836 and 5,266 genes were identified as differentially expressed in BAT and sWAT, respectively (**Table S7**). The heatmaps showed large expression changes in BAT and sWAT upon cold exposure (**Figure S8a** and **Figure S8b**).

A gene expression signature comparison analysis was performed using the gene expression signatures derived from adipose tissue upon cold exposure, compared to the gene expression signatures calculated from the curated SF perturbation datasets (see **Methods** section). Up-regulated genes in adipose tissue were compared to the up-/down-regulated genes in the SF perturbation datasets, and 22 SFs were shown to potentially regulate gene expression changes in BAT upon cold exposure (i.e., CD2BP2, CELF1, ESRP1, ESRP2, HNRNPK, HNRNPL, HNRNPU, MBNL1, MBNL2, METTL3, NOVA1, NOVA2, PQBP1, PRMT5, PRMT7, PTBP1, QK, RBM10, SF3A1, SF3B1, SRRM4, and U2AF1) (**Table S8**). In addition, 21 SFs were identified to potentially regulate gene expression changes in sWAT upon cold exposure (i.e., ACTA1, CELF1, CIRBP, EIF4A3, ESRP1, ESRP2, HNRNPA2B1, HNRNPU, MBNL1, MBNL2, MBNL3, PAF1, PHF5A, PRMT5, PRMT7, QK, RBFOX2, RBM17, RBM3, RBM8A, and U2AF1) (**Table S8**). These SFs potentially regulate the expression changes in adipose tissue upon cold exposure. We specifically evaluated SRSF1 in splicing signature comparison results for CIT, and a Seahorse metabolic flux analysis demonstrated a potential regulatory role of SRSF1 on mitochondria respiration in 3T3-L1 adipocytes (see **Methods** section). These adipocytes recently have been found to have characteristics of brown adipocytes, including high levels of uncoupled respiration^33, 34^. Such uncoupled respiration was assessed because it is a major component of CIT^35^. Knockdown of *Srsf1* in 3T3-L1 adipocytes by siRNA reduced *Srsf1* gene expression by approximately 95% (**Figure S9**; *p* < 0.0001) and reduced oxygen consumption rate (OCR) due to uncoupled respiration by approximately 35% (**Figure 5**; *p* = 0.02). This experimental finding supports the prediction that SRSF1 is an important factor in CIT. SRSF1 knockdown also reduced OCR coupled to ATP synthesis in these adipocytes (**Figure 5**, *p* < 0.001), suggesting that it has a broader role in regulating mitochondrial function. SRSF1 knockdown reduced basal adipocyte OCR, which is the sum of these two respiratory measures^36^ (**Figure 5**, *p* < 0.001). And our analysis showed no difference in extracellular acidification rate (ECAR), a proxy measure of glycolytic flux, in these adipocytes (**Figure S10**). Therefore, these data suggest a potential role of SRSF1 in regulating energy expenditure in adipose tissues specific to aspects of mitochondrial function.

**Figure 5.**
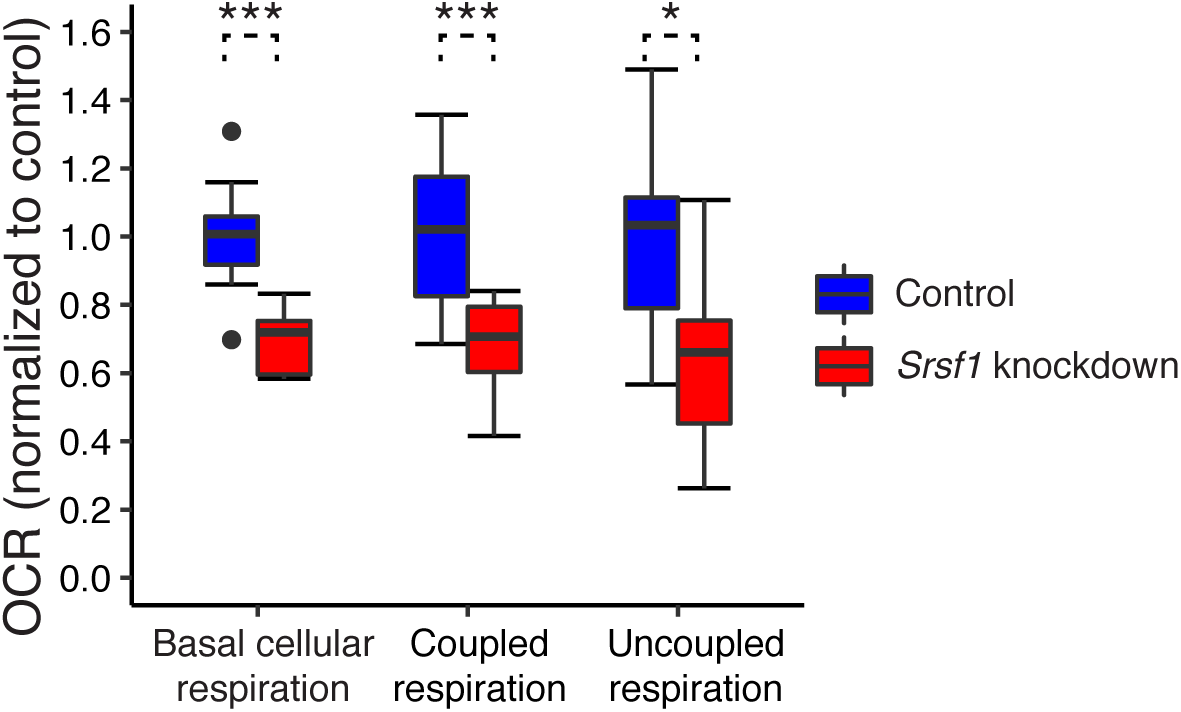
OCRs of mitochondrial respiration experiments with *Srsf1* knockdown. OCRs were recorded in Seahorse metabolic flux analysis. Three mitochondrial respiration OCRs (for basal cellular respiration, coupled respiration, and uncoupled respiration) were measured for both *Srsf1* knockdown adipocytes (red boxes) and controls (blue boxes). *Srsf1* knockdown adipocytes showed significantly reduced OCRs for all three measurements compared to controls (Unpaired *t-*test: *: *p*-value < 0.05, ***: *p*-value < 0.001, *n* = 8 to 10 in each group).

To determine whether PTBP1 is involved in adaptive thermogenesis, we performed *Ptbp1* knockdown using shRNA constructs in undifferentiated brown adipocytes (see **Methods** section)^37^. Partial *Ptbp1* knockdown was achieved with two independent shRNA constructs, as seen by western blot (**Figure S11a**). To investigate whether *Ptbp1* affected mitochondrial function, we performed *Ptbp1* knockdown in undifferentiated brown adipocytes before assessing mitochondrial respiration by Seahorse metabolic flux analysis. *Ptbp1* knockdown reduced cellular respiration and affected respiratory reserve capacity (**Figure S11b**)^38^. The decrease in respiration was caused mostly by coupled respiration. In agreement with the mitochondrial activity defect, we observed a decrease in mitochondrial complex abundance by western blot, especially for complexes I, II, and IV (**Figure S11c**). These results demonstrate that PTBP1 affects key components of brown adipocytes and may have a role during adaptive thermogenesis.

According to both the splicing signature analysis and gene expression signature analysis on cold-exposed adipose tissue, NOVA1 and NOVA2 were identified as key factors in CIT. Vernia et al.^17^ revealed that NOVA-deficient (both NOVA1 and NOVA2) mice have a significantly increased core body temperature compared to wild-type mice upon cold challenge. In addition, the expression of “browning” phenotype marker genes increased in subcutaneous adipocytes of NOVA-deficient mice. These findings indicate that NOVA proteins in adipocytes suppress adipose tissue thermogenesis^16^. Taken together, our results demonstrate the power of signature comparison analyses.

## Discussion

Technological advances have enabled RNA-Seq for the study of human diseases and BPs. However, RNA-Seq analyses have focused primarily on downstream expression changes, but the genes with changed expression do not necessarily play a critical role in regulating diseases or BPs. Standard RNA-Seq analysis with data limited to a specific biological context is unable to identify key factors in a disease or BP. We filled this void by generating a comprehensive compendium of RNA-Seq data for 56 SFs; these expression profiles were used in an integrated analysis to reveal key factors in diseases and BPs. DAS or DEG analysis alone only reveals genes whose expressions are changed in a disease or BP, but these genes do not necessarily regulate the disease process or BP. However, our signature comparison analyses aimed to reveal key factors that contribute to the regulation of the disease process or BP. As long as a significant portion of the SF targets derived in the SF perturbed datasets is maintained in the disease or BP, our integrated analysis is expected to reveal true factors that may not be ranked highly in DAS or DEG analysis. While the present study focused on an application in neuroscience and an application in metabolism, the integrated analysis described here can be generalized to other diseases and BPs. Thus, we expect that similar resources for other regulatory mechanisms and proteins, such as RNA-binding proteins^6^, as well as full-length RNA-Seq and proteogenomics data^39–41^, will serve as an important foundation for identifying key factors in human diseases and BPs.

A number of studies have been conducted to investigate splicing in RTT. For example, dozens of splicing changes were reported in a mouse model of RTT^42^ based on a splicing microarray study. A mutant gene in RTT, MECP2 physically interacts with dozens of proteins, including SFs PSIP1 and DHX9, and hundreds of alternative splicing events were misregulated in the cortex of *Mecp2* knockout mice^43^. However, few studies have identified key SFs in RTT development systematically. Our work provides an integrated analysis to fill the gap. For example, our identified factor CIRBP plays a critical role in controlling cellular responses to a variety of cellular stresses. Additionally, it has been shown that CIRBP migrates into stress granules under oxidative stress^44^. Interestingly, oxidative stress has been linked to RTT^45^. These facts suggest that CIRBP may affect RTT via the regulation of oxidative stress.

Some identified factors in CIT have been validated experimentally *in vivo* or *in vitro*. For example, NOVA1 has been validated previously *in vivo*^17^. In addition, evidence from *in vitro* experiments has been collected for other identified factors that may have roles in relevant processes related to CIT (**Online-only Table 1**). For example, they may affect adipogenesis (NOVA1^46^ and PRMT5^47^), the activity of BAT maker genes (CELF1^48^ and HNRNPU^49^), and lipid storage (HNRNPU^49^ and PQBP1^50^). In particular, CELF1 represses the expression of its targets by binding their 3’-UTR, such as *Ppargc1a* mRNA and BAT-enriched long noncoding RNA (lncRNA) 10 (lncBATE10). By repressing the expression of *Ppargc1a* mRNA and lncBATE10, thermogenesis was suppressed in brown adipocytes^48^. These experimental results corroborate our computational results, as CELF1 was predicted by our gene expression signature comparison instead of our splicing signature comparison. **Online-only Table 1** also records regulation directionality of the seven identified factors according to the gene expression changes of markers related to thermogenesis or other relevant BPs. The regulation directionality of three factors predicted by our method, CELF1, NOVA1, and PRMT5, were consistent with experimental results. An alternative direction for PRMT5 also was predicted (**Table S8**). This prediction is not surprising because PRMT5 is a protein arginine methyltransferase with many substrates^51^, and different conditions may lead to methylation of different substrates, which results in diversity in signatures. Such discrepancy can be resolved by additional experiments. The regulation directions of the remaining four factors, HNRNPU, HNRNPK, METTL3, and PQBP1, have not been determined because the corresponding references in **Online-only Table 1** do not contain expression data of marker genes.

In addition to the SFs supported by evidence of functional roles, those identified SFs without current literature support are connected to those with literature support according to the STRING database^52^ (**Figure S12**). For example, CIRBP, PTBP1, SF3A1, SRSF1, SRSF7, and U2AF1 share STRING associations with the SFs that have literature evidence (**Online-only Table 1**), suggesting that they also may affect thermogenesis in adipose tissue. Notably, some SFs are highly connected in the interaction network, such as PTBP1, SF3A1, SRSF1, and SRSF7. The *q*-value cutoff of 0.25 used in our integrated analysis may be relaxed. For example, SRSF1 in CIT has *p-*value = 0.045 and *q-*value = 0.59. These values indicate that additional SFs may be key factors as well. Specifically, we examined the functional roles of SRSF1 and PTBP1 in Seahorse metabolic flux analysis. *Srsf1* knockdown showed reduced OCR in multiple mitochondrial respiratory indices, suggesting a regulation role of SRSF1 for energy expenditure in adipose tissues. *Ptbp1* knockdown also reduced cellular respiration and affected the respiratory reserve capacity in brown adipocytes. Thus, the identified factors may be potentially critical candidates for future *in vivo* experiments that study CIT mechanisms.

Even though our integrated analysis identified many factors in CIT, SFs are not yet well-studied *in vivo*, with NOVA1 being the only one with *in vivo* experimental validation. Among more than 100 genes that enhance or suppress CIT supported by *in vivo* experiments^16^, SFs are not enriched (one-sided Fisher’s test, *p*-value = 0.8772). Thus, the importance of SFs is not appreciated by the CIT community fully. RNA-related BPs can be a fruitful direction for studying CIT mechanisms. Given that only approximately 15% of SFs currently have related RNA-Seq data^6^, more RNA-Seq data will be generated, which will fuel our integrated analysis to predict more key factors of CIT in the future.

Signatures generated from different tissues or cell types may share similarity for specific SFs. For example, four splicing and gene expression signatures of SRRM4 from different tissue/cell types shared significant similarities (**Figure S13a** and **Table S9**). However, there are cases in which signatures are different for the same SF perturbation in different cell types or tissues. For example, PRMT5 has two compact groups of splicing and gene expression signatures from different tissues (**Figure S13b**). Identification of multiple compact signatures for a given SF ensures that the signature comparison results will have a broad coverage of possible effects of the SF.

It is worth noting that although the data used to derive the SF signatures were not necessarily from neuronal and adipose tissues, they still could assist the identification of potential key regulatory factors of RTT and CIT, respectively. For RTT, the RNA-Seq data for CIRBP were from mouse embryonic fibroblasts (**Table S9**). Because our validation results for CIRBP indicated its potential role in RTT, it can be suggested that tissues other than neuronal tissues can be used to generate hypotheses related to neurological diseases. For CIT, cardiac tissues were used to generate the RNA-Seq data for CELF1 (**Table S9**). The heart has a connection to adipose tissue in that catecholamine signaling, which activates thermogenesis in BAT and browning of WAT^53^, also can lead to cardiomyopathy and heart failure^54^ when persistently activated in cardiac tissue. This potential suggests that the rich resources of publicly available gene expression data, despite the fact that they may not be from the tissue/cell type seemingly relevant to the biological problem at hand, should not be dismissed, and informative results can be derived from them. Our work is expected to extend beyond the current applications of neuroscience and metabolism, and the integrated analysis based on a compendium of SF RNA-Seq data is an efficient and economical approach to speed up the accurate identification of complex regulatory relationships in more disease and BP studies.

Some SFs belong to several protein types with different functions in the pre-mRNA-splicing process, including the SR family of splicing proteins, polypyrimidine tract-binding proteins, branch site-binding proteins, heterogeneous nuclear ribonucleoproteins (hnRNPs), and small nuclear ribonucleoproteins (snRNPs)^55^. **Table S10** shows classification of the identified SFs. To facilitate classification of the SFs, their Pfam family annotations were extracted from Uniprot because domain structures can elucidate the biological functions of proteins^56^. Although some SFs have clear functions according to the domains, there is still a subset of SFs that cannot be annotated unambiguously. For example, RBM10 has three Pfam domain family annotations (i.e., RNA recognition motif [PF00076], Zn-finger in Ran binding protein and others [PF00641], and G-patch domain [PF01585]). These SFs without a clear single domain classification were annotated as “Unclassified.” The classification result of the SFs provides a clue for a deeper understanding of mechanisms underlying their regulatory roles.

## Methods

### DAS analysis using RNA-Seq data

To identify the DAS events, we performed DAS analysis^57–61^. Briefly, the raw RNA-Seq reads first were aligned to mouse genome (mm9) using STAR^62^ with default settings, and those uniquely mapped reads were retained to calculate the counts of the reads for each exon and each exon-exon junction annotated in the UCSC knownGene (mm9) table^63^ using the Python package HTSeq^64^. UCSC mm9 annotation was downloaded from the table knownGene in the UCSC public MySQL database “mm9” hosting at “genome-mysql.cse.ucsc.edu” with the user “genome.” The mapped exon and exon-exon junction counts were used to construct a directed-acyclic graph representation of DAS events^65^. After modeling the counts in each alternative splicing event using Dirichlet-multinomial (DMN)^66^, the likelihood ratio test was used to test the significance of the changes in alternative splicing between the comparison conditions^67^. The Benjamini-Hochberg‒adjusted *q-*value was calculated from the *p-*values in the likelihood ratio test^68^. To integrate the effect size, PSI (Ψ) was calculated for the splicing events to examine the inclusion level of the variable exons over the total mature mRNAs^69^. The DAS events were identified under |ΔΨ| > 0.05 and *q* < 0.05.

### DEG analysis using RNA-Seq data

To identify the DEGs, we performed a DEG analysis using RNA-Seq data^70–73^. A count table was constructed by counting the number of reads aligned to each gene of each sample. The genes with low counts were filtered out for the downstream testing. Normalization and differential gene expression analysis were performed using DESeq2^74^. False discovery rate (FDR)‒adjusted *q*-values were calculated using the Benjamini-Hochberg procedure. The log_2_(fold change) also was calculated for each gene. The DEGs were identified under |log_2_(fold change)| > 0.5 and *q* < 0.05.

### Comparison of the number of DAS events and DEGs

Because of the crosstalk between splicing and transcription^19^, the DAS and DEG analysis results may reflect this relation. To demonstrate this relation, the difference in the experimental designs of the datasets must be accounted. Specifically, we formulated the numbers of DAS events and DEGs using linear models^67^.

### Linear models of the number of DAS events and DEGs

The number of DAS events and the number of DEGs were formulated as linear combinations of the main factors concerning experimental designs, i.e., the total number of reads (*T*), the effective read length (*L*), and the number of samples (*S*). The total number of reads refers to the number of reads of all the samples in comparison to the perturbed samples and the baseline samples. For single-end reads, the effective read length is just the read length; for paired-end reads, the effective read length is the sum of the length of each read in a pair. The number of samples is the sum of the perturbed and unperturbed samples in a comparison.

The number of DAS events was formulated by the following linear model:

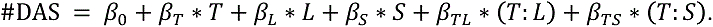

It includes the additive terms of the three metadata factors. More reads (*T*) results in more statistical power unless the number of reads is saturated. Given all the other conditions being the same, the greater the total number of reads (*T*), the more likely it is that the rare splicing junctions are covered. Longer effective read lengths (*L*) may result in more exon-exon junctions with reads mapped, leading to the detection of more DAS events. More samples (*S*) will result in more accurate estimates of variation, enabling more robust detection of DAS events. This linear model also includes two interaction terms. The interaction term *T:L* formulates the effect of *T* depending on the value of *L*. For example, increasing the number of reads (i.e., greater *T*) will make it more difficult to increase the number of detected DAS events in a short read length compared to a long read length because a short read length is more likely to have insufficient junction coverage. Similarly, the interaction term *T:S* formulates the effect of *T* depending on the value of *S*. However, our linear model does not include the interaction terms *L:S* and *T:L:S* because there is no prior knowledge that these two interaction terms should exist, and *t*-tests of the coefficients of these two terms were not statistically significant.

The number of DEGs was formulated by the following linear model:

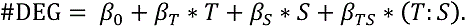

Different from splicing analysis, in which longer reads cover more splicing junctions enhancing the detection of DAS events, increasing read lengths beyond a reasonable length (e.g., 50 bps) will not increase read mappablility. Therefore, the effective length (*L*) is not expected to have an effect on the number of DEGs. Hence, the term *L* and the corresponding interaction term *T:L* were omitted in this linear model.

Information from the three factors was retrieved via the SRA Run Info CGI^75^. The coefficients of both linear models were estimated using `lm()` in R, and their significances were tested by *t-*test.

### Comparison of normalized numbers of DAS events and DEGs

Under the linear models, the number of DAS events and DEGs was normalized to a canonical experimental design for a fair comparison across multiple analyzed datasets. The normalization was performed by shifting standardized residuals in the fitted DAS and DEG models.

A subset of the comparisons was selected for examining the number of DAS events and DEGs. Some comparisons generated few DAS events because the original purpose to collect them was to study gene expression, which only needed relatively short reads. Much longer reads were needed to have enough coverage of exon-exon junctions for splicing analysis. These datasets designed for gene expression analysis performed poorly in detecting DAS events. Therefore, it was required to filter out these comparisons to produce a meaningful comparison between alternative splicing and gene expression analysis results.

The analysis of the linear model for DAS showed significant linear correlations of the number of DAS events with the total number of reads (*t-*test: *p-*value = 0.000172), the effective read length (*t-*test: *p-*value = 0.000825), and their interaction (*t-*test: *p-*value = 0.001188). We selected the high-quality comparisons with the total number of reads ≥ 100 million spots and the effective read length ≥ 150 bps because few DAS events were detected for the comparisons with < 100 millions of reads, and the three coefficients in the linear model became insignificant for comparing with effective read length < 150 bps. These two cut-off parameters were consistent with typical experimental design suggestions for splicing studies^76^.

### Splicing signature comparison analysis

To identify the regulatory role of key SFs in a BP, a splicing signature comparison analysis was performed among the splicing signatures derived from the experimental comparisons of SF perturbation datasets and the experiments in the BP, which was previously described by Li et al.^57^. A DAS event was considered positively regulated by an SF (notated as +) if the event had more inclusion of the variable exon (i.e., ΔΨ > 0.05) upon increased expression of the SF or if the event had less inclusion of the variable exon (i.e., ΔΨ < −0.05) upon decreased expression of the SF. Alternatively, an event was negatively regulated by an SF (notated as −) if the event had more inclusion of the variable exon (i.e., ΔΨ > 0.05) upon decreased expression of the SF or if the event had less inclusion of the variable exon (i.e., ΔΨ < −0.05) upon increased expression of the SF. Given the DAS events in one experimental comparison of the SF and the biological experiments, two vectors of +/−/0 were generated for both experimental comparisons, in which +/– indicated the event was positively or negatively regulated by the SF, and 0 meant that there was no evidence that the event was regulated by the SF. To compare two splicing signatures, a 3×3 contingency table was constructed with rows and columns named as +/–/0 that counted the number of events in the two splicing signatures. To determine the specific regulatory roles of an SF, the 3×3 table was collapsed into two 2×2 contingency tables so that two tables could be used to test the enrichment of + + events and − − events using Fisher’s exact test (with the null hypothesis H_0_: log-odds-ratio < 0.5), respectively. **Figure S14** illustrates how the splicing signature comparison analysis works with an example. FDR-adjusted *q*-values were calculated using the Benjamini-Hochberg procedure among *p-*values of all SF biological comparisons. Candidate SFs were identified by *q* < 0.25 ^77^.

### Gene expression signature comparison analysis

To determine the gene expression correlation relationship between SFs and DEGs in a BP, a gene expression signature comparison analysis was performed using gene expression signatures derived from the experimental comparisons of SF perturbation datasets and the BP. In the SF perturbation datasets, a DEG was up-regulated when log_2_(fold change) > 0.5 upon increased expression of the SF or when log_2_(fold change) < −0.5 upon decreased expression of the SF. In the opposite case, the DEG was down-regulated in the SF perturbation dataset. However, in the experimental comparison of the BP, the DEGs were up-/down-regulated when log_2_(fold change) > 0.5 or < −0.5, regardless of the expression changes of a specific SF. Given two gene sets of up-/down-regulated genes from experimental comparisons of the SF perturbation dataset and the BP, taking the expressed genes as background, Fisher’s exact test (with the null hypothesis H_0_: log-odds-ratio < 0.5) was used to test the significance of the genes shared by the two comparisons. Similar to the splicing signature comparison analysis, FDR-adjusted *q*-values were calculated using the Benjamini-Hochberg procedure among *p-* values of all SF biological comparisons. Candidate SFs were identified by *q* < 0.25.

### Generation of RNA-Seq datasets for *Mecp2* knockout mice

#### Animals

B6.129P2(C)-*Mecp2^tm1.1Bird^*/J mice were purchased from The Jackson Laboratory. Used to genotype the null allele (*Mecp2*^-^) were 5’-AAATTGGGTTACACCGCTGA-3’ (universal forward primer) and 5’-CCACCTAGCCYGCCTGTACT-3’ (knockout reverse primer). The universal forward primer and 5’-CTGTATCCTTGGGTCAAGCTG-3’ (wild-type reverse primer) were used to genotype the wild-type allele. Wild-type C57BL/6J male mice (The Jackson Laboratory) were bred with *Mecp2*^+/‒^ heterozygous females to generate *Mecp2*^‒/y^ mice and their wild-type littermates (*Mecp2*^+/y^). Mice for the experiments were euthanized by CO_2_ at 6 to 7 weeks old. Hippocampal dentate gyrus were dissected bilaterally and removed from the brain under a stereomicroscope^22^. All animal procedures were performed in accordance with the protocol approved by the Emory University Animal Care and Use Committee.

#### RNA isolation, RNA-Seq library preparation, and high-throughput sequencing

Total cellular RNA was purified from dentate gyrus using the TRIzol Reagent (Invitrogen), Phasemaker tubes (Invitrogen), and RNA Clean & Concentrator (Zymo Research) according to manufacturer instructions. DNase I treatment was included. RNA-Seq libraries were generated from 1 μg of total RNA from duplicated samples per condition using the TruSeq LT RNA Library Preparation Kit v2 (Illumina) following manufacturer protocol. Agilent 2100 BioAnalyzer and DNA1000 kit (Agilent) were used to quantify amplified complementary DNA (cDNA) and to control the quality of the libraries. Illumina HiSeq2500 was used to perform 100-cycle pair-end (PE100) sequencing. Image processing, sequence extraction, and adapter trimming were done using the standard cloud-based Illumina pipeline in BaseSpace.

#### RT-PCR for confirmation of alternative splicing changes in Mecp2 knockout mice

cDNA for RT-PCR was prepared from 120 ng of total RNA using SuperScript VILO MasterMix (Life Technologies) to verify gene expression levels. RT-PCR was performed using EmeraldAmp GT PCR Master Mix (Clontech) for 25 to 30 cycles with exon-specific primers as indicated^28^. PCR products were resolved on 2% agarose gel electrophoresis stained with ethidium bromide and visualized with an ultraviolet (UV) transilluminator. All events were tested in the dentate gyrus of three wild-type mice and five knockout mice (littermates) by RT-PCR. PSI was calculated by measuring the relative intensity of the PCR product, including the exon or retained intron, etc., divided by intensity of the PCR product, including the exon or retained intron, etc., plus the intensity of the PCR product, excluding the exon or retained intron, etc., multiplied by 100%. Statistical significance was calculated using a one-tailed Student’s *t*-test with unequal distribution of the variance and *p* smaller than 0.05 being considered significant.

#### RT-PCR confirmation of DAS events in Mecp2 knockout mice

To confirm DAS analysis in *Mecp2* knockout mice, a subset of DAS events was selected for RT-PCR experiment. Because *Cirbp* was increased significantly in *Mecp2* knockout mice (fold change: 2.04), and *Cirbp* was up-regulated in RTT whole-brain samples in a northern blot analysis^23^, we hypothesized that *Cirbp* may play a regulatory role in RTT. Therefore, we overlapped DAS events between *Mecp2* knockout mice and a dataset of *Cirbp* knockdown mouse embryonic fibroblasts (GSE40468). A total of 12 DAS events were commonly identified in two datasets (**Data S2**). Primer sequences of 11 DAS events were designed for RT-PCR experiments (**Data S3**), with the DAS event in *Vcam1* excluded because of low expressions of its variable exon in the event.

#### qRT-PCR for Cirbp RNA expression in Mecp2 knockout mice

Total RNA was isolated from mouse cortex using TRIzol Reagent (Invitrogen, Carlsbad, CA). cDNA was synthesized according to manufacturer protocol, using the QuantiTect Reverse Transcription Kit (Qiagen). Real-time qRT-PCR was performed using Sybr Green (Bioline) on an ABi ViiiA 7 in 384-well format using primers to Cirbp (F-GTCTTCTCCAAGTATGGGCAGAT, R-TCCTTAGCGTCATCGATATTTTC), with results normalized to GAPDH. Fold change was calculated relative to wild-type. Reactions were performed as three biological replicates.

### Neuronal morphology experiment

#### Primary neuron culture

Procedures for the dissociation and maintenance of primary neuron cultures were performed as described in previous work^24^. Briefly, time-mated Sprague-Dawley dams were euthanized via carbon dioxide asphyxiation, in accordance with guidelines set out by the SingHealth Institutional Animal Care and Use Committee. Embryonic day-18 embryos were extracted from the uterus and decapitated in order to remove their brains.

The harvested brains were placed in ice-cold Earle’s Balanced Salt Solution (EBSS) containing 10-mM HEPES. The hippocampi were dissected out, minced, and digested using papain (in EBSS) for 30 minutes at 37 °C. The digested tissues were resuspended in a neuronal plating medium (minimum essential medium containing 10% fetal bovine serum, 1× N2 supplement, 1× penicillin/streptomycin, and 3.6-mg/mL glucose). The tissue suspension was passed through a 70-μm cell strainer to sieve out tissue clamps. To obtain neuronal cells, the tissue suspension then was passed through a 7.5% bovine serum albumin (in phosphate-buffered saline [PBS]) layer by centrifuging at 200× g for 5 minutes. The resultant cell pellet was resuspended in a neuronal plating medium and seeded onto poly-l-lysine-coated glass coverslips (for immunohistochemistry) or culture plates (for RNA extraction). On the following day, the plating medium was exchanged for a maintenance medium (Neurobasal medium supplemented with 1× B27 supplement, 0.5× L-glutamine, and 1× penicillin/streptomycin). The cells were maintained in a humidified incubator at 37 °C and 5% CO_2_ level.

#### shRNA cloning and vector transduction

shRNAs against different regions of rat *Cirbp* were cloned into FUGW lentiviral vectors. The shRNA target sequence is GCAGGTCTTCTCCAAGTAT. The shRNA sequence against rat *Mecp2,* and the control sequence were cloned into PLL lentiviral vectors—shMeCP2: GGGAAACTTCTCGTCAAGA and shCtrl: AGTTCCAGTACGGCTCCAA^25^. shRNA expression was under the control of the human U6 promoter, while their fluorescent reporters (GFP or mCherry) were co-expressed under the control of the human ubiquitin C promoter in the same vector. Calcium phosphate precipitation was used to transfect the plasmids, and packaging and envelope proteins vectors into HEK293 cells for the production of lentiviruses, as previously described in previous work^26^. Viruses were collected via ultracentrifugation and resuspended in sterile PBS for use in transduction of primary neurons. The shRNA viruses were added to the cultures at days *in vitro* 1 (DIV 1). Thereafter, the cells were monitored for the expression of fluorescent tags to verify efficient expression of the shRNAs.

#### RNA extraction, cDNA conversion, and semiquantitative PCR

DIV 7‒cultured neurons first were washed twice with ice-cold PBS. Total RNA was extracted from the cells using an RNeasy Mini Kit (QIAGEN) according to manufacturer instructions. The RNA was eluted with nuclease-free water and stored at ‒-80 °C until use. cDNA was synthesized from 1-μg RNA using SuperScript^®^ III First-Strand Synthesis System (Invitrogen) with oligo(dT) primers. Conventional RT-PCR was performed on a DNA Engine Peltier Thermal Cycler (Bio-Rad Inc.). Primer sequences used were as follows—Cirbp: TCAGCTTCGACACCAATGAG (forward [F]), GTATCCTCGGGACCGGTTAT (reverse [R]) and Gapdh: CATCACTGCCACTCAGAAGA [F], CAACGGATACATTGGGGGTA [R]. PCR products were separated via agarose gel electrophoresis, and their band intensities were quantitated using ImageJ.

#### Immunohistochemistry

DIV 7 neurons grown on glass coverslips were washed twice with PBS and fixed with 4% paraformaldehyde (with 4% sucrose added to maintain osmolality) for 15 minutes at room temperature. Afterward, the cells were washed twice (5 minutes per wash) with Tris-buffered saline (TBS) to remove traces of the fixative. They then were permeabilized for 5 minutes with 0.1% Triton X-100 in TBS (TBS-Tx). Donkey serum (5%) in TBS-Tx was used to block the cells for 2 hours at room temperature. They then were incubated with primary antibodies overnight at 4 °C. Primary antibodies used were as follows—anti-GFP (1:3000, Rockland) and anti-Map2 (1:500, Sigma). On the following day, the primary antibodies were removed, and the cells were washed thrice with TBS-Tx. They then were incubated with Alexa Fluor^®^ secondary antibodies diluted 1:500 in TBS-Tx for 2 hours at room temperature. After removing the secondary antibodies, the cells were washed three times in TBS-Tx and then stained with DAPI (1:5000 in TBS-Tx) for 10 minutes at room temperature. After three final washes, two with TBS and one with phosphate buffer, the coverslips were mounted onto glass microscope slides and allowed to dry before imaging.

#### Image acquisition and analysis

Images were taken with a Zeiss LSM 710 confocal microscope. For examining neuronal morphology, images were taken at a single plane. Length measurements and tracings of neurites were made based on Map2 immunofluorescence using LSM Image Browser. To evaluate branch complexity, a Sholl analysis was performed on the tracings of the neurite arbors using an ImageJ plugin^27^.

#### Statistical analysis

At least 50 cells from three independent cultures were analyzed for each condition. Statistical testing was performed using GraphPad PRISM 5. One-way analysis of variance (ANOVA) with a Bonferroni test *post hoc* was used for comparing the conditions. Statistical significance was set at *p-*value < 0.05.

### RNA-Seq data generated from adipose tissues in cold treated mice

#### Animals

The study of adipose tissue upon cold exposure was performed on 20 male C57/Bl6 wild-type mice aged 8 to 12 weeks that were divided into two groups. The first group (n = 10) was exposed to thermoneutral temperatures (30 °C) for 72 hours, and the second group (n = 10) was exposed to cold (4 °C) conditions for 72 hours. All experimental mice were housed in a barrier animal facility with a 12-hour dark-light cycle, with free access to water and food. All animal experiments conducted in this study were approved by the Institutional Animal Care and Research Advisory Committee at the University of Texas Southwestern Medical Center (APN# 2015–101207).

#### RNA extraction and quantitative and quality RNA controls

sWAT and BAT were harvested and immediately snap-frozen in liquid nitrogen. Total RNA extraction was performed utilizing Trizol reagent (Invitrogen, Carlsbad, CA) and an RNeasy RNA extraction kit (#74106, Qiagen, Valencia, CA). Briefly, after homogenizing the tissues using a TissueLyser (Qiagen), RNA was isolated following manufacturer protocol (Qiagen, Valencia, CA). RNA quality and concentration were determined using a Nanodrop Spectrophotometer (N1-1000, Thermo Scientific, Wilmington, DE). RNA quality was confirmed using an Agilent 2100 Bioanalyzer following manufacturer protocol. The nanochip used for evaluating RNA quality produces electrophoresis peaks, from which the RNA integrity number (RIN) is calculated. RIN is the best predictor for assessing the integrity of the mRNA molecules. The RIN algorithm was calculated for all the normal and tumor tissue samples. The RIN is a decimal number ranging from 1 to 10, where 1 is attributed to completely degraded samples and 10 to intact RNA samples with very good quality. The main features taken into consideration for RNA quality evaluation are the size of the 18S and 28S peaks, the shape of these two peaks, the stability of the baseline, the appearance of additional peaks on the electropherogram, and the elevation of the baseline between the two peaks. A total of 1,000 ng of RNA was used to prepare libraries following Illumina TruSeq protocol. The criteria included the following: total RNA-Seq, 35M (4/lane) reads per samples, 100PE, long reads, full regular MC pipeline analysis. We performed the experiment on 20 mice. Ten mice were exposed to thermoneutrality, and 10 mice were exposed to cold temperatures (4 °C), both for 72 hours.

### Adipocyte mitochondrial respiration experiments

#### Cell culture and reverse transfection

Mouse-immortalized 3T3-L1 fibroblasts (ATCC) were maintained in growth media consisting of high-glucose DMEM (4.5-g/L glucose; Life Technologies) supplemented with 10% heat-inactivated fetal bovine serum (HI-FBS; Thermo Scientific). Cells were maintained in 10-cm dishes and differentiated into adipocytes upon reaching confluence, as previously described^78^. Briefly, cells were induced to differentiate by supplementing growth media with 3 nM insulin (Humulin R; Eli Lilly), 0.25-µM dexamethasone (Sigma-Aldrich), and 0.5 mM isobutyl-1-methyl xanthine (Sigma-Aldrich) for 3 days before being exposed to growth media supplemented with 3 nM insulin only for an additional 4 days.

Adipocytes then were maintained in normal growth media for 24 hours before undergoing reverse transfection of control and *Srsf1* siRNA^79^. For respiration analyses, Seahorse V7 plates were coated with ECM (Sigma-Aldrich), and siRNA was prepared for transfection using OPTI-MEM and RNAiMAX (Life Technologies). Per well, 0.45 μL of RNAiMAX was added to 7.5 μL of OPTI-MEM and 0.45 μL of 10-μM ON-TARGETplus SMARTpool siRNA (Dharmacon) directed against *Srsf1*, or nontargeting control siRNA was added to 7.5 μL of OPTI-MEM. The diluted RNAiMAX was mixed with diluted siRNA, and 15 μL per well was added to ECM-coated Seahorse plates and incubated at room temperature for 25 minutes. Adipocytes were trypsin-digested from their 10-cm dish before being seeded into the Seahorse plate at 10,000 cells per well. For gene expression analysis, the differentiation and reverse transfection protocol was followed as described above, but adipocytes were seeded into 12 well plates.

#### Respiration analysis

Cells were maintained for 3 days in reverse transfection media prior to respiration analysis. One hour prior to the assay, media were replaced with assay media consisting of unbuffered DMEM (Life Technologies), 25-mM glucose (Sigma-Aldrich), 1-mM sodium pyruvate, and 1-mM Glutamax (Life Technologies), a pH of 7.4, and were incubated at 37 °C in a non-CO_2_ incubator for 60 minutes. Cellular respiration was assessed, and mitochondrial function parameters were calculated as previously described^34^ using the Seahorse XF24 analyzer. Specifically, OCR was measured before and after injection of inhibitors to derive mitochondrial respiration parameters. Initially, 3 basal measurement cycles consisting of 3-minute mix, 4-minute wait, and 2-minute measure periods were performed, and basal respiration was derived by subtracting nonmitochondrial respiration from the baseline cellular OCR. Next, oligomycin, an inhibitor of ATP synthase (mitochondria complex V) was injected (1 μM final; Sigma-Aldrich), with results able to be used to derive the coupled respiration (also called ATP-linked respiration) and uncoupled respiration (i.e., proton leak). This step was followed by another 3 measurement cycles before the injection of rotenone and antimycin A (both 1 μM final; Sigma-Aldrich). Rotenone is a mitochondria complex I inhibitor, and antimycin A is a mitochondria complex III inhibitor. They shut down mitochondrial respiration, enabling the calculation of nonmitochondrial respiration. Respiration coupled to ATP-linked respiration was defined as the difference between respiration under basal and oligomycin conditions, and proton leak was calculated as the difference in respiration between oligomycin and rotenone/antimycin A conditions. Two independent experiments consisting of five biological replicates were pooled by normalizing all respiration indices to control group values within each experiment.

#### Gene expression analysis

Cells were maintained for 3 days in reverse transfection media prior to collection. Media was aspirated from cells and replaced with 500-μL Trizol (Thermo Fisher), which was collected and frozen at ‒80 °C. Samples were thawed later, and RNA was extracted using RNeasy columns (Qiagen). RNA was reverse-transcribed to cDNA using the SuperScript III First-Strand Synthesis System (Life Technologies), and real-time RT-PCR was performed as previously described^80^, with primers for *Srsf1* (Forward 5’-GGC TAC GAC TAC GAC GG TA-3’ Reverse 5’-GGA GGC AGT CCA GAG ACA AC-3’) and using cyclophilin (Forward 5’-CCC ACC GTG TTC TTC GAC A-3’ Reverse 5’-CCA GTG CTC AGA GCT CGA AA-3’) as the housekeeping gene.

#### Statistics

All data were expressed as mean ± SEM and were analyzed by unpaired *t*-test. Statistical significant differences were identified where *p* < 0.05.

### *Ptbp1* knockdown in brown adipocytes

#### Cell culture

An established mouse brown adipocyte cell line was obtained from Dr. Bruce Spiegelman (Dana-Farber Cancer Institute, Boston, US). Four mouse shRNA constructs against *Ptbp1* and a scrambled shRNA in pGFP-V-RS vector were purchased from Origene. shRNA constructs B (TTCTCTAAGTTTGGCACCGTCCTGAAGAT) and D (ACAATGATAAGAGCAGAGACTACACTCGA) were efficient at knocking down endogenous *Ptbp1* expression and were selected to perform experiments. Plasmids were transfected with BioT reagent (Bioland) according to the manufacturer protocol. Cells were analyzed or collected 2 days post-transfection.

#### Cellular bioenergetics

Cellular respiration was measured using a Seahorse XF24 analyzer (Agilent), as previously published^37^. Transfected brown pre-adipocytes were replated in the XF24 plates at a density of 30,000 cells per well using trypsin. Measurements were obtained before and after the sequential injection of 0.75 µM oligomycin, 0.75 µM FCCP, and 0.75 µM rotenone/myxothiazol. Results were normalized to total protein. Maximal respiration was determined after FCCP injection. Coupled respiration corresponded to the oligomycin response.

#### Immunoblot analysis

Cells were lysed in 10 mM Tris pH 7.5, 10 mM NaCl, 1 mM EDTA and 0.5% Triton X-100, supplemented with complete mini EDTA-free protease (Roche Diagnostics) and phosphatase (Cocktail 2 and 3, Sigma) inhibitors, followed by 10 second sonication. Protein lysates were separated by SDS-PAGE (4-12% Bis-Tris, Invitrogen) and transferred to a nitrocellulose membrane. Transfer was confirmed by Ponceau staining (P7170, Sigma). After blocking in 5% milk, 0.1% Tween-20 in Tris-buffered saline (TBS), primary antibody was incubated overnight at 4°C in 5% bovine serum albumin and 0.1% Tween-20 in TBS. Primary antibodies against PTBP1 (gift from Douglas L. Black, UCLA) and electron transport chain protein complexes (Total OXPHOS rodent WB antibody cocktail ab110413, Abcam) were used at 1:2000. Peroxidase goat anti-rabbit (sc-2030, Santa Cruz Biotechnology, Inc) or rabbit anti-mouse (A9044, Sigma) secondary antibody was used at a 1:10,000 dilution for 1 hour at room temperature in 5% milk and 0.1% Tween-20 in TBS. Immunoreactive bands were revealed with ECL Prime (Amersham) and visualized with a Bio-Rad Gel-doc imager.

#### Statistical analyses

Statistical analyses were performed by an unpaired two-tailed Student’s *t*-test. A value of *p* < 0.05 was considered significant.

## Data availability

The metadata of analyzed datasets are available in SFMetaDB (http://SFMetaDB.yubiolab.org). The analyzed datasets are listed in Data Citations^81–158^, and the processed splicing and gene expression signature data from this study are available at Figshare ^156^.

## Code availability

The scripts and source codes of raw DAS and DEG analyses are deployed in Docker image and deposited at Docker Hub with the public tag sfrs/dasdegdocker:latest^157^. The docker image for signature comparison analysis workflow also was deposited at Docker Hub with the public tag sfrs/sfsigdb:latest ^158^.

## Acknowledgements

The authors would like to thank Mike Zwick and Ben Isett at the Emory Integrated Genomics Core (EIGC), Nolwenn Joffin and Philipp E. Scherer at the University of Texas Southwestern Medical Center for assistance with high-throughput sequencing. The authors would also like to thank Wan Ying Leong for generating and determining efficiencies of shCirbp lentiviral constructs and Douglas L. Black for the primary antibodies against PTBP1. The research was supported partly by the Eunice Kennedy Shriver National Institute of Child Health and Human Development of the National Institutes of Health under award number R01HD037109 to C.C.M. The funders had no role in study design, data collection and analysis, decision to publish, or preparation of the manuscript.

## Author contributions

P.Y., J.L., and S.P.D. carried out the analyses and wrote the manuscript. F.Z., P.N.G., E.W.M.C., S.D.M., L.V., M.S.I., J.M.L., S.L.M., E.G., C.C.M., and P.J. performed validation experiments. All the authors reviewed and approved the final manuscript.

## Additional information

Competing interests: The authors declare no competing interests.

