## Supplemental figures for "Integrated analysis of a compendium of RNA-Seq datasets for splicing factors"


**Figure S1. The massively increasing number of GEO datasets.** ........ 4
**Figure S2. DAS events and DEGs of RNA-Seq datasets curated for SFMetaDB.** ........ 5
**Figure S3. DAS events of Mecp2 knockout mice.** ........ 7
**Figure S4. Cirbp expression is increased in Mecp2-null mice and was knocked down efficiently in hippocampal neurons from rats.** ........ 8
**Figure S5. Confirmation of splicing changes in Mecp2 knockout mice.** ........ 9
**Figure S6. DEGs of Mecp2 knockout mice.** ........ 10
**Figure S7. DAS events of sWAT.** ........ 11
**Figure S8. DEGs of BAT and sWAT.** ........ 12
**Figure S9. Srsf1 knockdown is efficient in 3T3-L1 adipocytes.** ........ 13
**Figure S10. ECARs of glycolytic flux experiments with Srsf1 knockdown.** 14
**Figure S11. Effect of Ptbp1 knockdown in immortalized brown adipocytes.** ........ 15
**Figure S12. Protein-protein associations among MSFs for CIT from the STRING database.** ........ 16
**Figure S13. Similarity among the splicing signatures and among the gene expression signatures of SRRM4 and PRMT5 from different tissue/cell types.** ........ 18
**Figure S14. A hypothetic example of splicing signature comparison analysis.** ........ 20

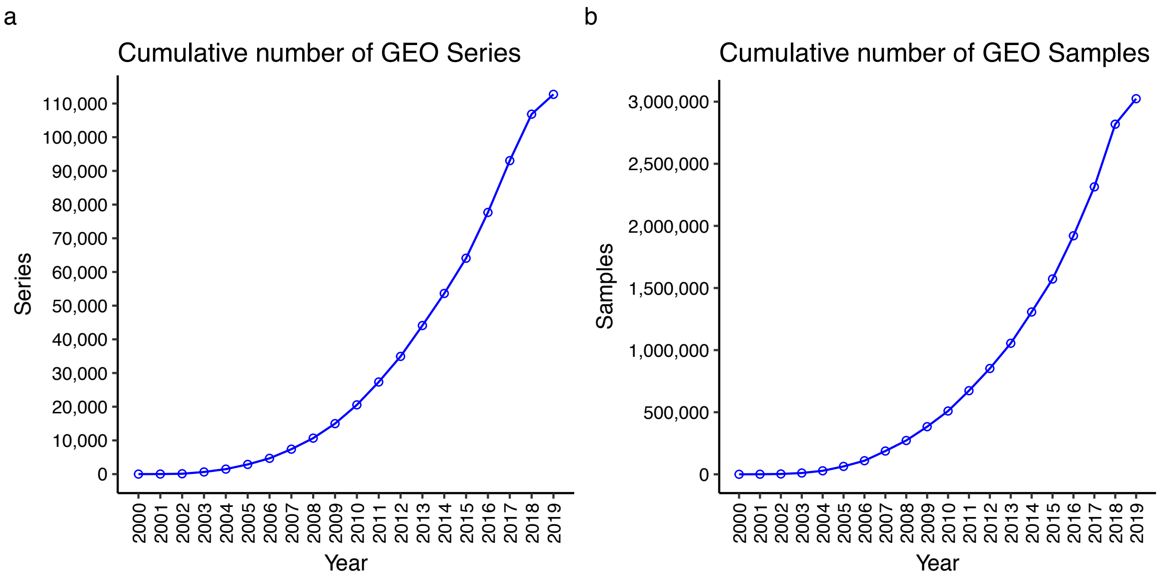


**Figure S1. The massively increasing number of GEO datasets.**

To evaluate the trend of generation of public biological datasets, the number of GEO data was extracted from “GEO summary-history.” From year 2000 to the second quarter of 2019, the points depict an increasing trend of generation of public biological datasets. (a) The cumulative number of GEO series. (b) The cumulative number of GEO samples.

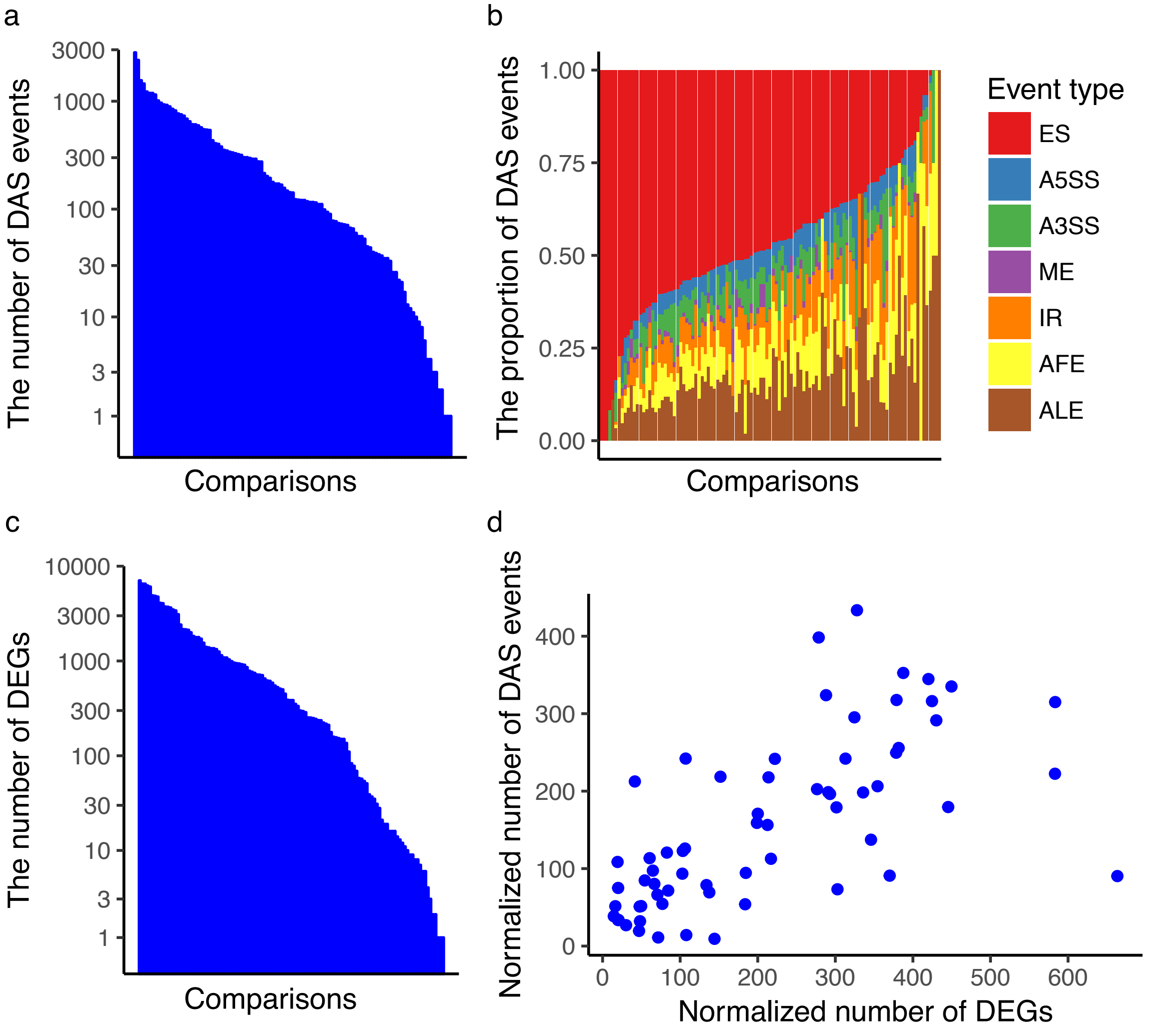


**Figure S2. DAS events and DEGs of RNA-Seq datasets curated for SFMetaDB.** (a) The number of DAS events in the RNA-Seq datasets. A total of 111 comparisons have nonzero DAS events. Each bar represents a comparison derived from an RNA-Seq dataset, and comparisons are ordered along the x-axis by the number of DAS events. (b) The proportion of seven common DAS event types in the analysis results. The types are exon skipping (ES), alternative 5′ splice sites (A5SSs), alternative 3′ splice sites (A3SSs), mutually exclusive (ME) exons, intron retention (IR), alternative first exons (AFEs), and alternative last exons (ALEs). The colored bars at a given position in the x-axis represent the proportions of the seven DAS events. (c) The number of DEGs in the RNA-Seq datasets. A total of 120 comparisons have nonzero DEGs. The comparisons are ordered along the x-axis by the number of DEGs. (d) The scatterplot of the normalized number of DEGs and the normalized number of DAS events in the high-quality comparisons derived from the RNA-Seq datasets curated for SFMetaDB (where the total number of reads is $\geq 100$ million spots and the effective read length is $\geq 150$ bps). Each point represents a comparison in a dataset.

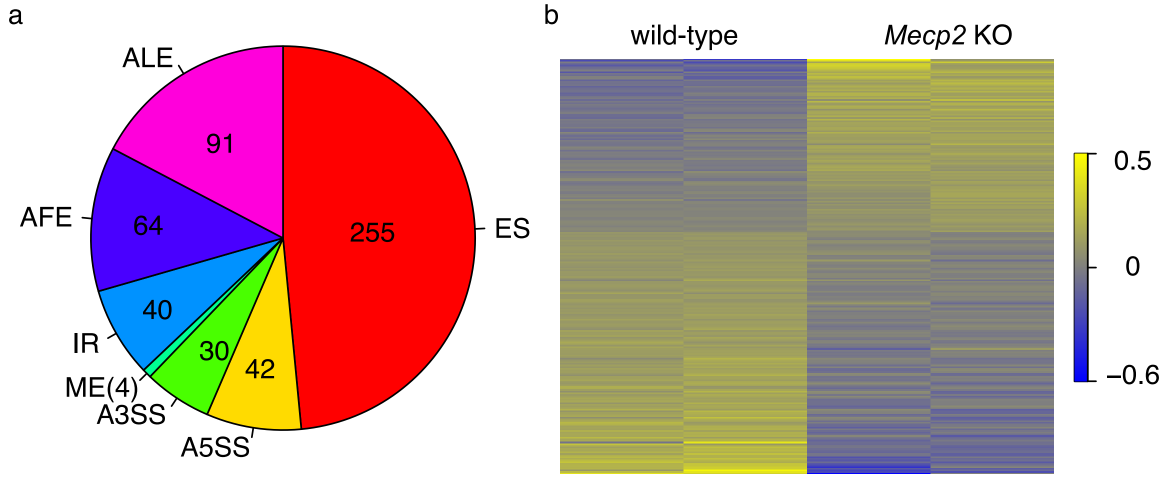


**Figure S3. DAS events of *Mecp2* knockout mice.**

DAS analysis was performed on *Mecp2* knockout mice. The seven alternative splicing event types consist of exon skipping (ES), alternative 5' splice sites (A5SSs), alternative 3' splice sites (A3SSs), mutually exclusive (ME) exons , intron retention (IR), alternative first exons (AFEs), and alternative last exons (ALEs). (a) The pie chart depicts the number of DAS events of the seven splicing event types. (b) The heatmap shows the PSI values (scaled by standard deviation) for the differential alternative ES events in *Mecp2* knockout mice. Yellow: high PSI value; blue: low PSI value.

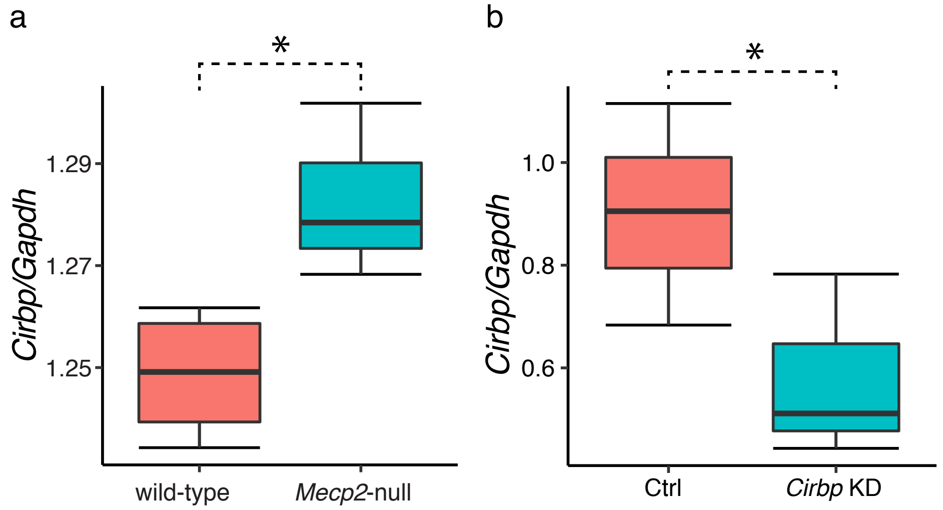


**Figure S4. *Cirbp* expression was increased in *Mecp2*-null mice and was knocked down efficiently in hippocampal neurons from rats.**

(a) The expression level of *Cirbp* (normalized to *Gapdh*, a housekeeping gene) in *Mecp2*-null mice was increased compared to wild-type mice according to qRT-PCR analysis. (One-way ANOVA test. *:$p$-value < 0.05, $n$ = 3 in wild-type and $n$ = 5 in *Mecp2*-null.) (b) The expression level of *Cirbp* (normalized to *Gapdh*, a housekeeping gene) in *Cirbp* knocked-down neurons was significantly reduced compared to control rats according to qRT-PCR analysis. (One way ANOVA test. *:$p$-value < 0.05, $n$ = 3 in each group.)

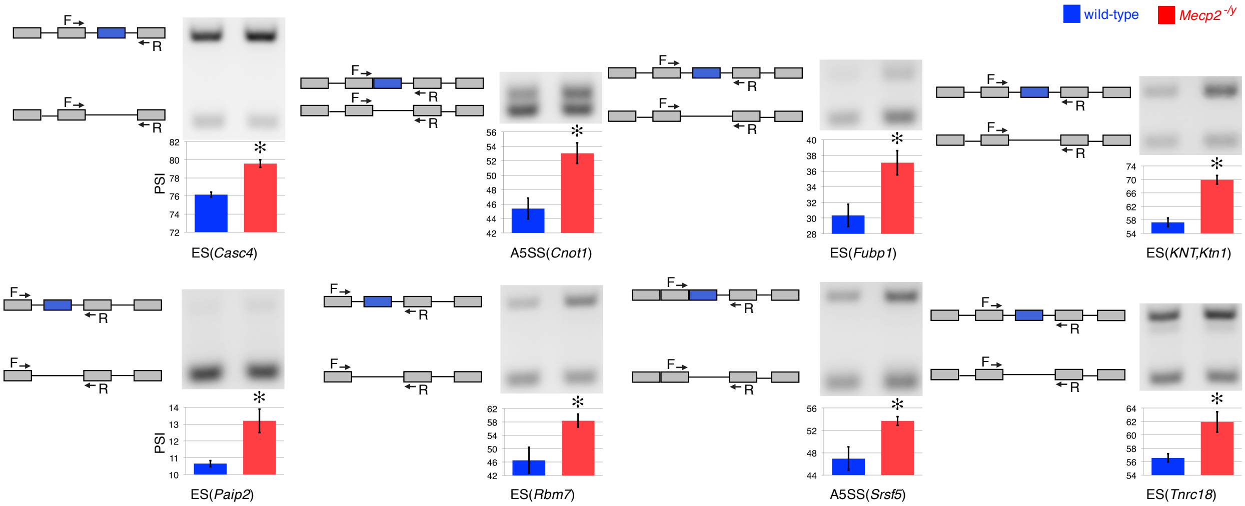


**Figure S5. Confirmation of splicing changes in *Mecp2* knockout mice.** RT-PCR was performed on the selected differential alternative splicing events to confirm splicing changes in the dentate gyrus of *Mecp2* knockout mice. The confirmed DAS events were of two types: ES and A5SS events. Agarose gel electrophoresis of PCR products for splicing events were obtained in wild-type and *Mecp2*$^{-/y}$ mice. Gene models of DAS events representing forward and reverse primer sequences are associated with gel images. The band sizes are shown to the right of the gel images. PSI values calculated from the scanned gel are shown under each lane. The bar graph depicts the PSI value (y-axis) of each event in *Mecp2* knockout (red bar) compared to wild-type (blue bar) mice. Asterisks (*) indicate significant splicing changes between 5 *Mecp2* knockout and 3 wild-type male mice ($p < 0.05$ by a one-tailed $t$-test).

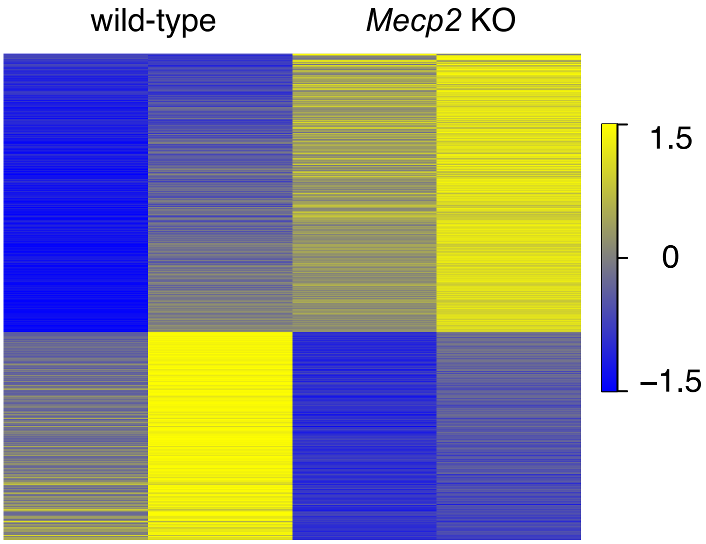


**Figure S4. DEGs of *Mecp2* knockout mice.**

DEG analysis was performed in the dentate gyrus of *Mecp2* knockout and wild-type mice. The heatmap shows the expression levels scaled by standard deviation for each gene in *Mecp2* knockout mice. Yellow: high expression level; blue: low expression level.

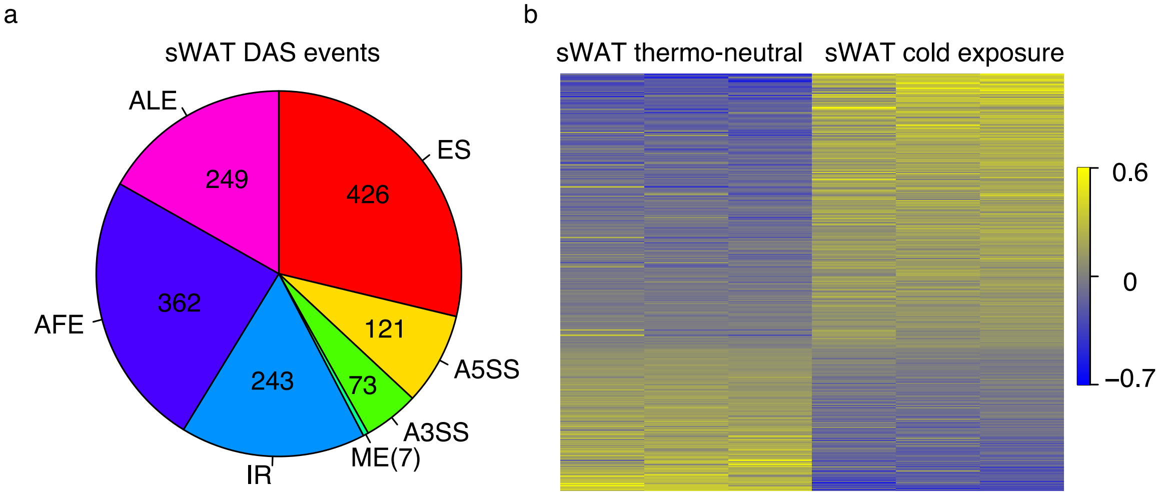


**Figure S5. DAS events of sWAT.**

(a) DAS analysis identified seven alternative splicing event types, i.e., ES, A5SS, A3SS, ME, IR, AFE, and ALE. The pie chart depicts the number of DAS events of the seven splicing event types in sWAT. (b) The heatmaps show the PSI values (scaled by standard deviation) for the differential alternative ES events in sWAT. Yellow: high PSI value; blue: low PSI value.

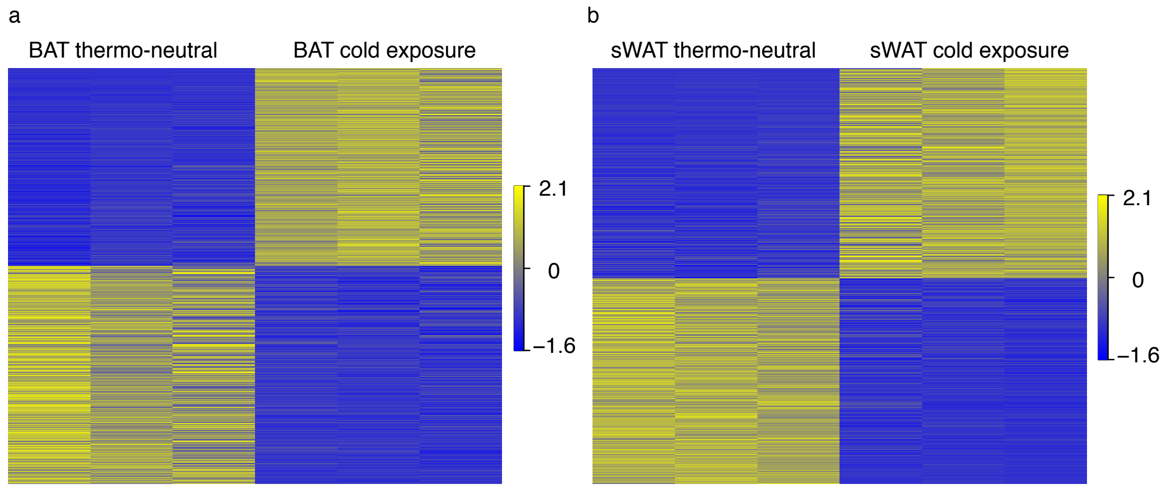


**Figure S6. DEGs of BAT and sWAT.**

DEG analysis was performed on BAT and sWAT derived from cold-exposed mice. The heatmaps show the expression levels scaled by standard deviation for each gene in BAT and sWAT. Yellow: high expression level; blue: low expression level. (a) The expression levels of DEGs in BAT. (b) The expression levels of DEGs in sWAT.

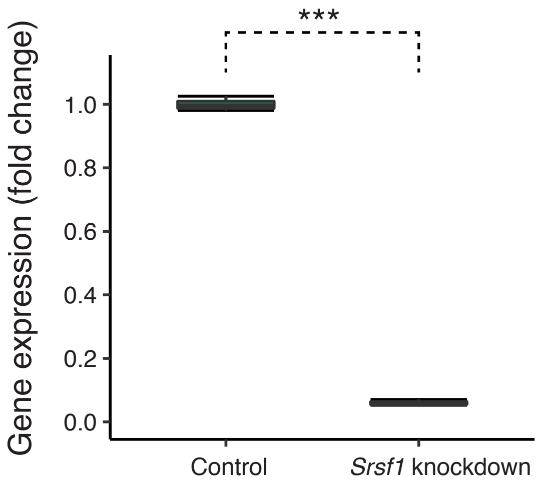


**Figure S7. *Srsf1* knockdown is efficient in 3T3-L1 adipocytes.**

The normalized expression level of *Srsf1* to cyclophilin (a housekeeping gene) in *Srsf1* knockdown adipocytes was reduced significantly compared to control cells according to RT-PCR analysis. (Unpaired *t*-test. ***:$p$-value $< 0.0001$, $n = 3$ in each group.)

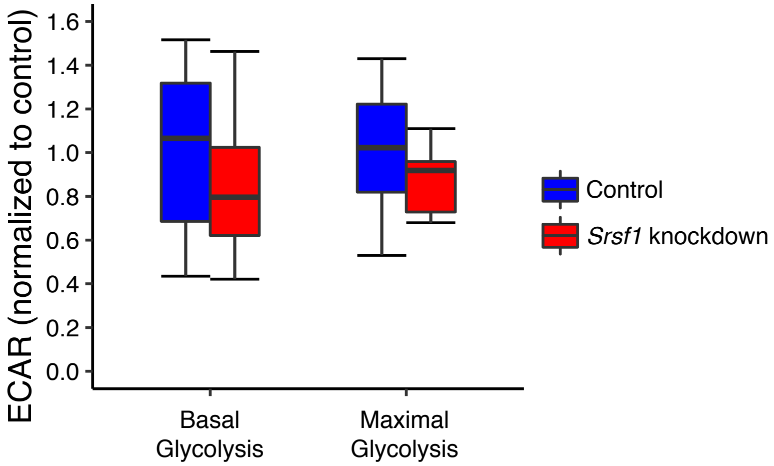


**Figure S8. ECARs of glycolytic flux experiments with *Srsf1* knockdown.**
ECARs were measured for *Srsf1* knockdown adipocytes (red boxes) and controls (blue boxes). There was no significant difference between *Srsf1* knockdown adipocytes and controls (*n* = 9 to 10 biological replicates in each group).

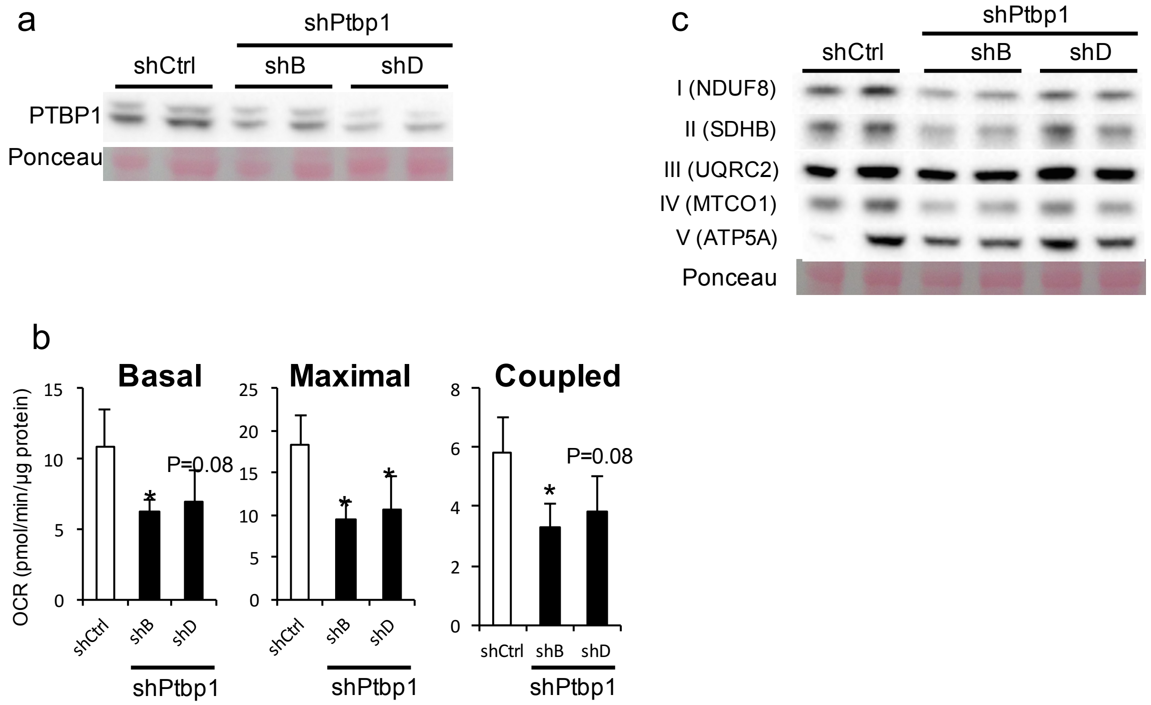


**Figure S9. Effect of *Ptbp1* knockdown in immortalized brown adipocytes.** Cells were transfected with shRNA vectors containing scrambled negative control (shCtrl) or *Ptbp1*-specific shRNA (shB and shD), and analyzed 2 days later. (a) PTBP1 detection by western blot. Ponceau staining represents a loading control. (b) Cellular bioenergetics. Cellular respiration was measured with a Seahorse XF24 analyzer (n = 4 in each group). (c) Electron transport chain protein complexes detected by western blot. Ponceau staining represents a loading control.

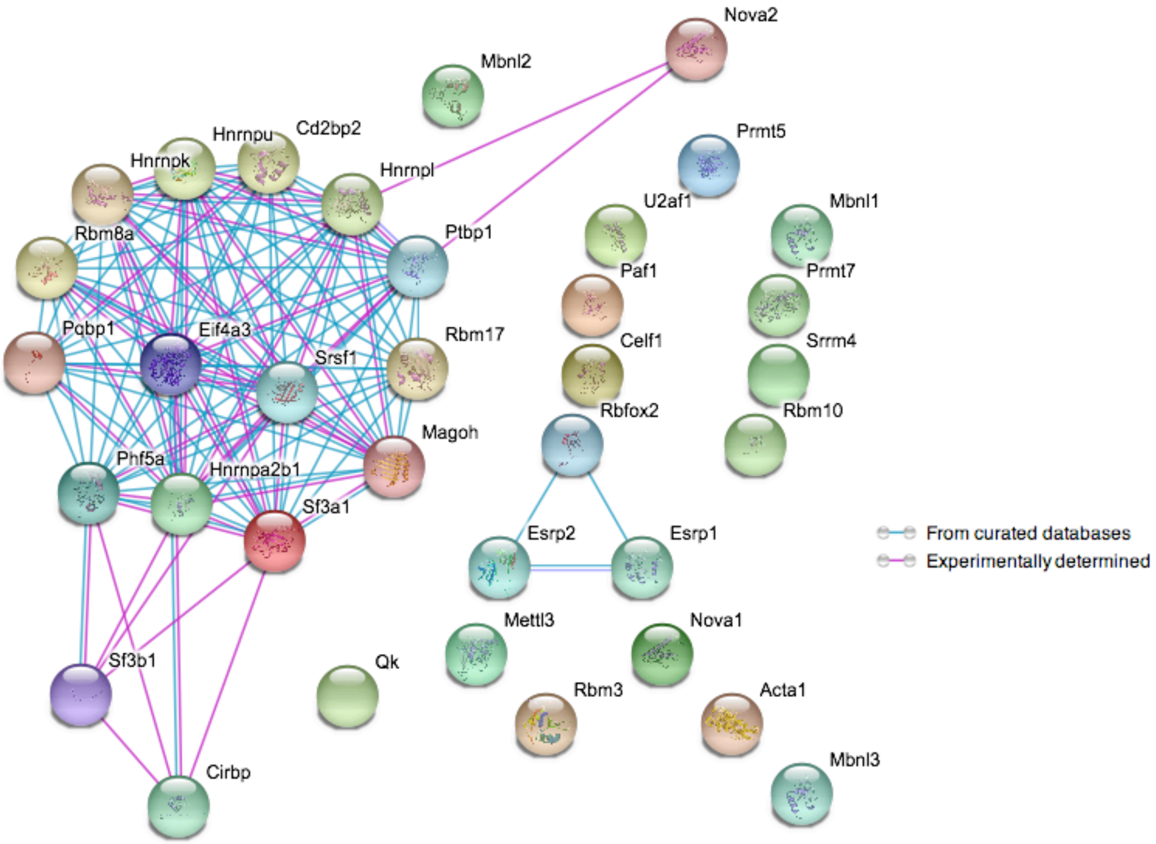


**Figure S10. Protein-protein associations among MSFs for CIT from the STRING database.**

To examine the functional relation among the candidate MSFs in CIT, 35 factors were queried in the STRING database for evidence about the protein-protein associations. The nodes with the gene symbols correspond to the candidate MSFs. There are two edge types denoting the two types of protein-protein association evidence from the STRING. The purple edge represents the protein-protein associations supported by the direct experimental evidence, while the light blue edge means that proteins are grouped into metabolic, signaling, or transcriptional pathways. The figure depicts the protein-protein associations for the MSFs identified for CIT.

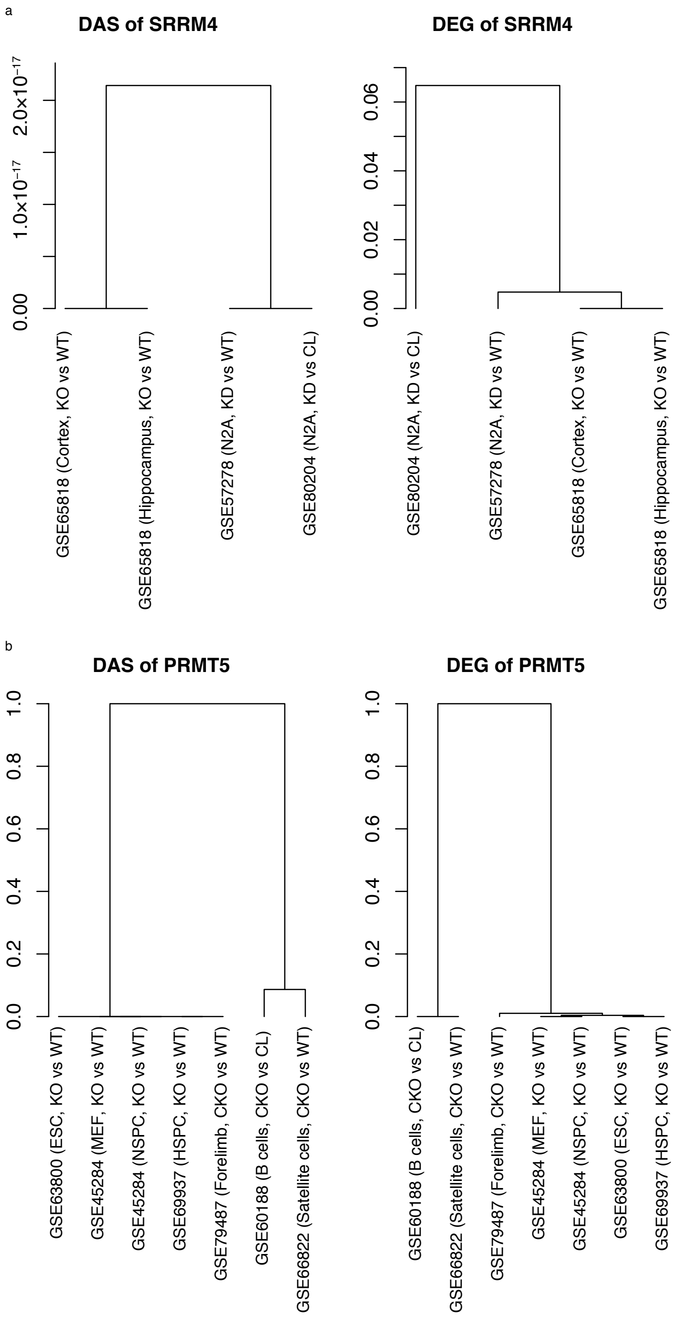


**Figure S11. Similarity among the splicing signatures and among the gene expression signatures of SRRM4 and PRMT5 from different tissue/cell types.**

To evaluate the similarity among SRRM4 and PRMT5 signatures from different tissue/cell types, Fisher's exact test was used to calculate the significance of the overlap between each pair of splicing and gene expression signatures. The resultant *p*-values were used as distances to cluster the signatures using complete-linkage hierarchical clustering. The dendrograms show that (a) SRRM4 has similar splicing and gene expression signatures among comparisons, while (b) PRMT5 has two groups of splicing and gene expression signatures from different tissues. Acronyms in the figure are as follows—CKO: conditional knockout; KO: knockout; KD: knockdown; WT: wild-type; CL: control; N2A: neuroblastoma cell line; ESC: embryonic stem cells; MEF: mouse embryonic fibroblasts; NSPC: neural stem and progenitor cells; HSPC: hematopoietic stem and progenitor cells.

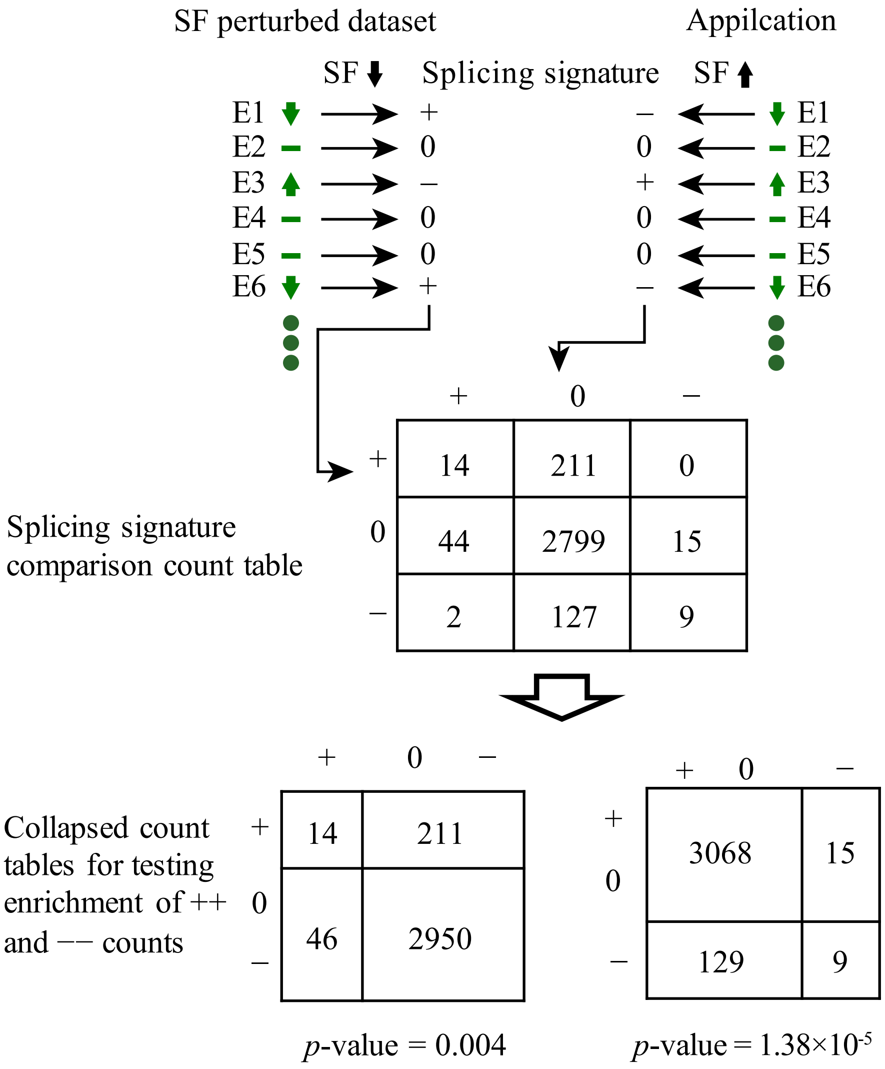


**Figure S12. A hypothetic example of splicing signature comparison analysis.** To illustrate the splicing signature comparison workflow, a hypothetic SF splicing signature was used to compare to a signature from an application. In the SF perturbed dataset, the inclusion and exclusion of DAS events, and the nonsignificantly-changed events are indicated with up arrows, down arrows, and horizontal bars. The potential regulatory direction of the DAS events by the SF then can be determined. In this example, the down arrows and up arrows of DAS events were potentially positively regulated by the SF (denoted by +) and potentially negatively regulated by the SF (denoted by $-$), respectively, because the expression of the SF is down. The rest of the events had no evidence of regulation by the SF (denoted by 0). This vector of $+/-/0$ values was the splicing signature of the SF perturbed dataset. Similarly, a signature also was calculated for an application dataset. A $3\times3$ count table was tabulated by counting the number of pairs of $+/-/0$ events in two signatures. The $3\times3$ count table was collapsed further into two $2\times2$ count tables to test the enrichment of $++$ events (left matrix) and $--$ events (right matrix). Fisher's exact test *p*-values were 0.004 and $1.38\times10^{-5}$ for the two tables, respectively. The $+-$ events and $-+$ events in the antidiagonal corners were not tested because they represent opposite regulation directions in the SF perturbed dataset and the application, leading to no interpretable regulation of the SF in the application.
