## Supplementary material for "Integrated analysis of a compendium of RNA-Seq datasets for splicing factors": Table 1

**Online-only Table 1. Evidence for identified SFs in CIT from the literature.**

| **Gene symbol** | **Relevance to processes related to thermogenesis** | **Dir^a^** | **PMID** |
| --- | --- | --- | --- |
| *Celf1* | *Celf1* represses the activity of *Ppargc1a* mRNA by binding to its 3’UTR in primary brown preadipocytes. As *Ppargc1a* has been demonstrated to be a master regulator of the UCP1-mediated thermogenesis in BAT based on an *in vivo* experiment using *Ppargc1a* knockout mice, BAT thermogenesis can be activated by inhibiting *Celf1* ^1^. | S | 28763438 |
| *Hnrnpu* | The siRNAs knockdown of *Hnrnpu* in differentiating brown adipocytes significantly impairs lipid droplet accumulation and decreases the gene expression of BAT markers including *Ucp1*, *Dio2*, *Ppargc1a*, *Prdm16*, *Pparg,* and *Cox8b*, indicating that *Hnrnpu* is needed for brown adipogenesis and is critical for thermogenesis in BAT ^2^. | NA | 25921091 |
| *Nova1* | *Nova1* can function as a brown-adipogenic repressor. Transient overexpression of FLAG-tagged NOVA1 protein reduces the gene expression of brown adipocyte‒specific markers *Prdm16* and *Ucp1* compared to the controls ^3^. In contrast, shRNA-induced knockdown of *Nova1* increases *Prdm16* and *Ucp1* transcript levels in the differentiating cells. *Nova1*-overexpressing cells exhibit less lipid accumulation compared to controls. These results have identified the repressive effect of *Nova1* on the brown adipocyte development and metabolism. Therefore, *Nova1* may affect thermogenesis in BAT. | S | 26857472 |
| *Pqbp1* | The shRNA knockdown of *Pqbp1* reduces lipid storage in mammalian white adipocyte, indicating that *Pqbp1* is critical for thermogenesis ^4^. | NA | 19119319 |
| *Prmt5* | *Prmt5* proliferates the activation of adipogenic gene expression and promotes adipogenic differentiation. *Prmt5* is required for adipogenesis via an *in vitro* experiment in preadipocytes, indicating that *Prmt5* is critical for thermogenesis ^5^. | E | 22361822 |
| *Hnrnpk* | Overexpression of hnRNP K protein modulates insulin-activated mitochondrial gene expression, and mitochondrial biogenesis and remodeling are inherent to adipose differentiation *per se* and are influenced by the actions of insulin sensitizers, suggesting the potential role of *Hnrnpk* in CIT ^6^. | NA | 16519889 |
| *Mettl3* | Knockdown of METTL3 increases lipid accumulation during adipogenesis ^7^. | NA | 25412662 |

^a^The column “Regulation direction of thermogenesis (Dir)” indicates whether an MSF enhances or suppresses thermogenesis based on mRNA-level changes of markers of relevant processes related to thermogenesis, such as adipogenesis and lipid accumulation. The “Enhancing (E)” regulation direction represents increased mRNA levels of markers given the overexpression of the MSF or decreased levels under knockdown of the MSF. On the contrary, the “Suppressive (S)” regulation direction means decreased mRNA levels of markers given the overexpression of the MSF or increased levels under knockdown of the MSF. “NA” represents that the regulation direction cannot be determined because mRNA levels of markers are not evaluated in the reference.
