## Supplementary material for "Integrated analysis of a compendium of RNA-Seq datasets for splicing factors": Data S3

**Data S3. Primer sequences for RT-PCR validations**

NM_001205370.1

Casc4	Sequence (5'->3')	Template strand	Length

>Casc4F	CTTGGATCGGGAACCCAGAA	Plus	20

>Casc4R	ATCACTGAAGCGCTGTTTGC	Minus	20

NM_001205226.1

Cnot1	Sequence (5'->3')	Template strand	Length

>Cnot1F	AGCATTCTCTGCGTTTGTTGG	Plus	21

>Cnot1R	ATCATGGTAGGGTTAGCCGC	Minus	20

NM_057172.3

Fubp1	Sequence (5'->3')	Template strand	Length

>Fubp1F	GGGGGTGATGCTGGTACATC	Plus	20

>Fubp1R	TGGGGAGGTACTTTCTTAGCATC	Minus	23

NM_001039129.4

Hnrnpa1	Sequence (5'->3')	Template strand	Length

>Hnrnpa1F	AGTAGCTATGGCAGTGGCAGG	Plus	21

>Hnrnpa1R	TCTGTTGTAACCTGTAGCTTCCC	Minus	23

Hnrnpa1	Sequence (5'->3')	Template strand	Length

>Hnrnpa1F	AGTAGCTATGGCAGTGGCAGG	Plus	21

>Hnrnpa1R1	TGGCCCAGAATAAAGGCTGC	Minus	20

NM_001293636.1

Ktn1	Sequence (5'->3')	Template strand	Length

>Ktn1F	GCGAAGTGTGGAGCAAGAAG	Plus	20

>Ktn1R	TCACGTAAGTCGATCGCTCC	Minus	20

NM_026420.2

Paip2	Sequence (5'->3')	Template strand	Length

>Paip2F	GATTCGTTGGCTACCGTCCC	Plus	20

>Paip2R	ATGCTTGGGCTAGTACTGCTG	Minus	21

NM_018814.3

Pcnx	Sequence (5'->3')	Template strand	Length

>PcnxF	CCAAGCCAGGTTGCATTTCC	Plus	20

>PcnxR	CGTCTCATAGAGGGTGGACG	Minus	20

uc009grj.1

Ptov	Sequence (5'->3')	Template strand	Length

>PtovF	CACAGTCCCAGACGAGGC	Plus	18

>PtovR	AACATACAGGGGCAAGGACC	Minus	20

NM_144948.5

RbmF	Sequence (5'->3')	Template strand	Length

>Rbm7F	GGTAACCTGGAGACGAAGGTG	Plus	21

>Rbm7R	AAGTTCACGAATGCAAACTGC	Minus	21

NM_009159.2

Srsf5	Sequence (5'->3')	Template strand	Length

>Srsf5F	ACAGCTCCGTCGCAGACTA	Plus	19

>Srsf5R	TGAACACTCGACAGCCACTC	Minus	20

NM_001122730.2

Tnrc18	Sequence (5'->3')	Template strand	Length

>Tnrc18F	TTGGTCAAGCTCCAGAGACG	Plus	20

>Tnrc18R	GATGAGGAGTCGTGATCGTCC	Minus	21
