## Supplementary figures and images for "Integrated analysis of a compendium of RNA-Seq datasets for splicing factors"

### rbpCirbpMmJunYan_DEG_CVsH.pdf

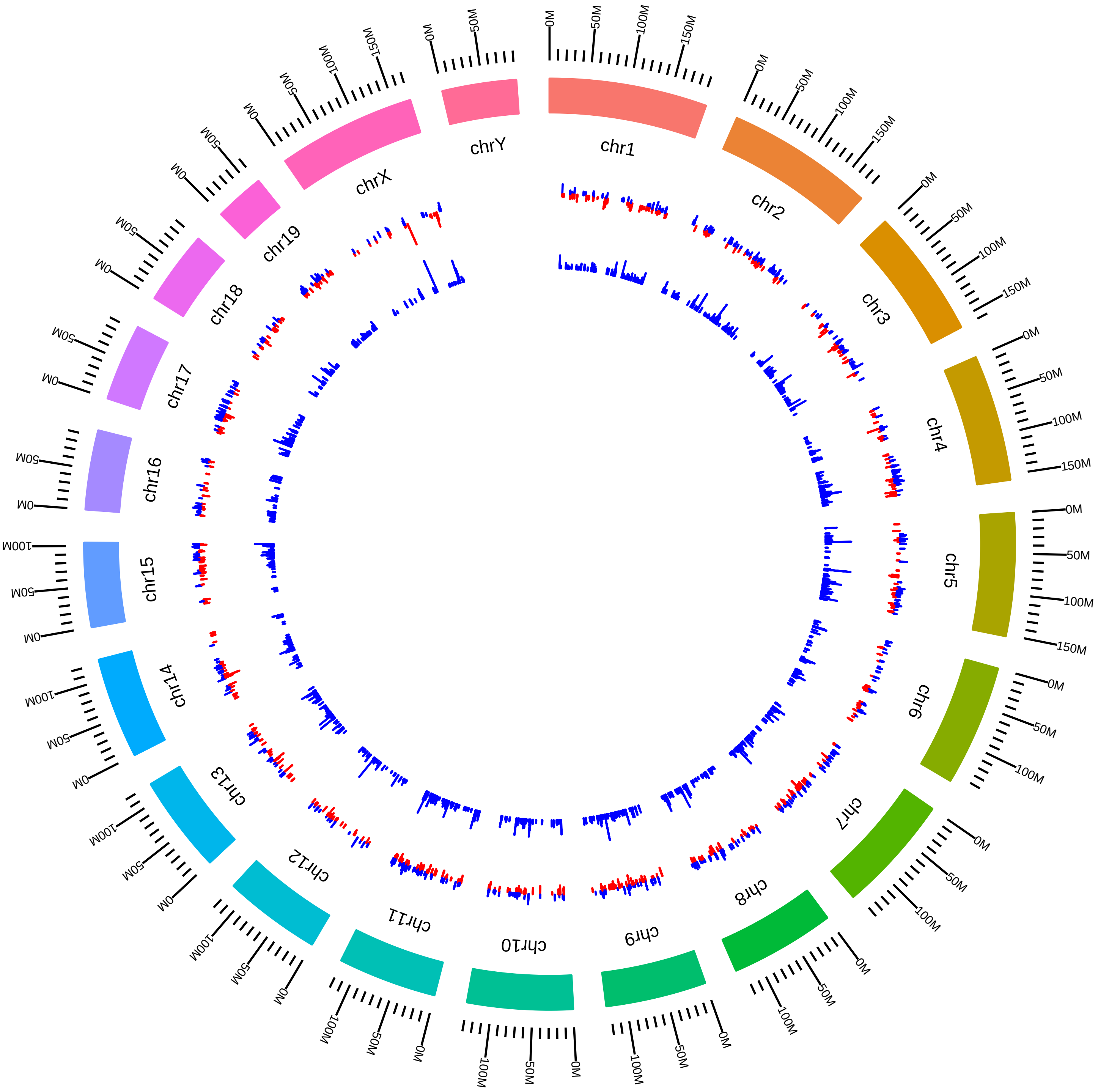

### rbpCirbpMmJunYan_DEG_RVsH.pdf

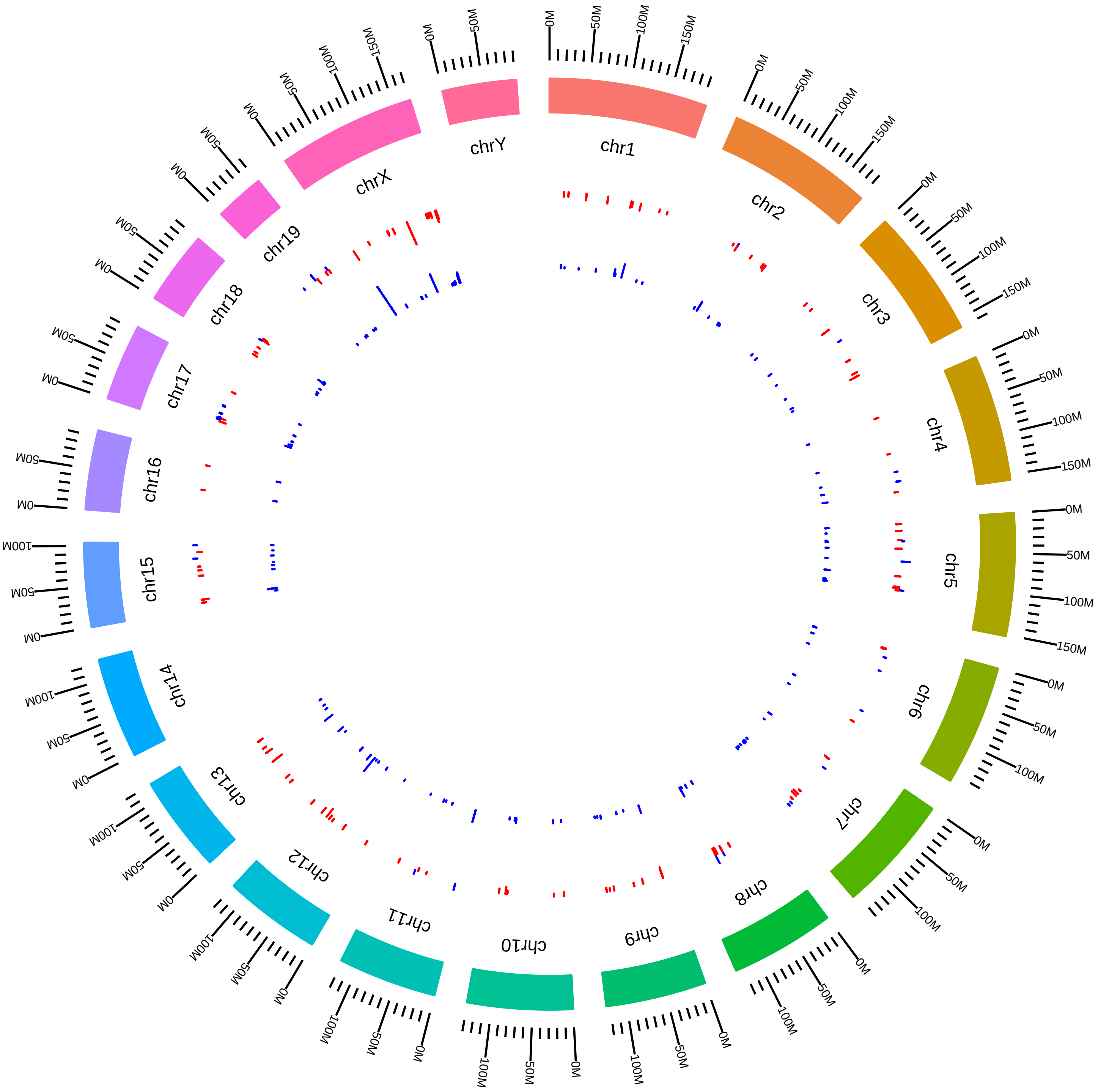

### rsfCd2bp2MmCFreund2015_KVsW.pdf

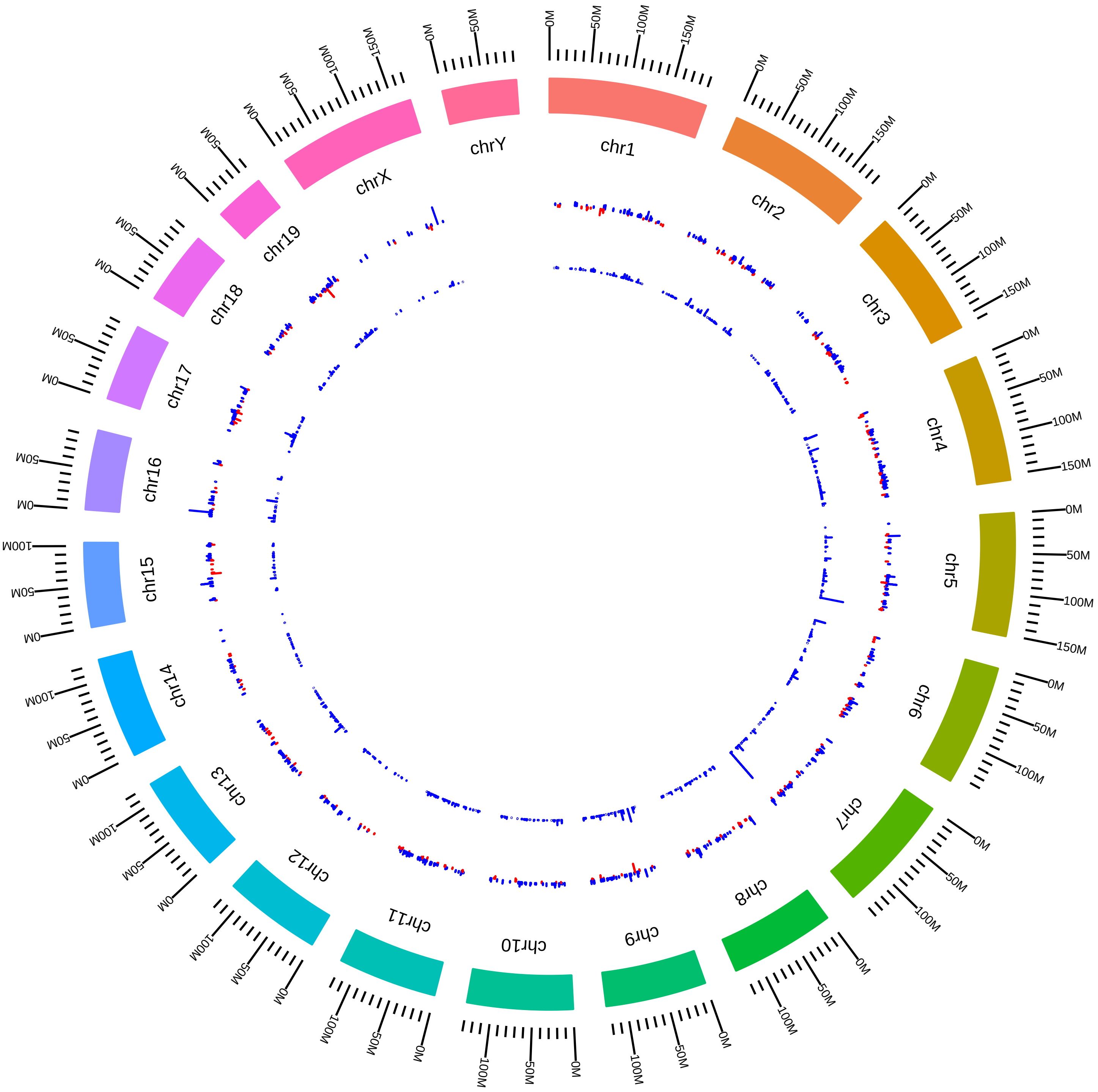

### rsfCelf1MmCBurgeHeart_DEG_LVsS.pdf

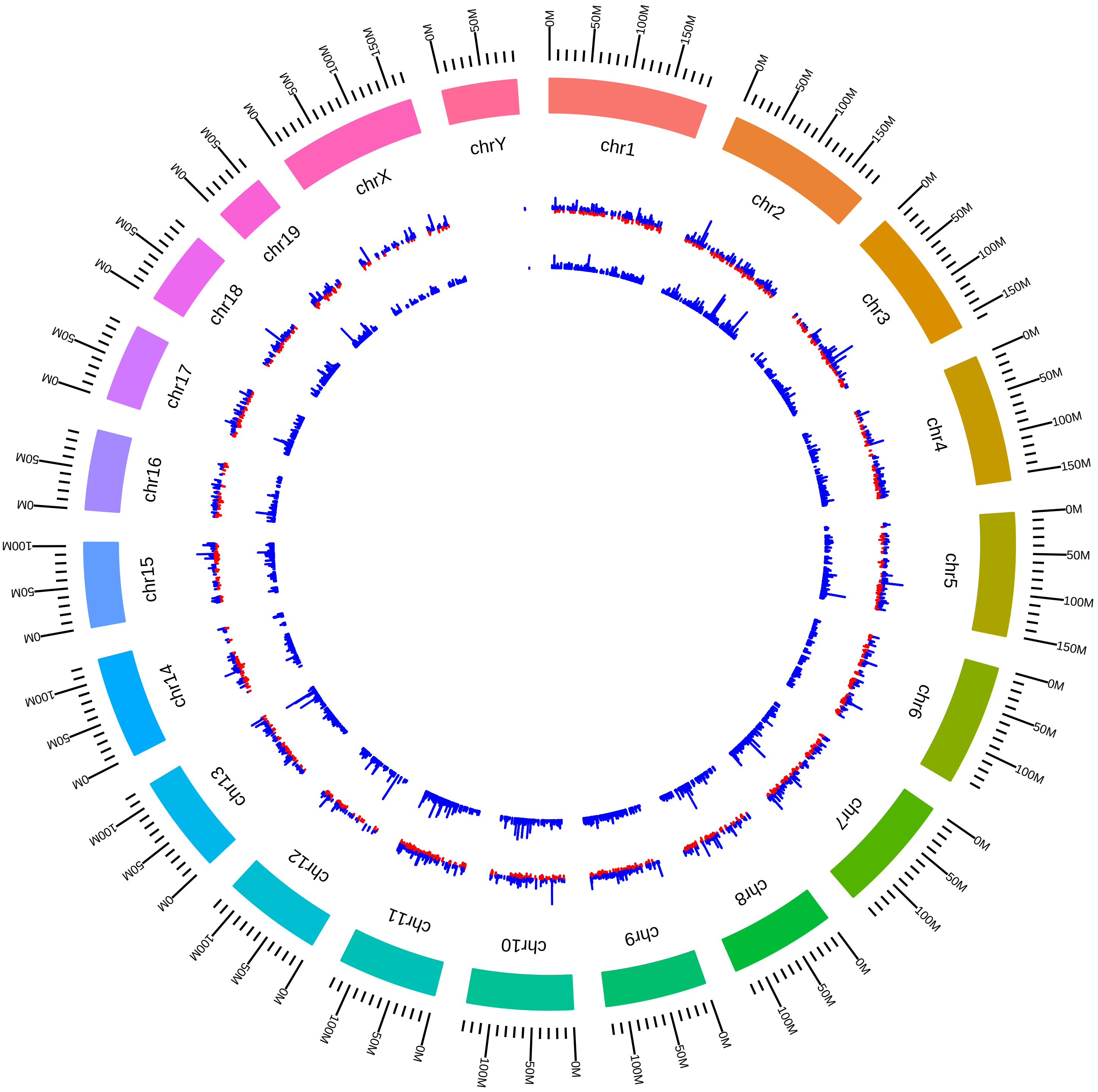

### rsfCelf1MmCBurgeMuscle_DEG_LVsS.pdf

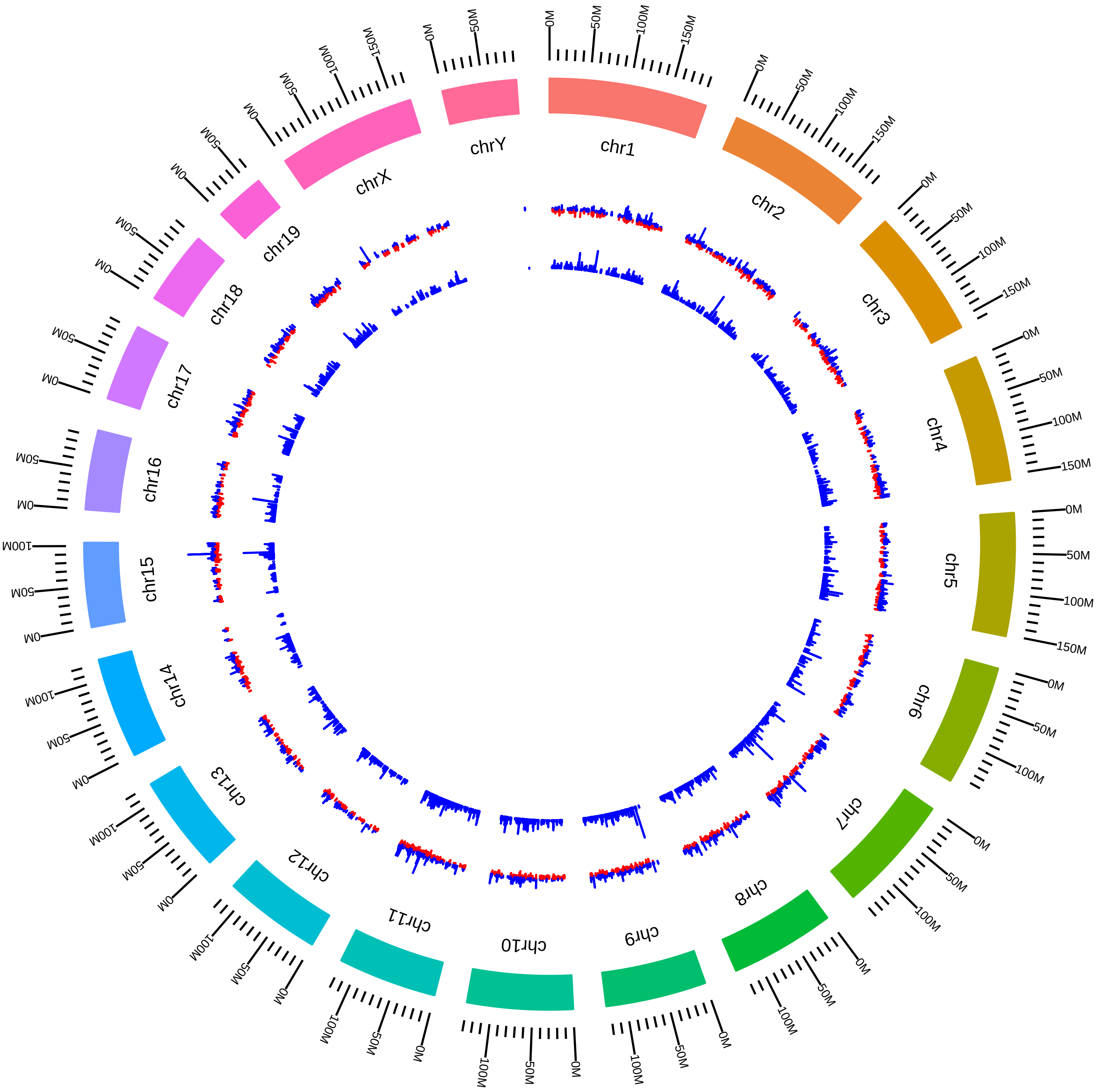

### rsfCelf1MmTCooper_DEG_KVsW.pdf

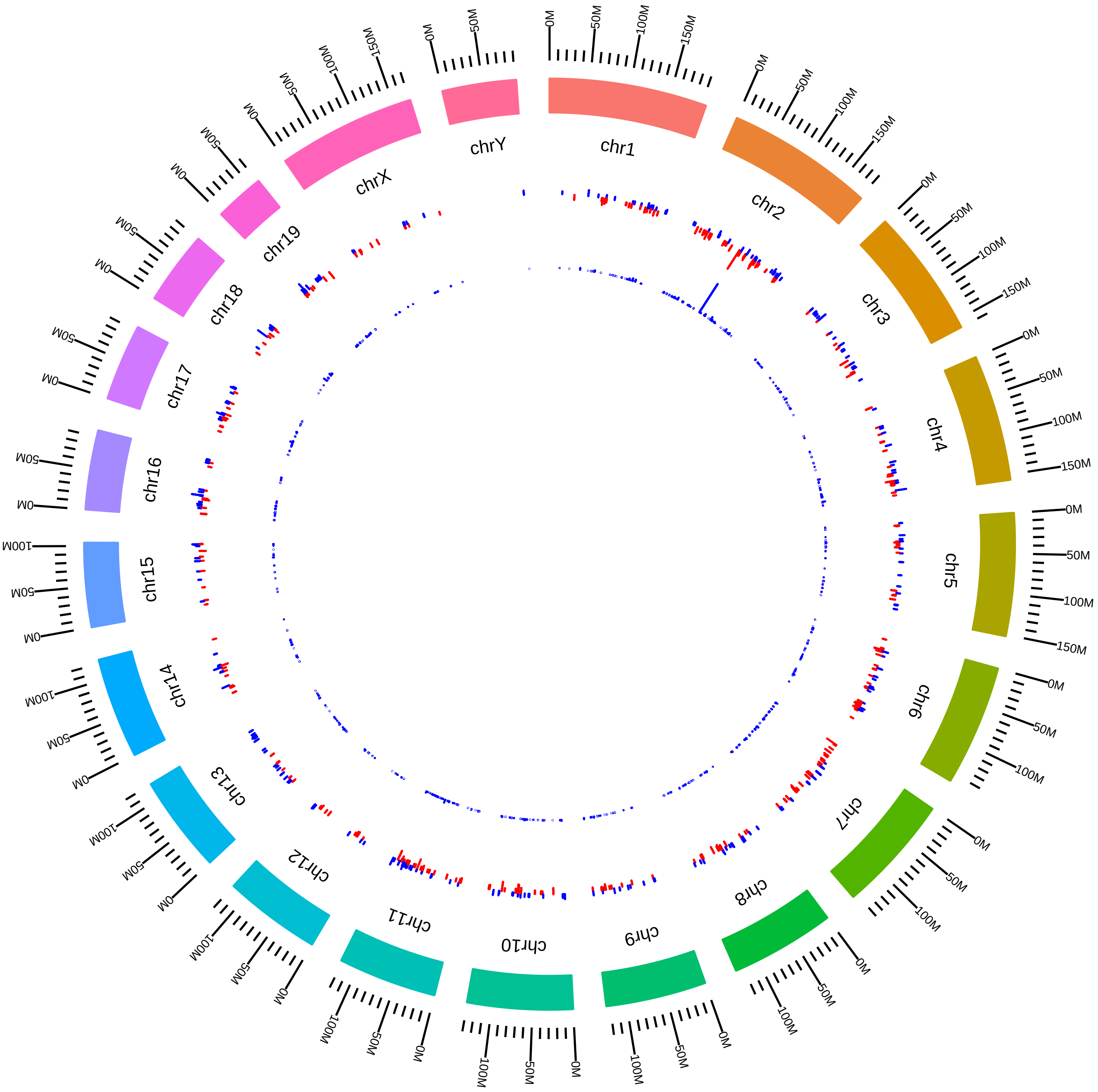

### rsfCelf2MmCBurge2015_DEG_LVsS.pdf

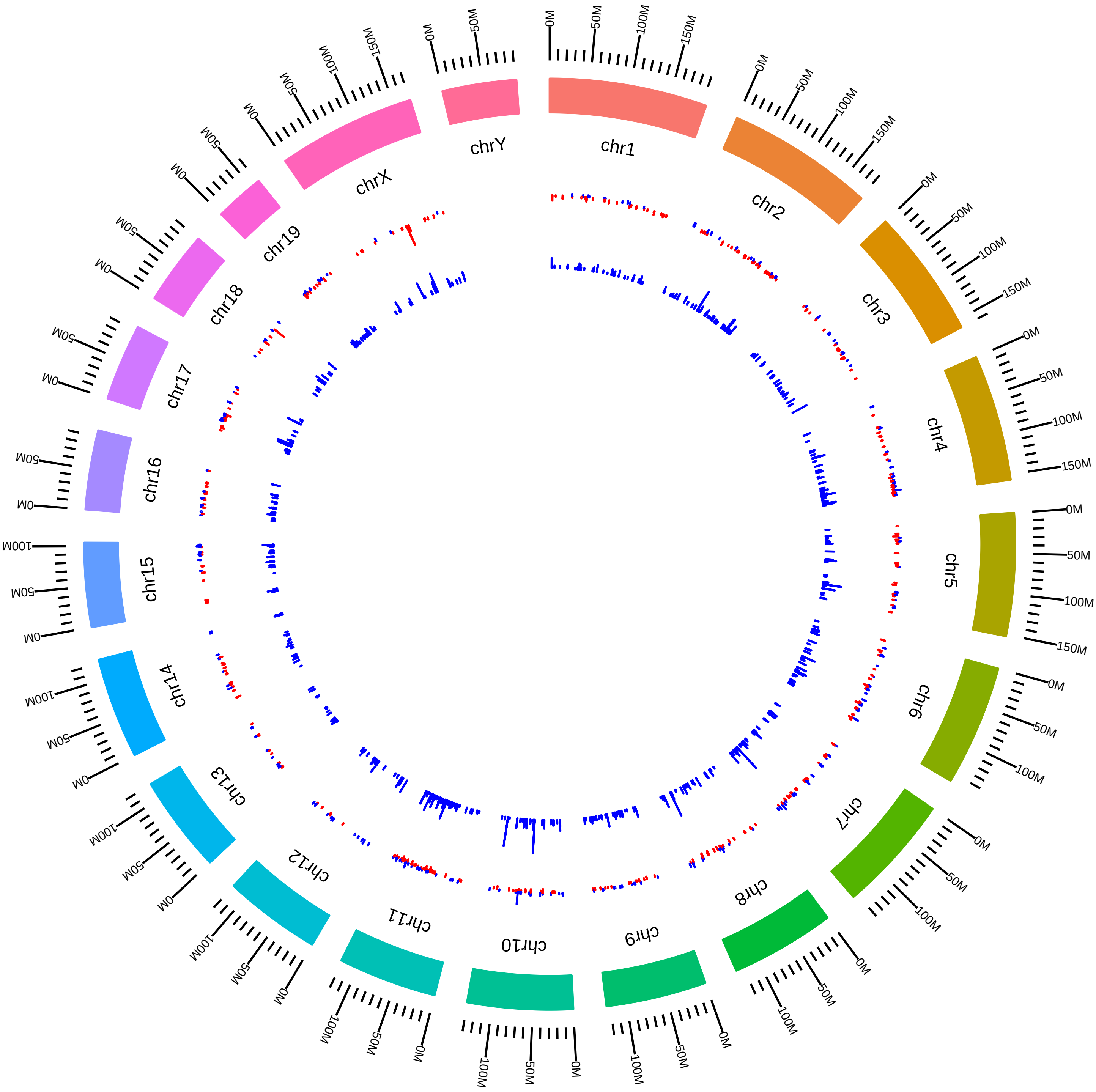

### rsfCirbpRbm3MmJunYan2013_CVsH.pdf

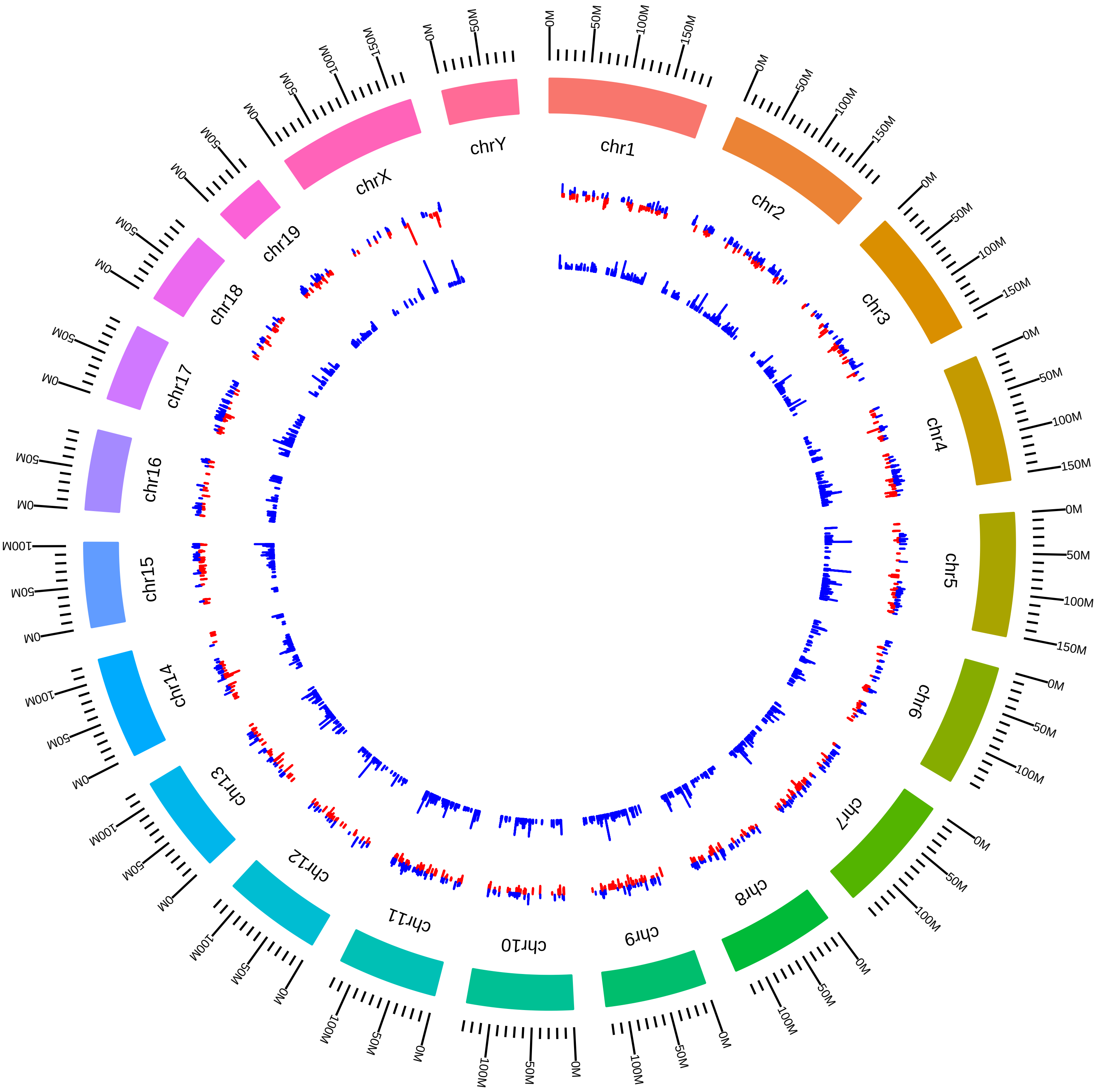

### rsfCirbpRbm3MmJunYan2013_RVsH.pdf

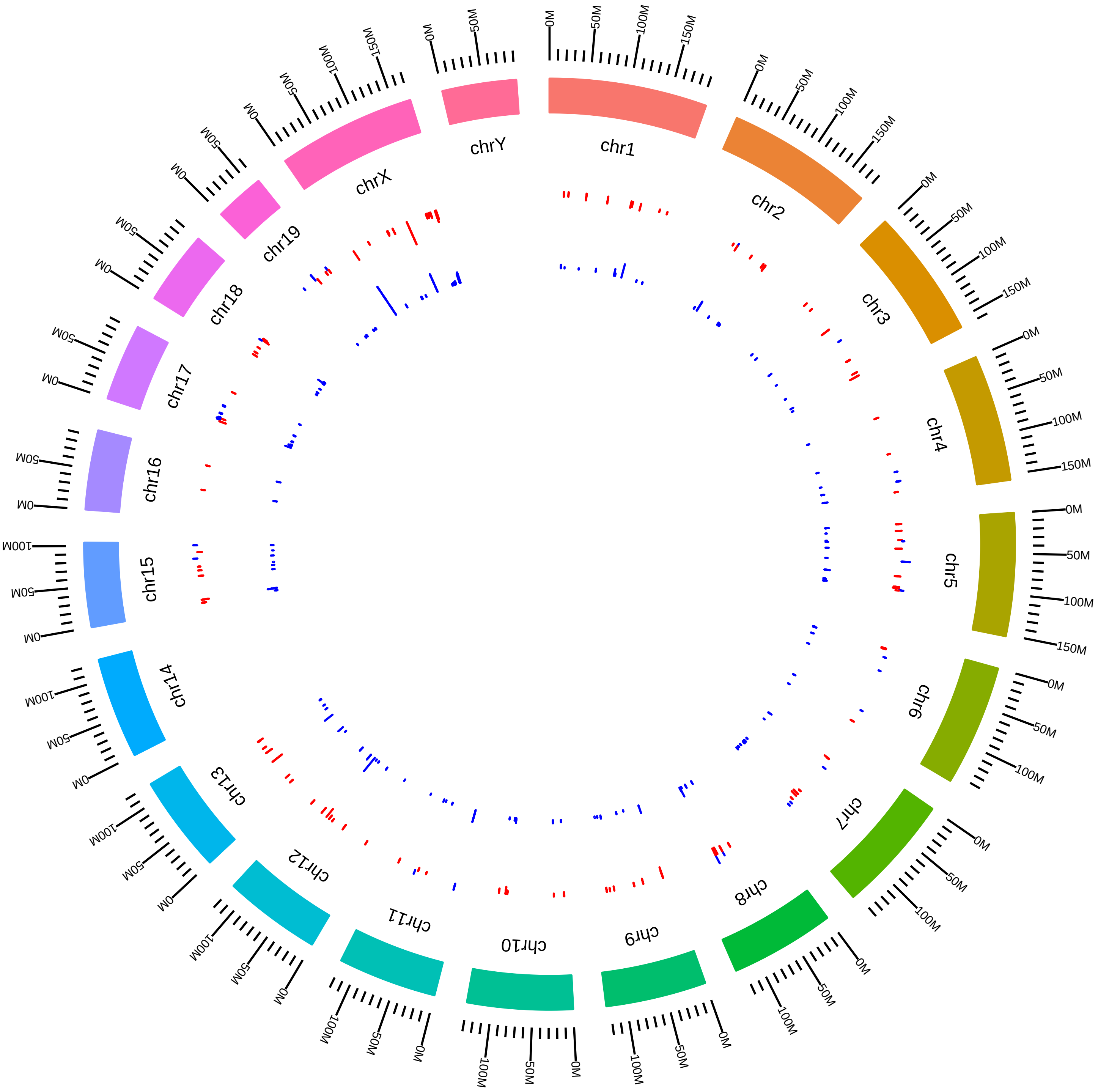

### rsfDdx5MmBStillman2014_DEG_DVsC.pdf

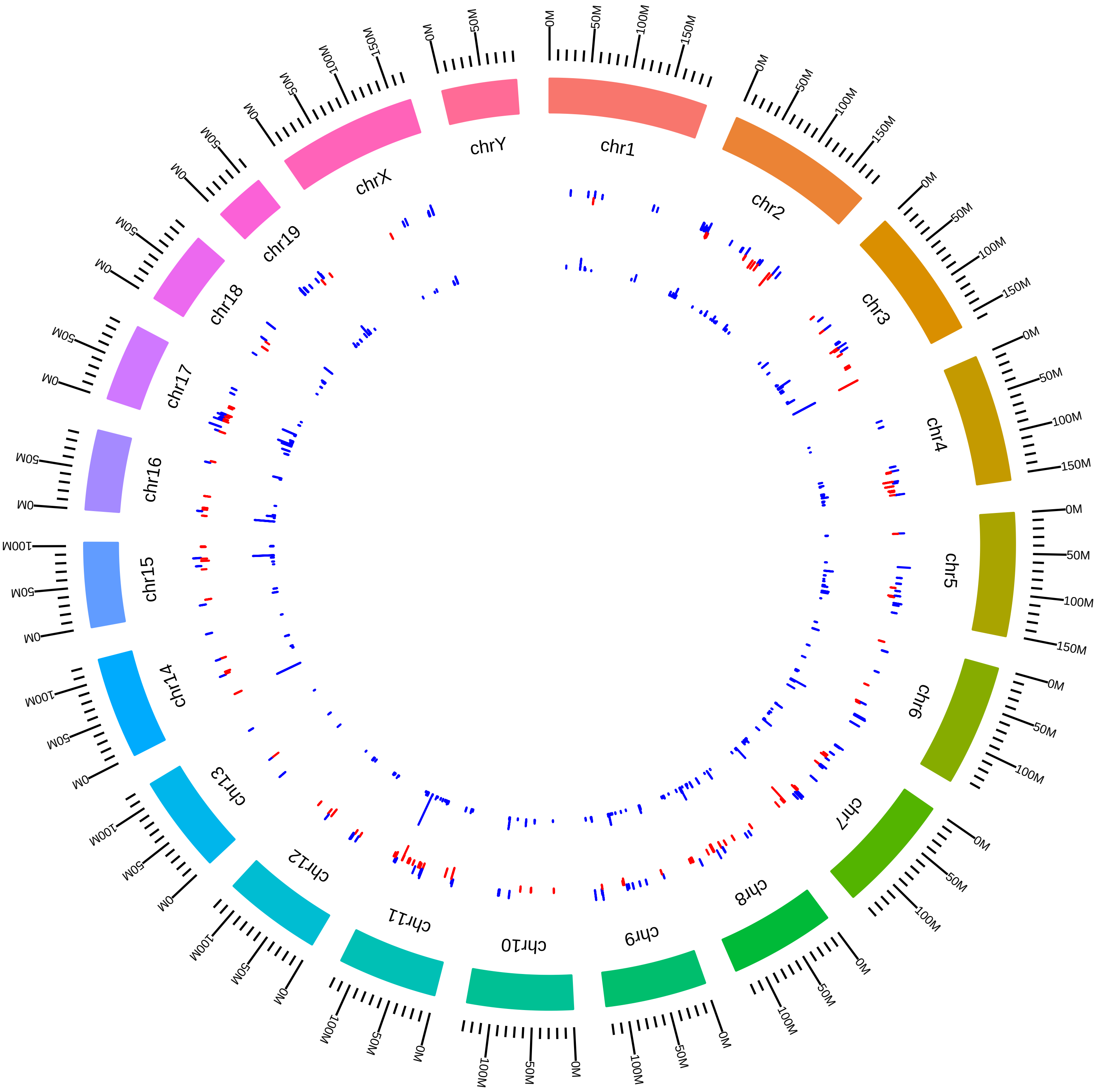

### rsfDdx5MmDRLittman2015_DEG_KVsW.pdf

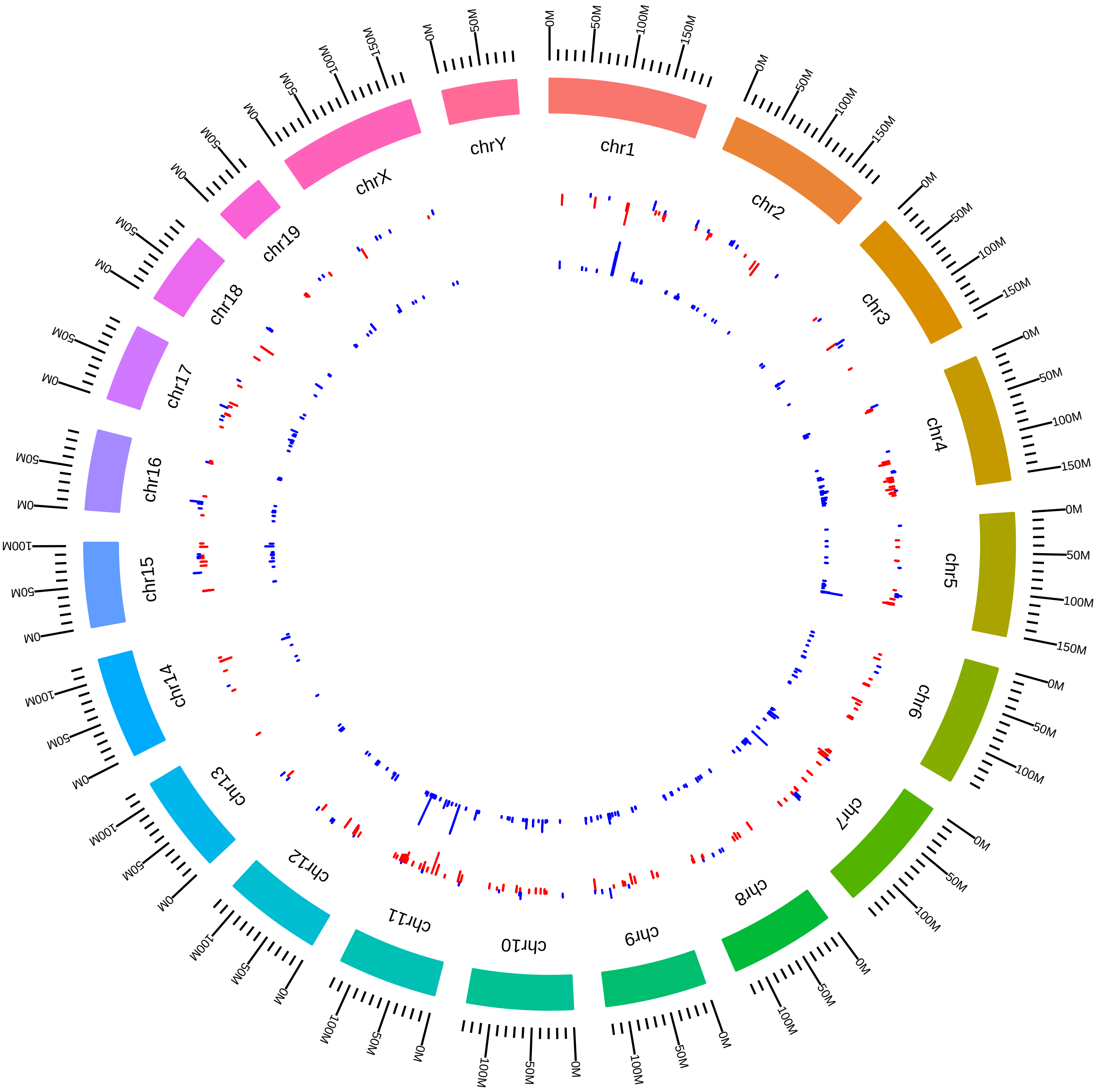

### rsfDdx5MmHongjieYao2017_KVsW.pdf

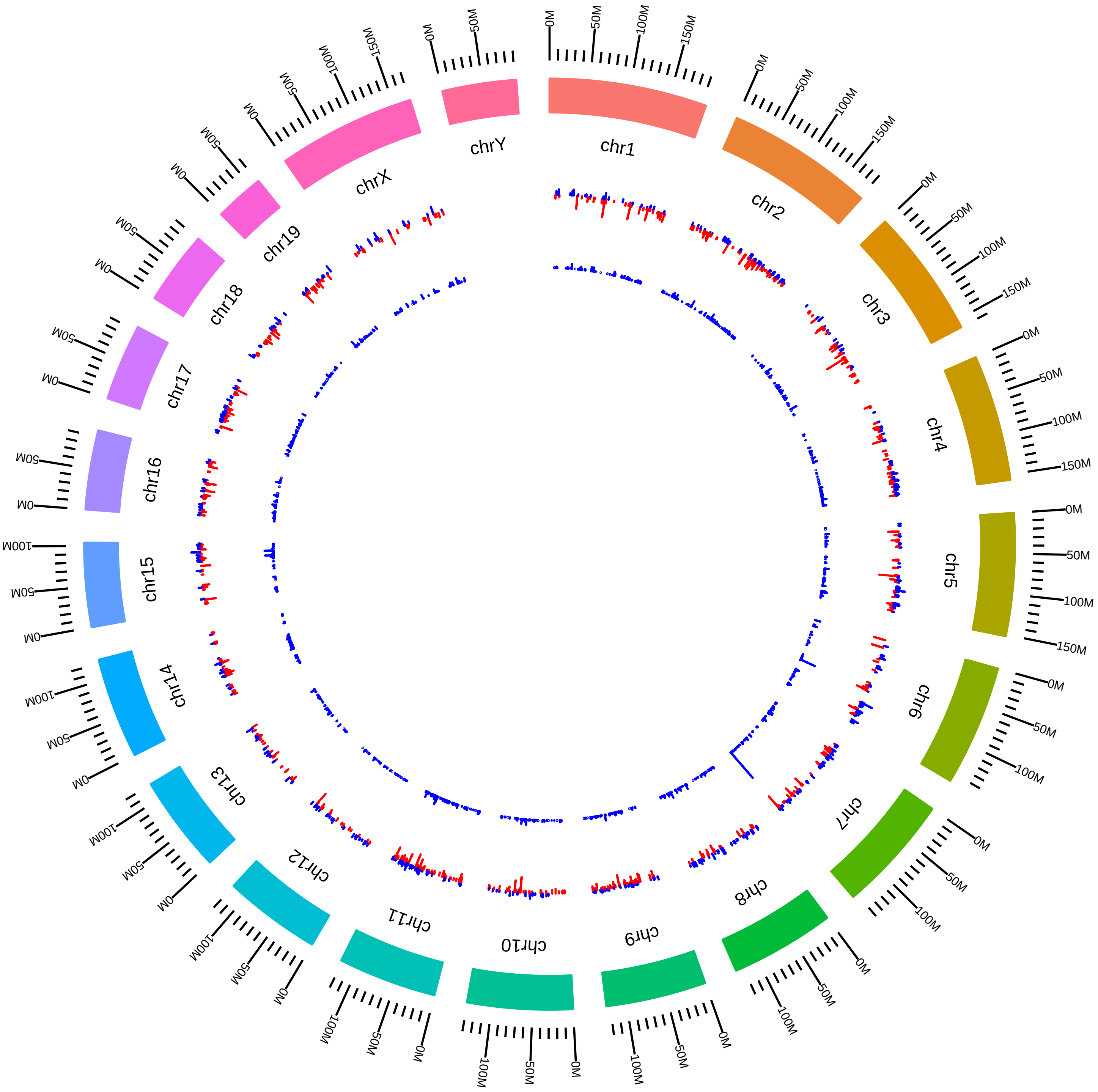

### rsfDyrk1aMmRXavier2015_DVsC.pdf

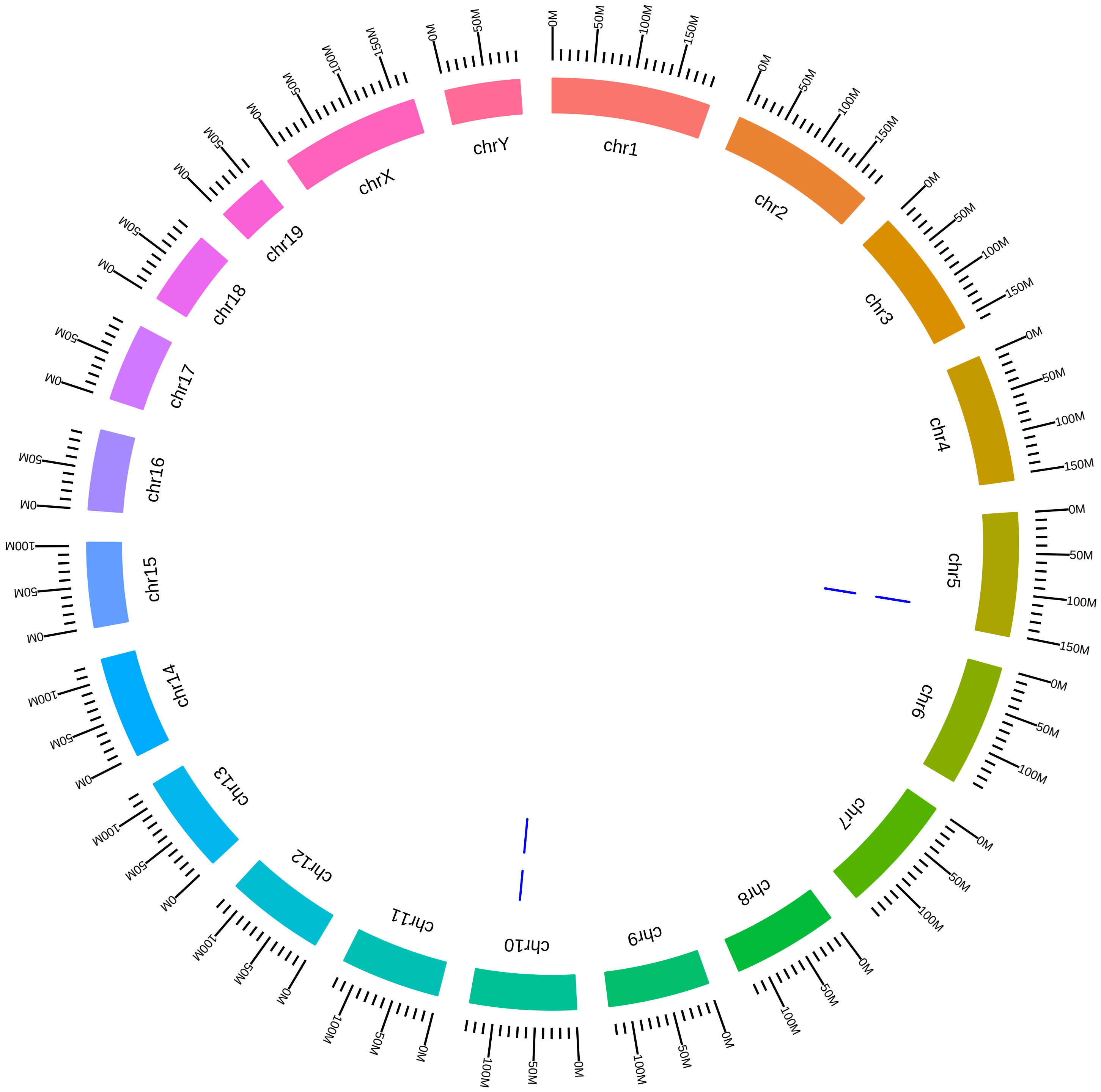

### rsfDyrk1aMmRXavier2015_IVsC.pdf

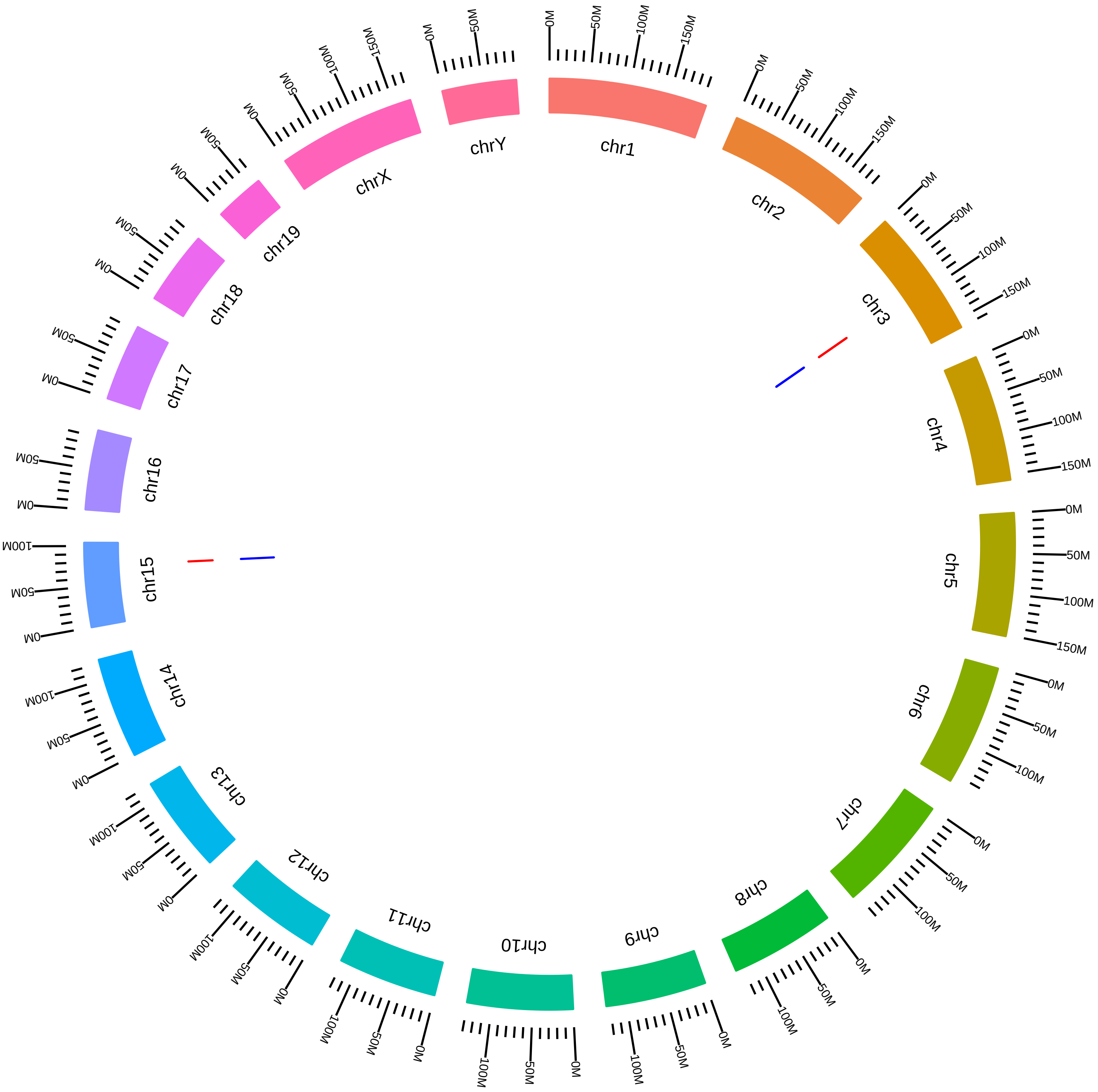

### rsfEif4a3MagohRbm8aMmDSilver2016_EVsC.pdf

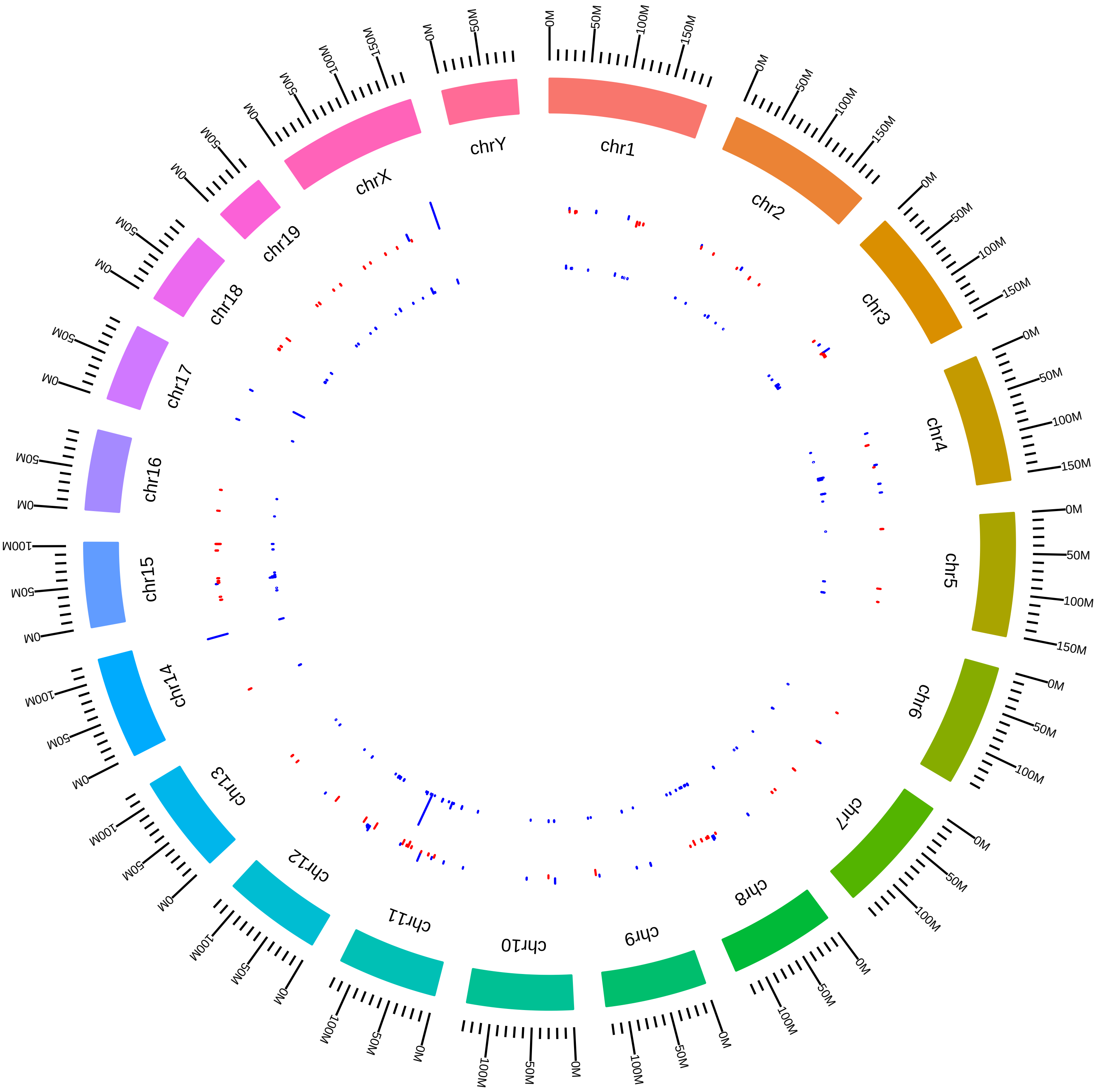

### rsfEif4a3MagohRbm8aMmDSilver2016_MVsC.pdf

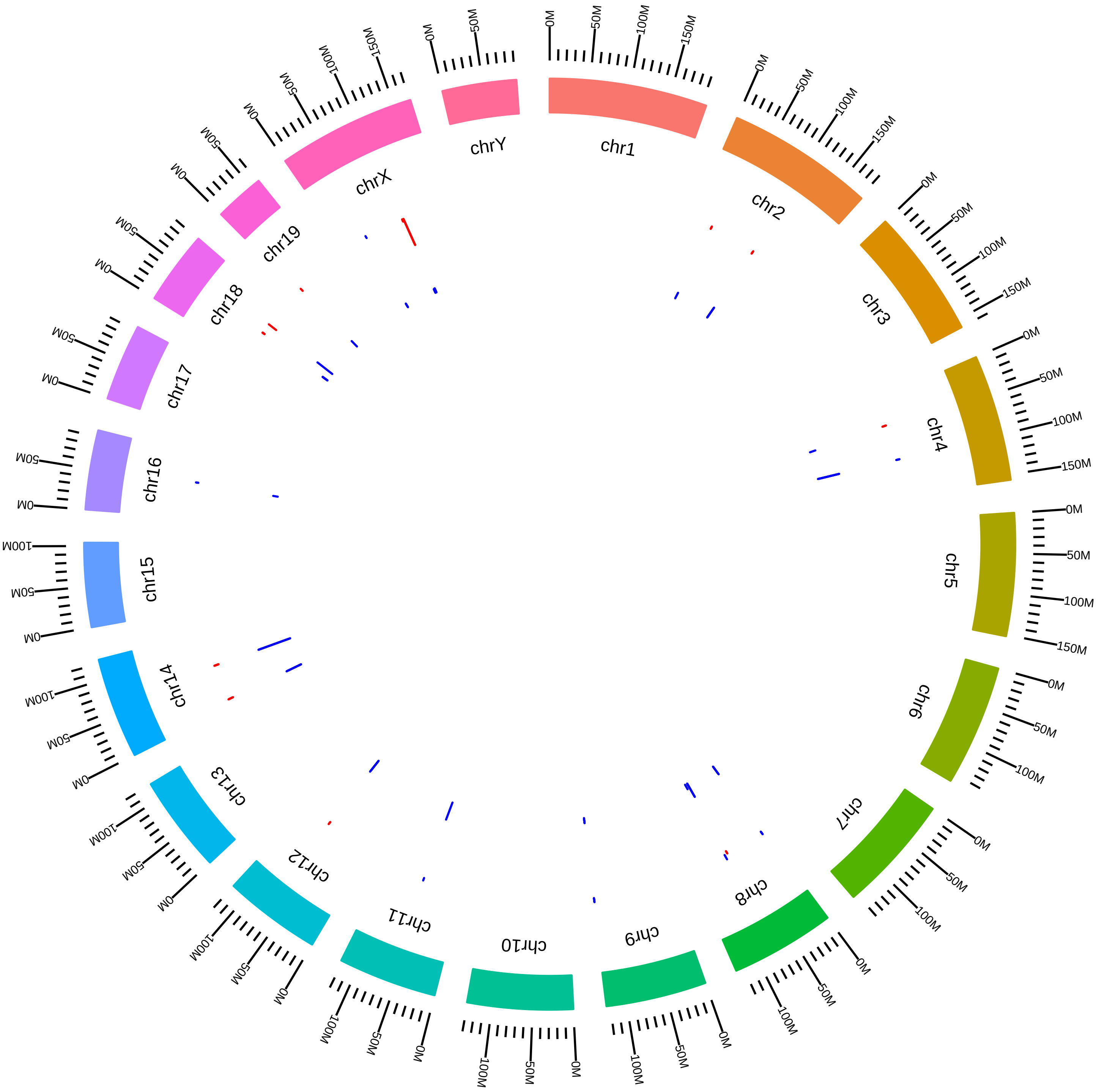

### rsfEif4a3MagohRbm8aMmDSilver2016_RVsC.pdf

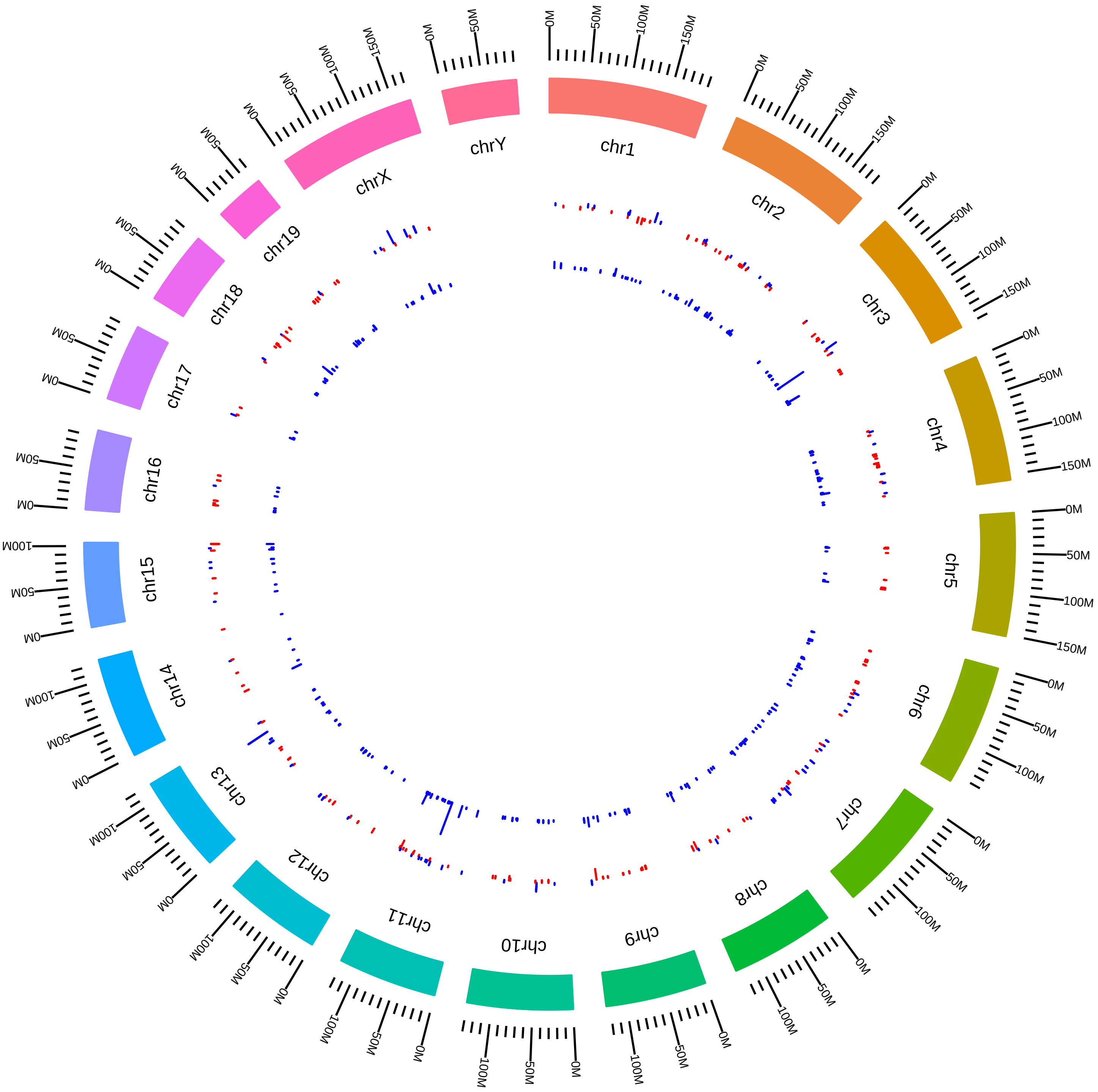

### rsfEsrp1Esrp2MmYiXing2015_DVsH.pdf

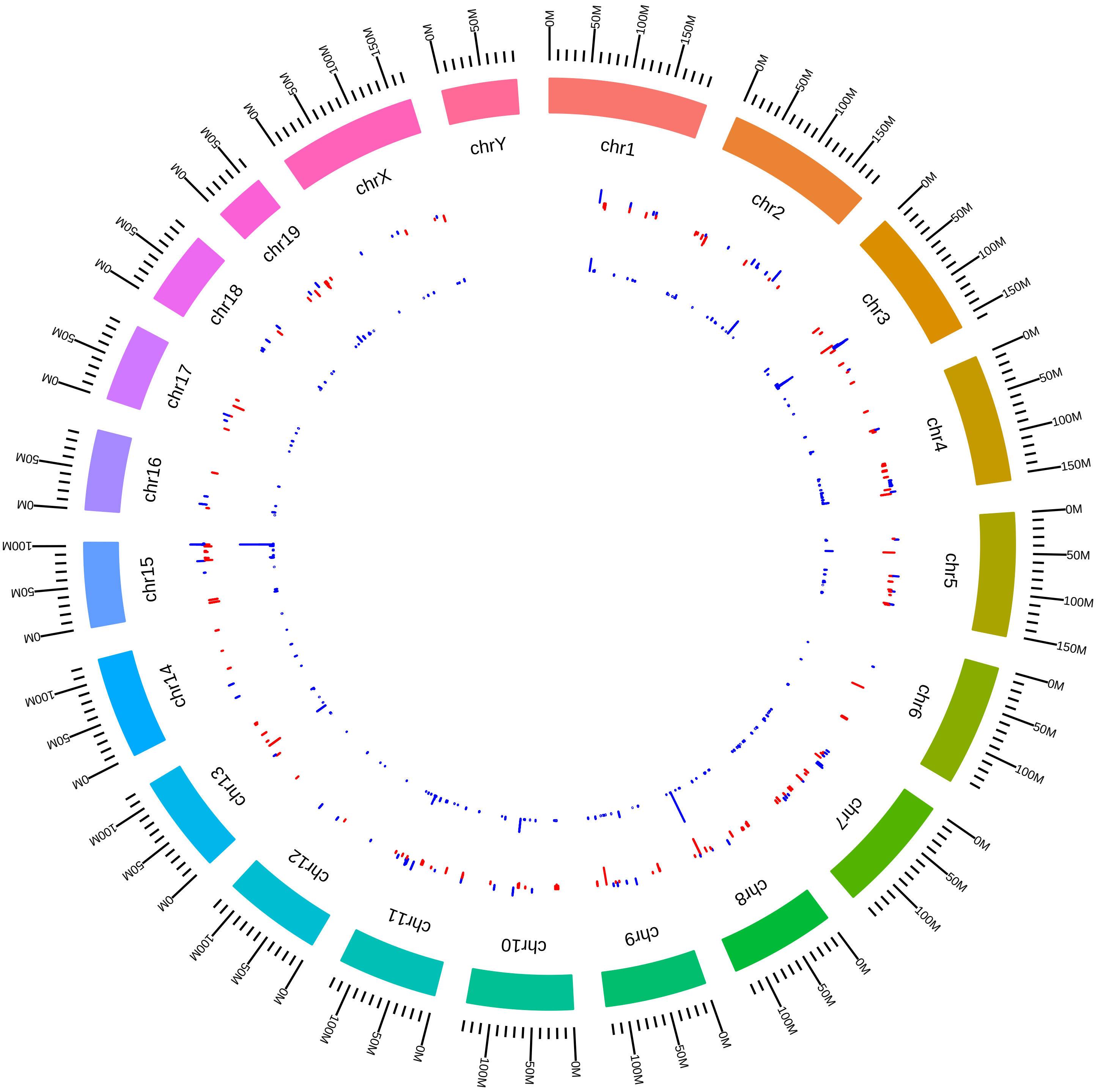

### rsfEsrp1Esrp2MmYiXing2015_DVsK.pdf

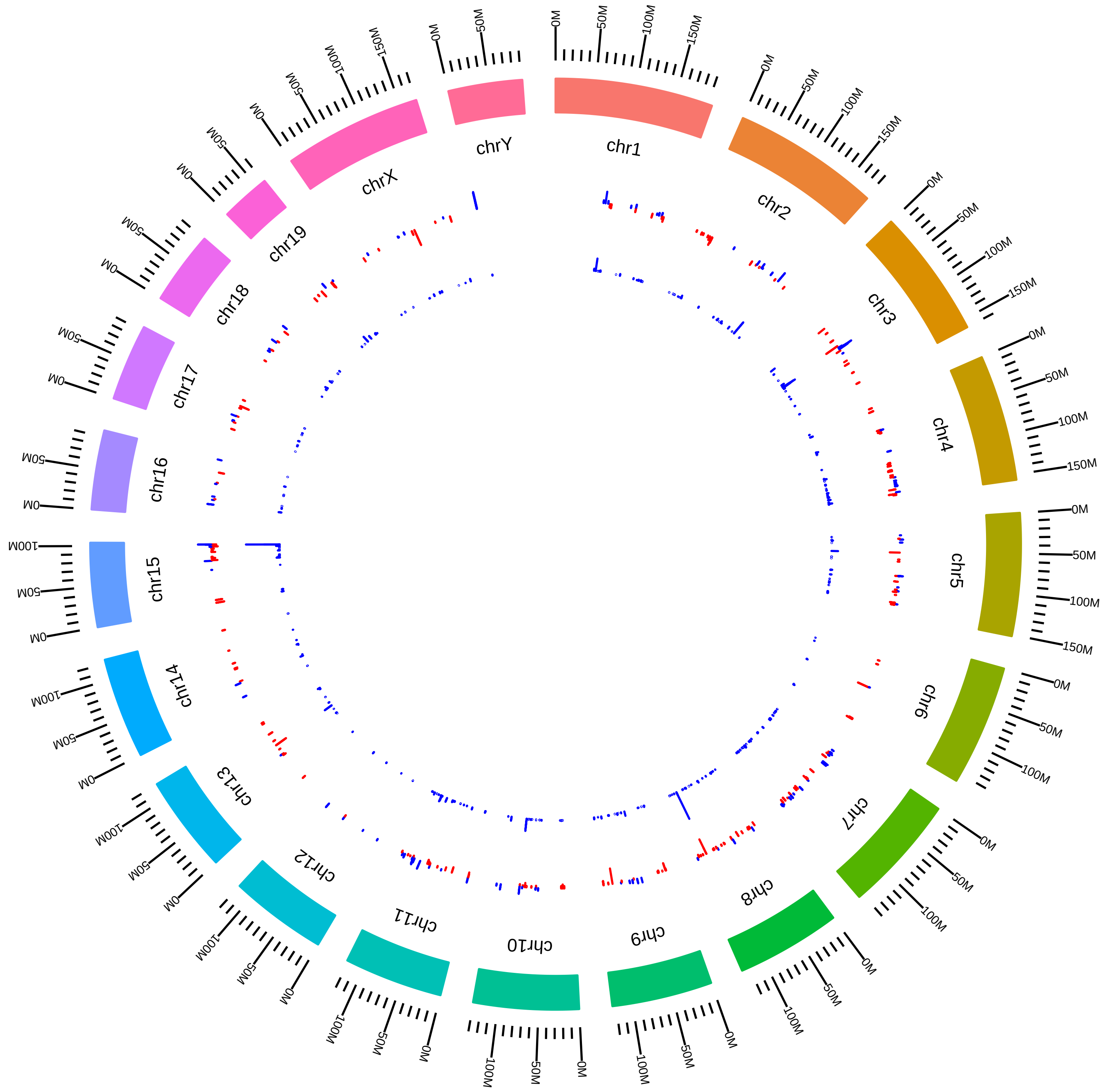

### rsfEsrp1Esrp2MmYiXing2015_DVsW.pdf

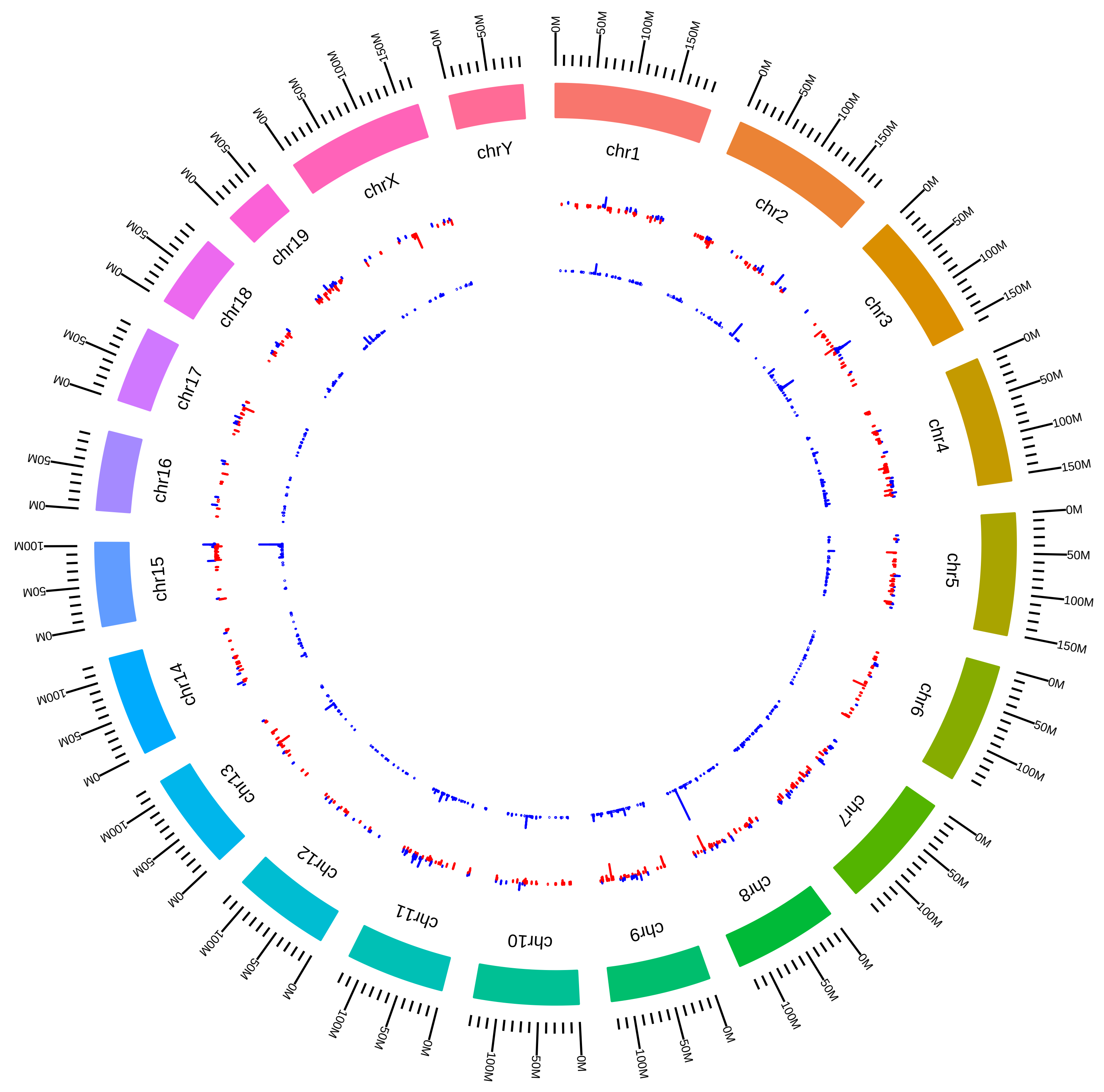

### rsfEsrp1Esrp2MmYiXing2015_HVsK.pdf

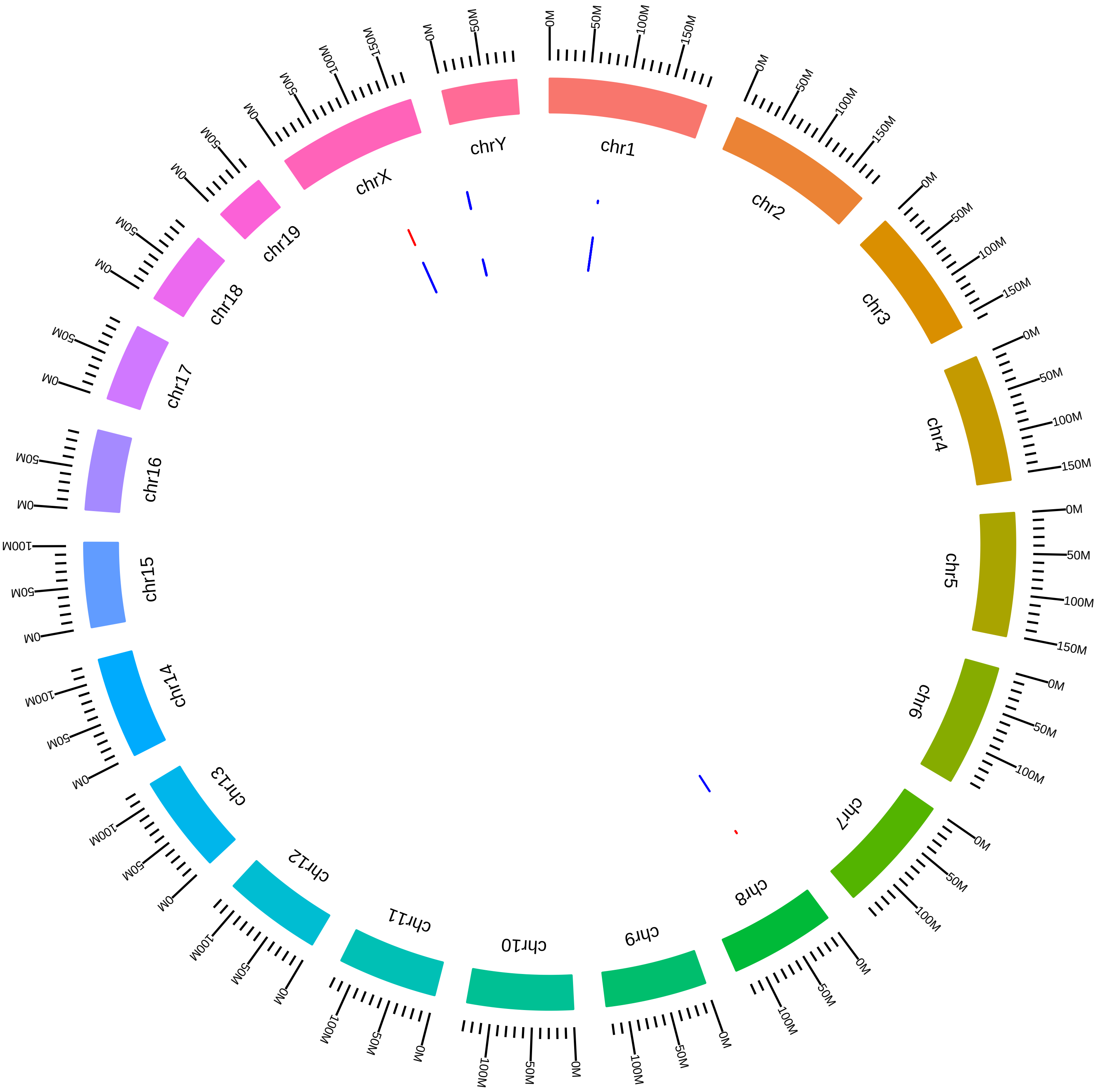

### rsfEsrp1Esrp2MmYiXing2015_HVsW.pdf

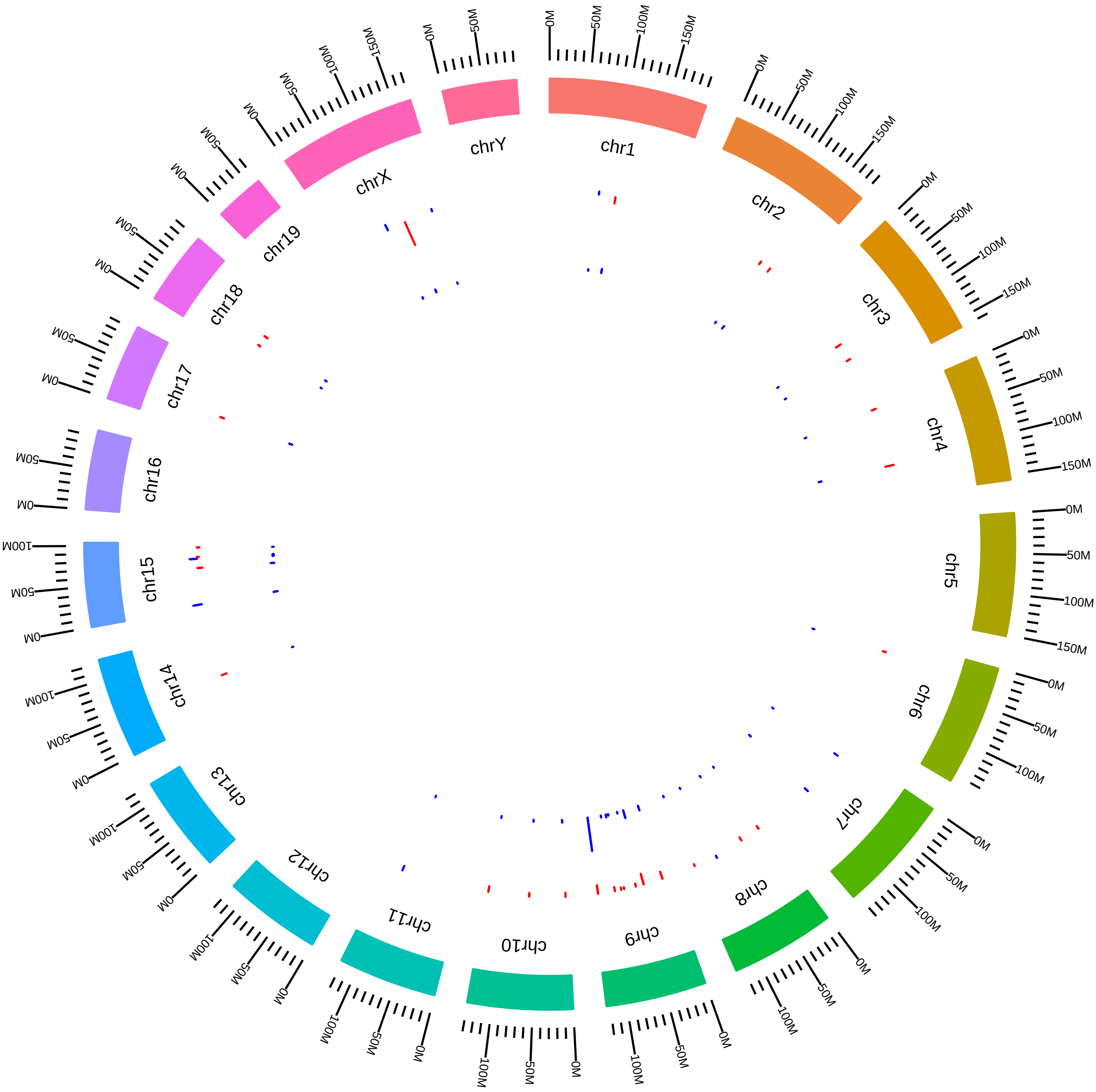

### rsfEsrp1Esrp2MmYiXing2015_KVsW.pdf

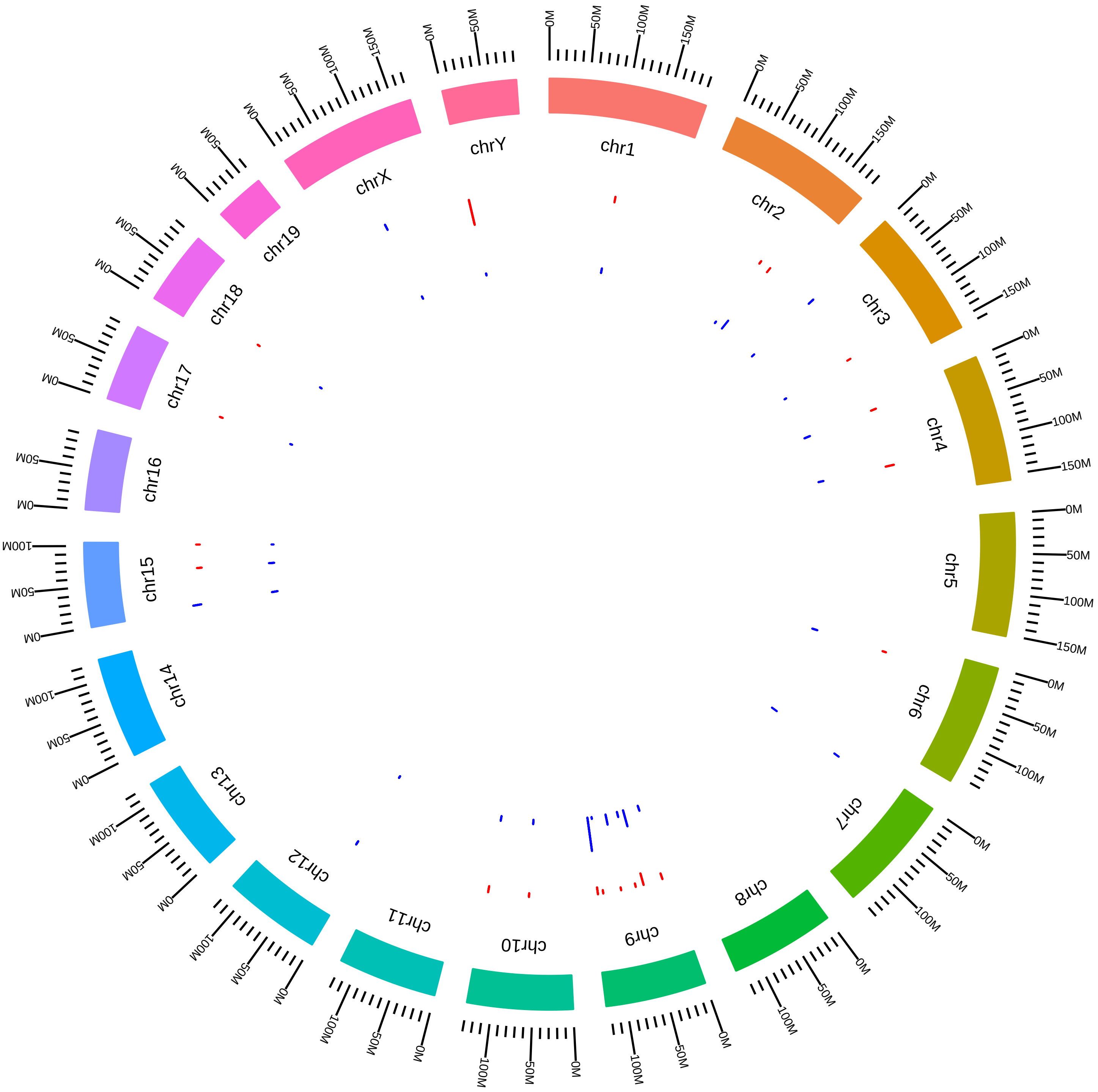

### rsfEsrp1MmRCarstens2016_DEG_KVsC.pdf

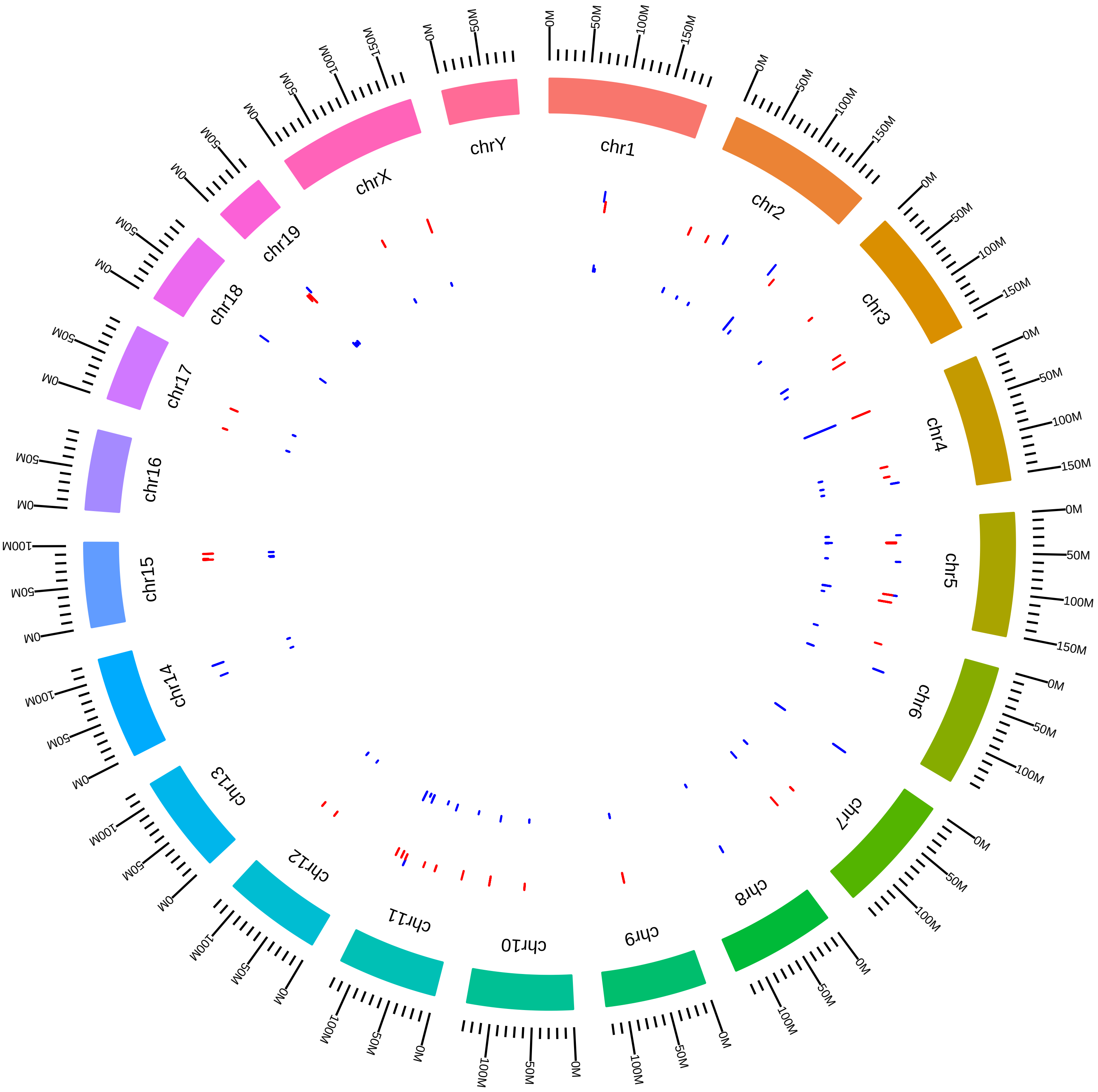

### rsfEsrp1MmRCarstens2016_KVsC.pdf

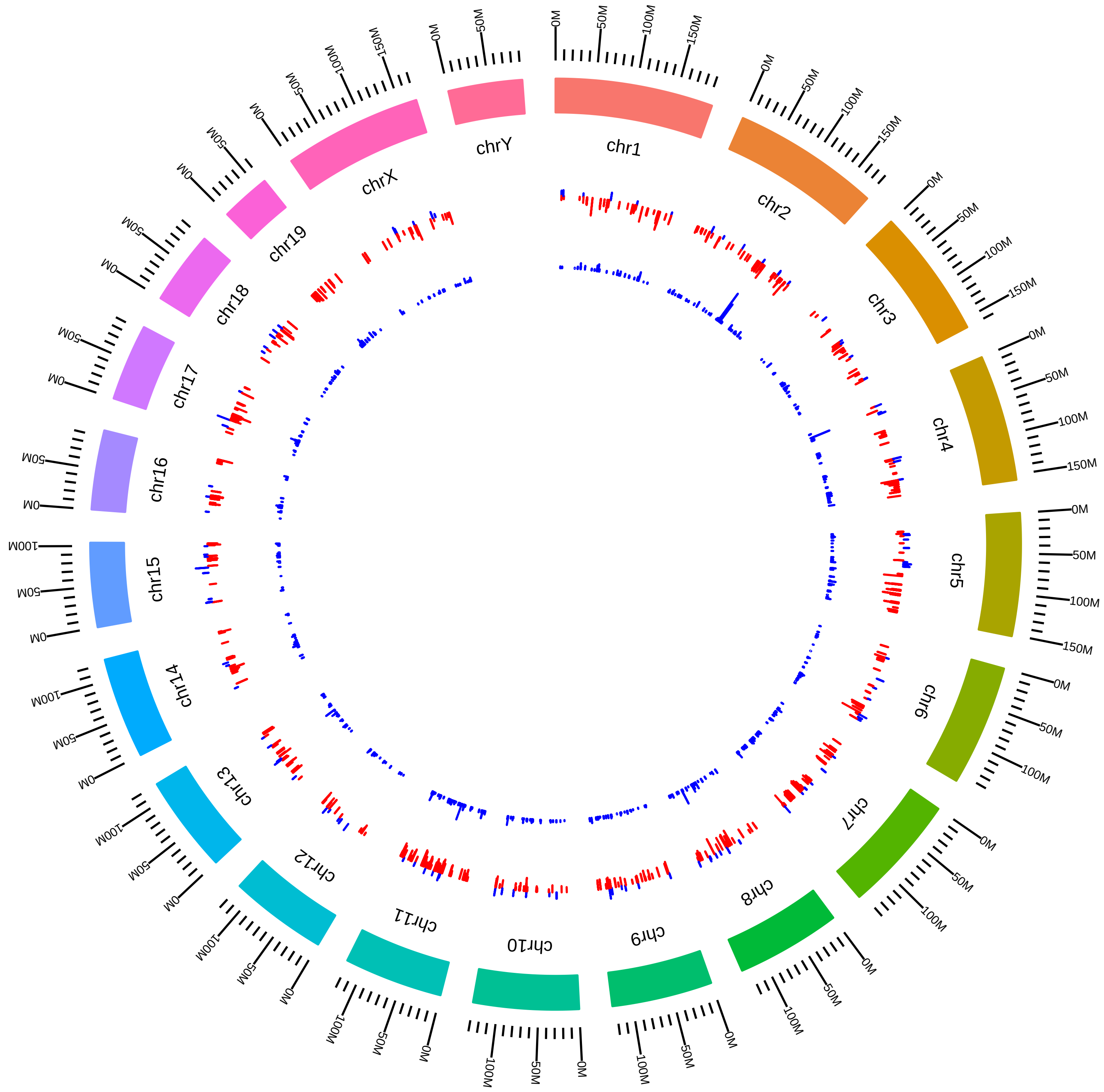

### rsfHnrnpa2b1MmGYeo2016_HVsC.pdf

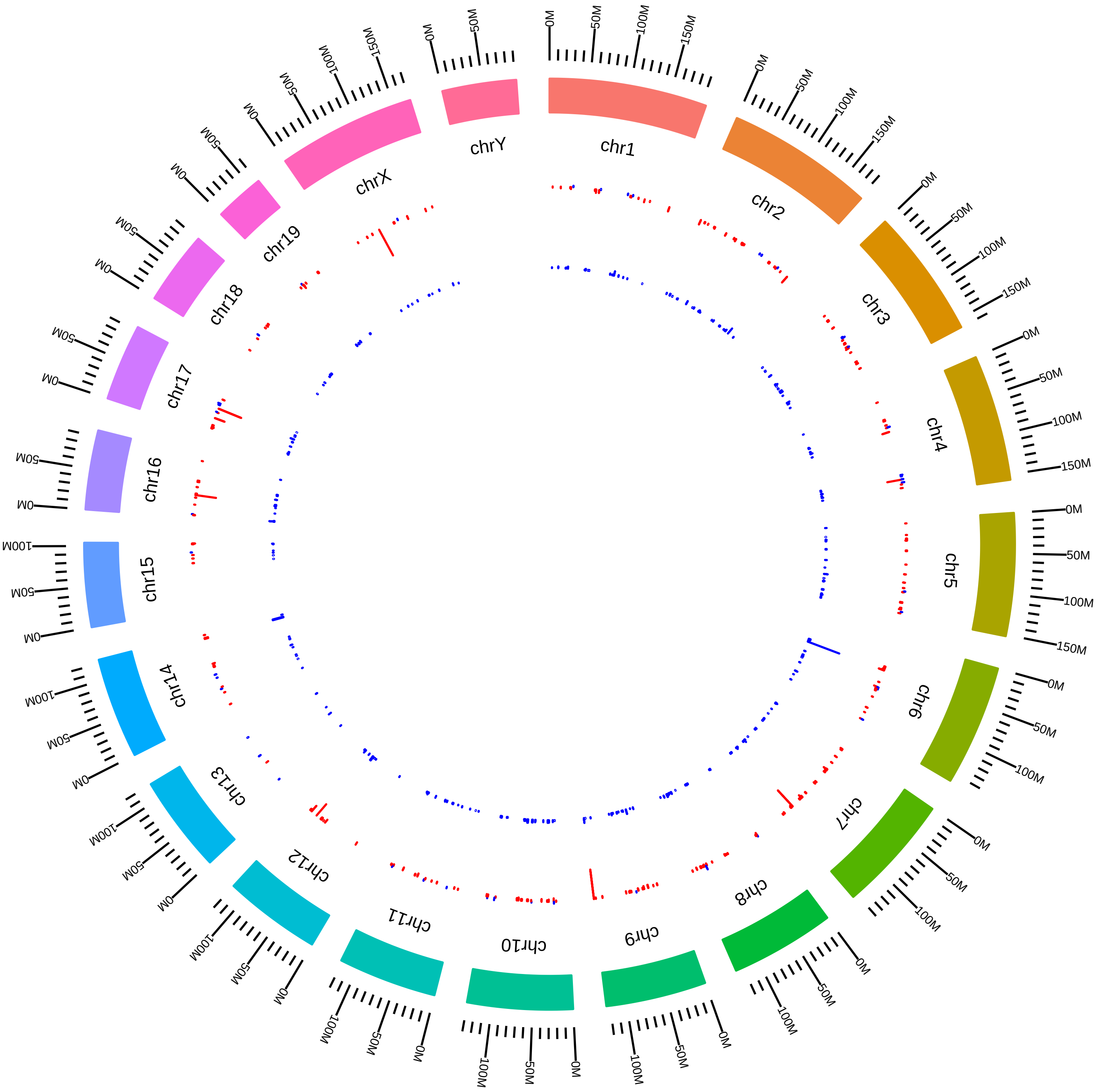

### rsfHnrnph1MmCBryant_MKVsMW.pdf

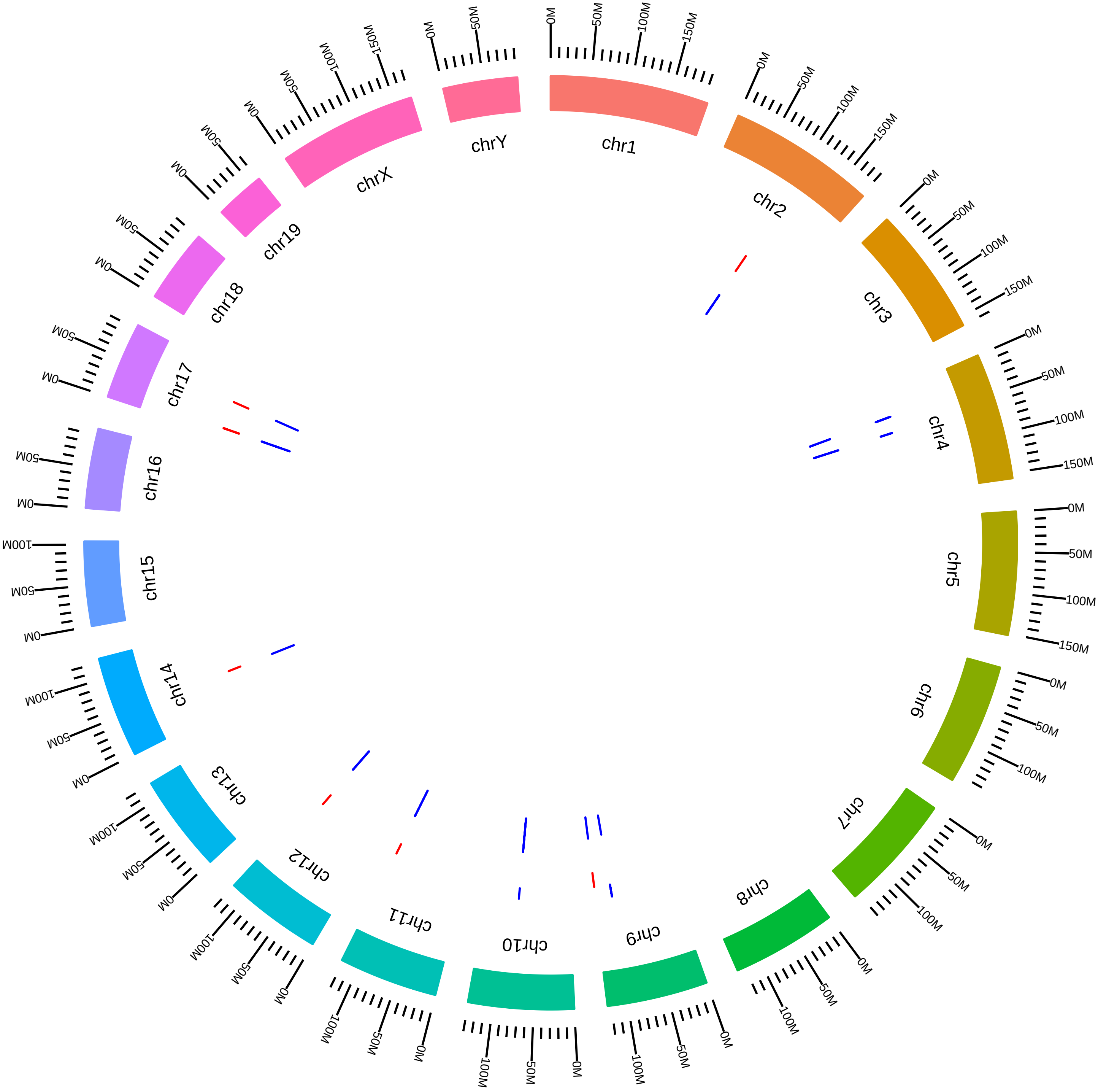

### rsfHnrnpkMmMLorincz2015_DVsC.pdf

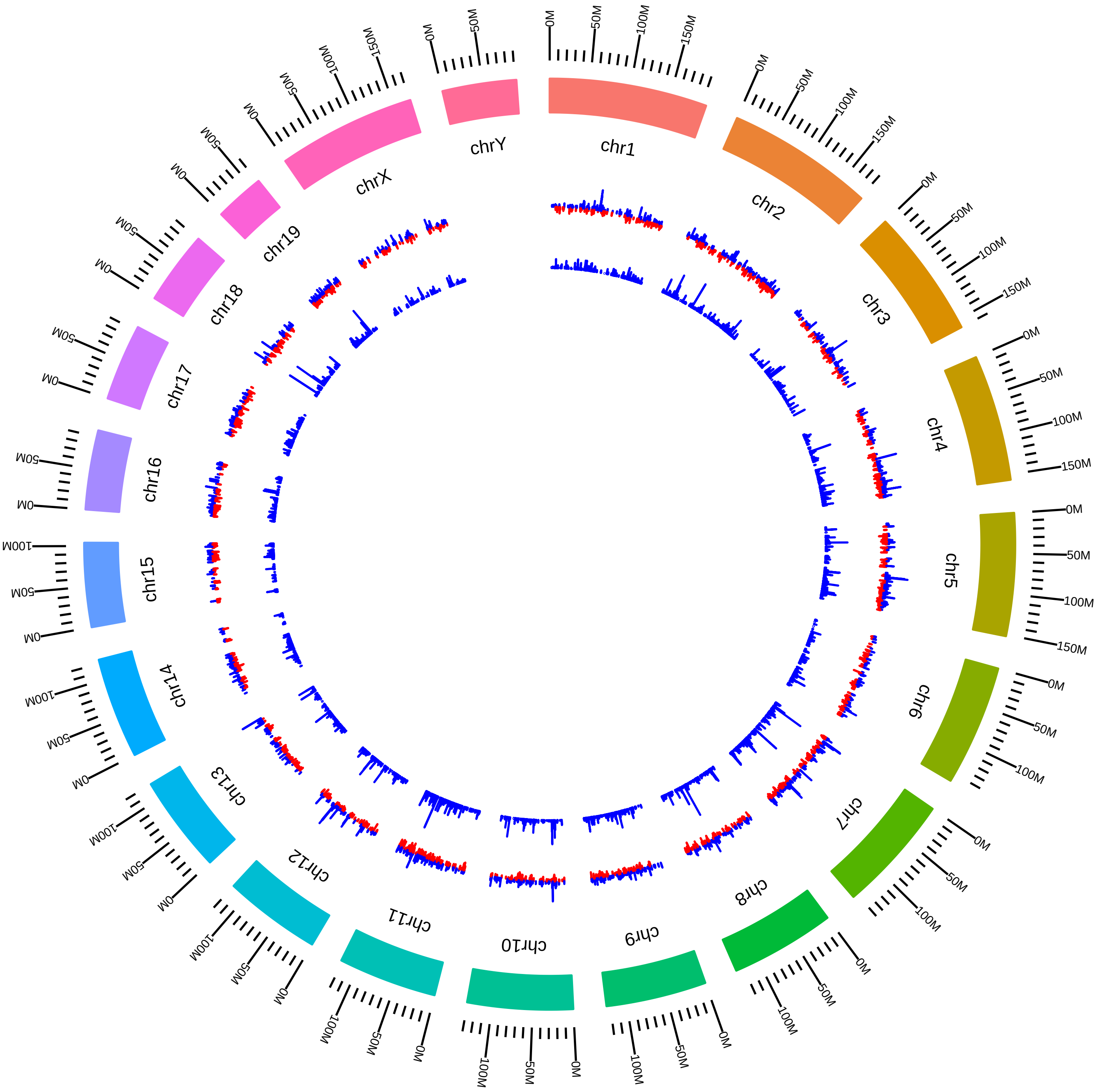

### rsfHnrnplMmTMoroy2016_KVsW.pdf

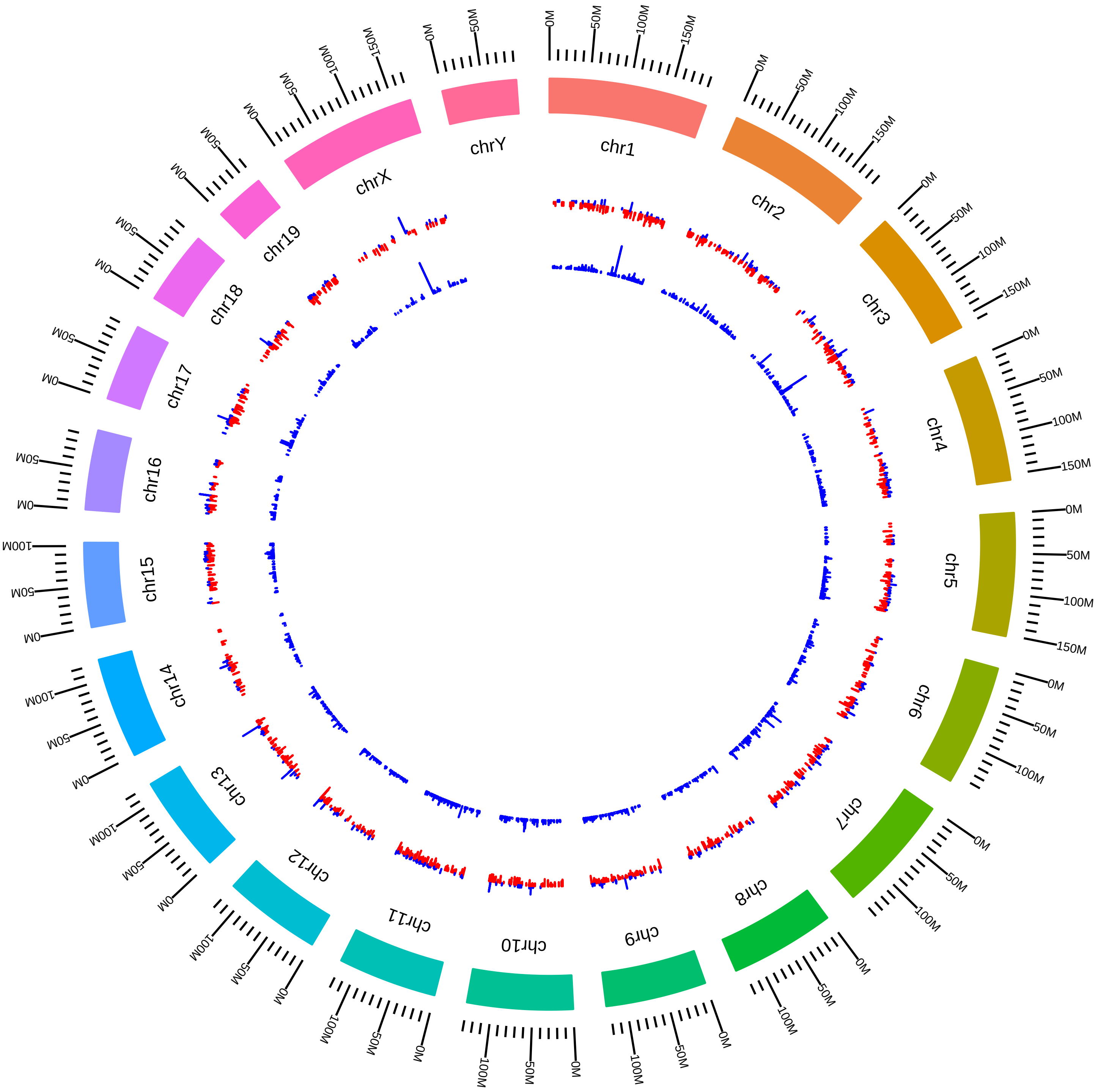

### rsfHnrnpuMmTManiatis_DEG_KVsW.pdf

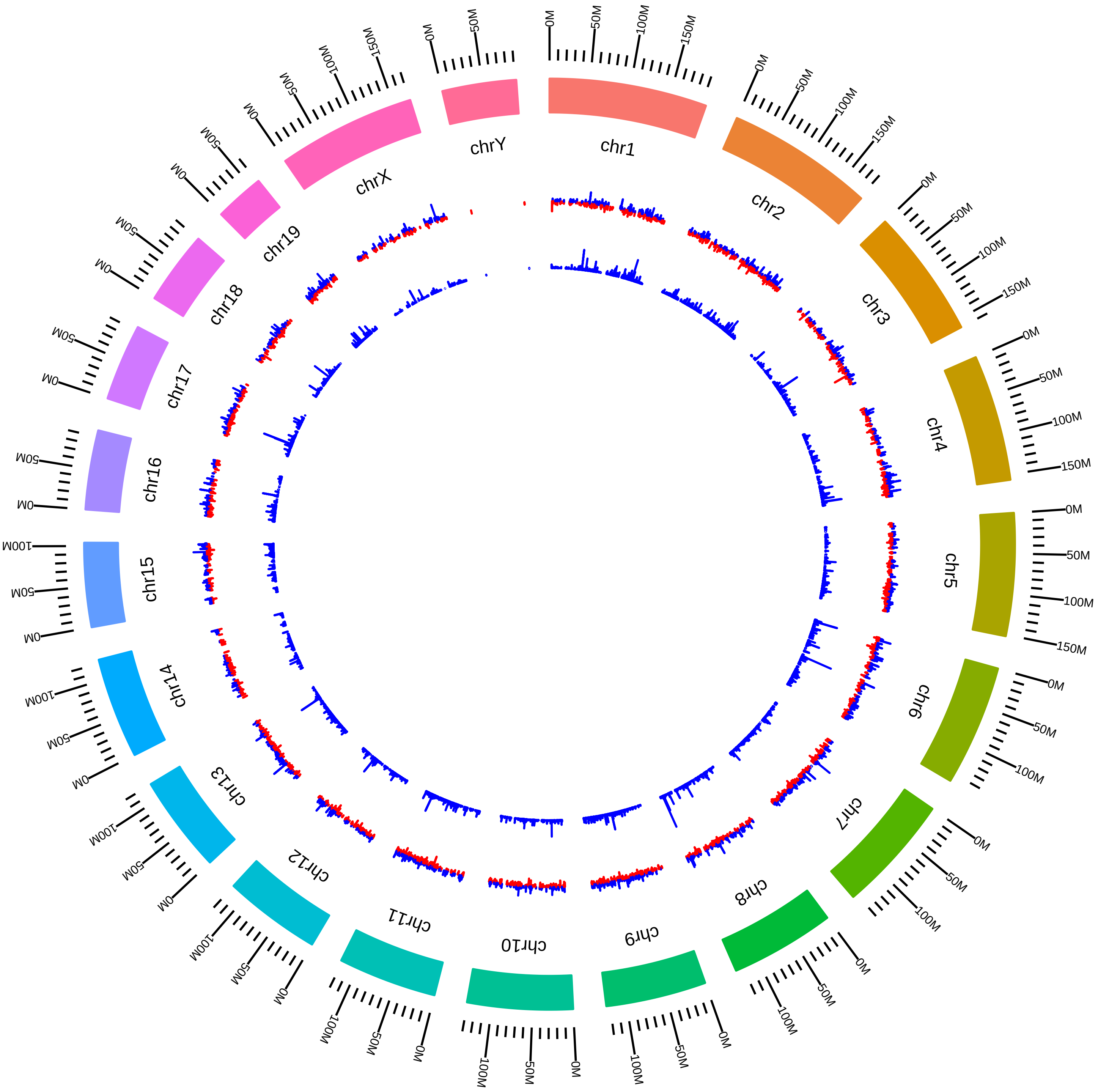

### rsfJmjd6MmDLittman_DEG_DVsW.pdf

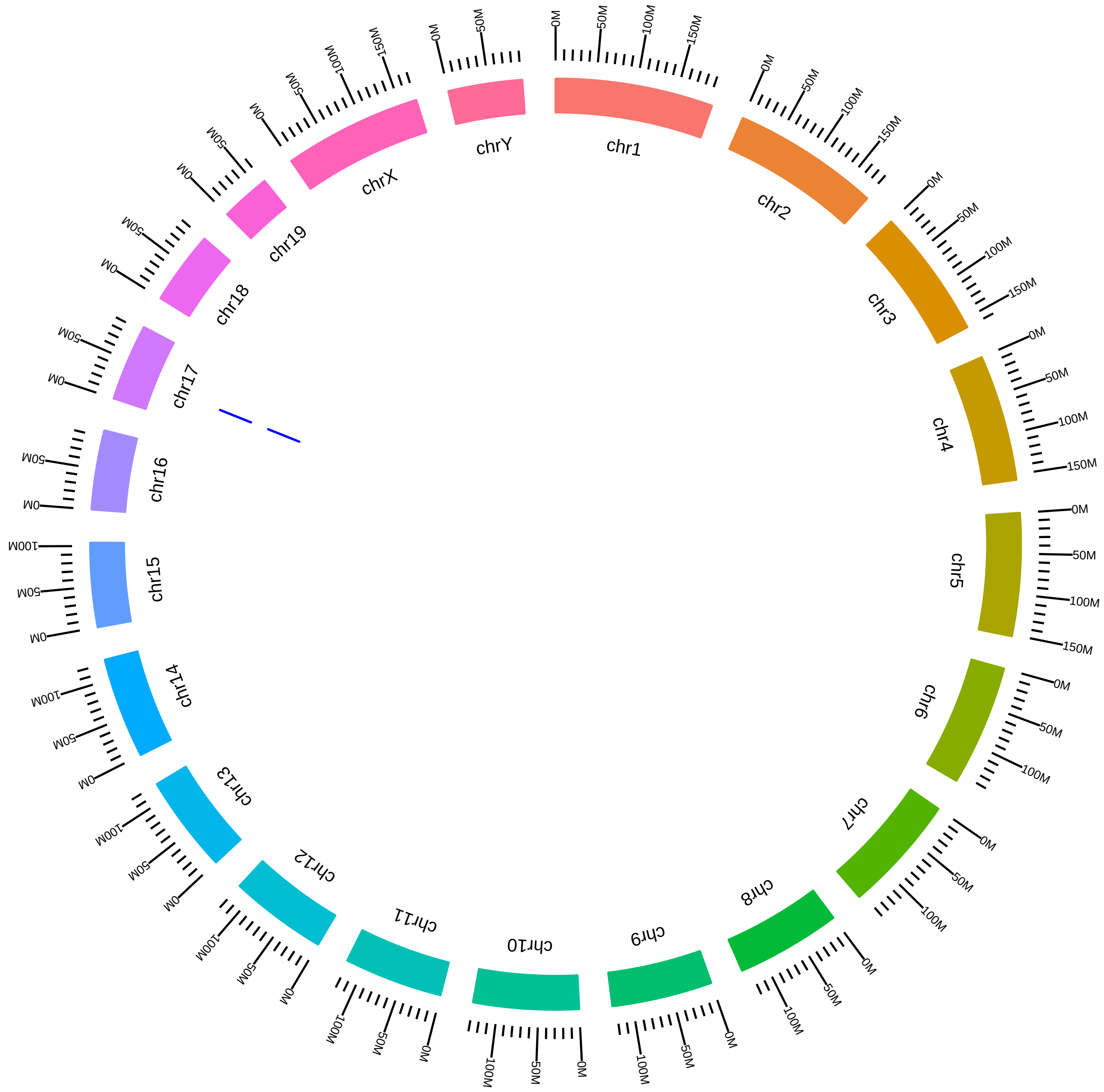

### rsfJmjd6MmFYoshinori2015_KVsW.pdf

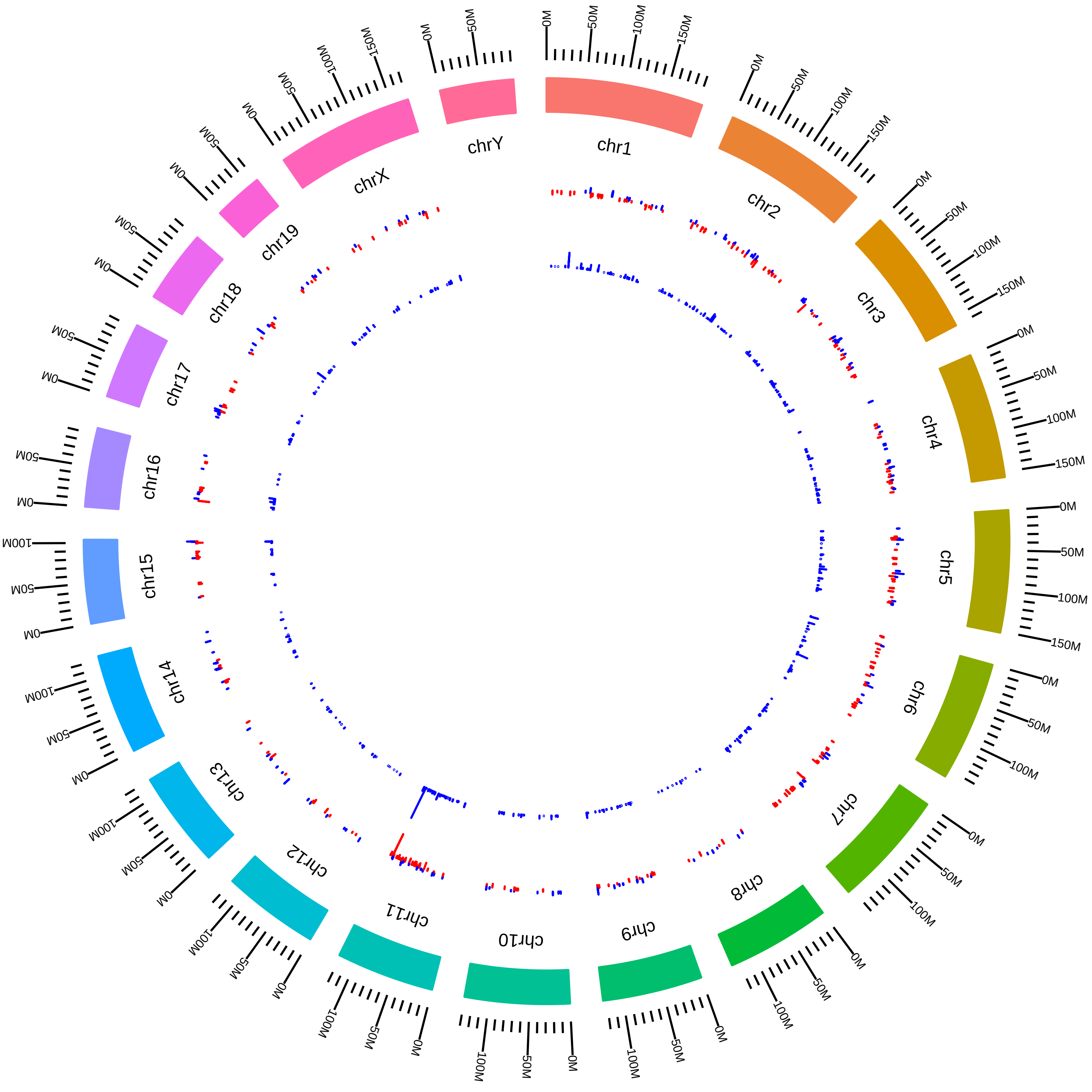

### rsfKhdrbs3MmDElliott2016_KVsW.pdf

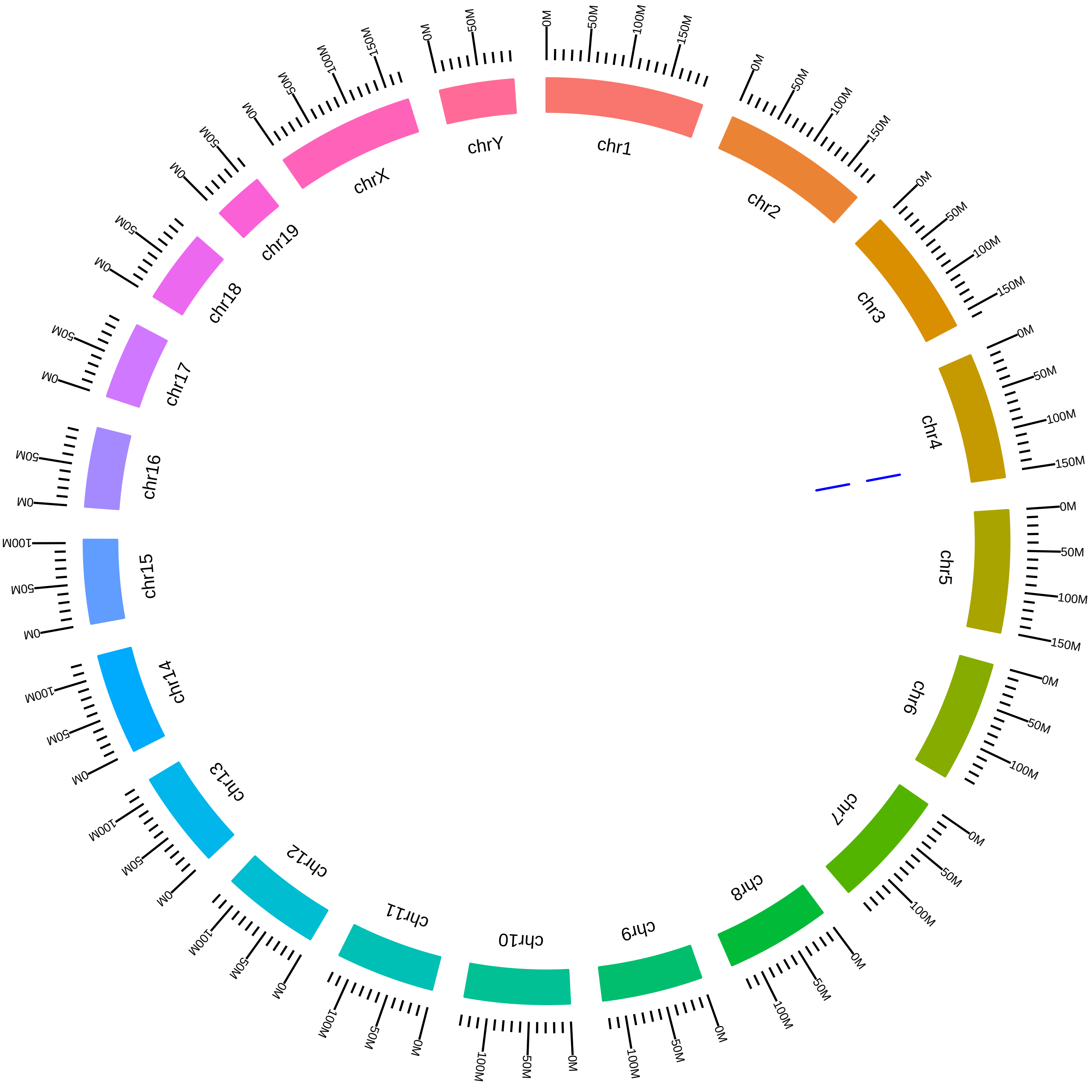

### rsfMagohMutMmDSilver2016_HVsW.pdf

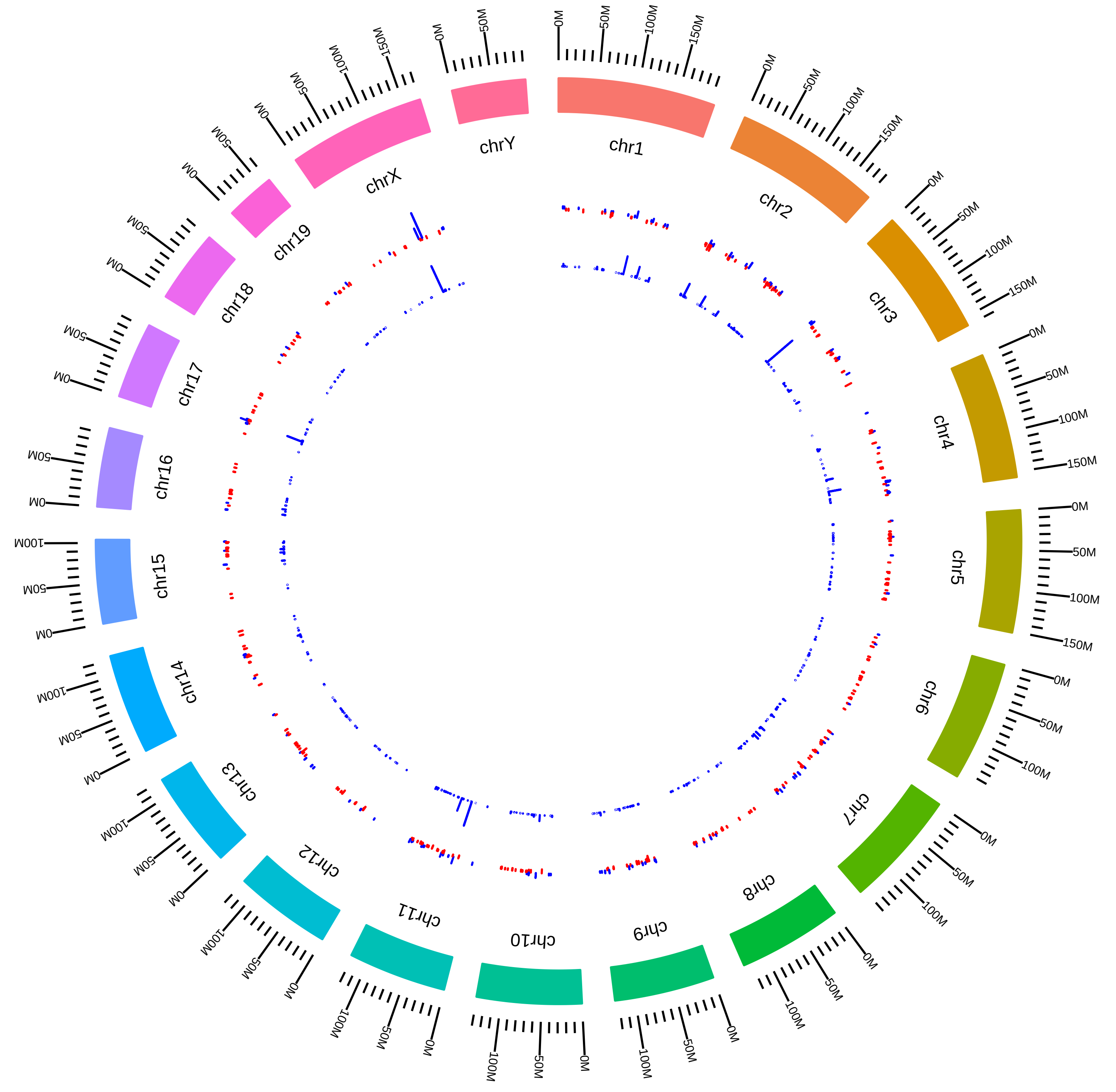

### rsfMalat1MmDLSpector_DEG_OTVsOC.pdf

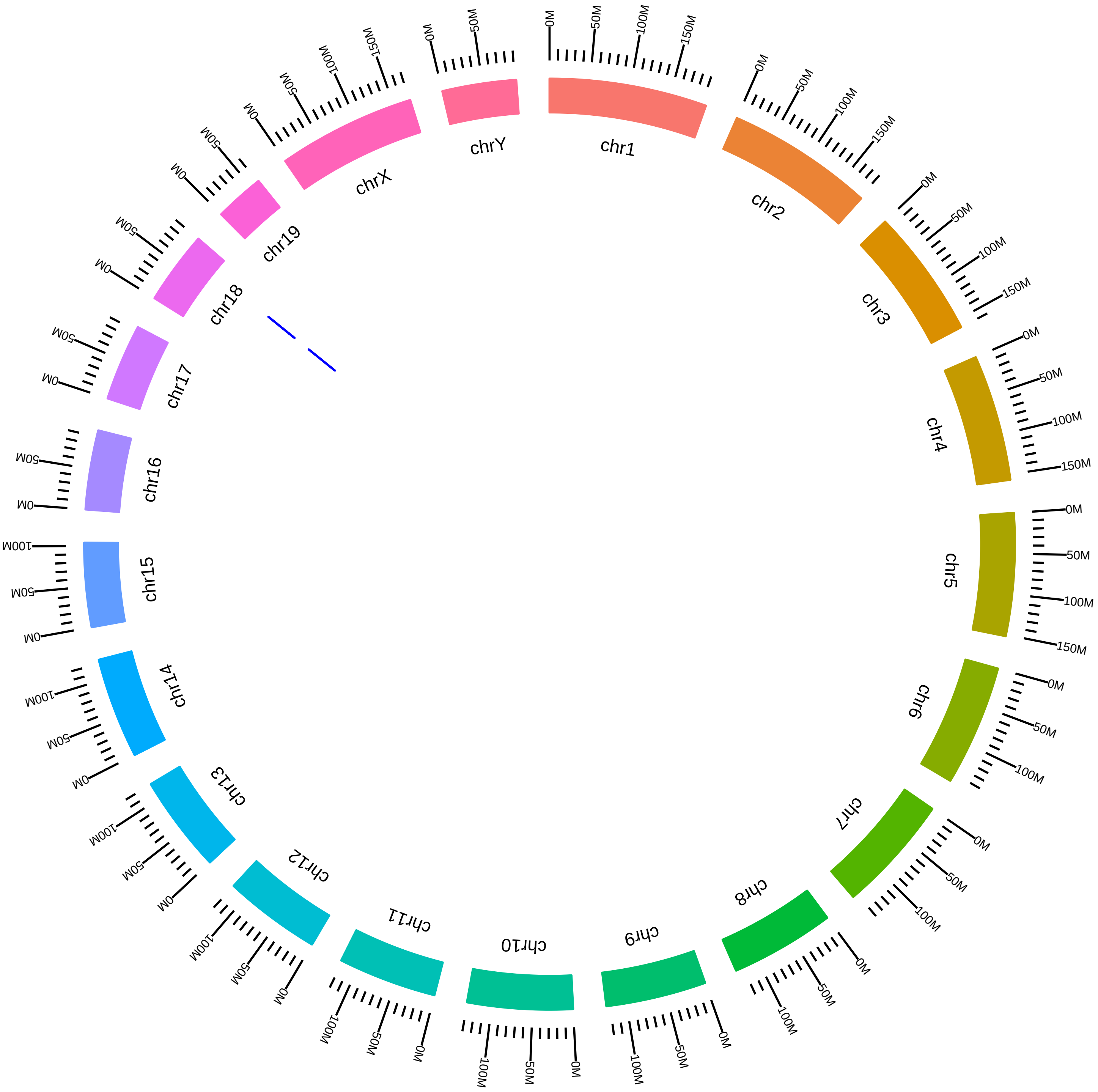

### rsfMalat1MmDLSpector_DEG_TTVsTC.pdf

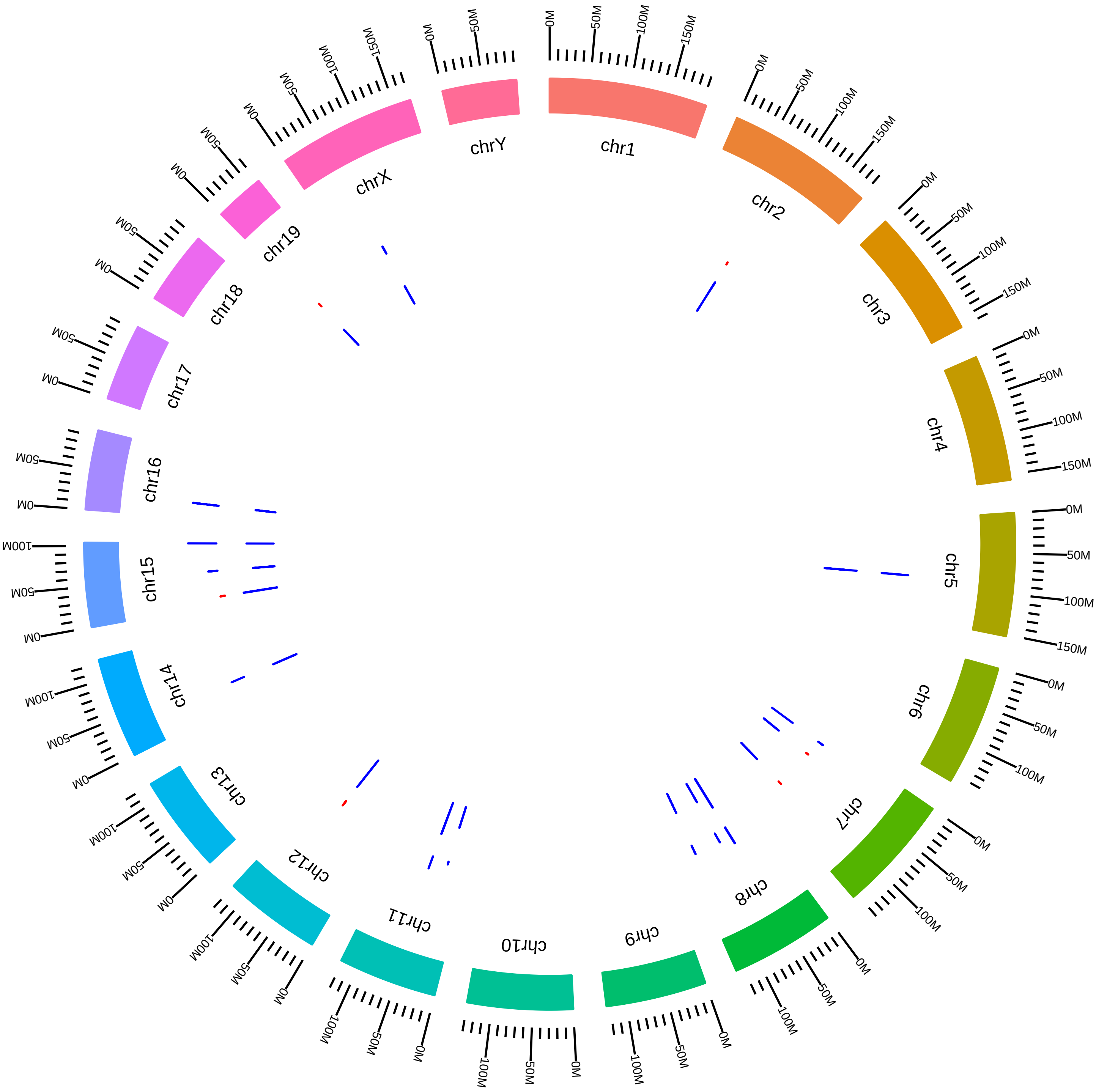

### rsfMbnl1Mbnl2Mbnl3MefMmMSwanson2014_DVsC.pdf

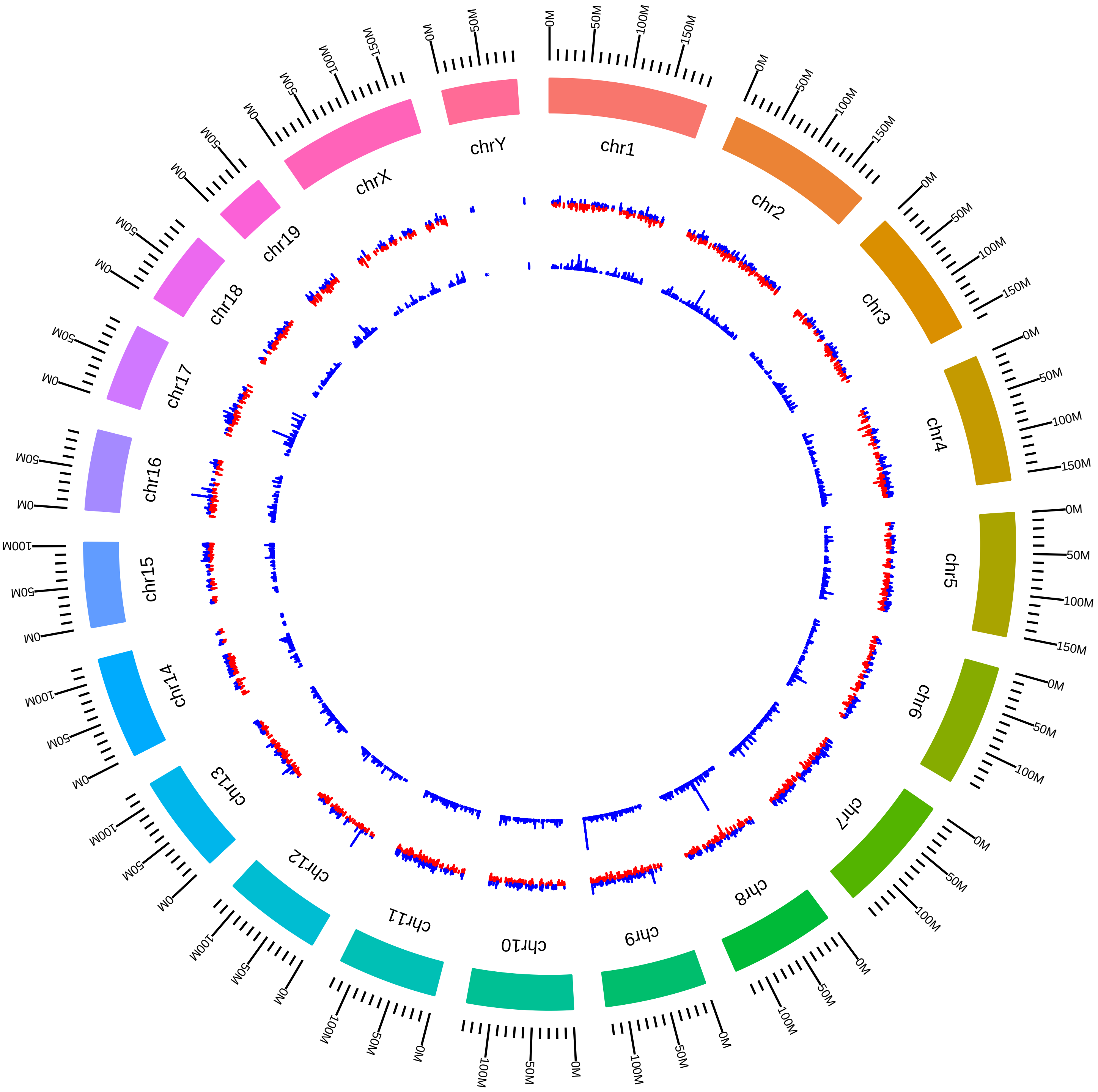

### rsfMbnl1Mbnl2Mbnl3MefMmMSwanson2014_DVsK.pdf

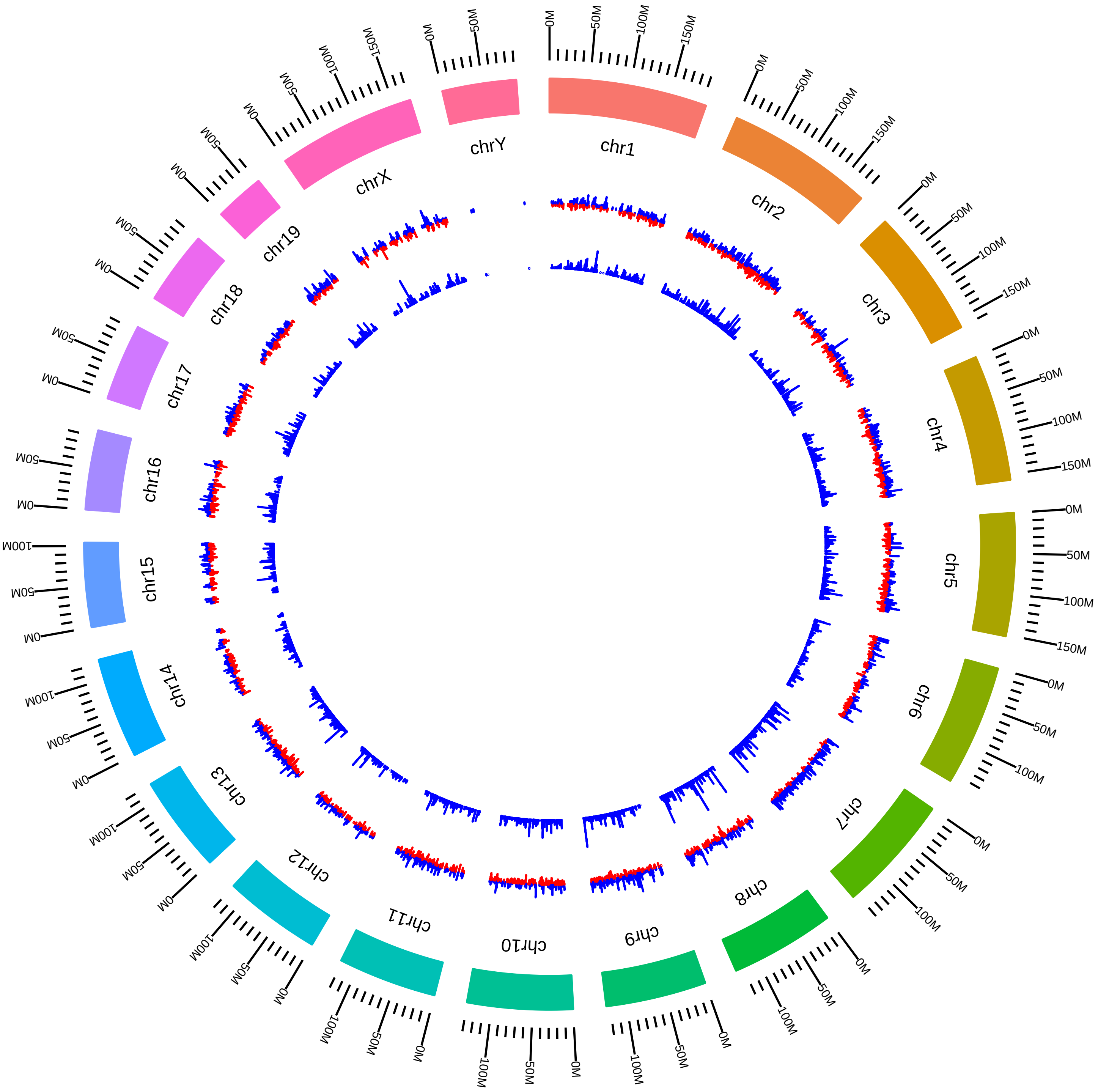

### rsfMbnl1Mbnl2Mbnl3MefMmMSwanson2014_DVsW.pdf

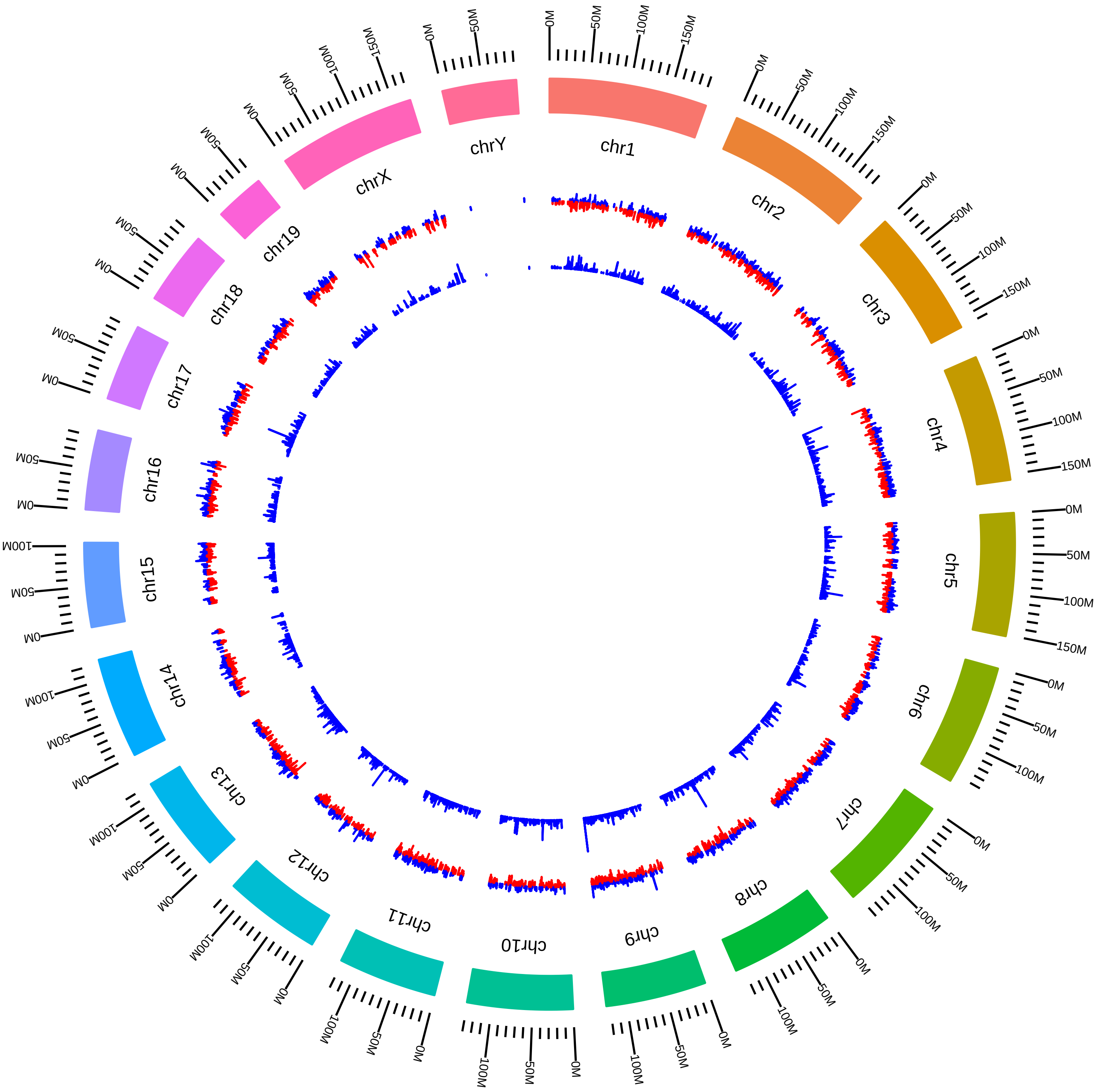

### rsfMbnl1Mbnl2Mbnl3MefMmMSwanson2014_KVsW.pdf

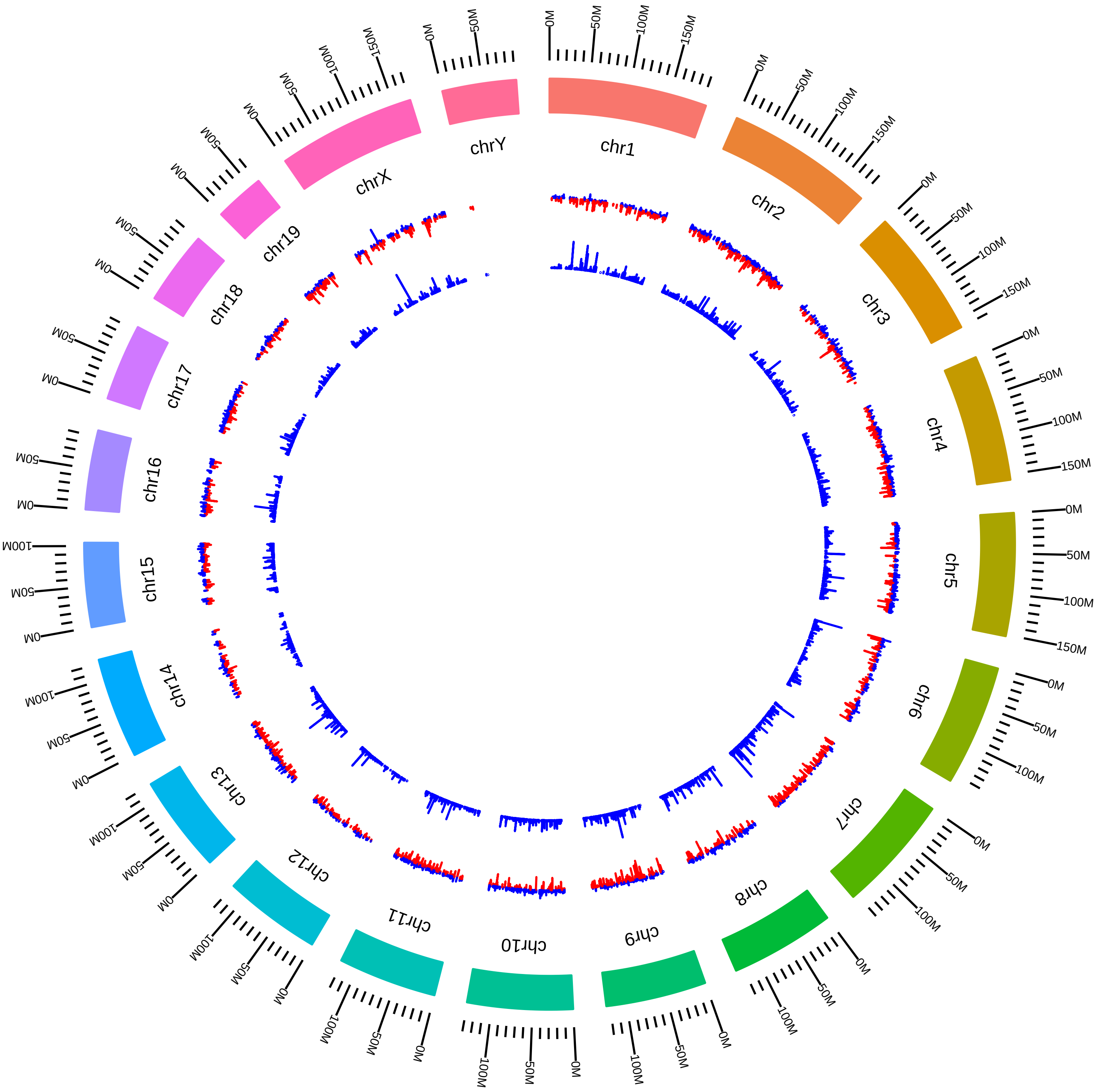

### rsfMbnl1Mbnl2Mbnl3MusMmMSwanson2014_M4VsW4.pdf

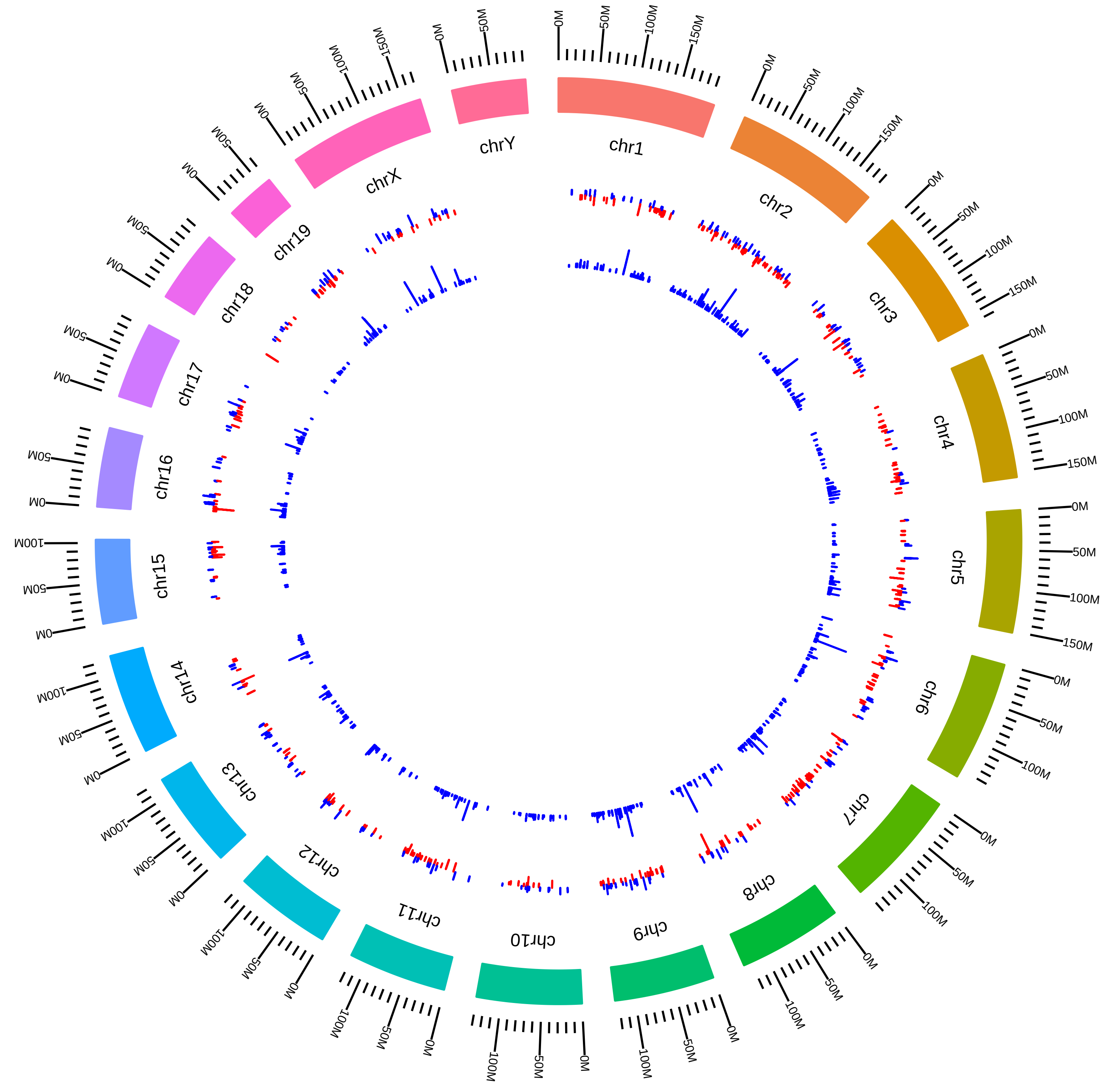

### rsfMbnl1Mbnl2MmCBurge2016_DEG_CN1VsCNW.pdf

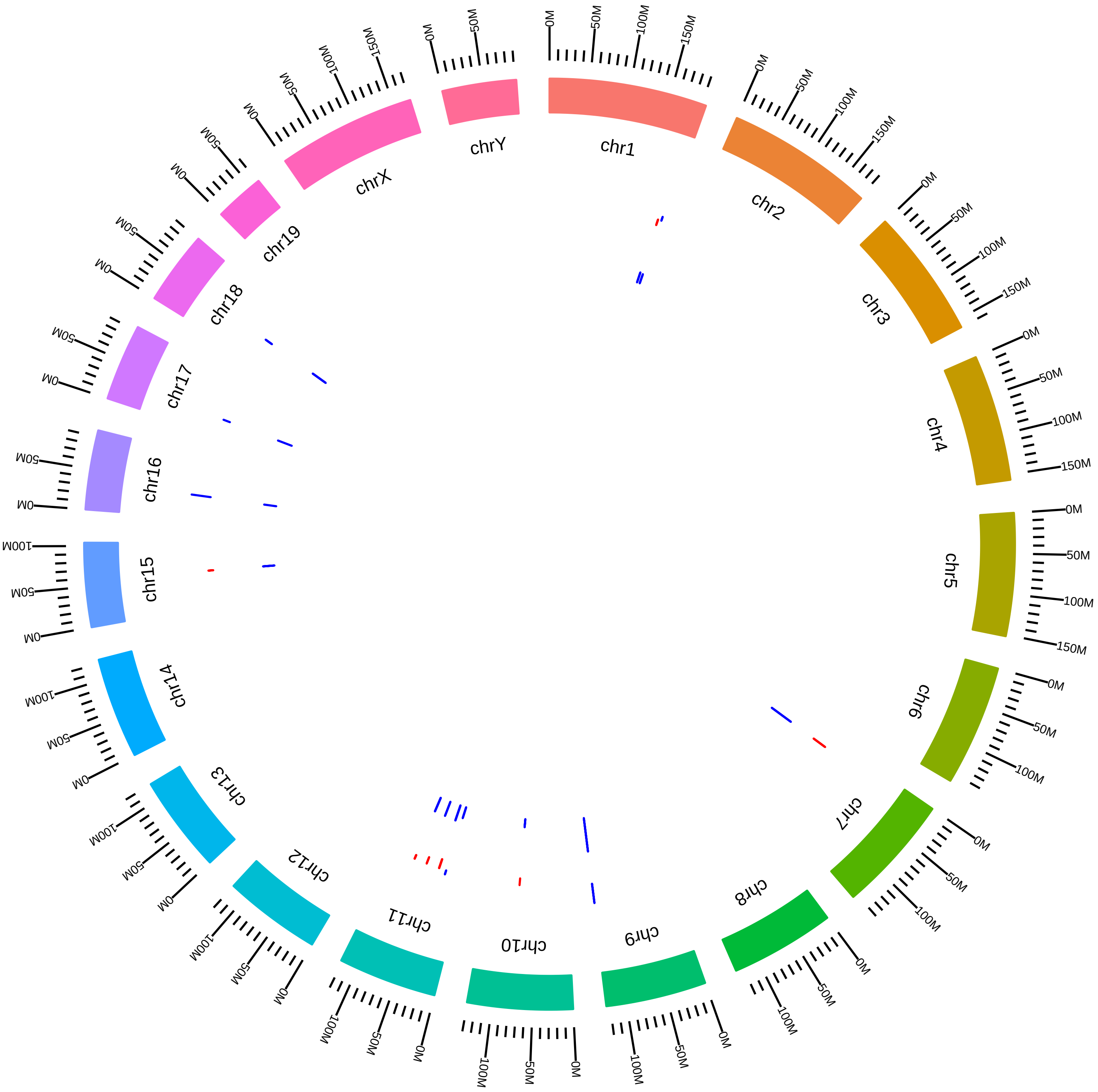

### rsfMbnl1Mbnl2MmCBurge2016_DEG_CN2VsCNW.pdf

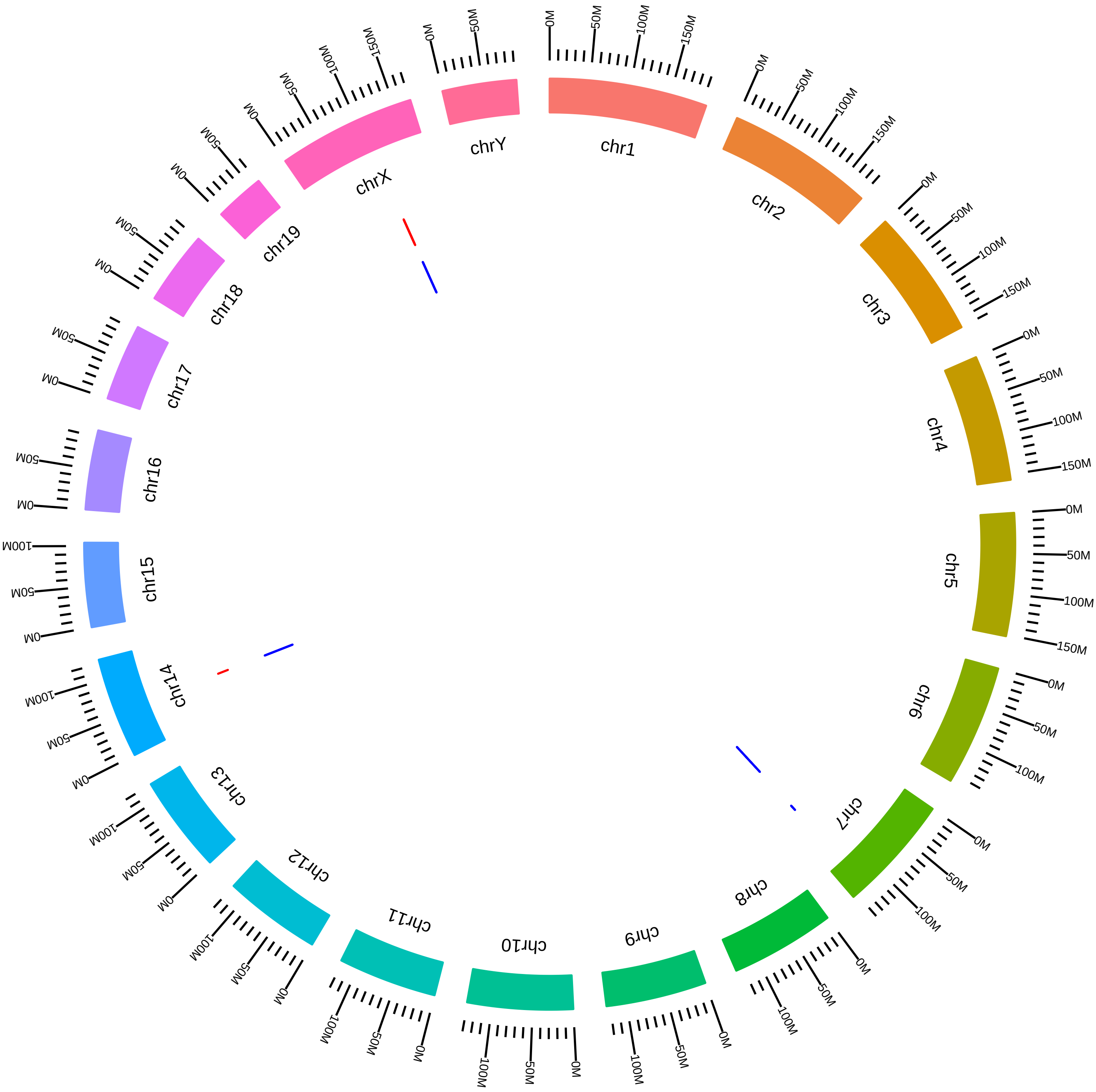

### rsfMbnl1Mbnl2MmCBurge2016_DEG_CS1VsCSW.pdf

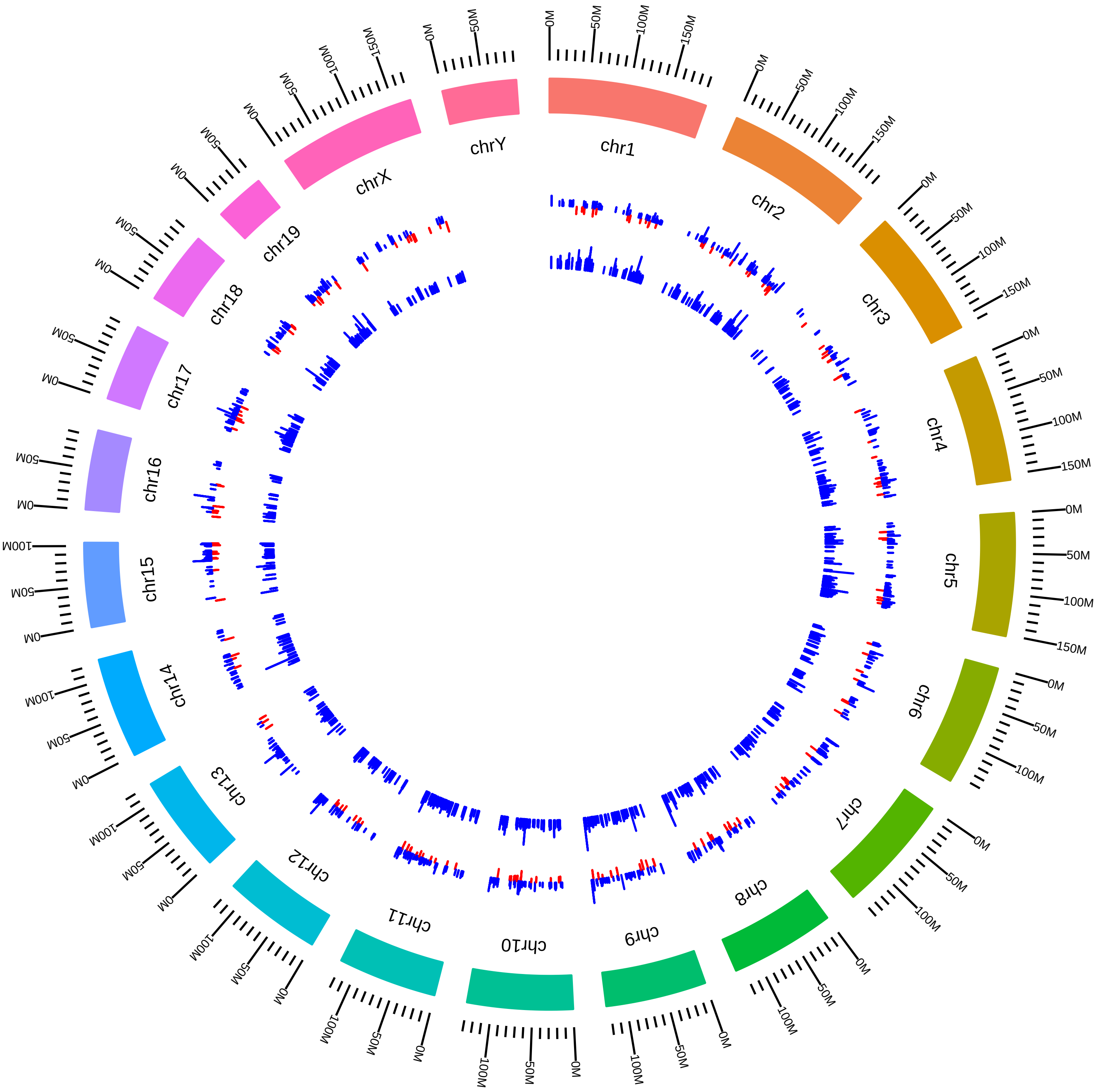

### rsfMbnl1Mbnl2MmCBurge2016_DEG_CS2VsCSW.pdf

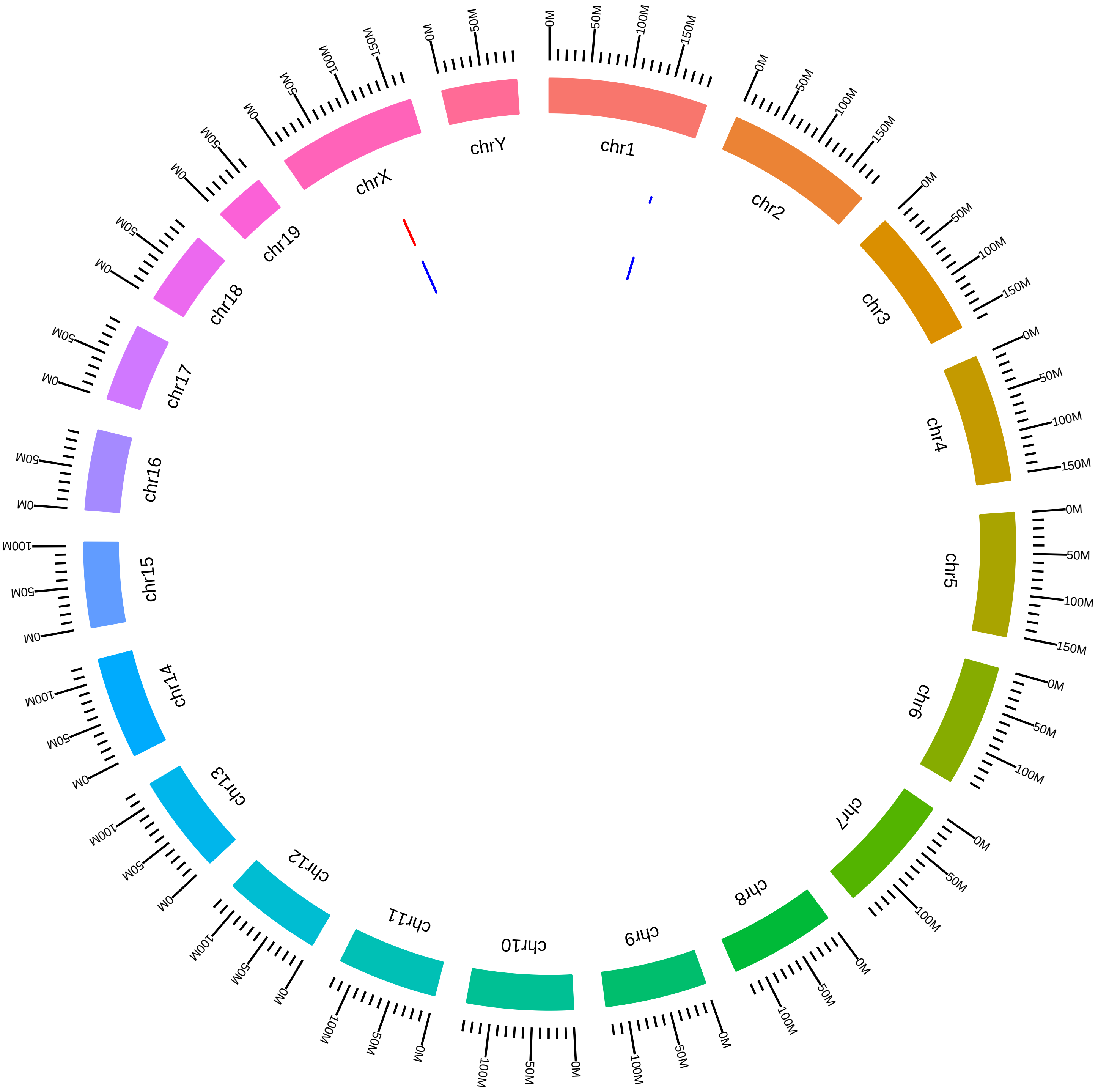

### rsfMbnl1Mbnl2MmCBurge2016_DEG_DNDVsDNW.pdf

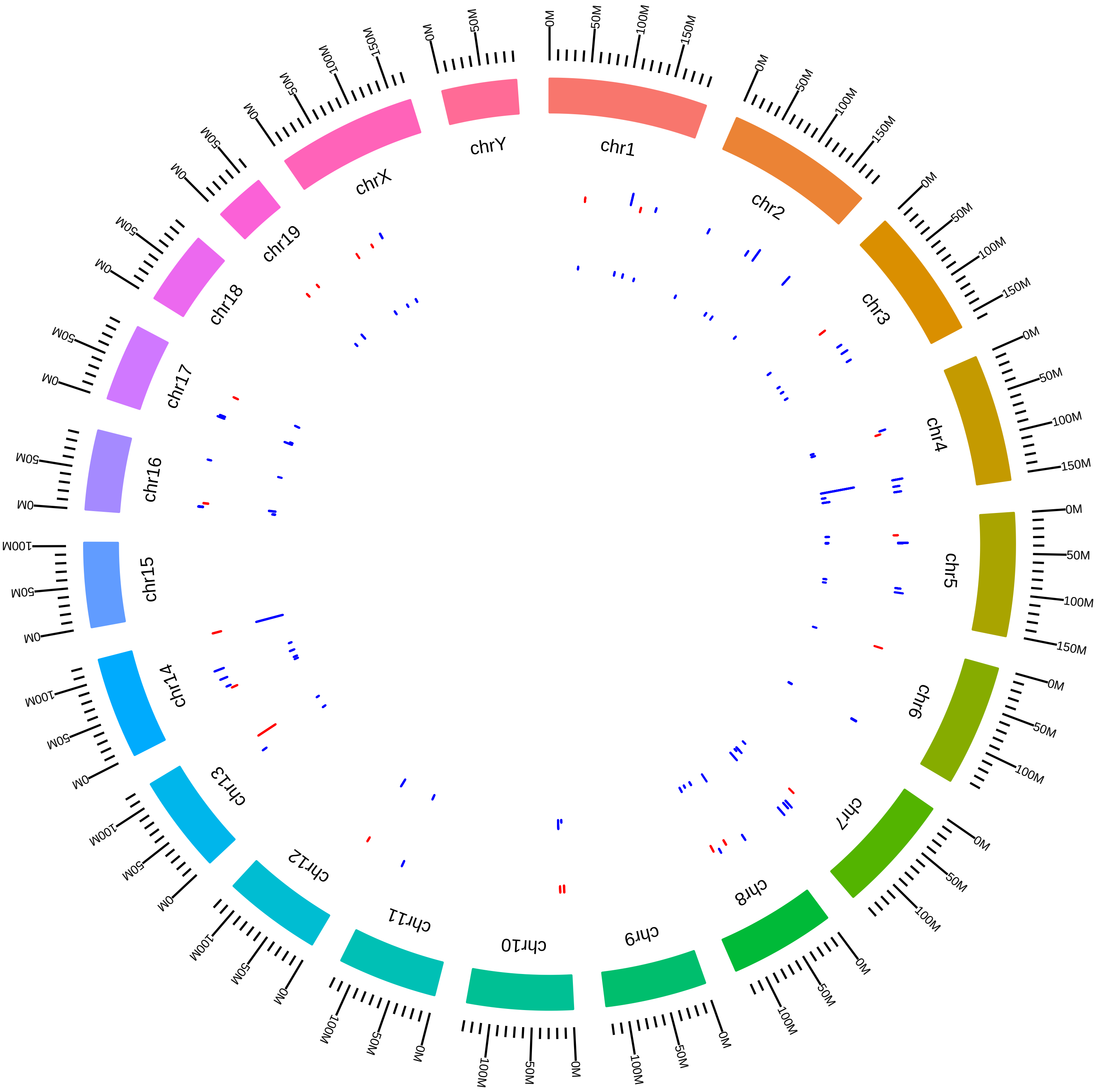

### rsfMbnl1Mbnl2MmCBurge2016_DEG_DSDVsDSW.pdf

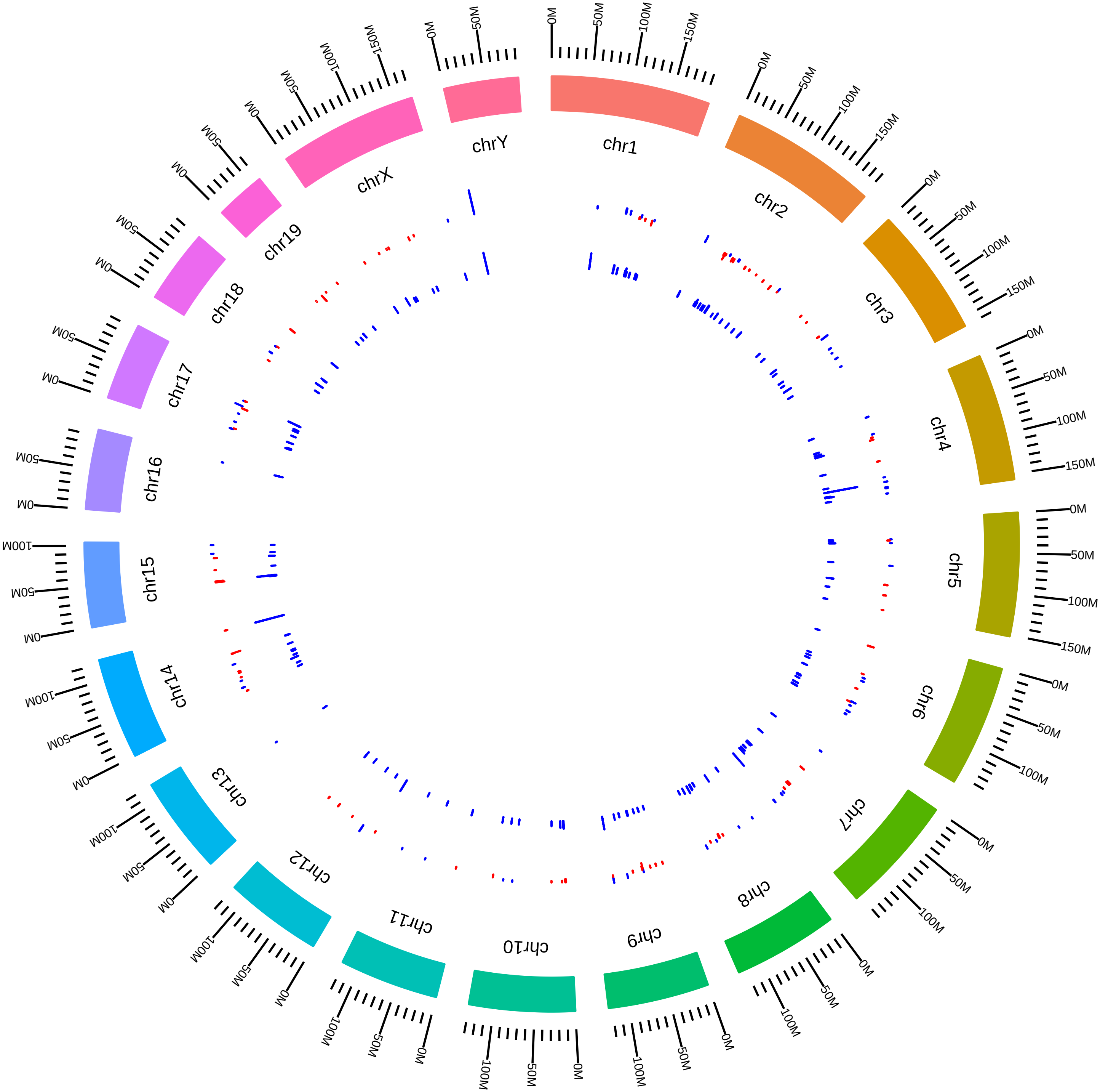

### rsfMbnl1Mbnl2MmEWang_DEG_KVsW.pdf

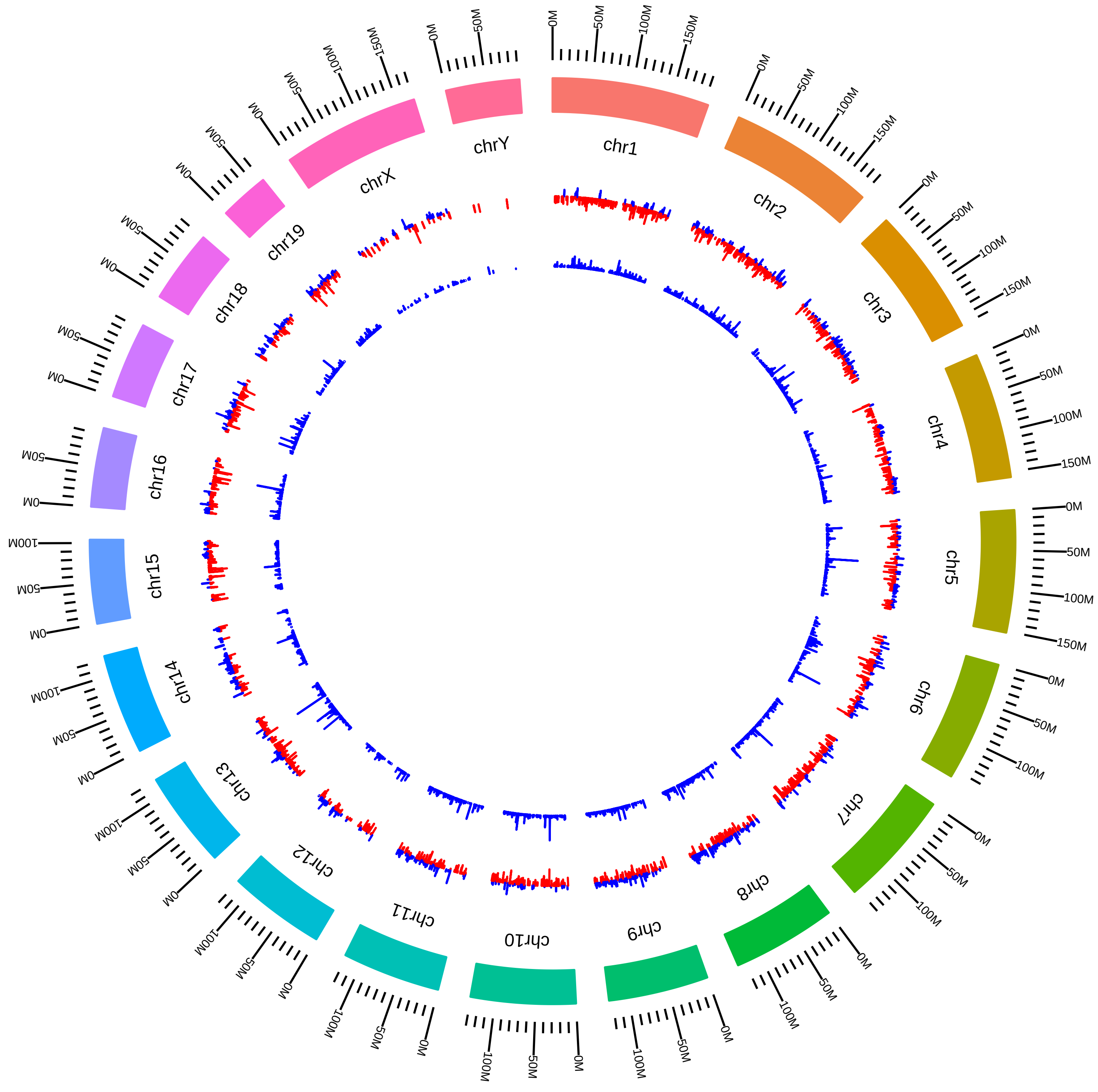

### rsfMbnl1Mbnl2MmMSwanson2015_KVsW.pdf

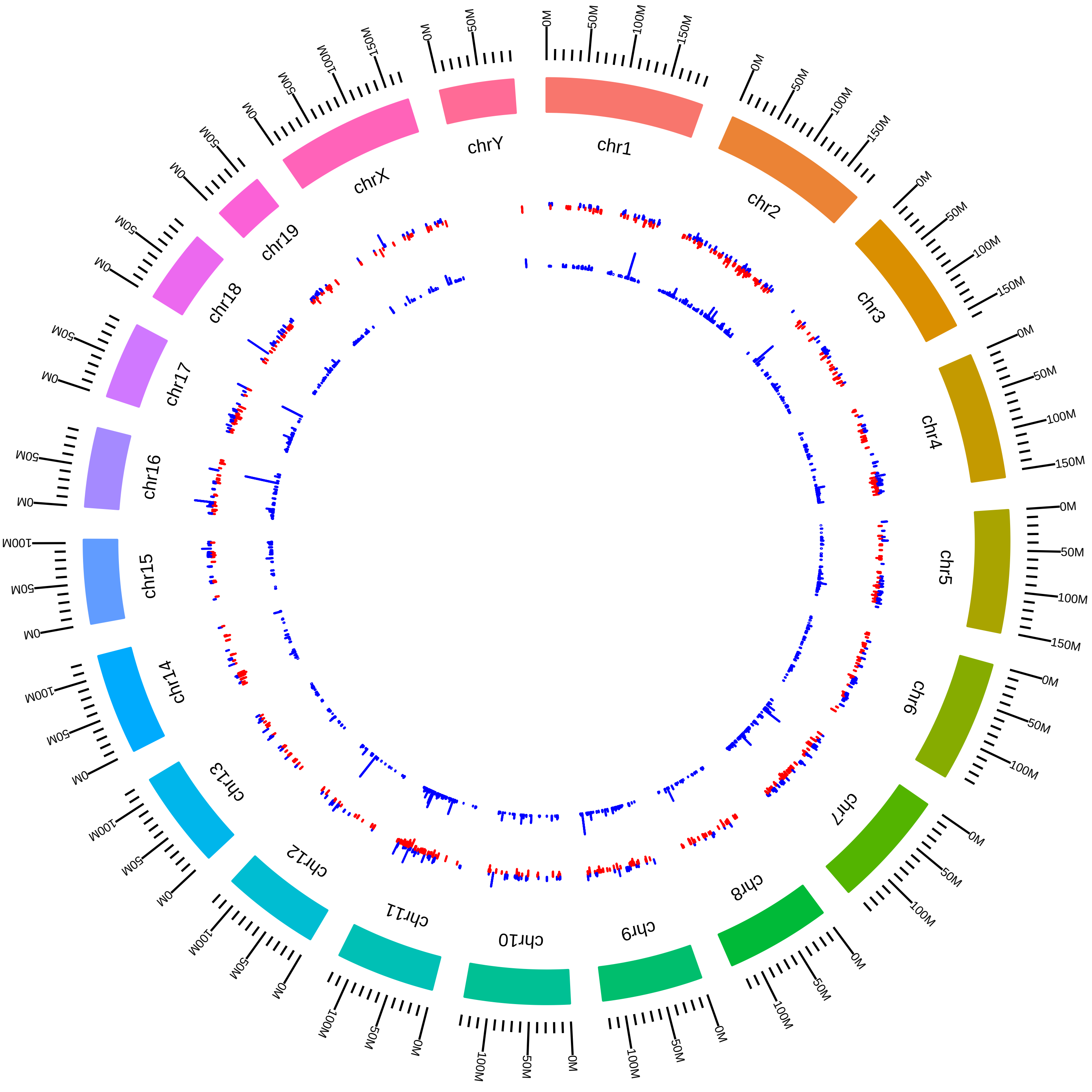

### rsfMbnl1MmCBurge_DEG_BKVsBW.pdf

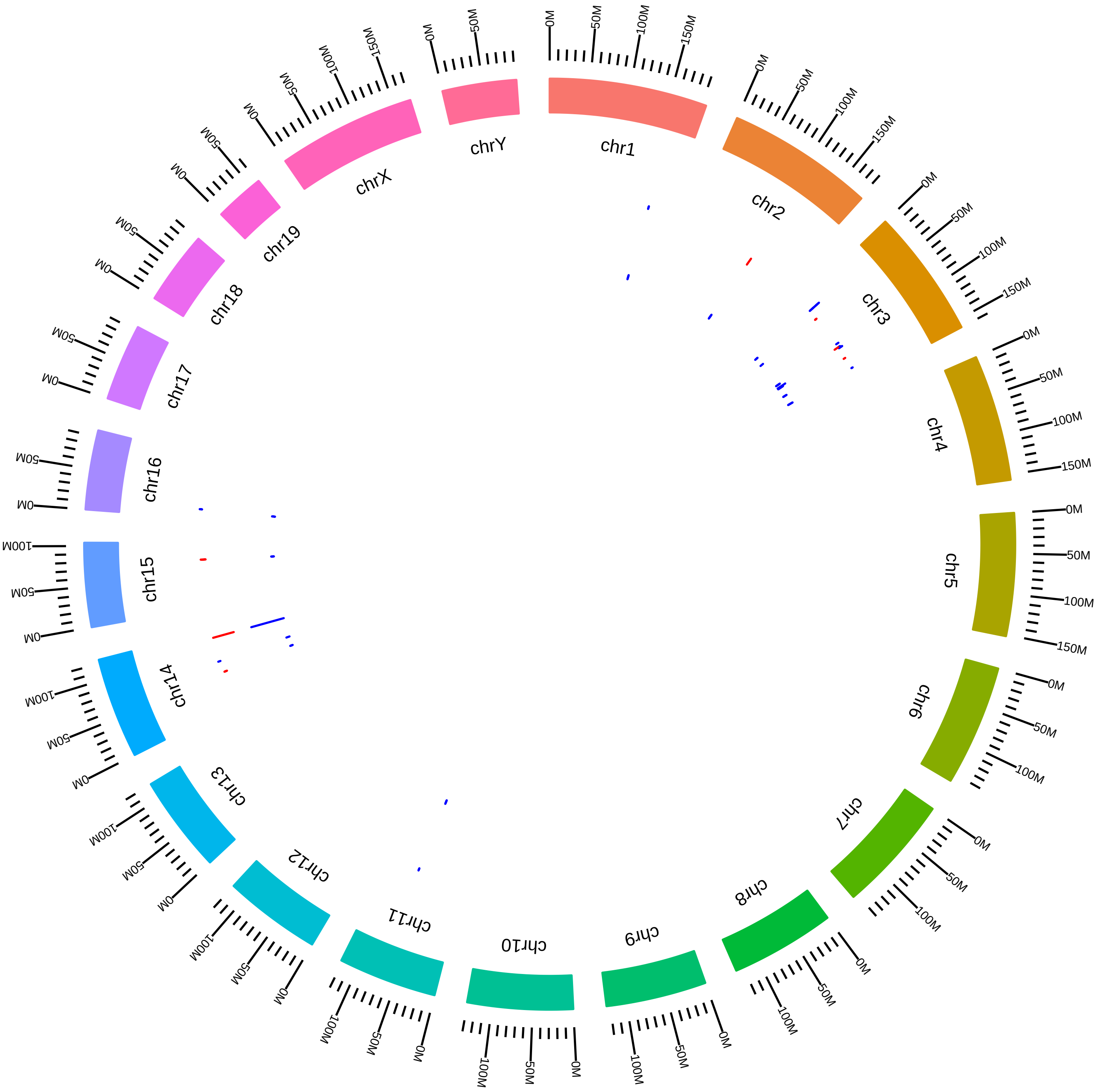

### rsfMbnl1MmCBurge_DEG_HKVsHW.pdf

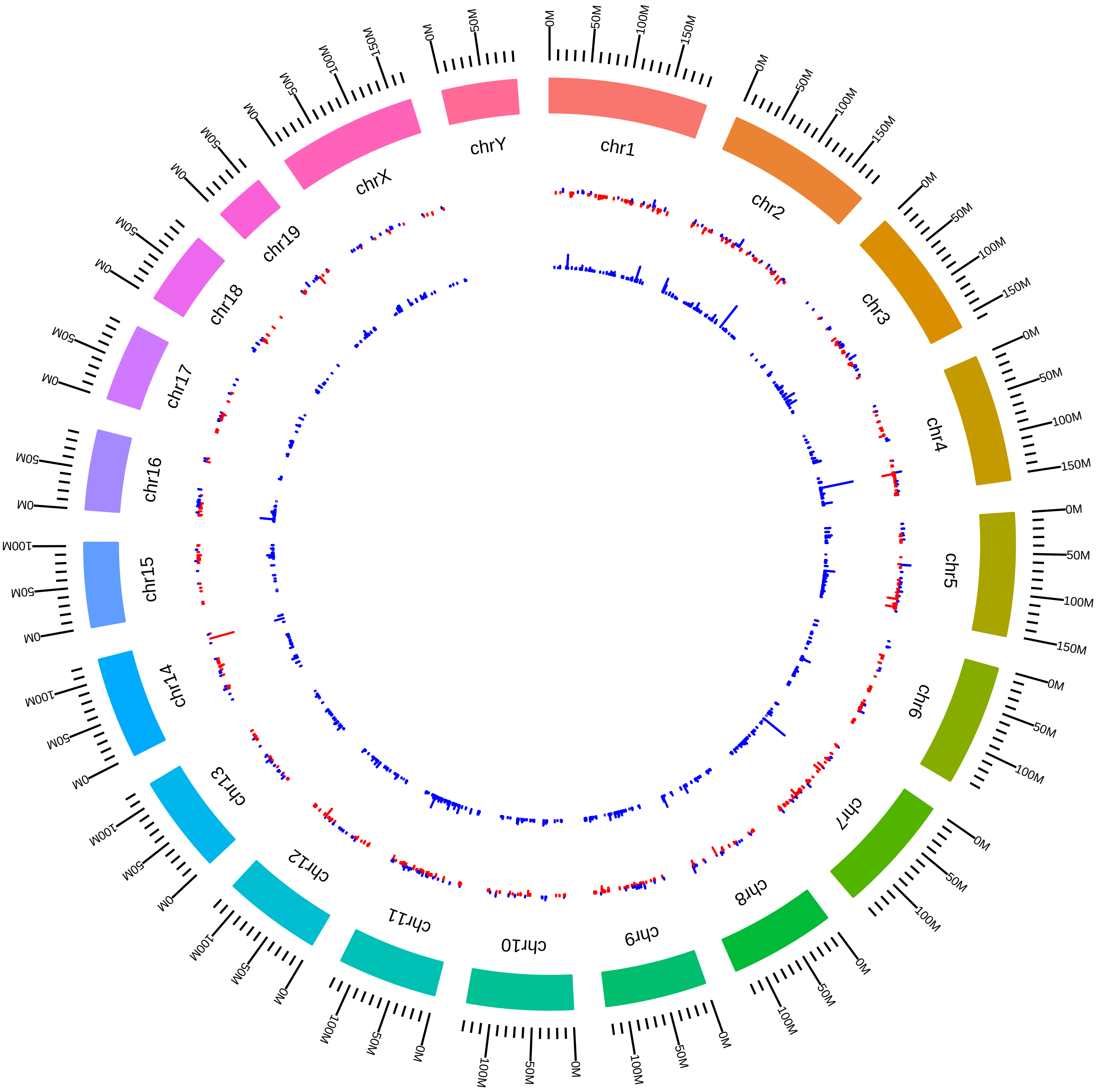

### rsfMbnl1MmCBurge_DEG_MKVsMW.pdf

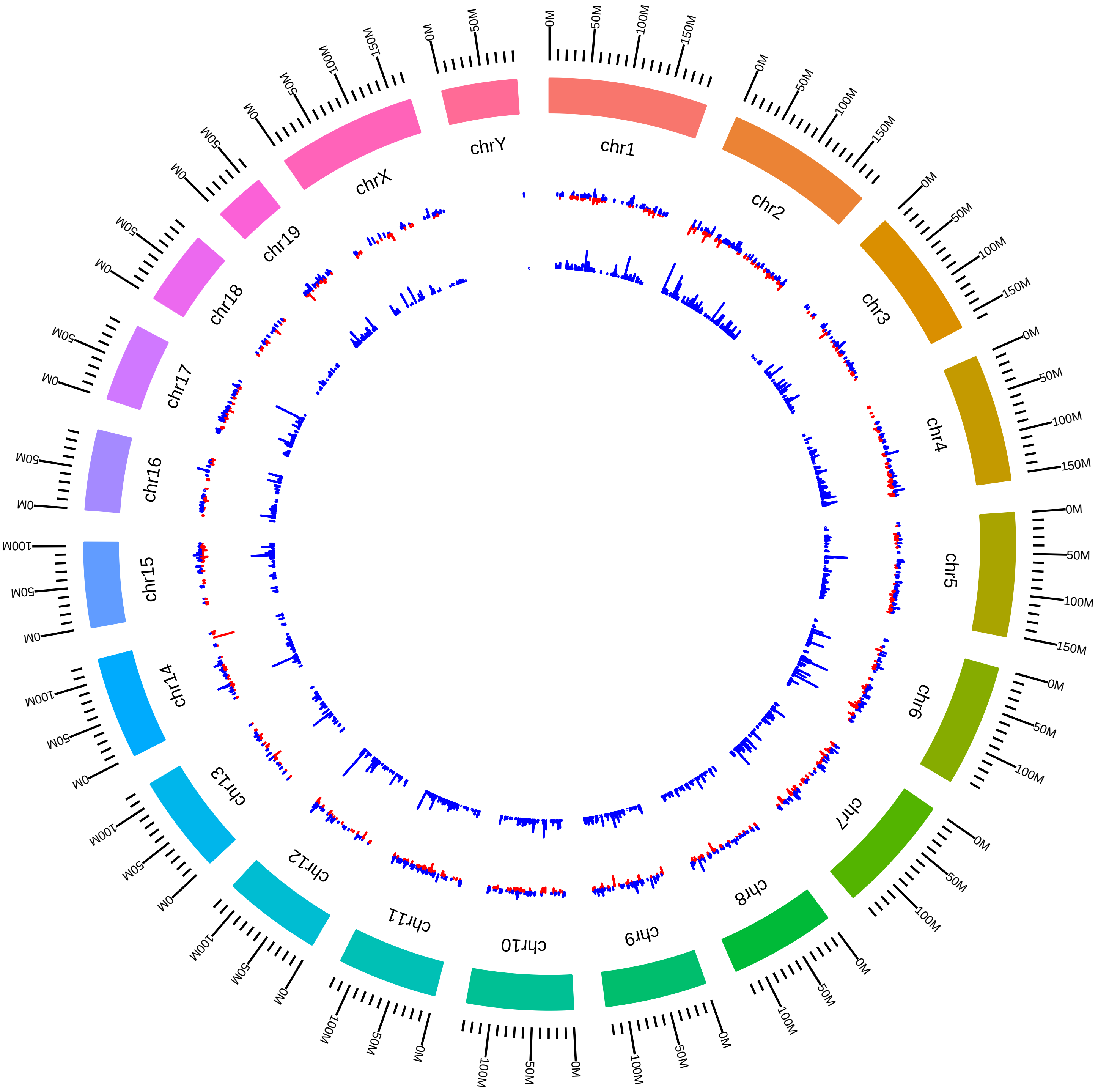

### rsfMbnl2MmMSwanson_DEG_KVsW.pdf

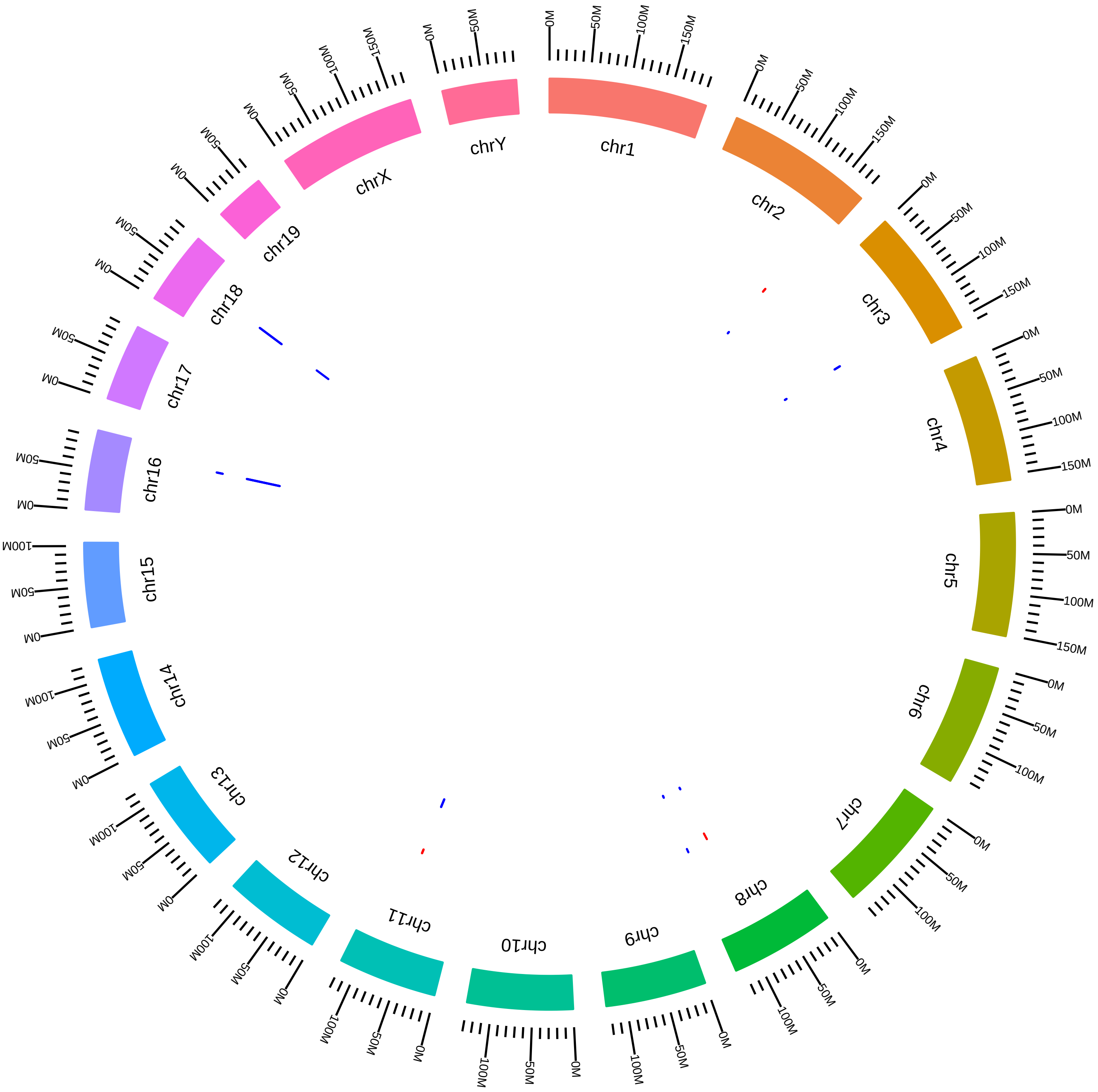

### rsfMettl3MmJHanna2015_BKVsBW.pdf

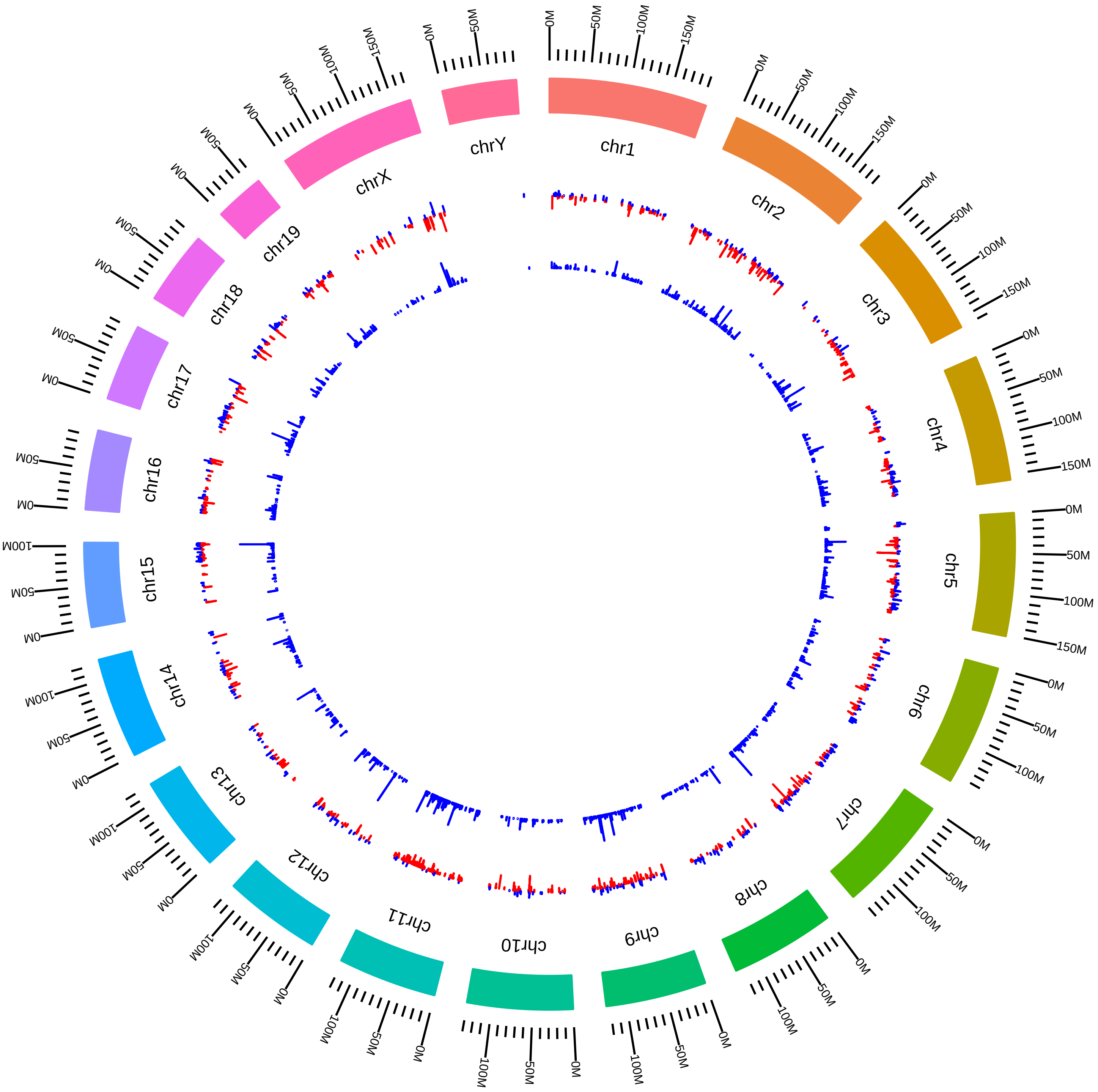

### rsfMettl3MmJHanna2015_SKVsSW.pdf

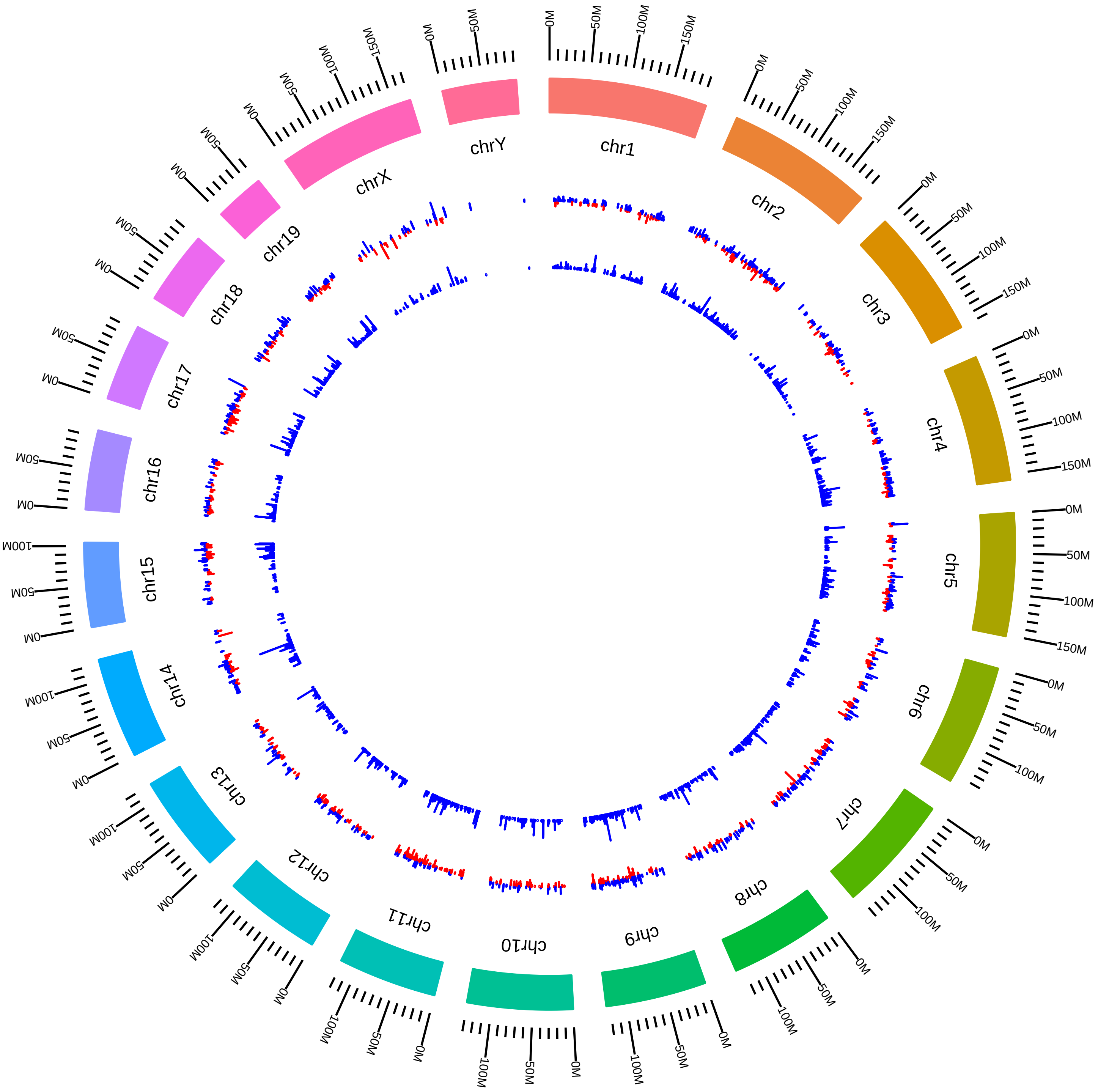

### rsfMettl3MmYunguiYang2014_DVsC.pdf

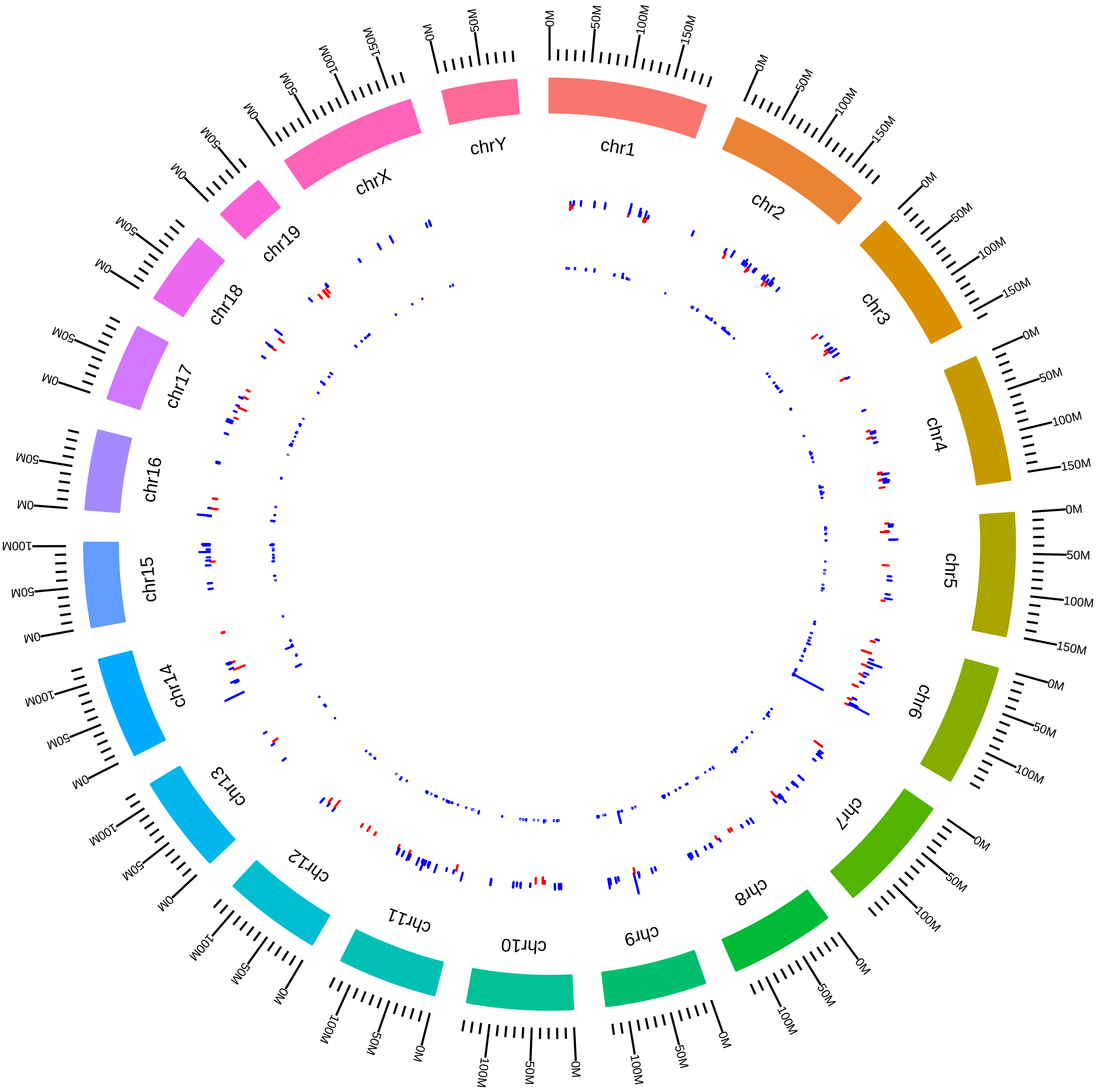

### rsfNonoMm2015SBrown2015_KVsW.pdf

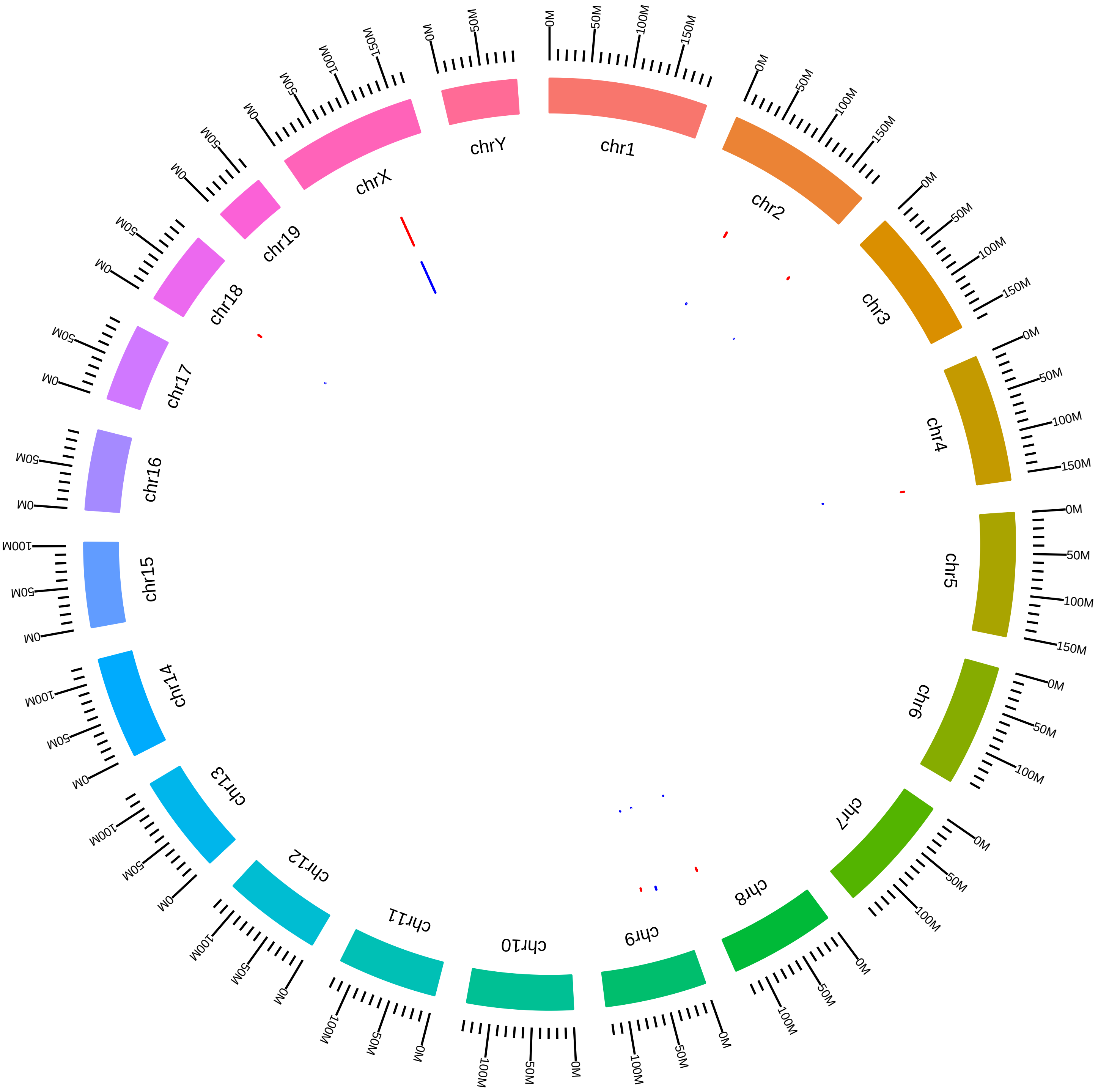

### rsfNova1Nova2MmRDarnell_DEG_B1KVsB1W.pdf

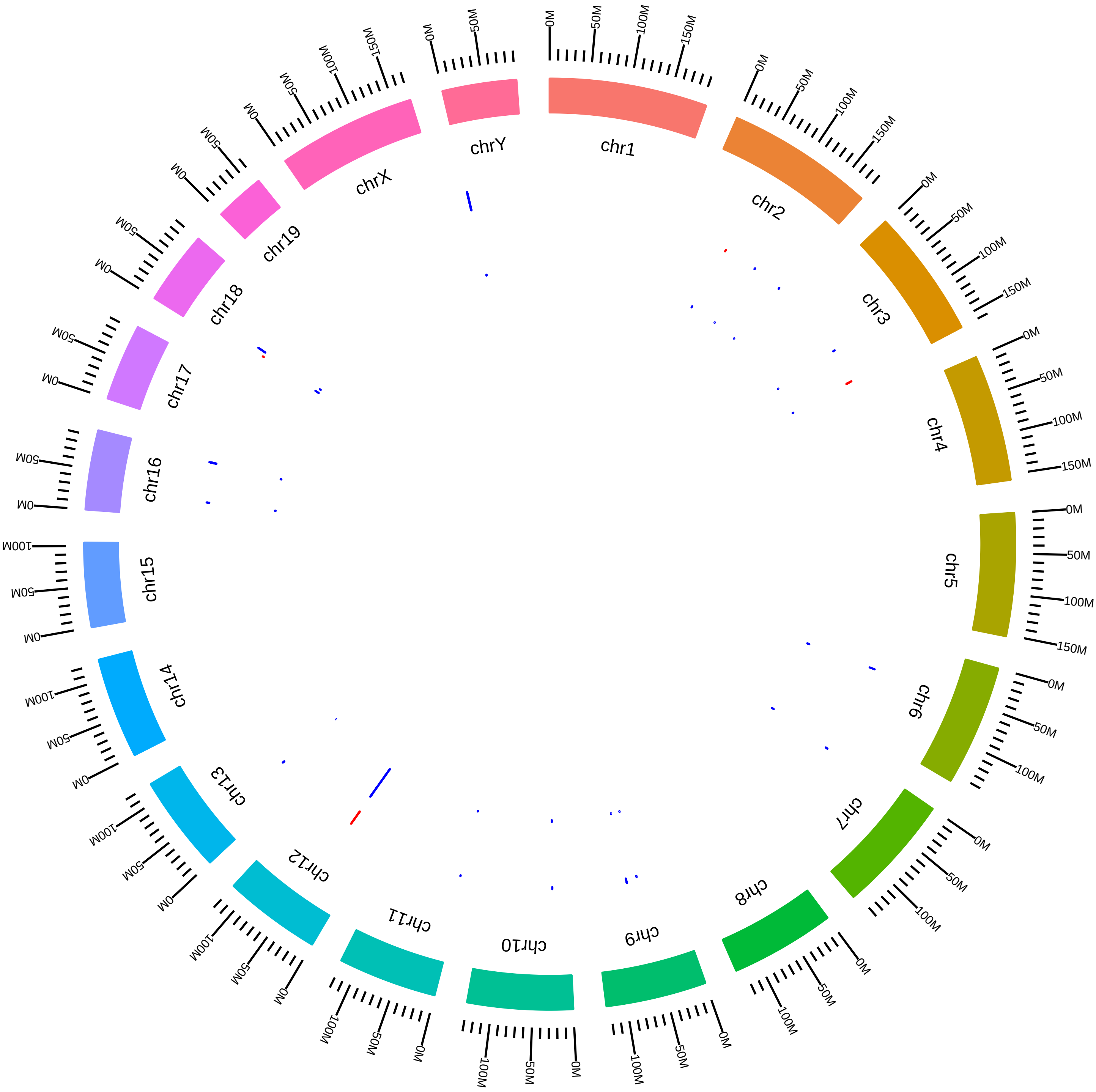

### rsfNova1Nova2MmRDarnell_DEG_C1KVsC1W.pdf

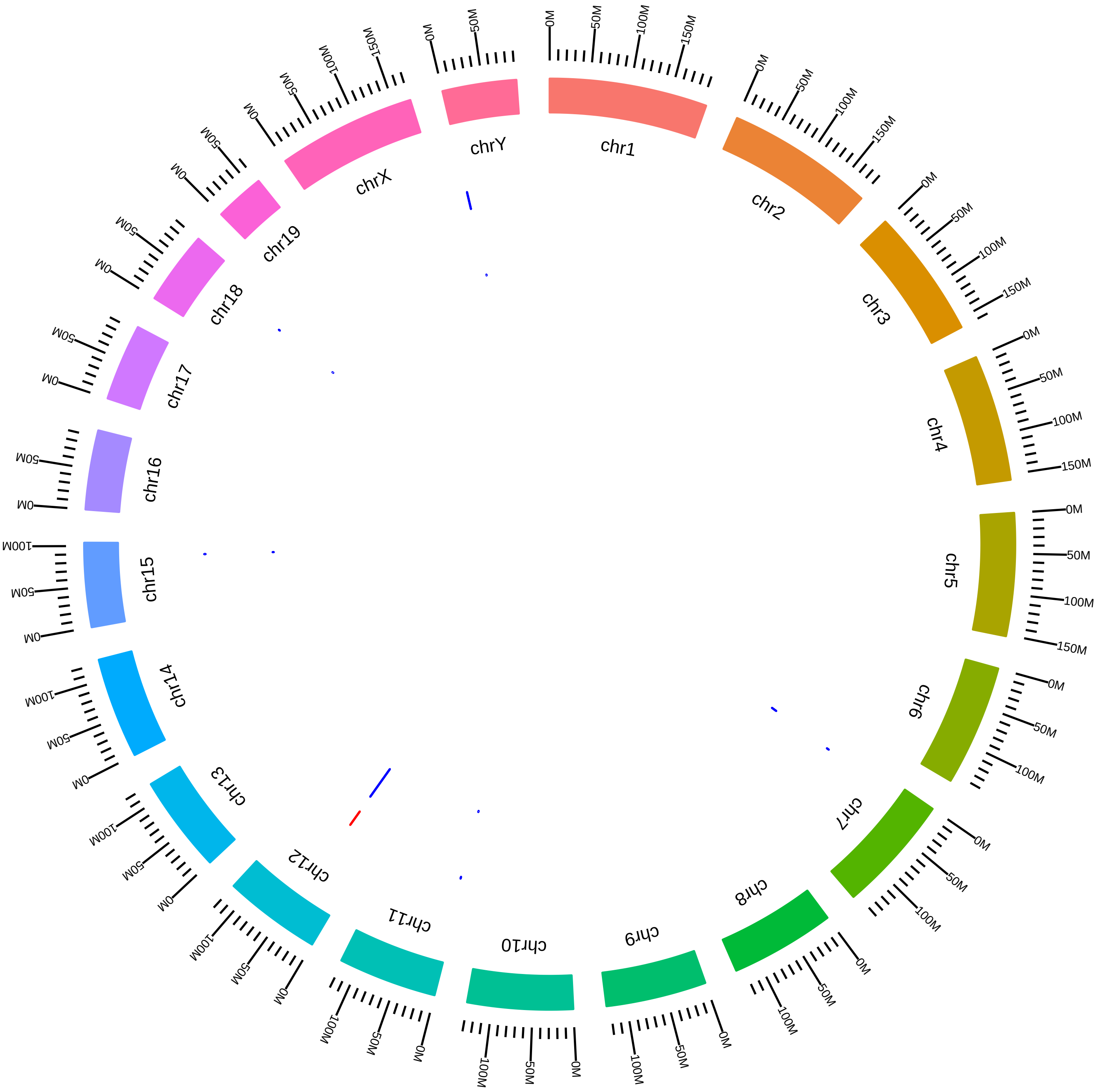

### rsfNova1Nova2MmRDarnell_DEG_C2KVsC2W.pdf

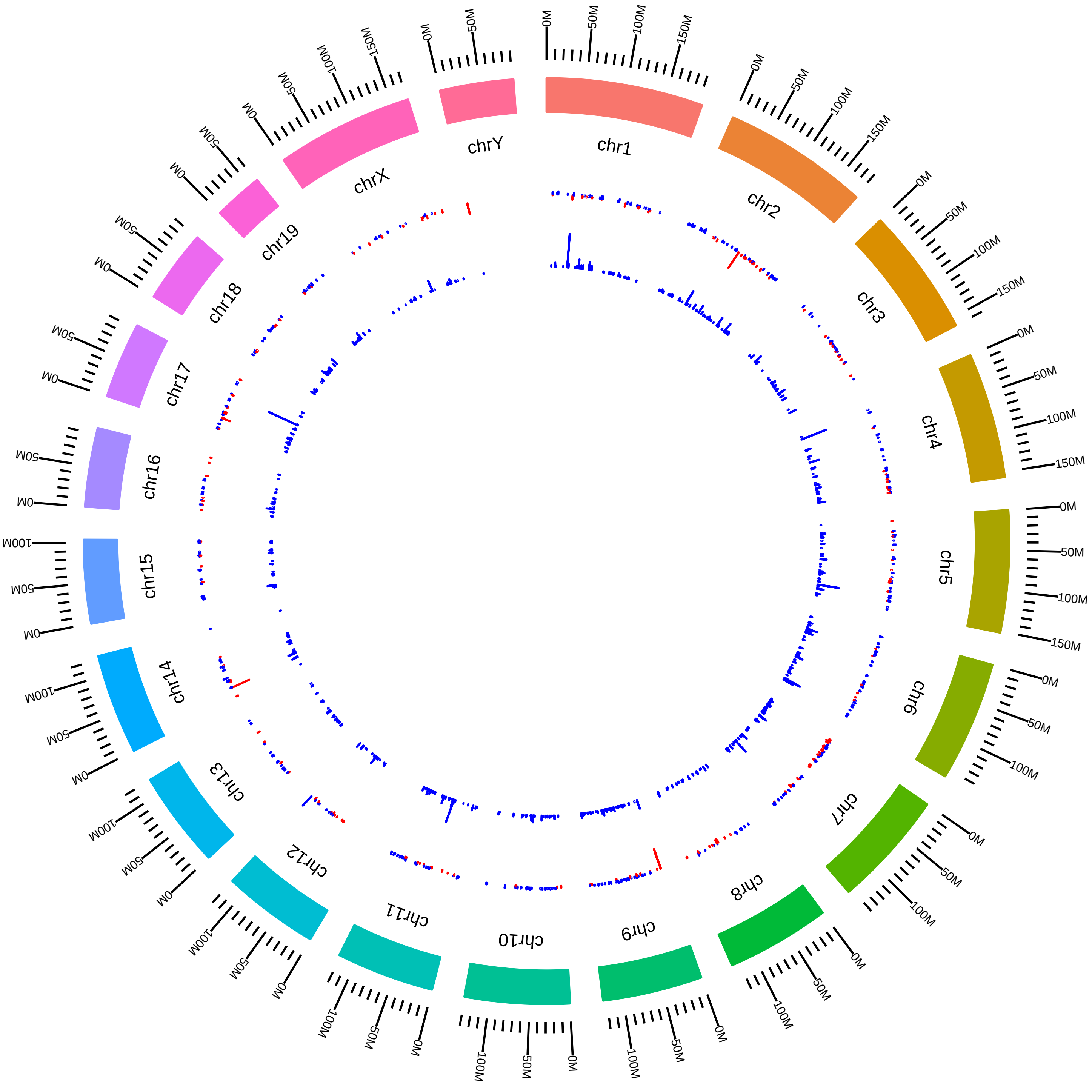

### rsfNova1Nova2MmRDavis_DEG_K1VsWT.pdf

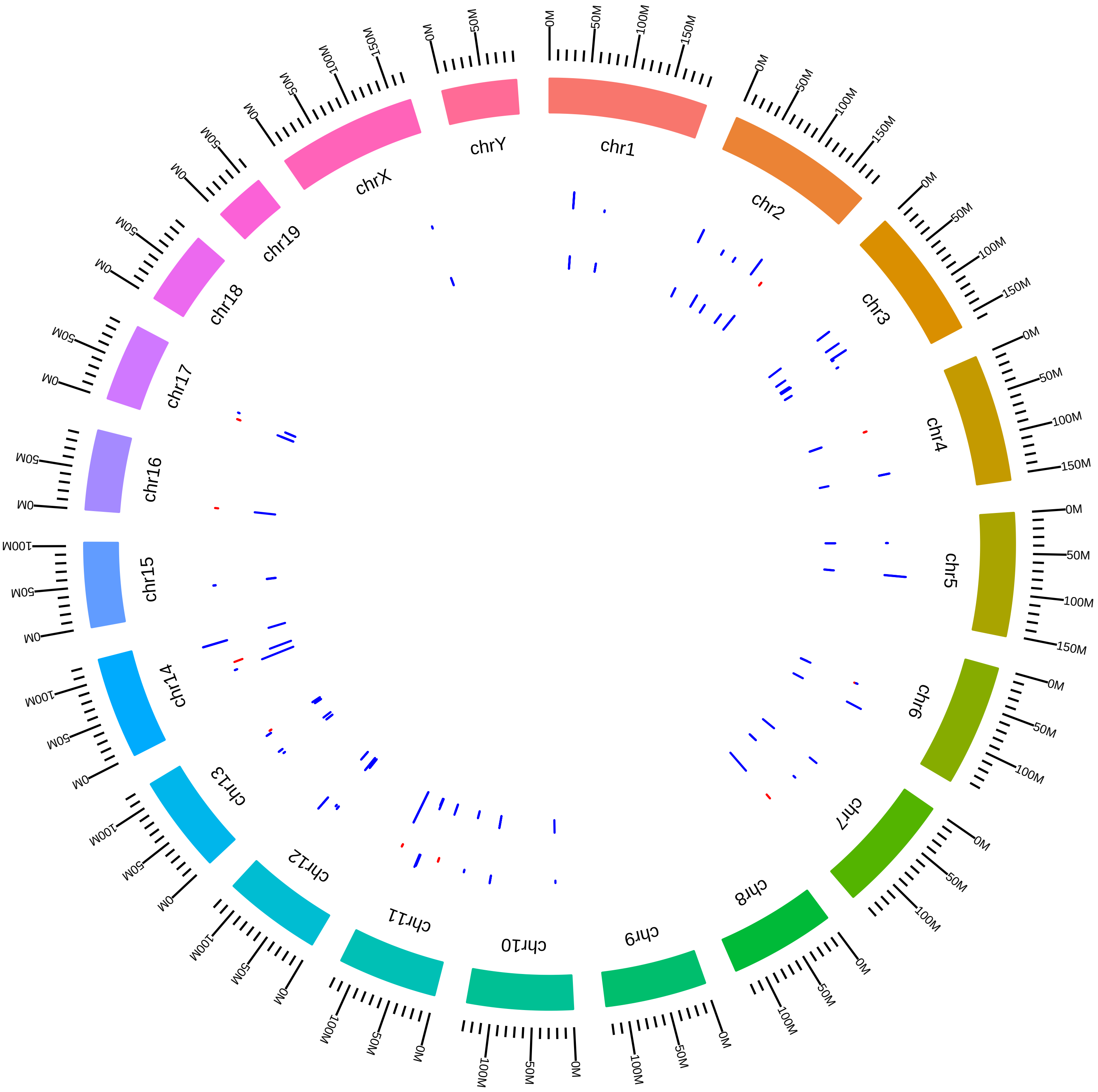

### rsfNova1Nova2MmRDavis_DEG_K2VsWT.pdf

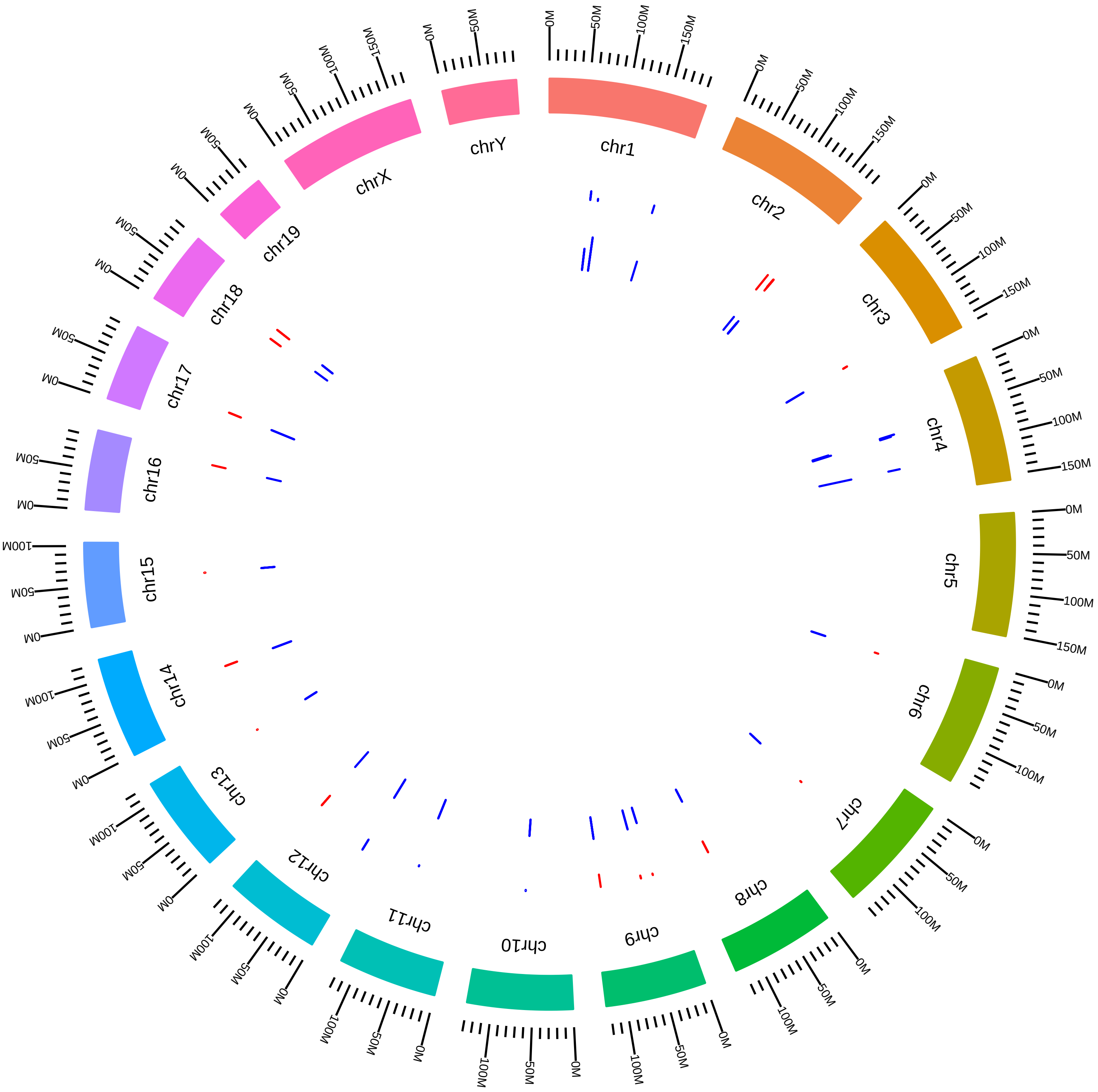

### rsfNova1Nova2MmRDavisIII_DEG_KVsW.pdf

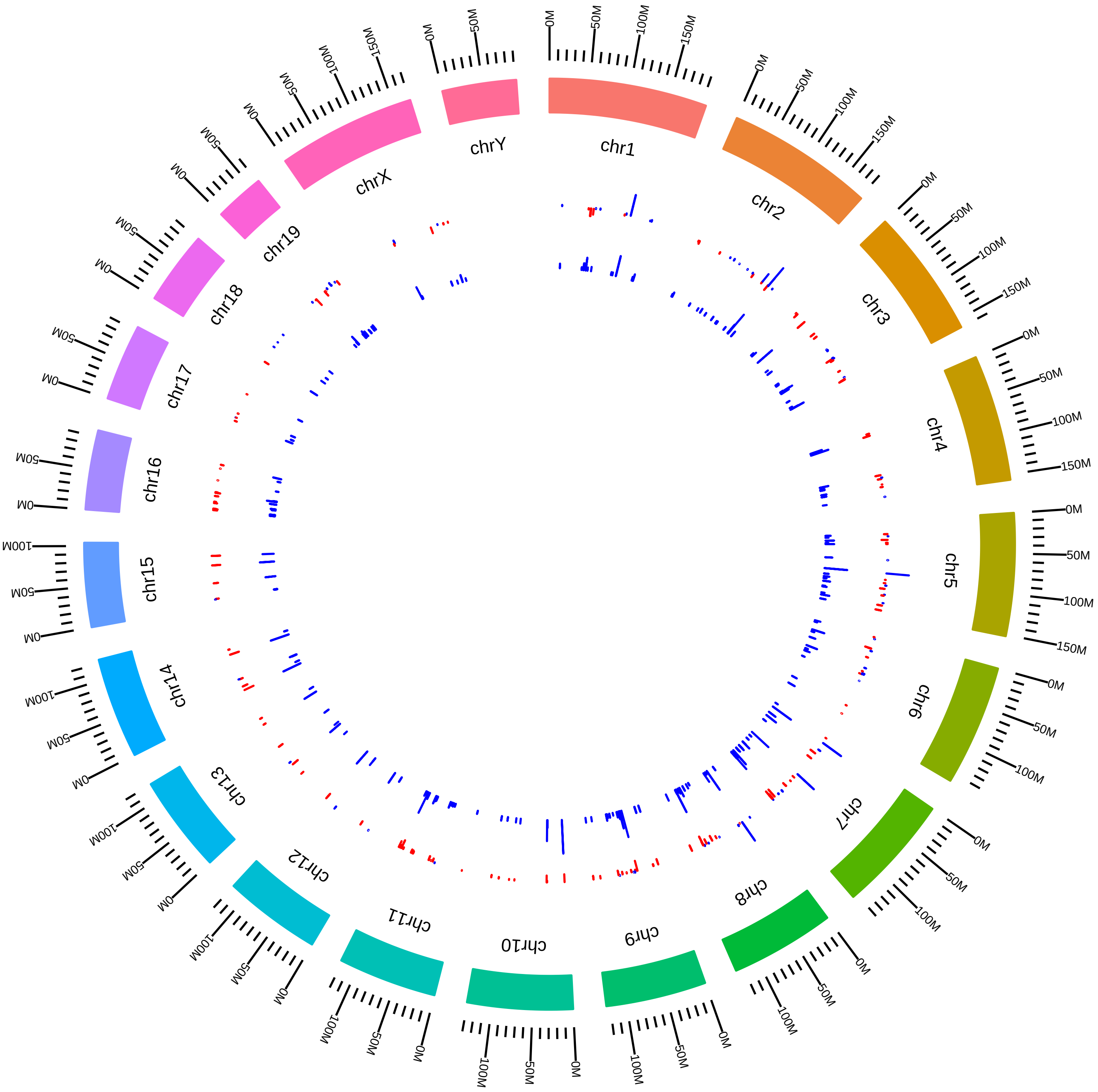

### rsfPhf5aPaf1MmIAifantis2016_AVsC.pdf

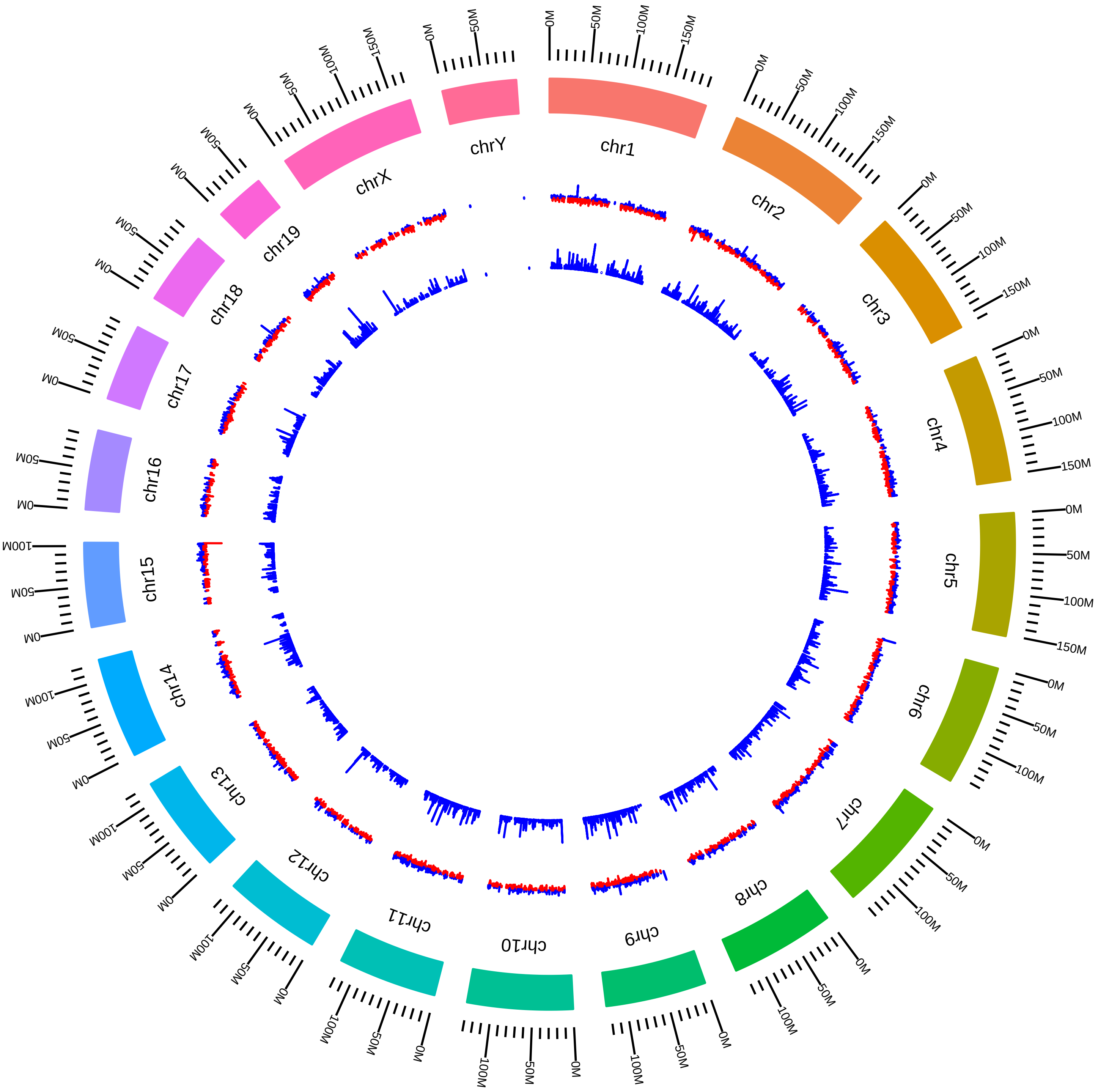

### rsfPhf5aPaf1MmIAifantis2016_HVsC.pdf

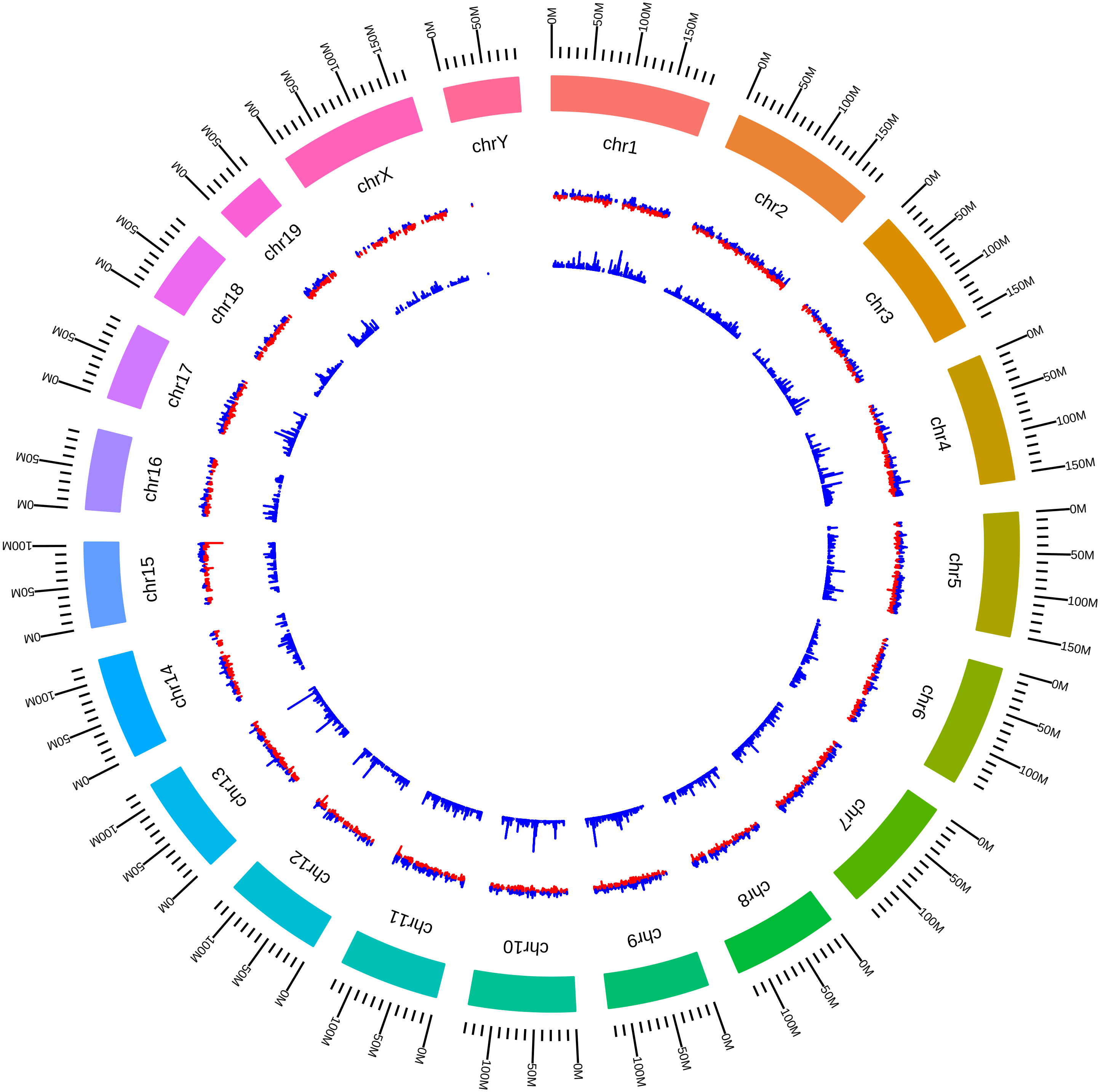

### rsfPqbp1MmPSilver_DEG_DVsW.pdf

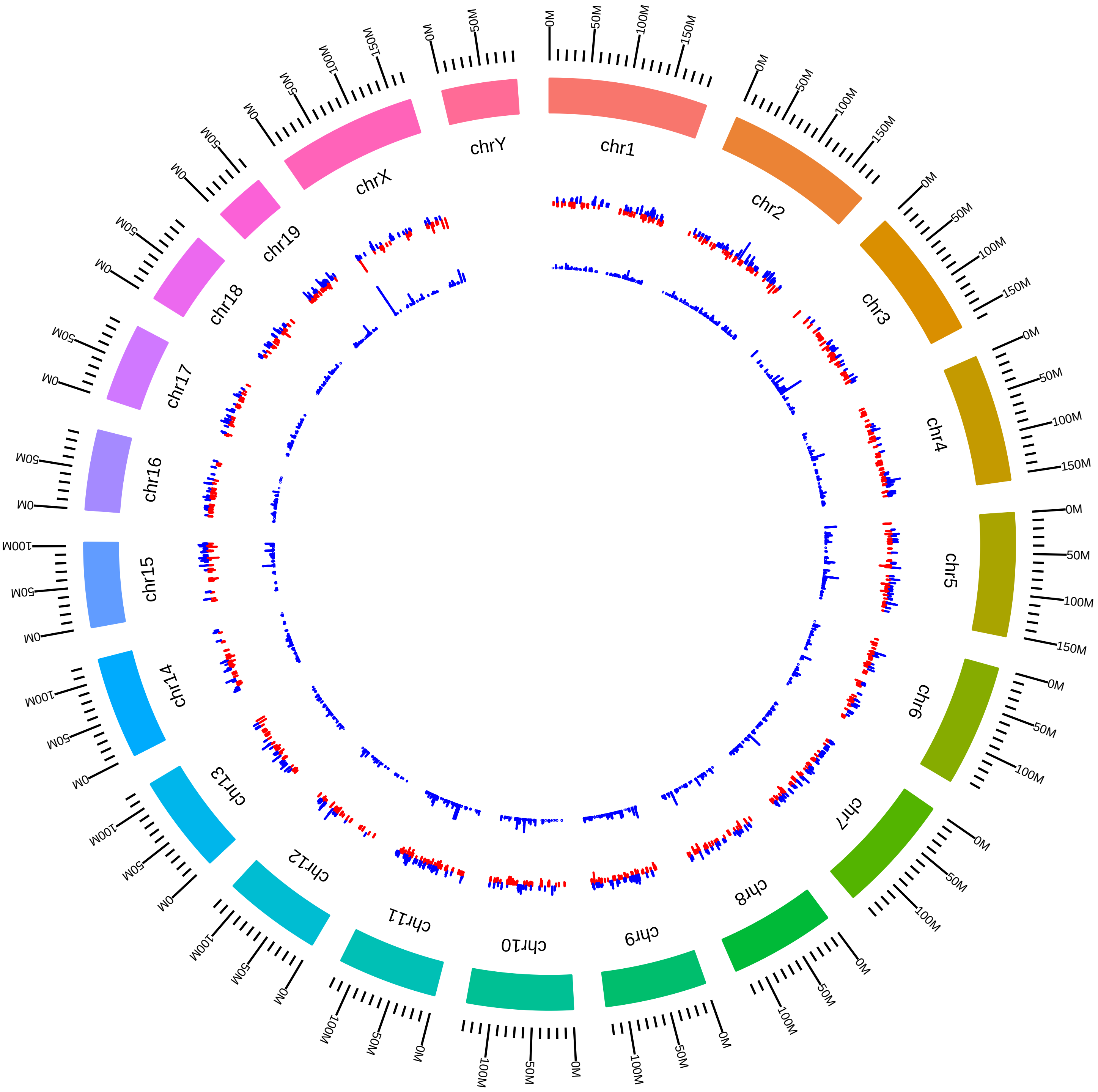

### rsfPrmt5MmATClark2015_DEG_KVsW.pdf

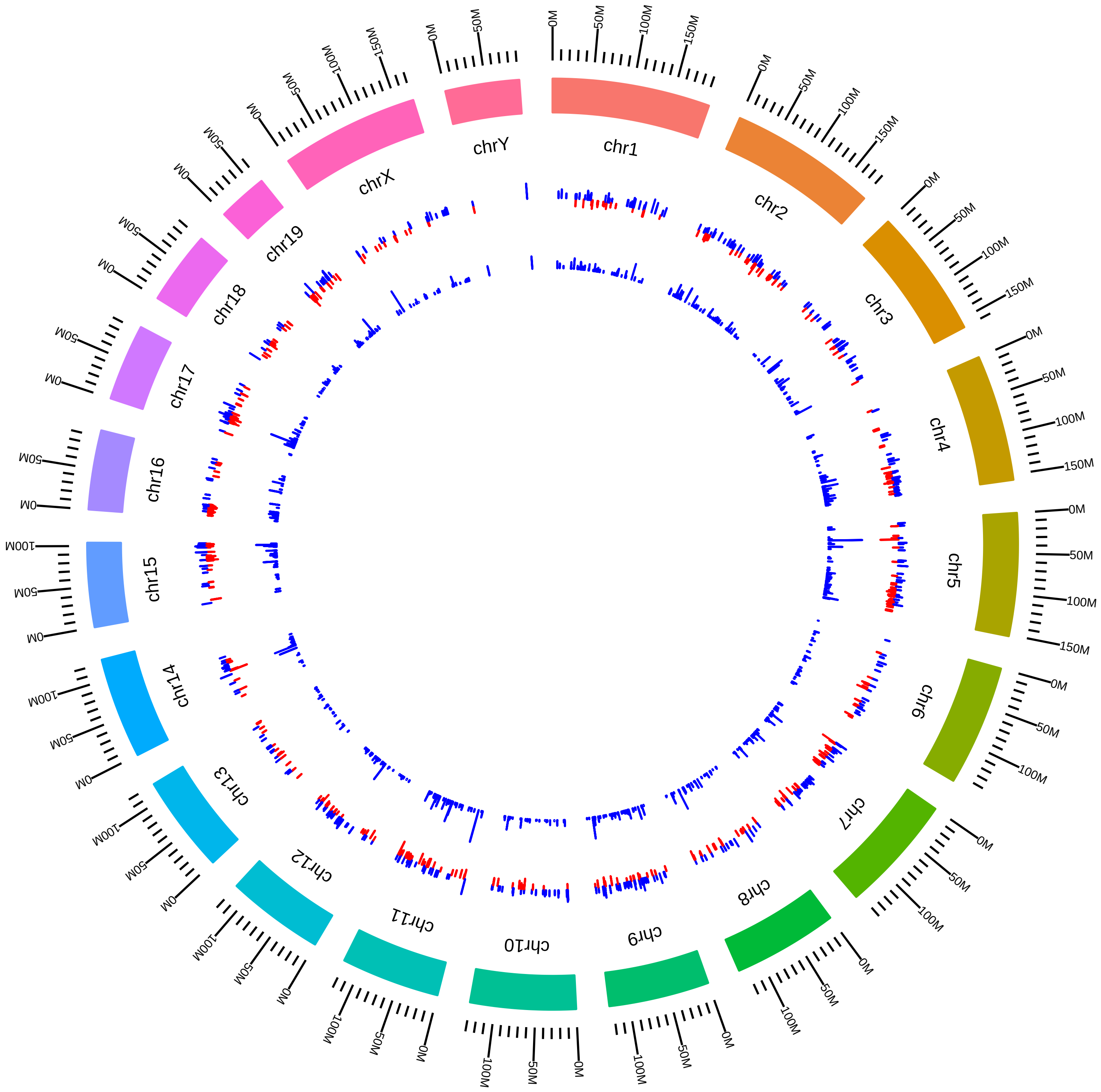

### rsfPrmt5MmEGuccione2015_KVsC.pdf

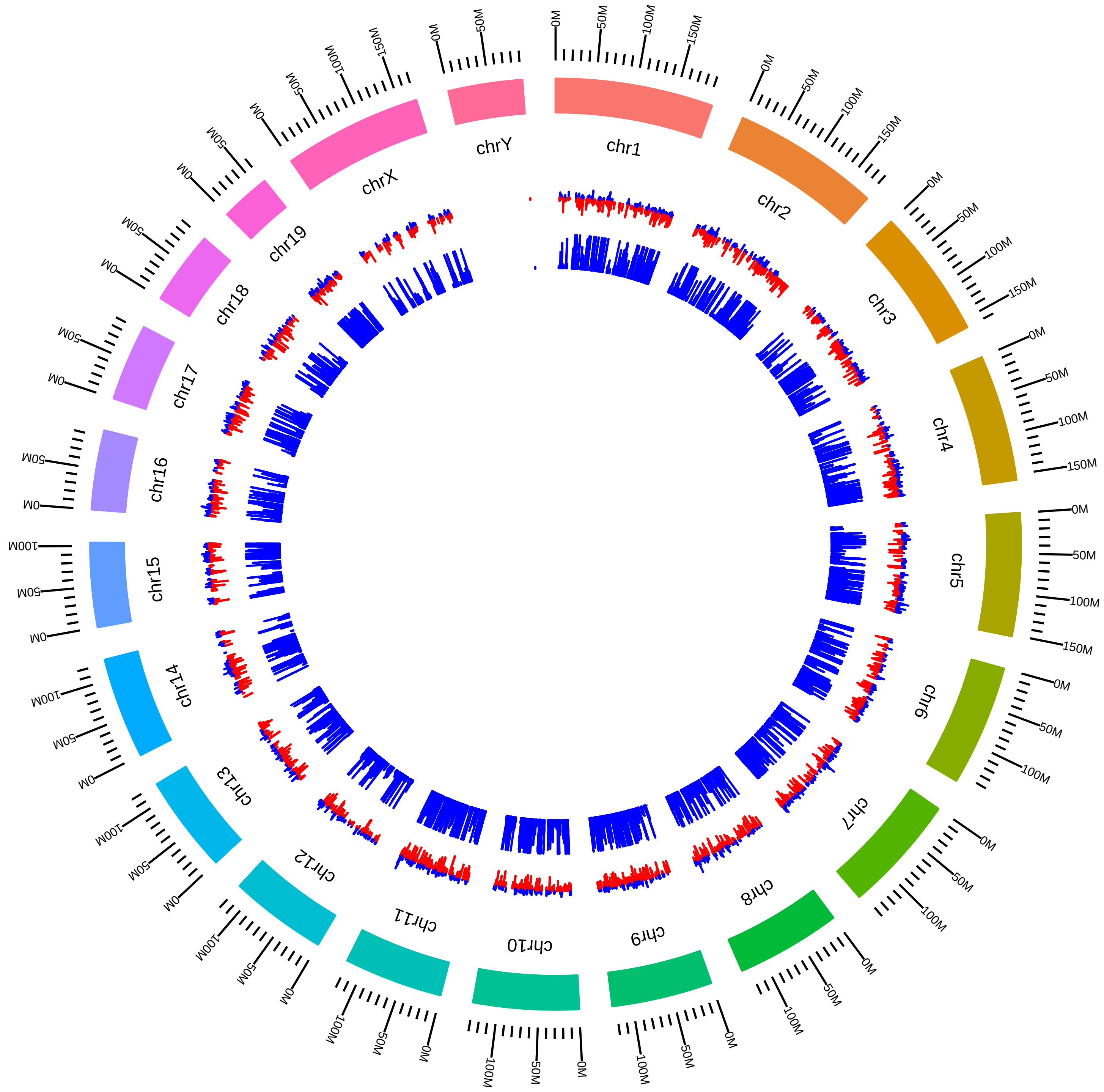

### rsfPrmt5MmEGuccione_DEG_MKVsMW.pdf

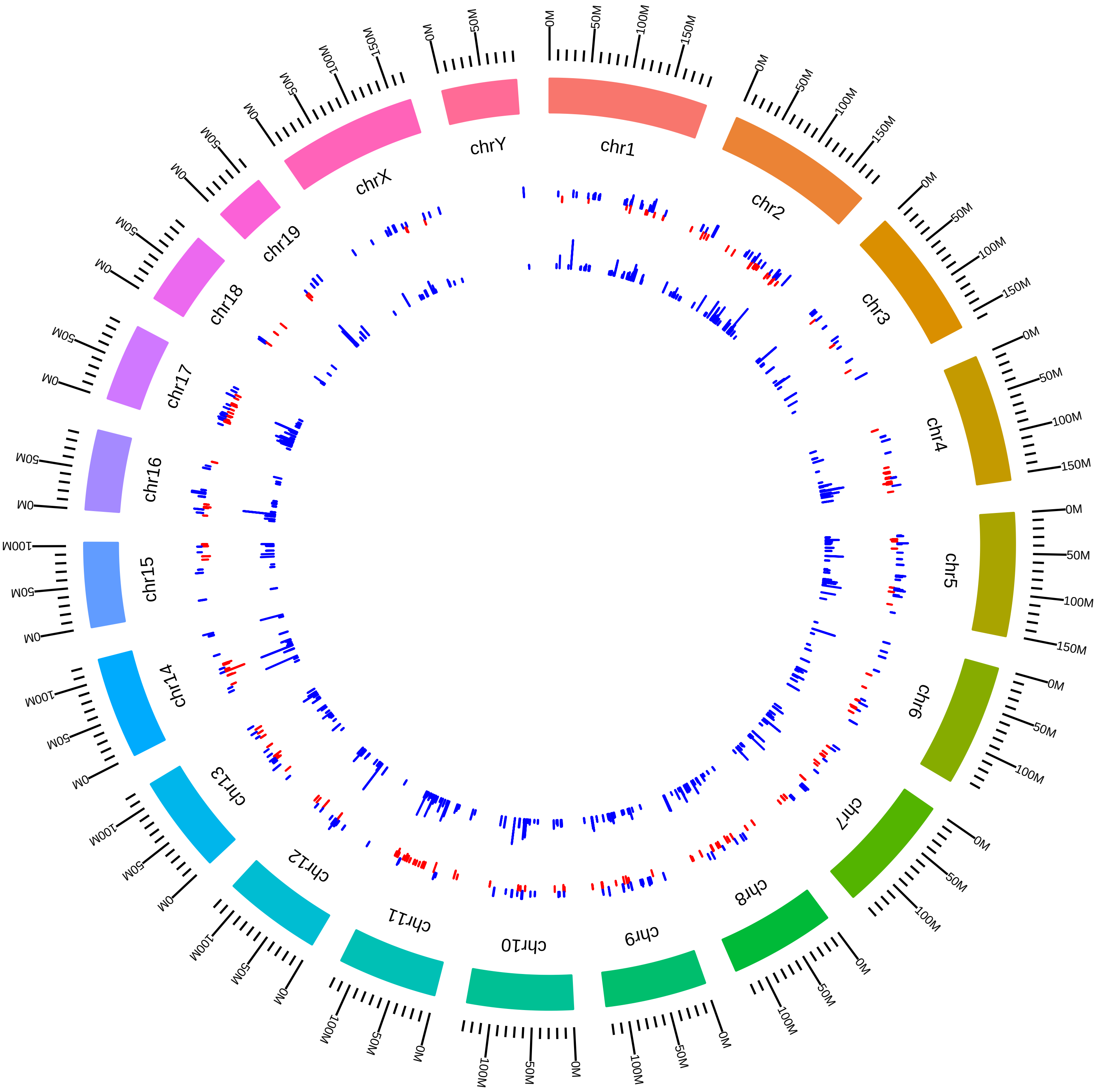

### rsfPrmt5MmEGuccione_DEG_NKVsNW.pdf

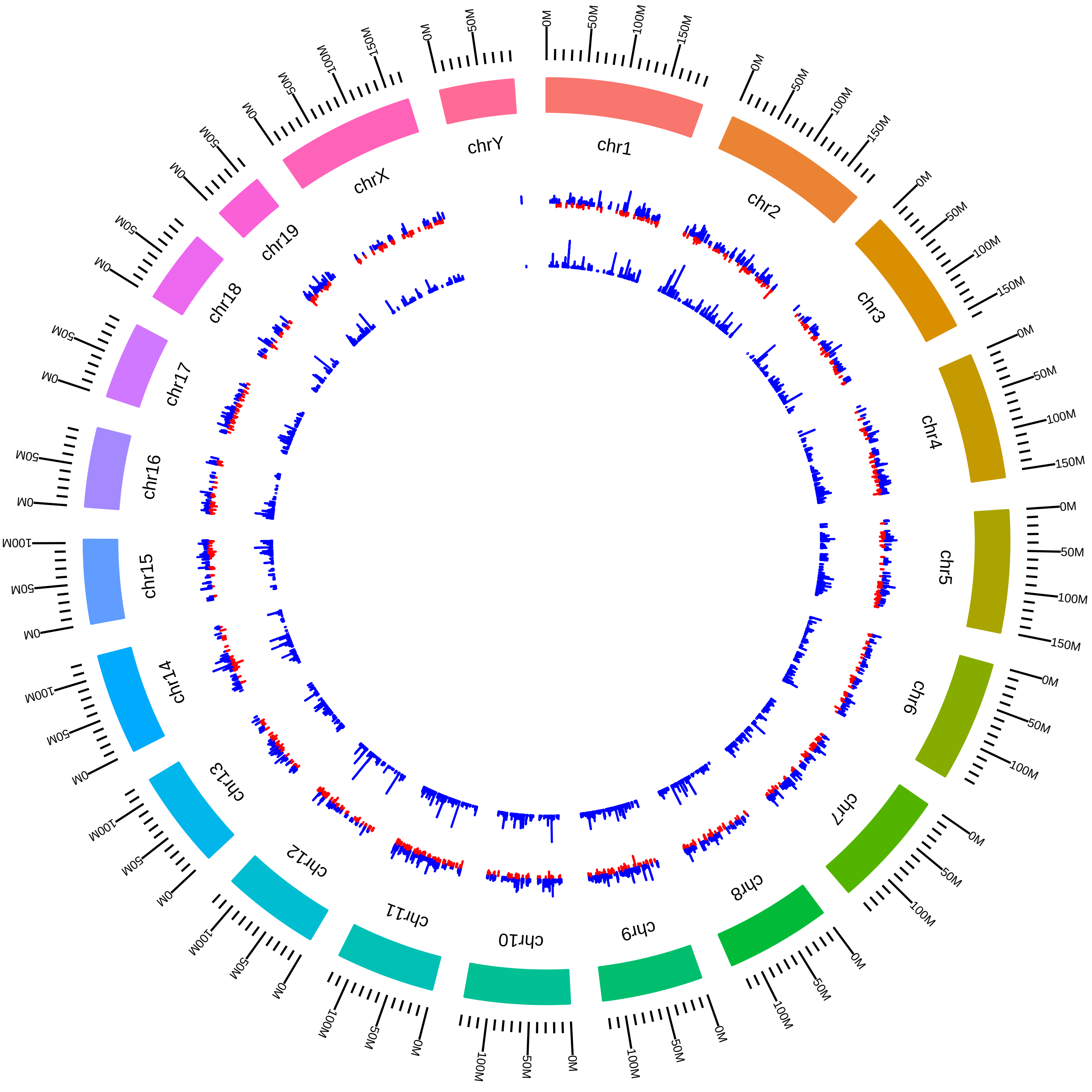

### rsfPrmt5MmJKim2015_KVsW.pdf

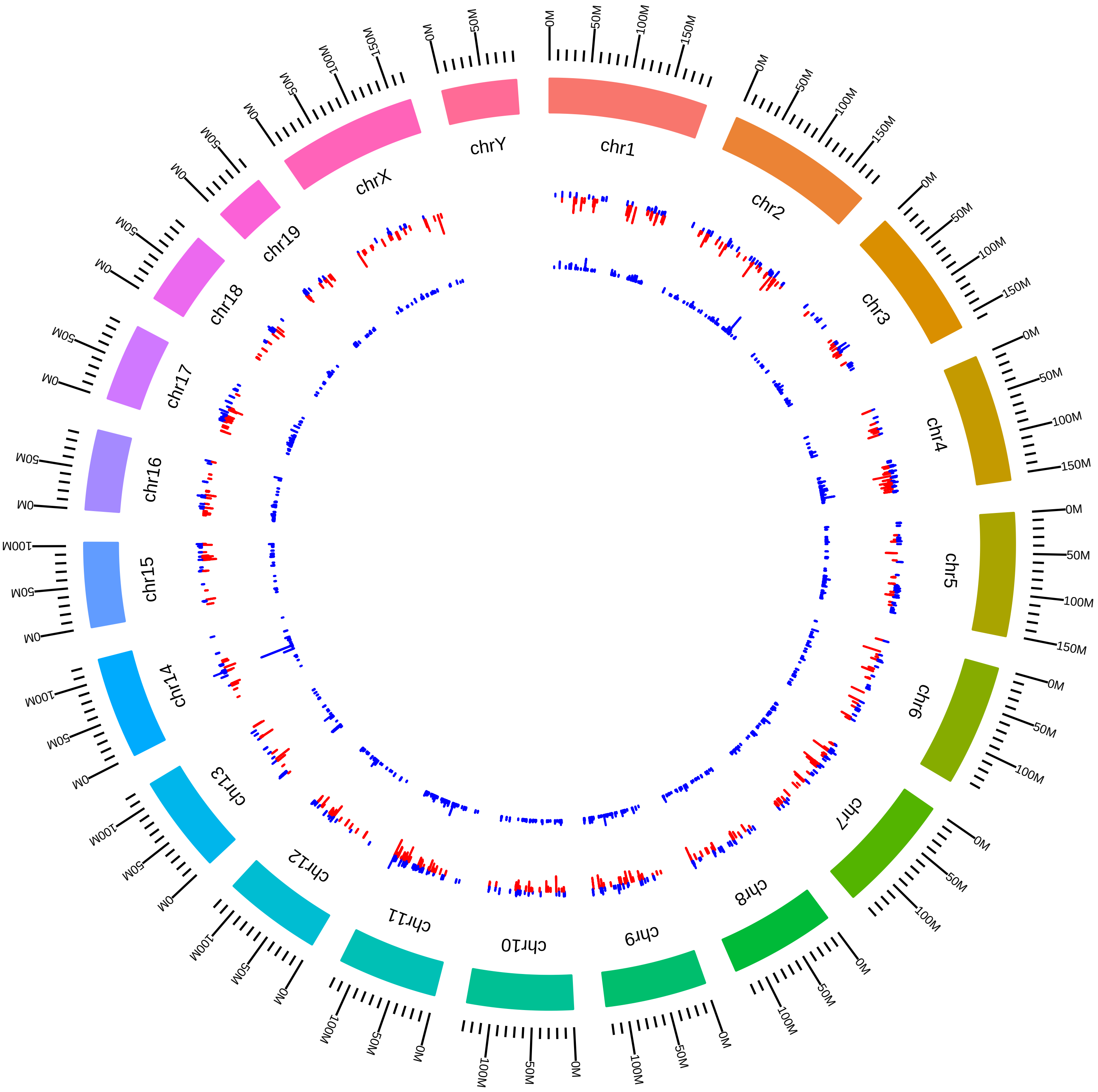

### rsfPrmt5MmSDNimer_DEG_KVsW.pdf

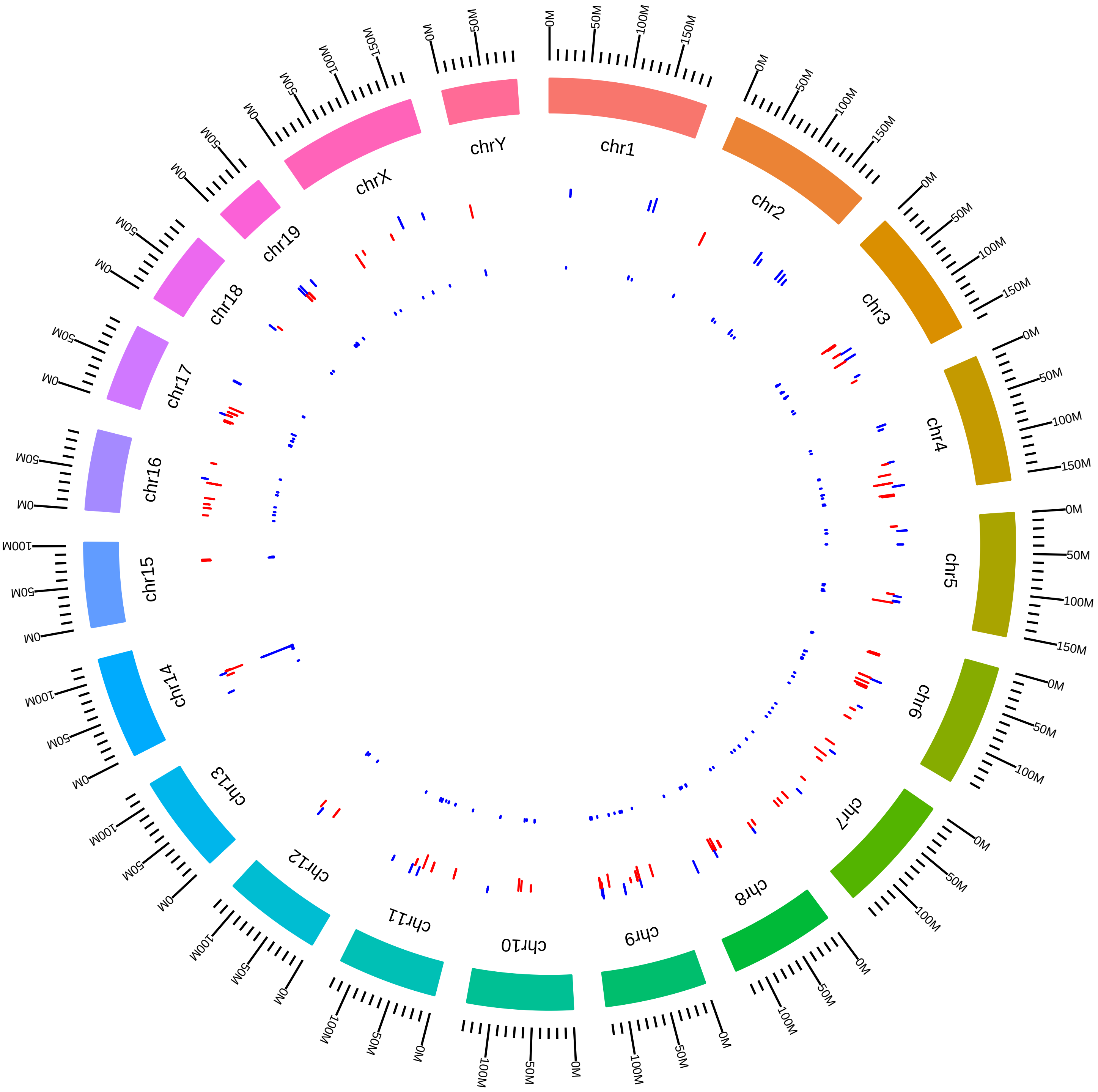

### rsfPrmt5MmSVokes2016_KVsW.pdf

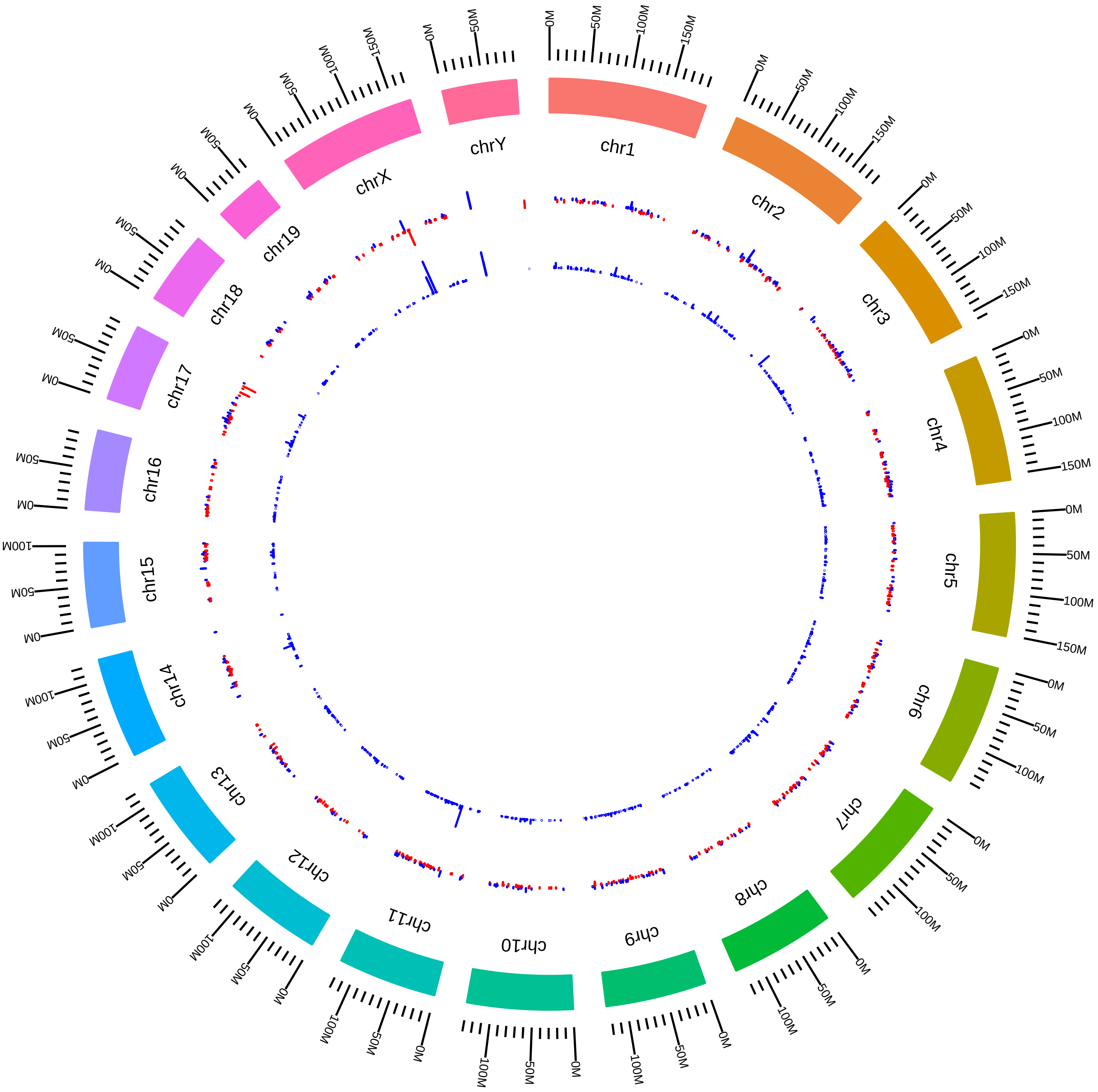

### rsfPrmt7MmSRichard2016_DEG_KVsW.pdf

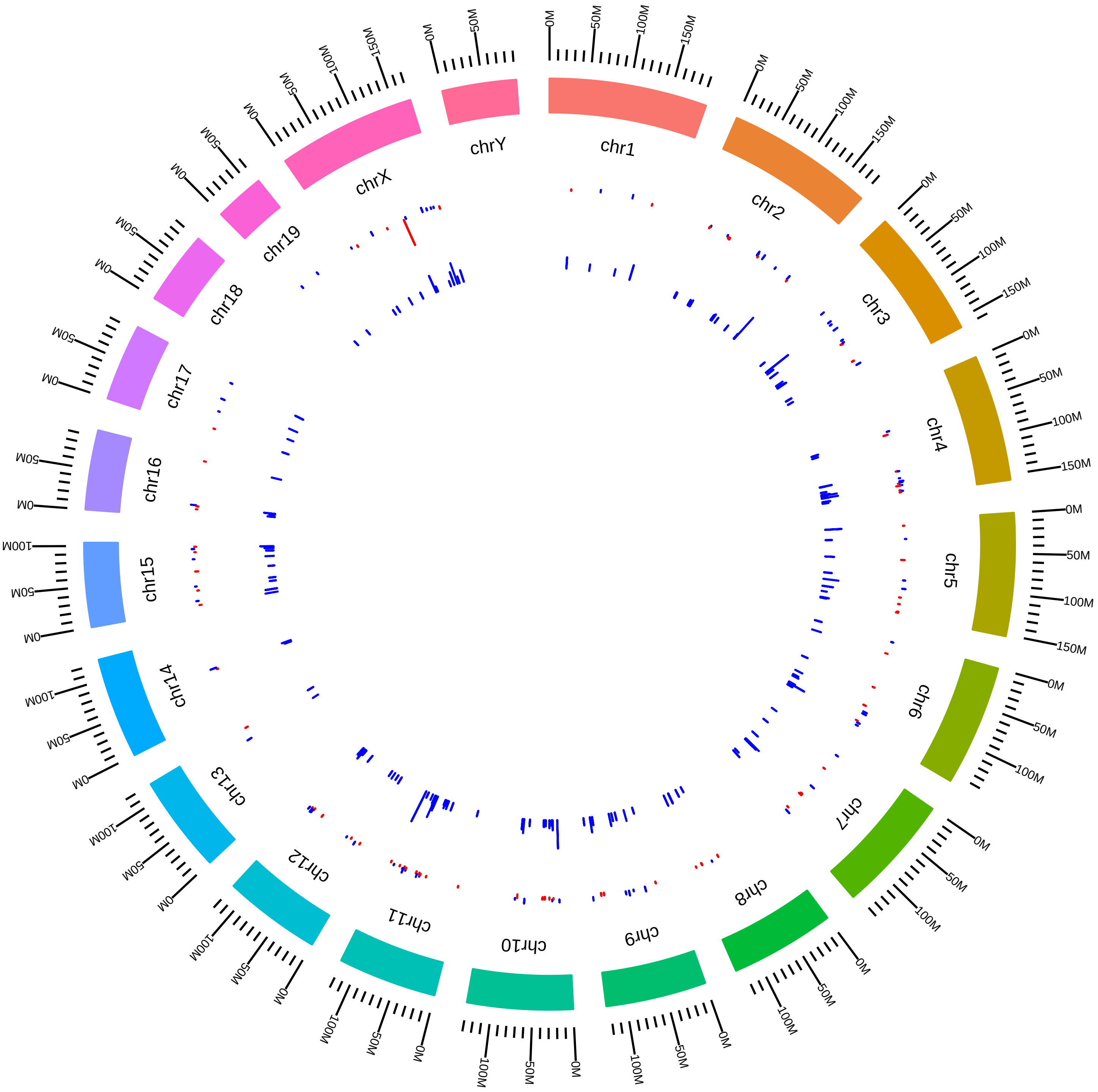

### rsfPtbp1MmDLim2015_DEG_DVsW.pdf

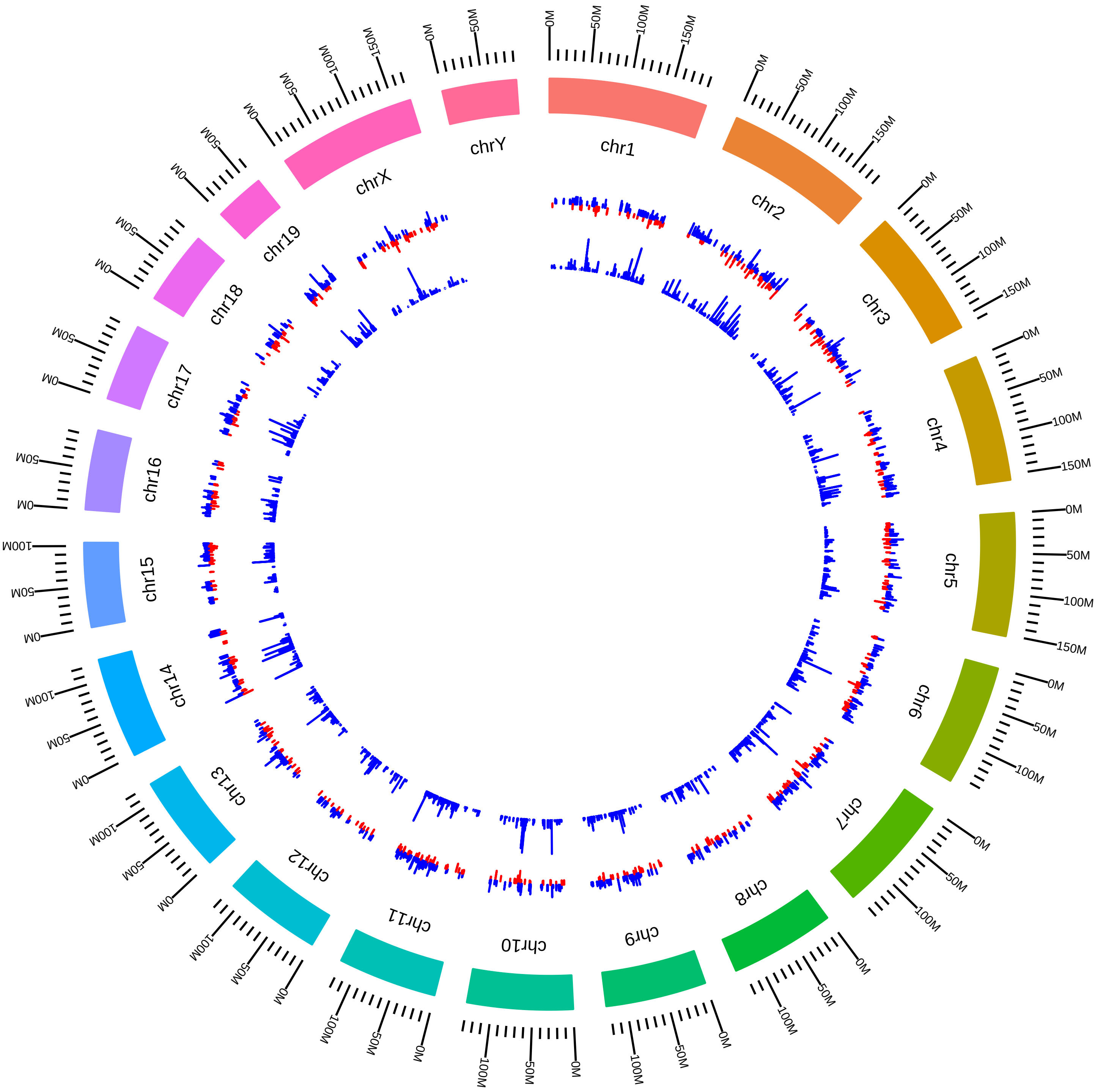

### rsfPtbp1Ptbp2MmDBlackmESC2015_DEG_DVsW.pdf

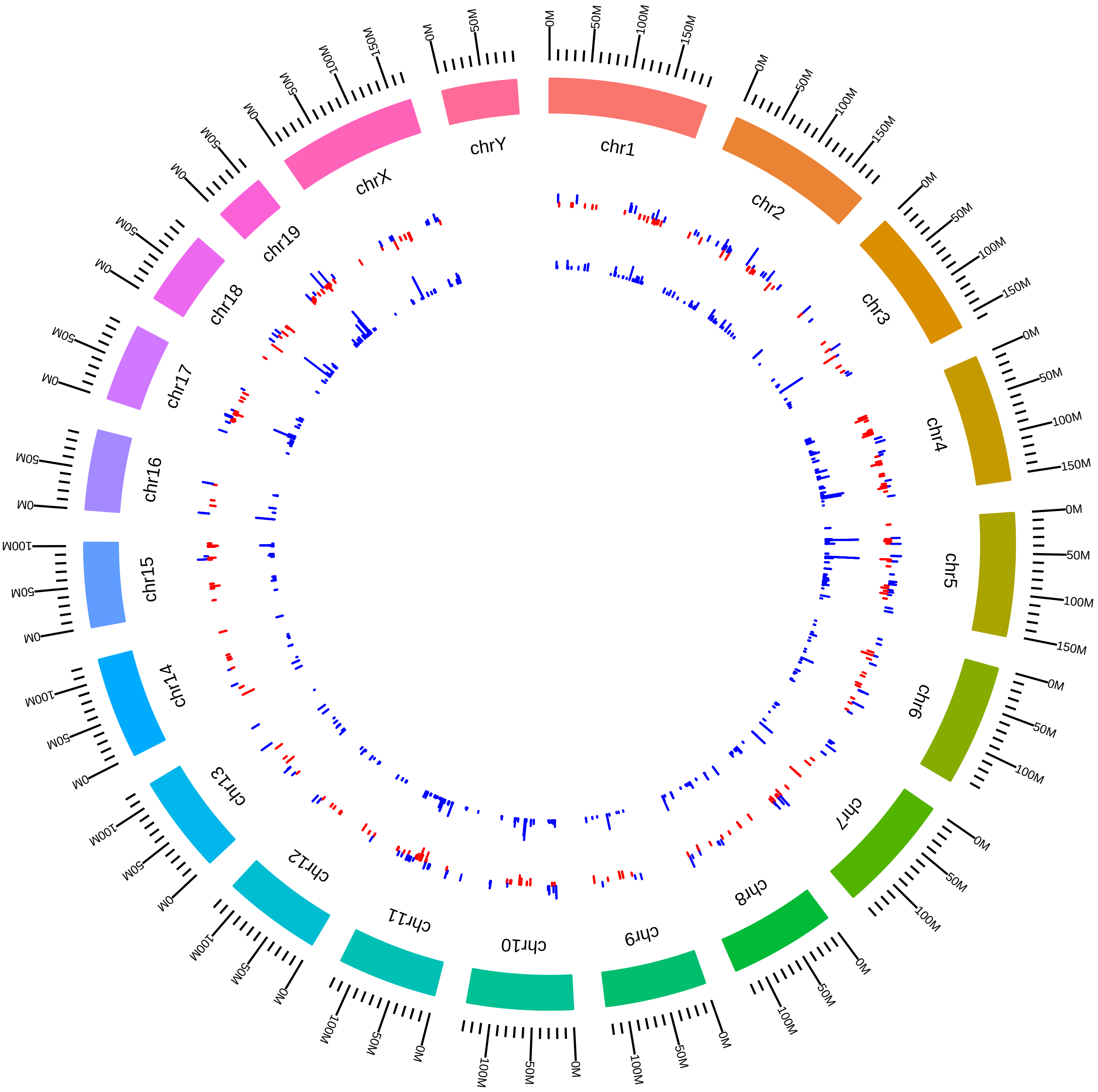

### rsfPtbp1Ptbp2MmDBlackmESC2015_DEG_KVsW.pdf

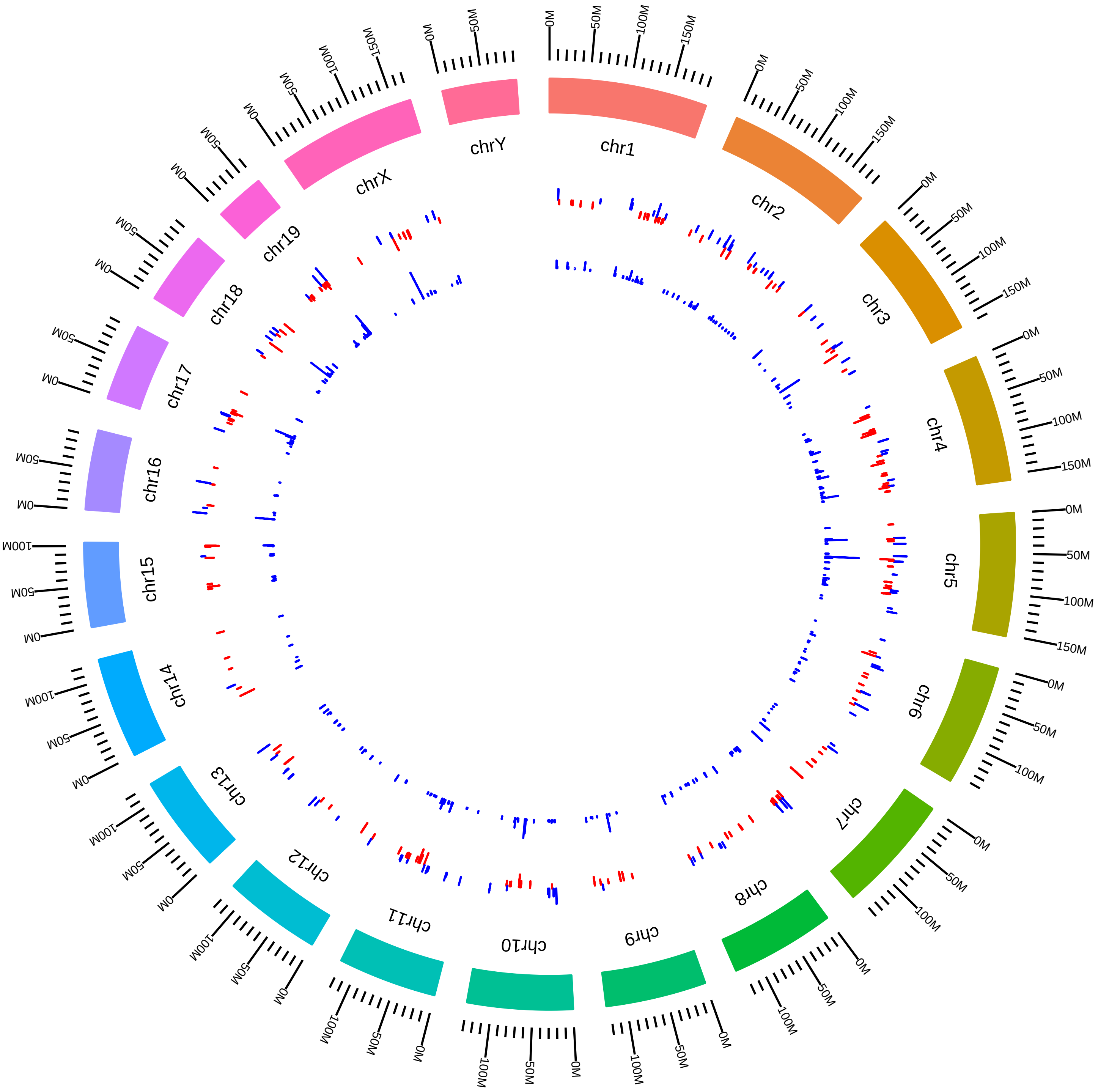

### rsfPtbp1Ptbp2MmDBlackmNPC2015_DEG_DKVsWT.pdf

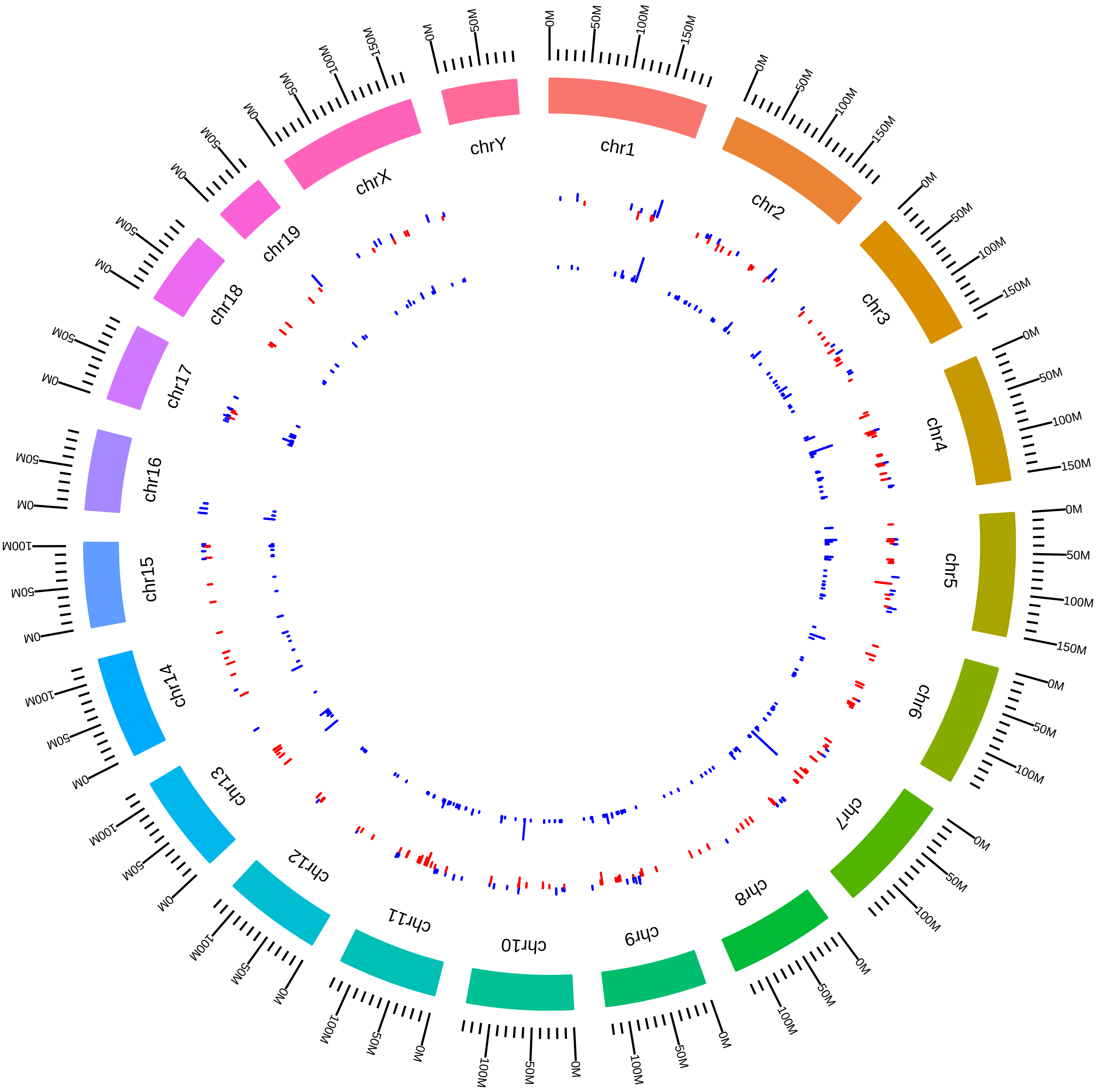

### rsfSrrm4Zfp871MmBBlencowe2017_SVsC.pdf

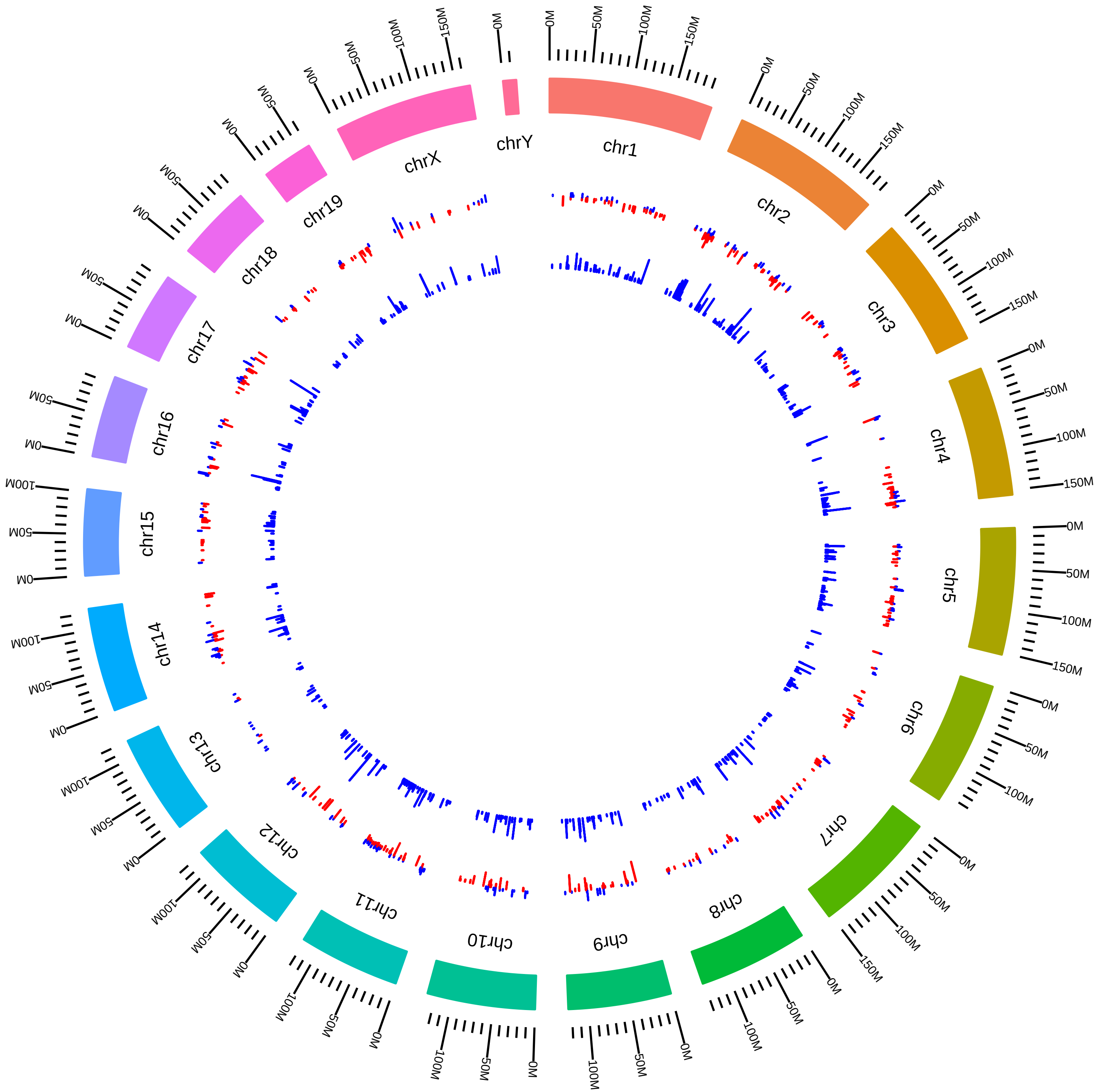

### rsfSrrm4Zfp871MmBBlencowe2017_ZVsC.pdf

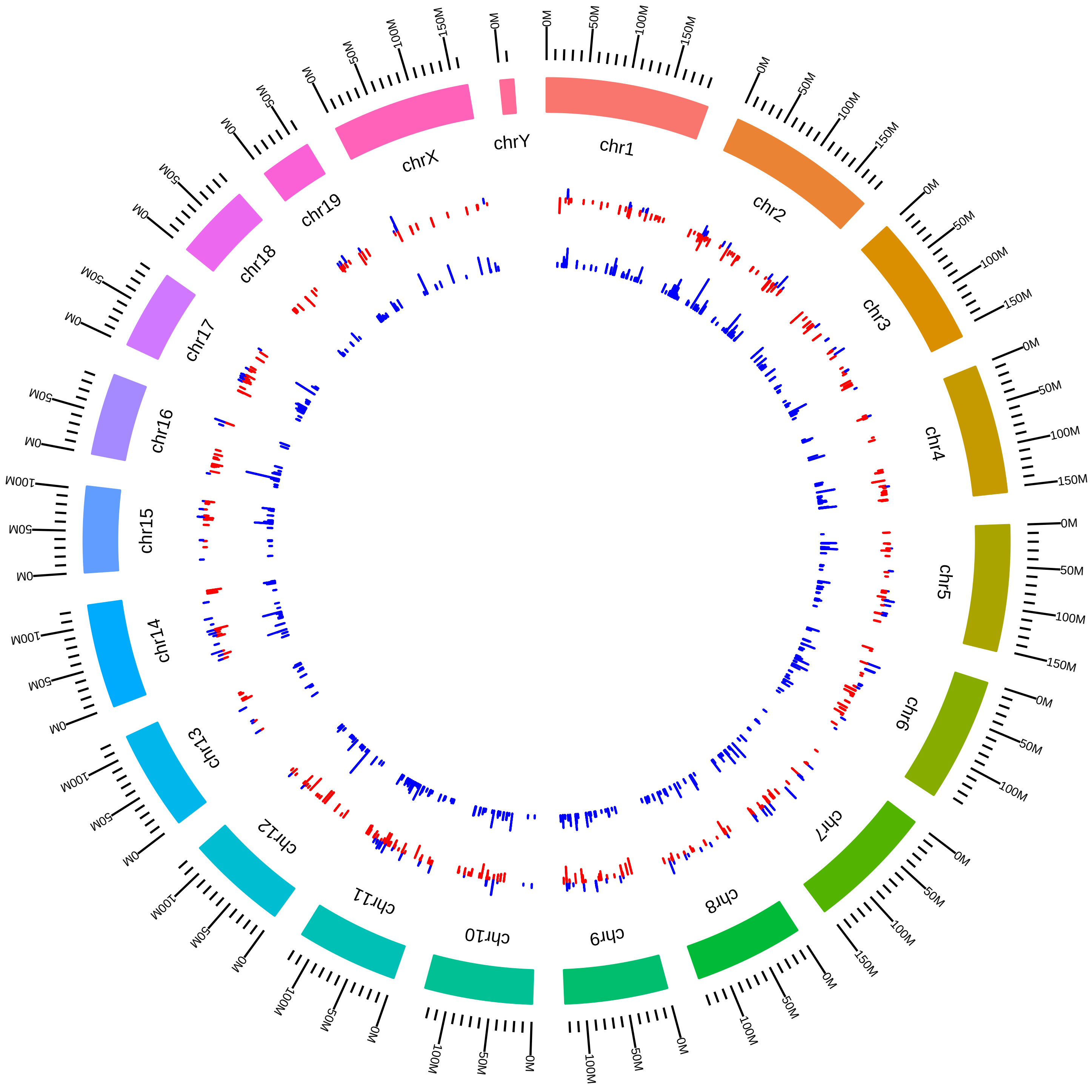

### rsfSrsf1234567MmKNeugebauerCyto_Srsf1KD_vs_WT.pdf

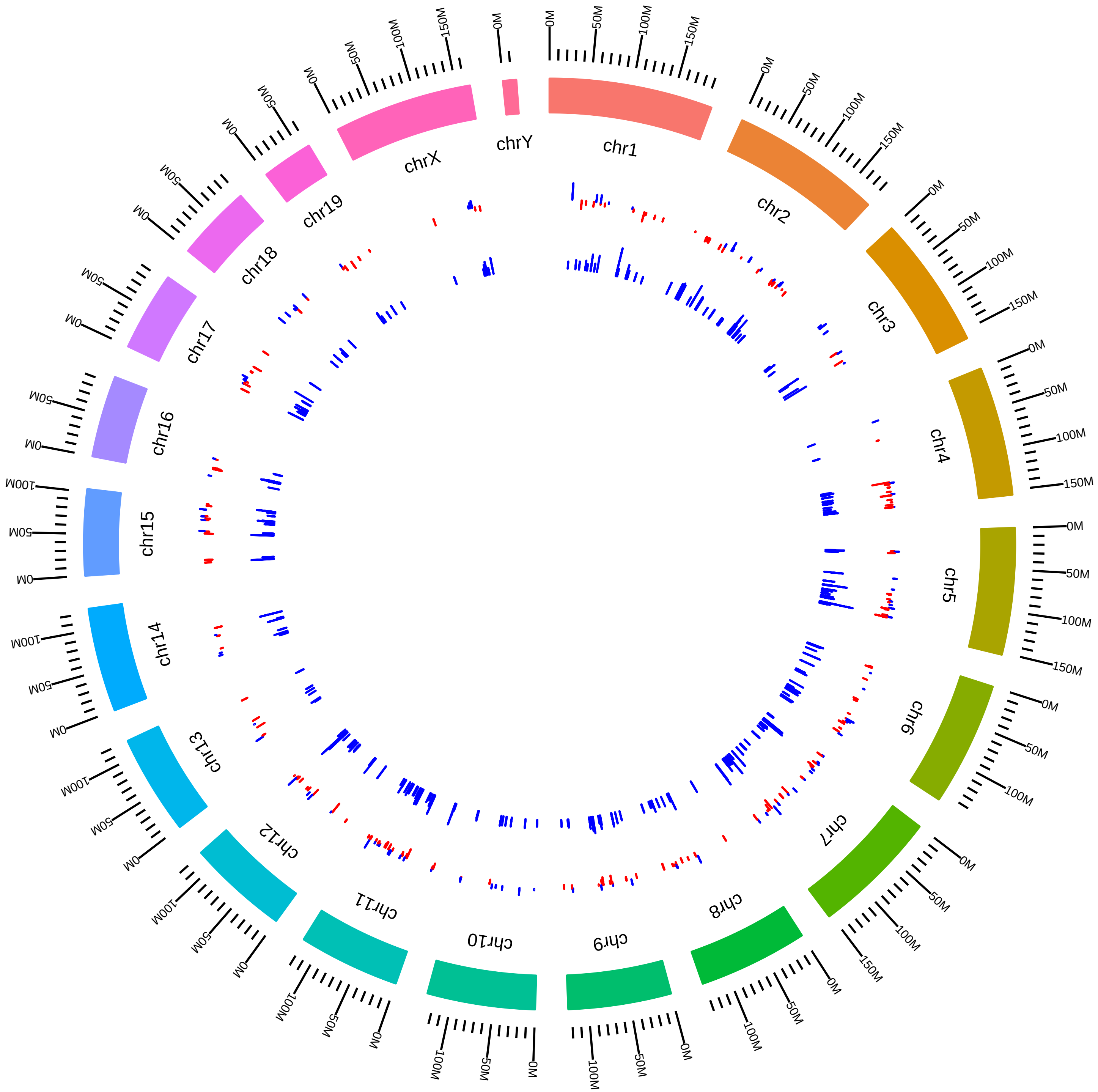

### rsfSrsf1234567MmKNeugebauerCyto_Srsf2KD_vs_WT.pdf

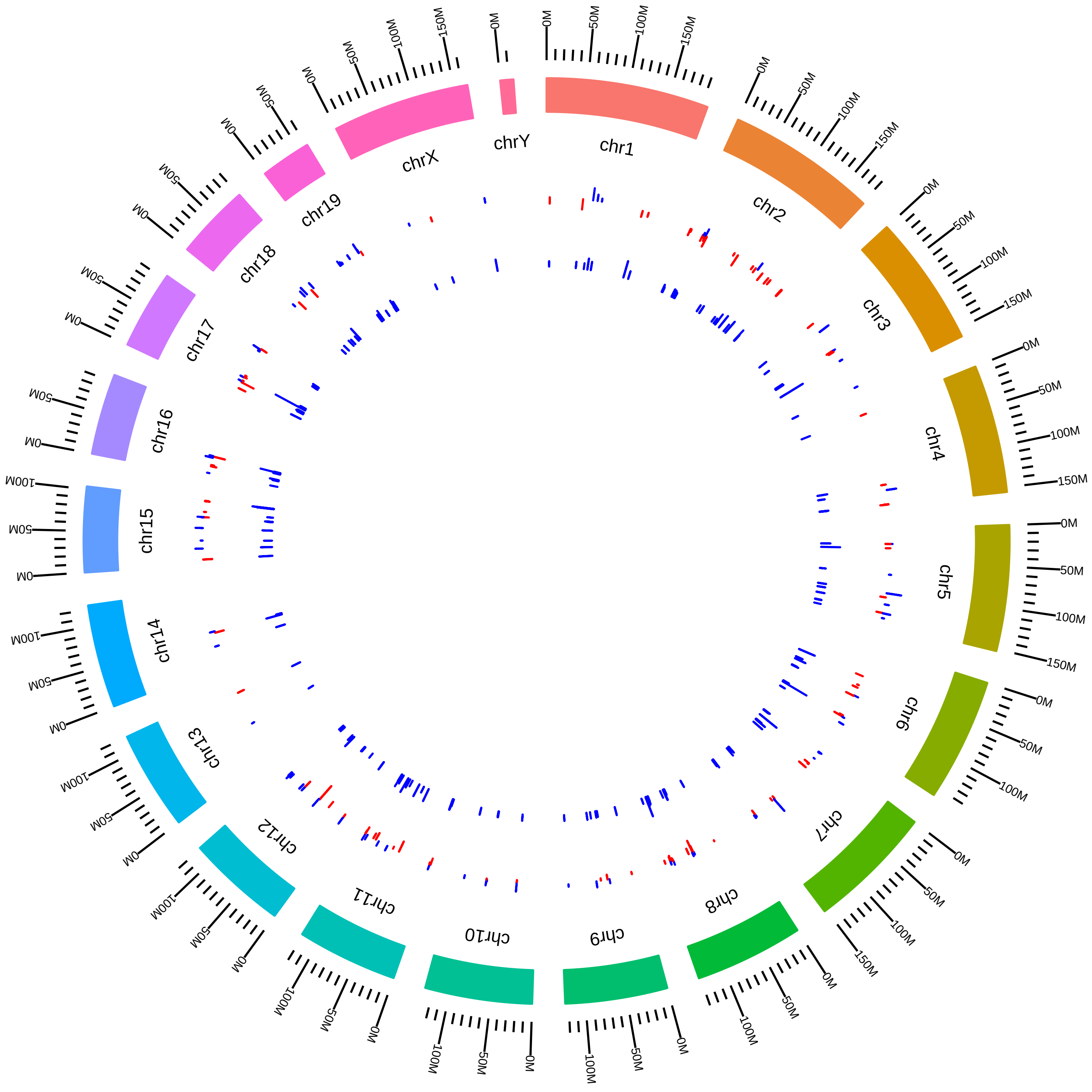

### rsfSrsf1234567MmKNeugebauerCyto_Srsf3KD_vs_WT.pdf

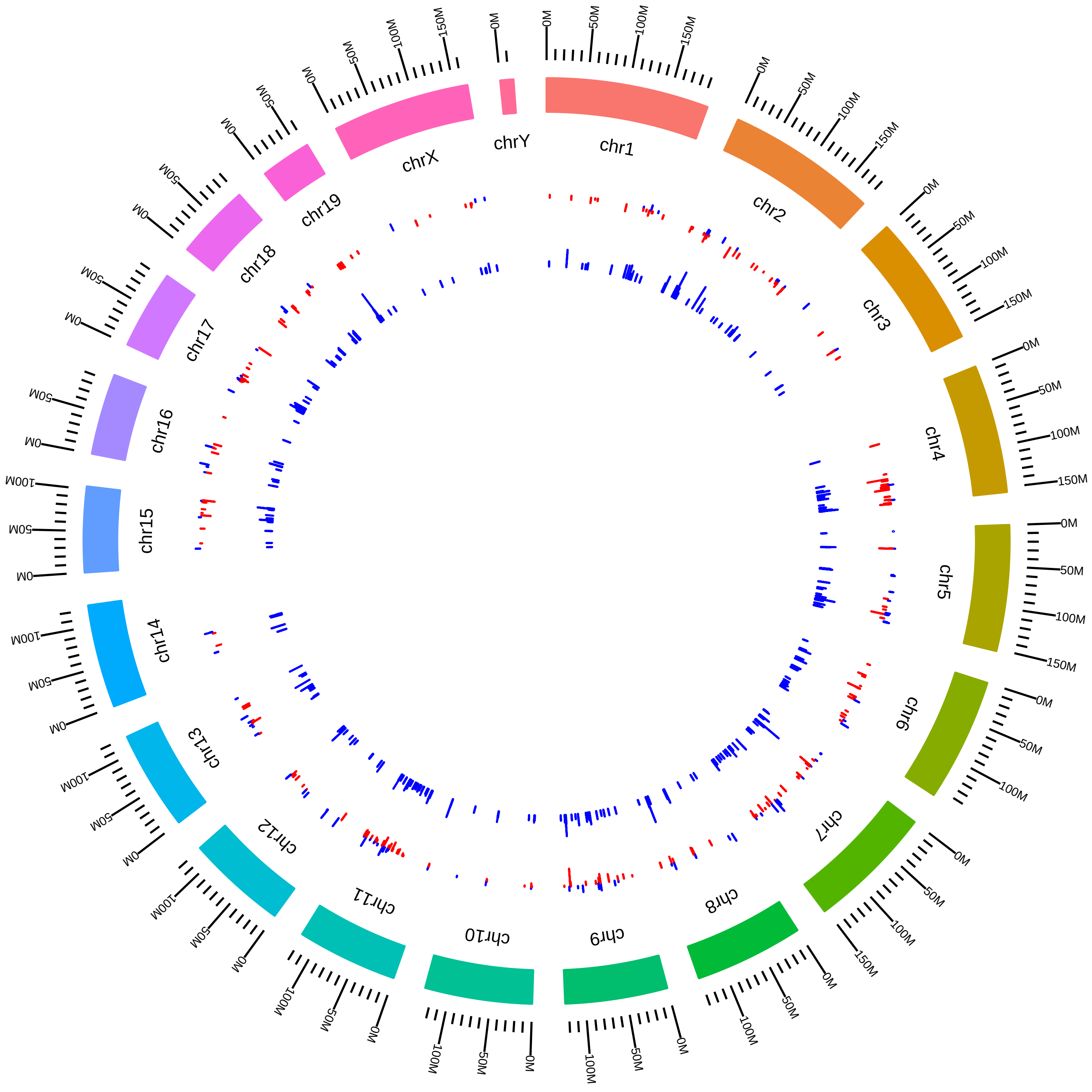

### rsfSrsf1234567MmKNeugebauerCyto_Srsf4KD_vs_WT.pdf

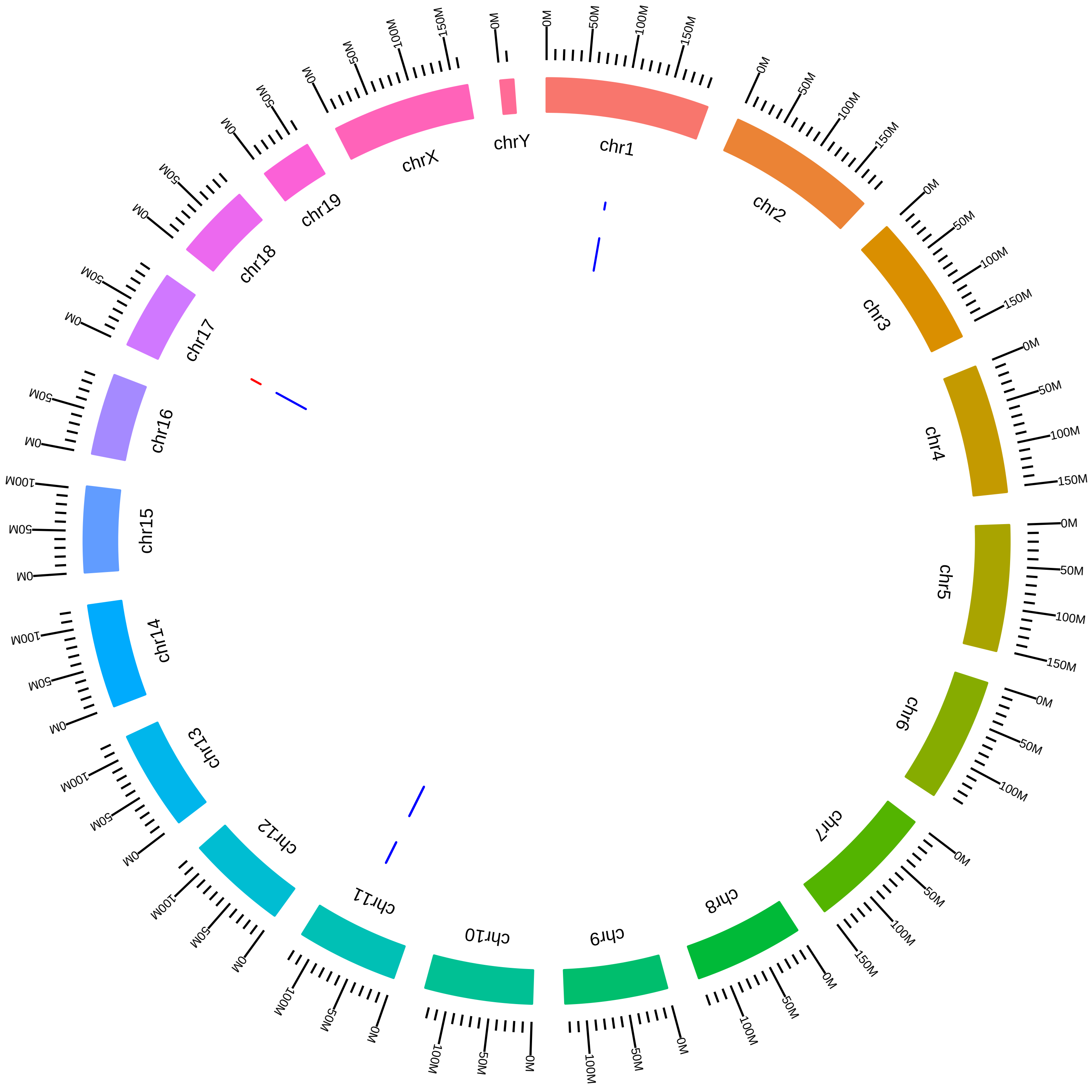

### rsfSrsf1234567MmKNeugebauerCyto_Srsf6KD_vs_WT.pdf

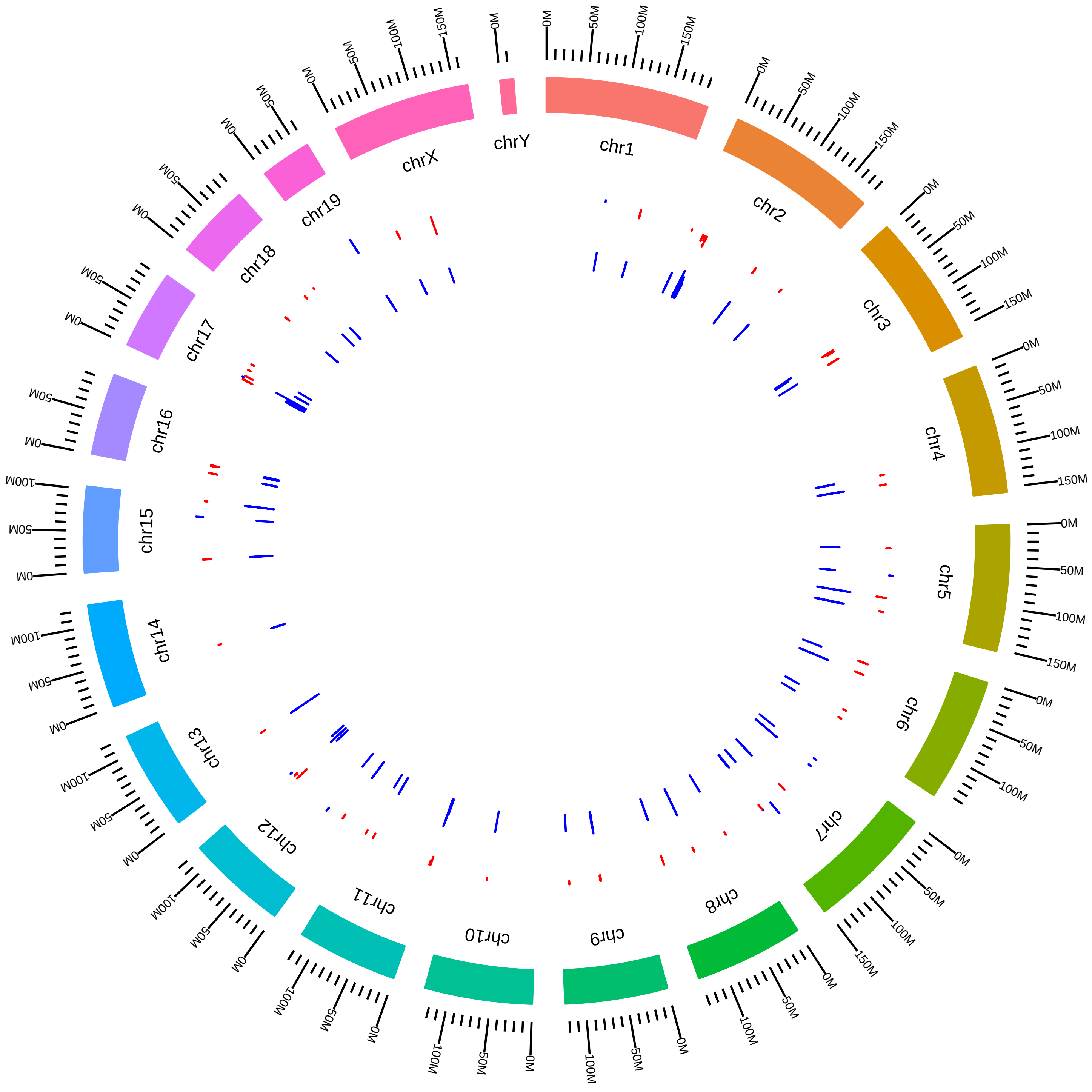

### rsfSrsf1234567MmKNeugebauerCyto_Srsf7KD_vs_WT.pdf

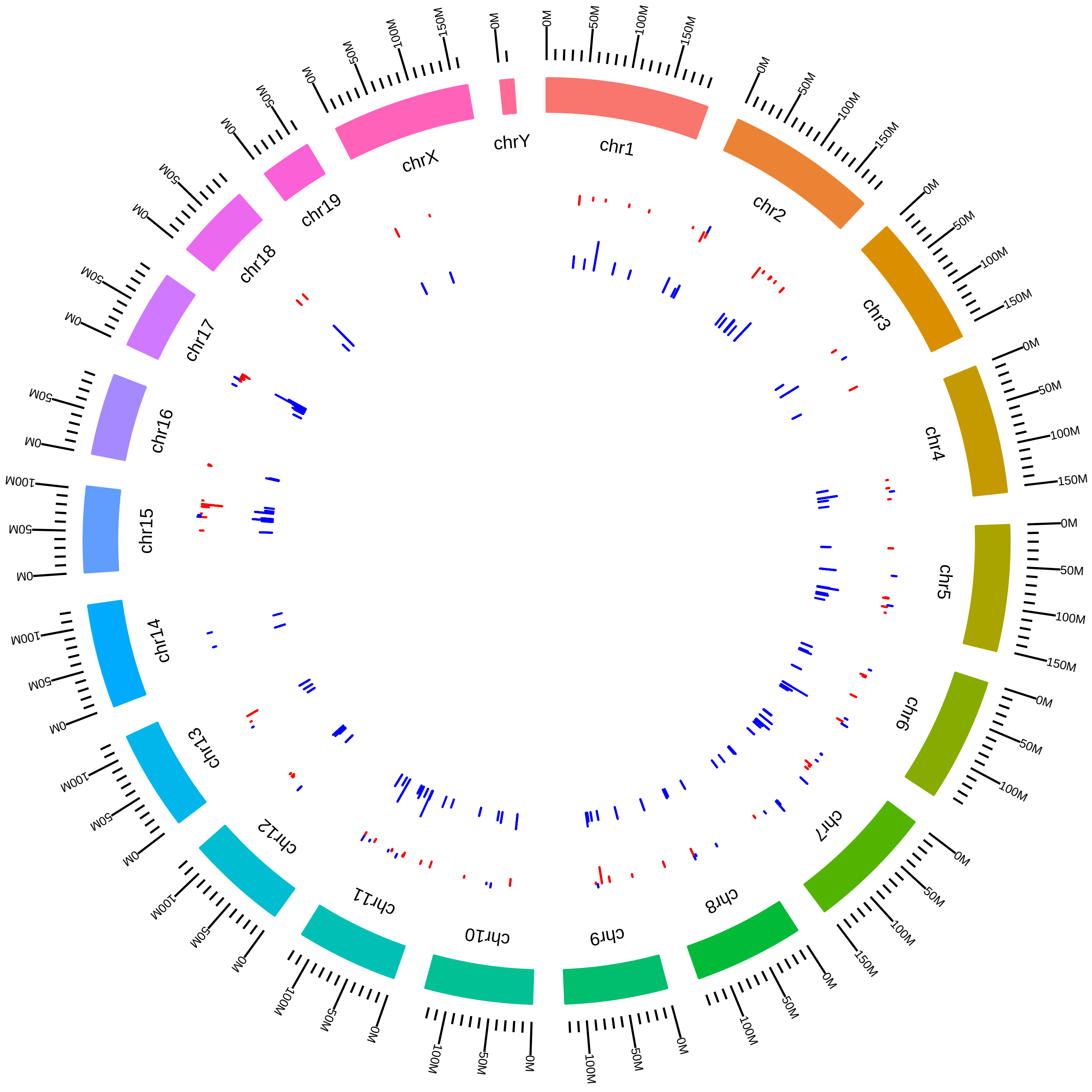

### rsfSrsf1234567MmKNeugebauerWhole_Srsf1KD_vs_WT.pdf

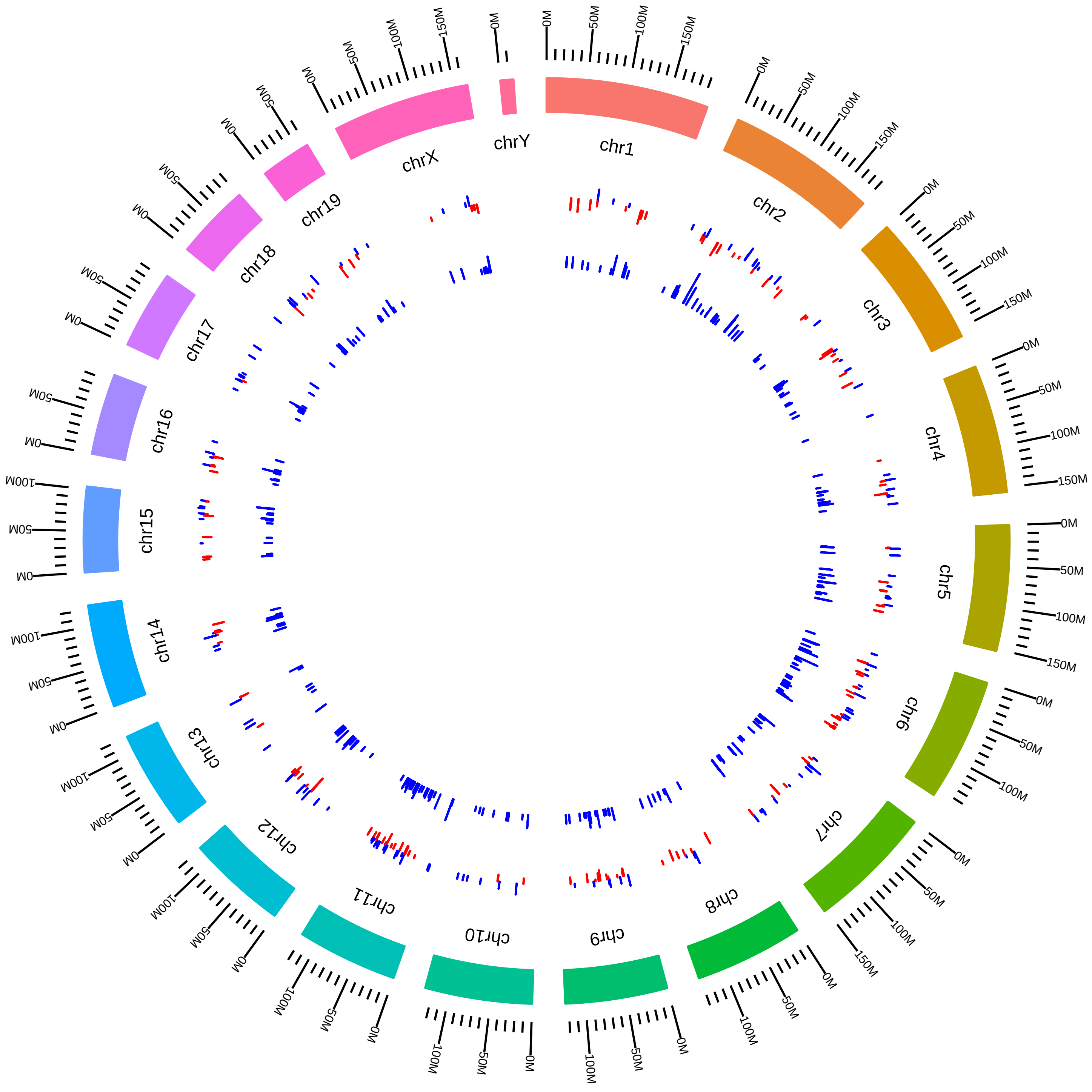

### rsfSrsf1234567MmKNeugebauerWhole_Srsf2KD_vs_WT.pdf

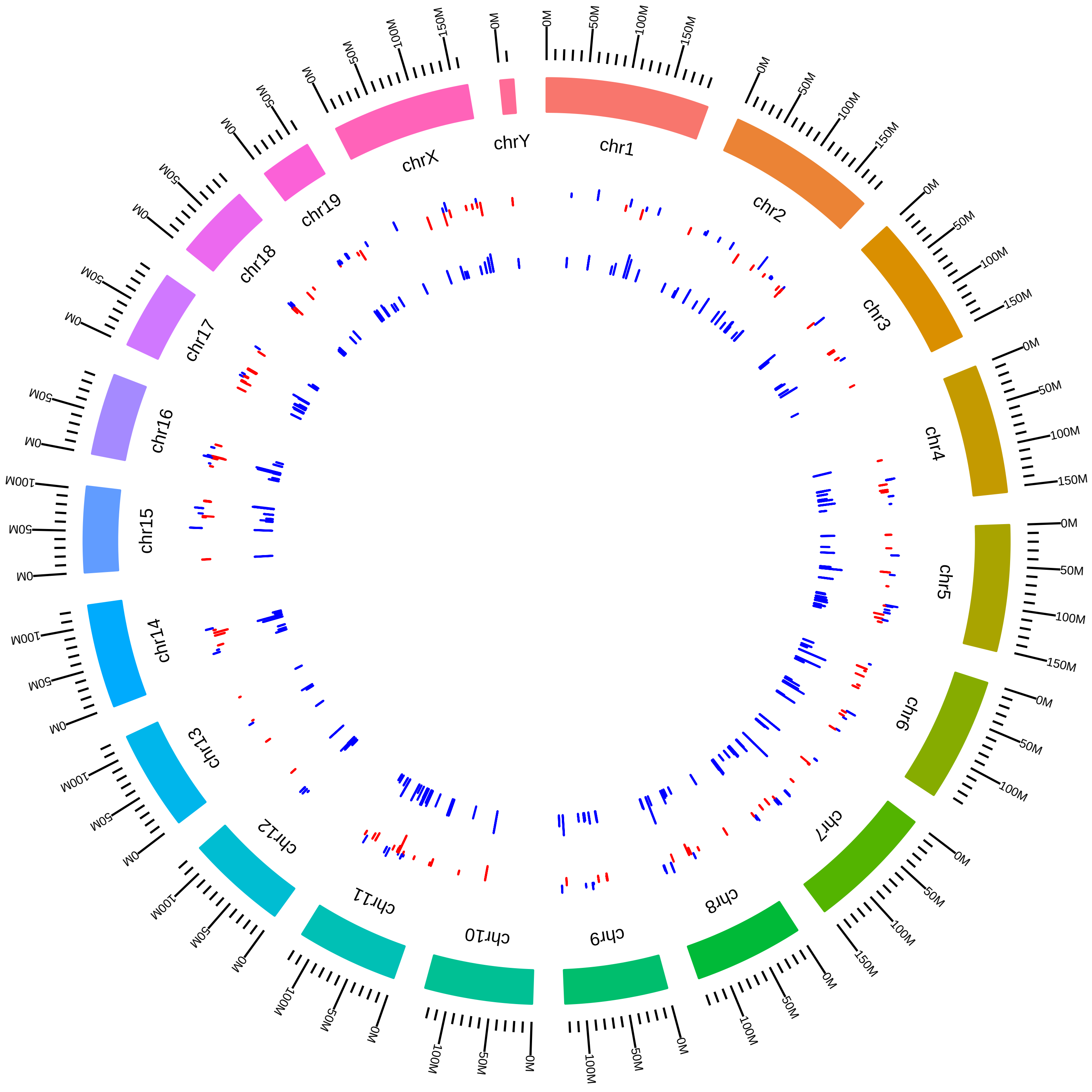

### rsfSrsf1234567MmKNeugebauerWhole_Srsf3KD_vs_WT.pdf

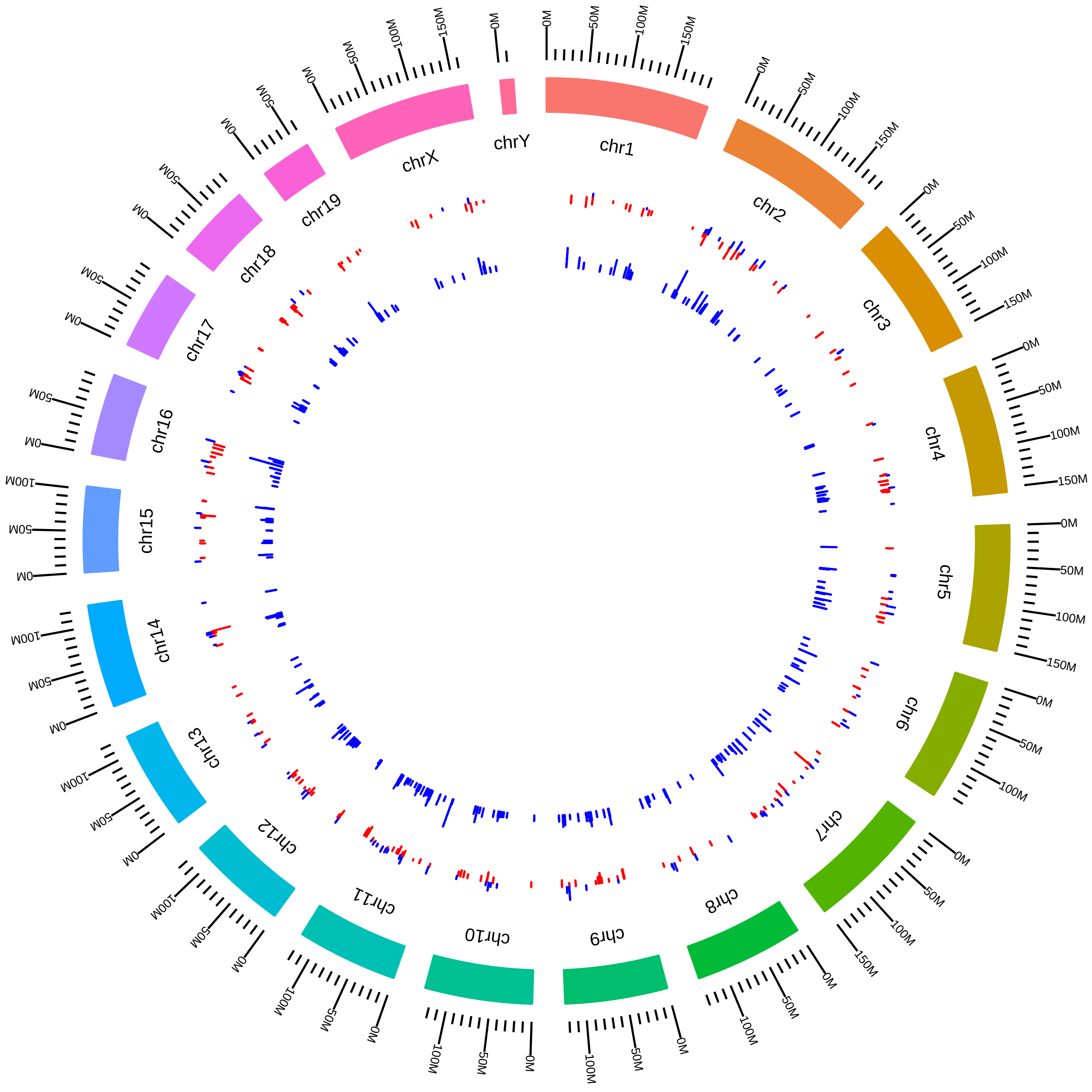

### rsfSrsf1234567MmKNeugebauerWhole_Srsf5KD_vs_WT.pdf

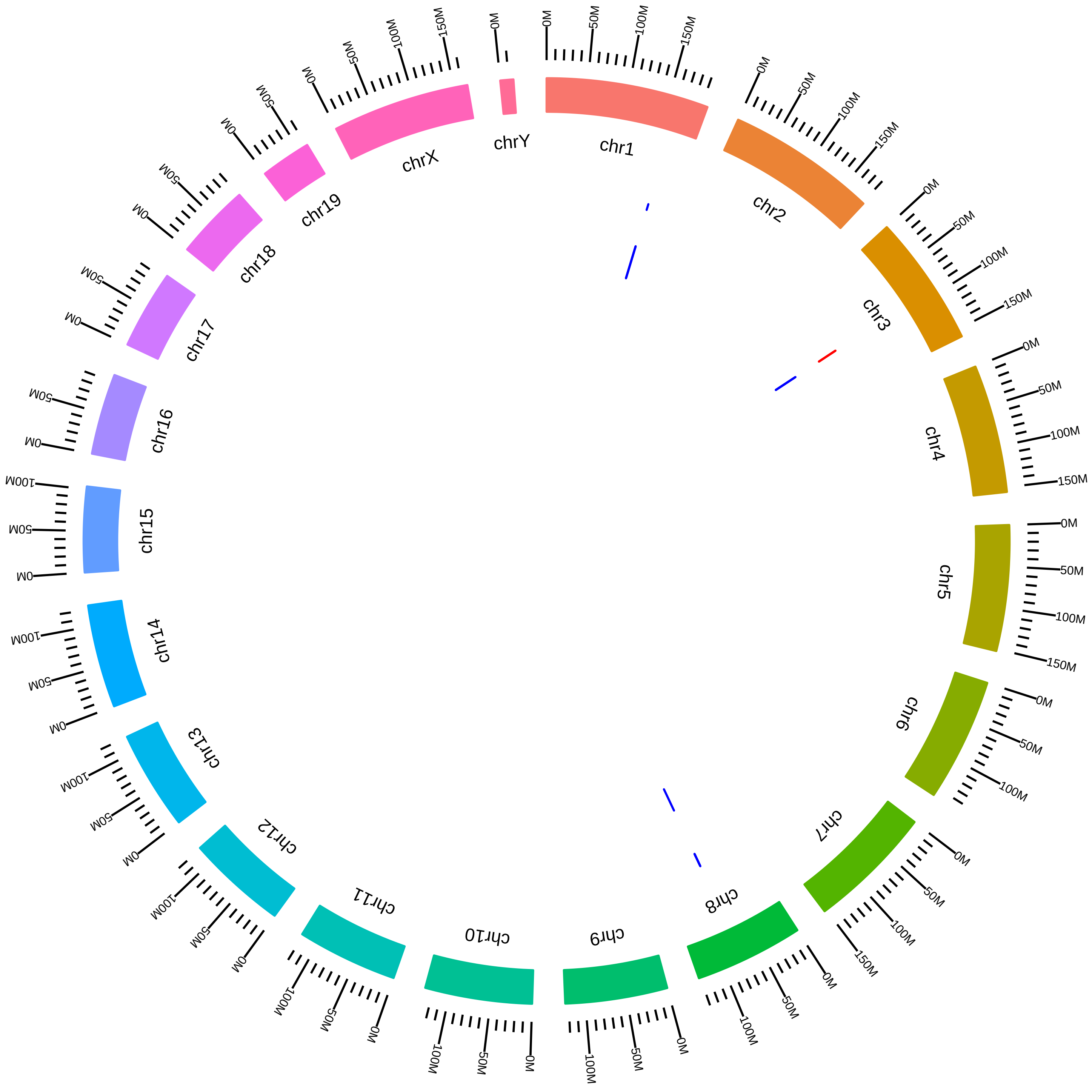

### rsfSrsf1234567MmKNeugebauerWhole_Srsf6KD_vs_WT.pdf

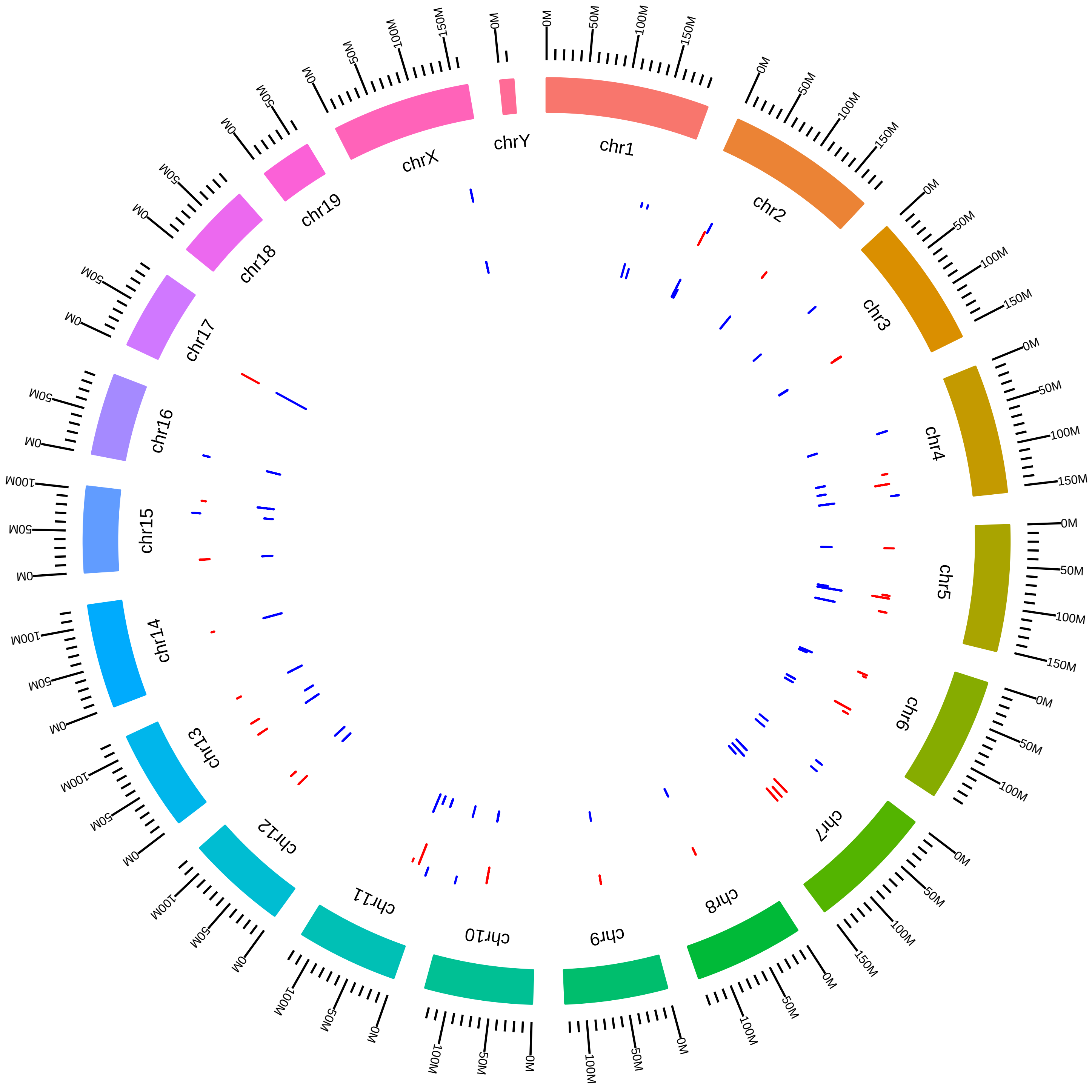

### rsfSrsf1234567MmKNeugebauerWhole_Srsf7KD_vs_WT.pdf

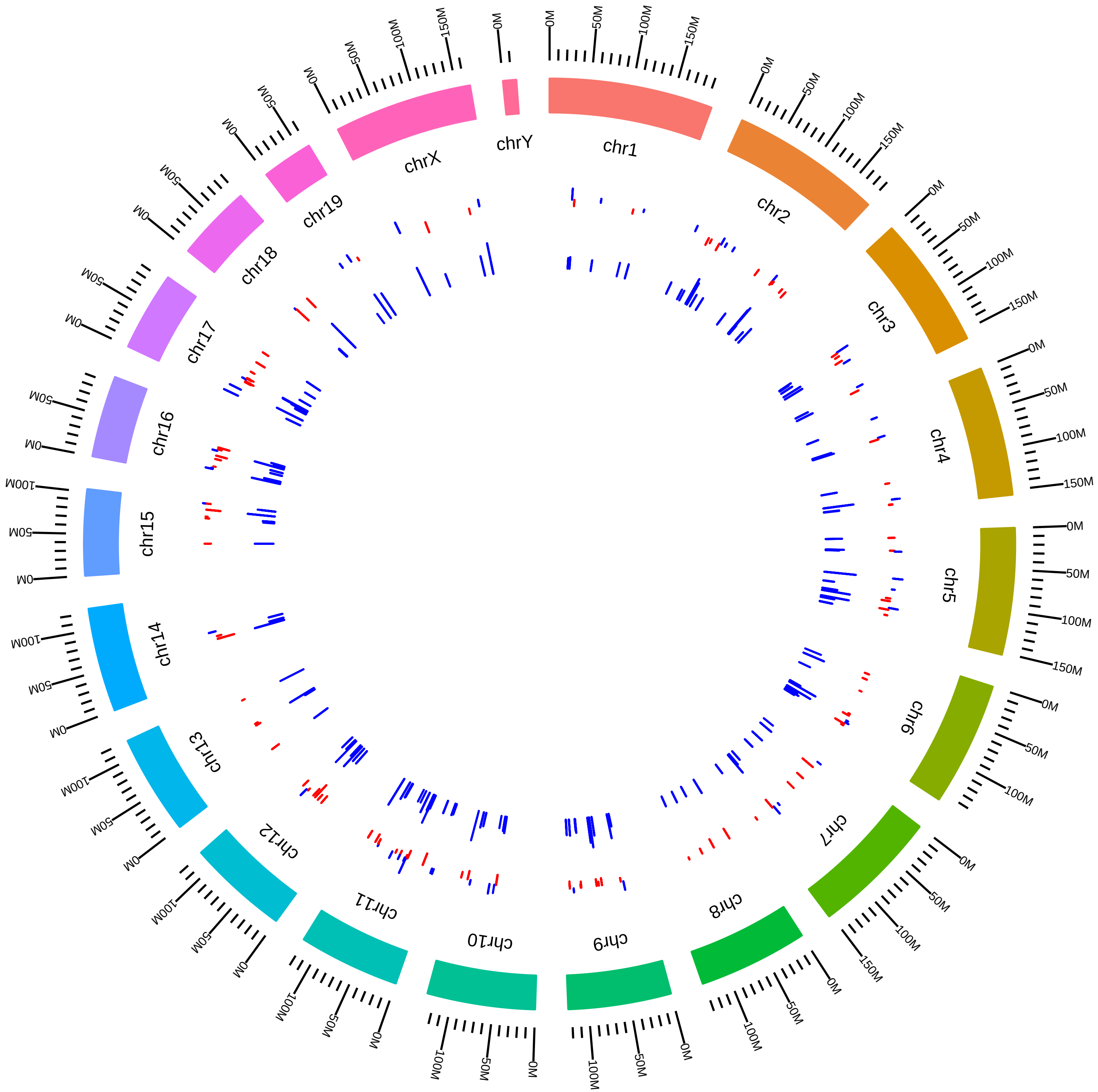

### rsfSyncripMmMKharas2017_DVsC.pdf

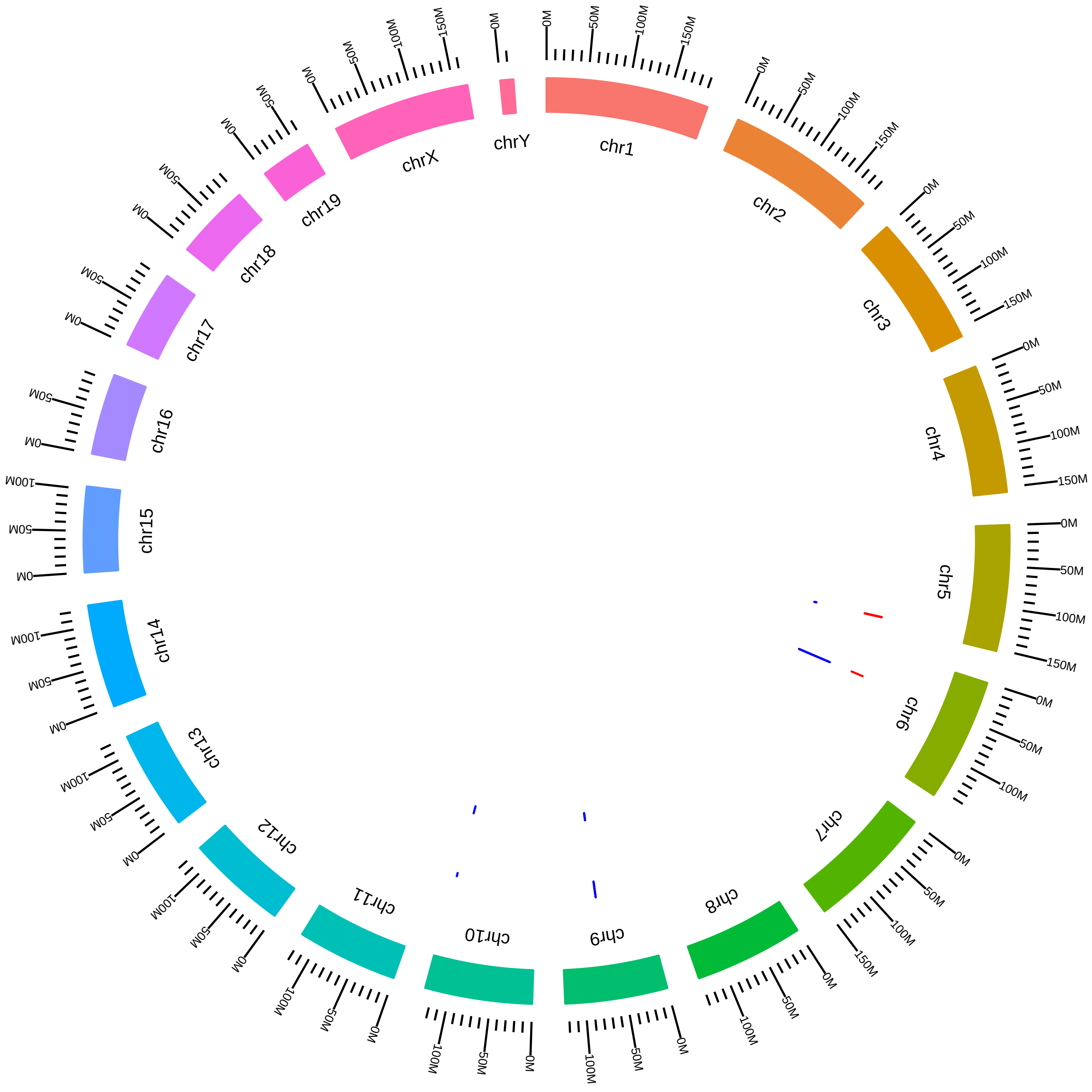

### rsfSyncripMmMKharas2017_KVsC.pdf

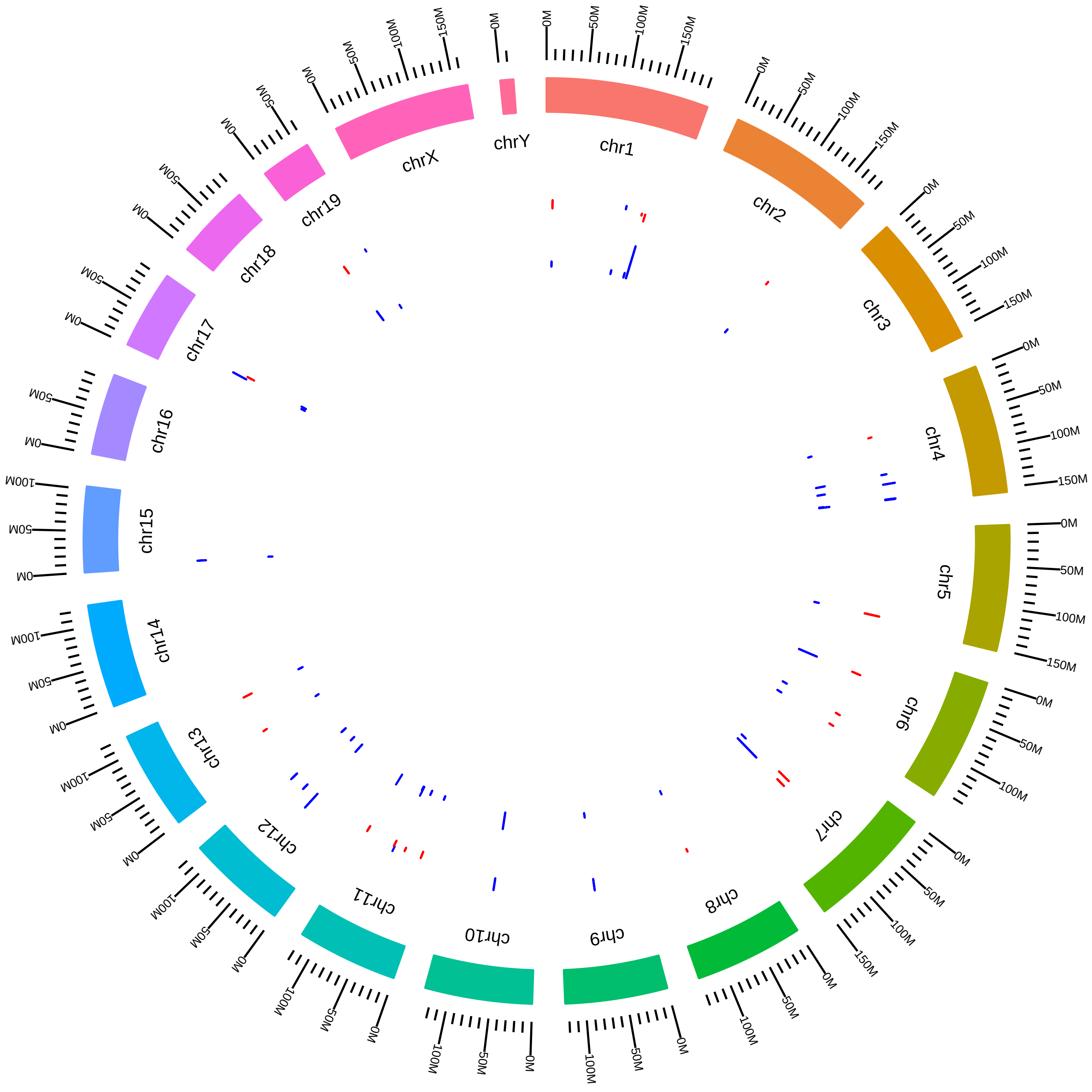

### rsfTardbpMmPCWong_KO_vs_WT.pdf

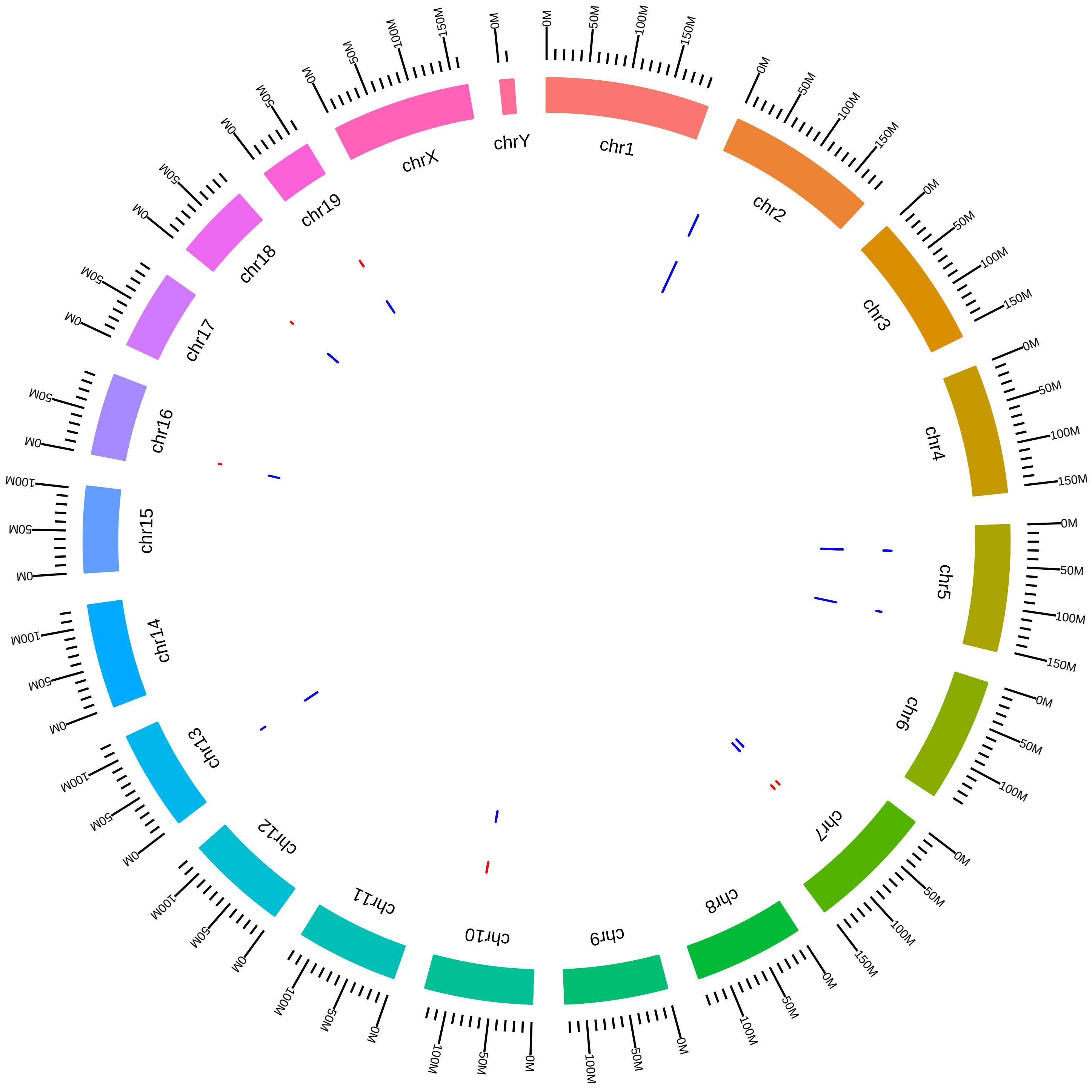

### rsfU2af1MmMWalter2017_SUVsSW.pdf

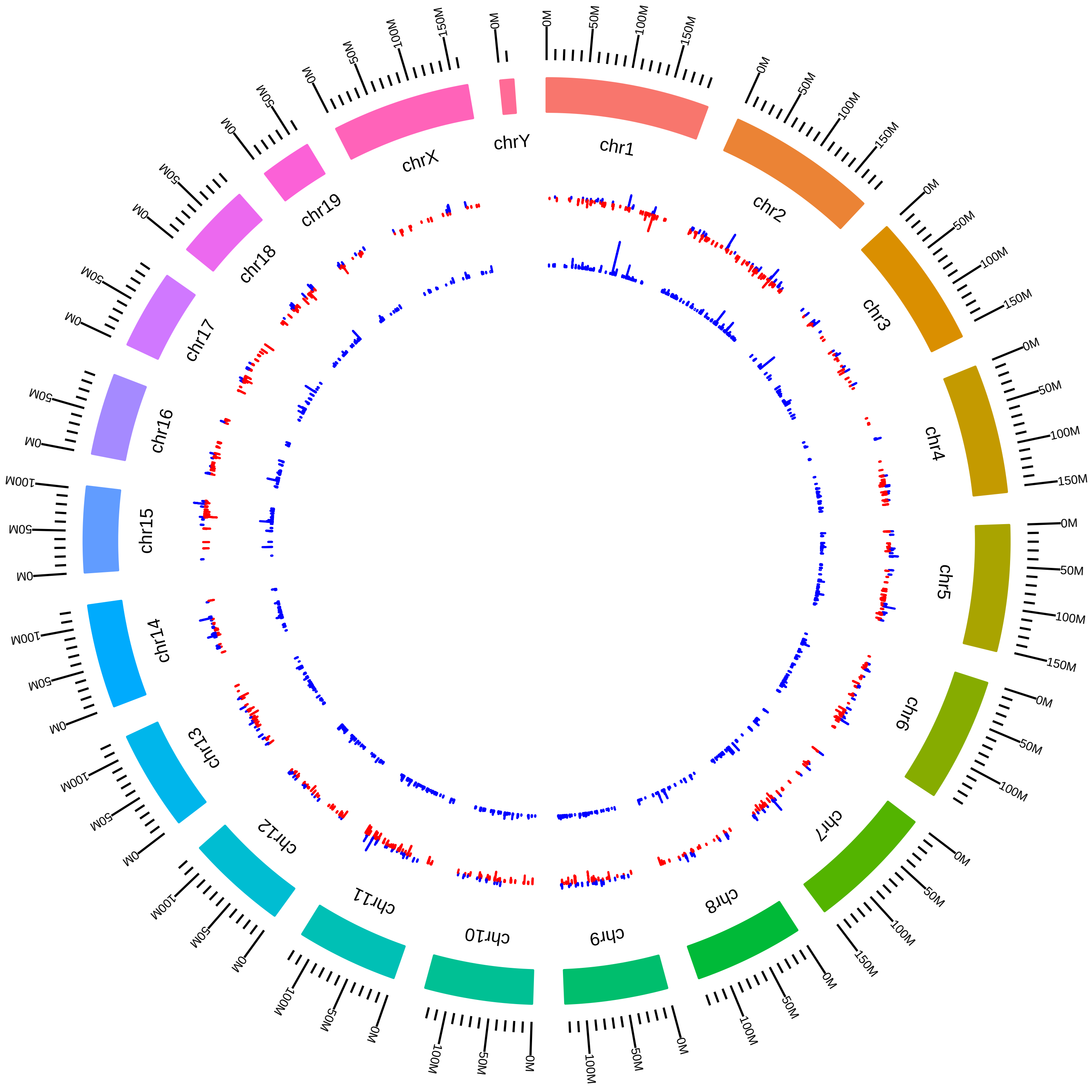

### rsfU2af1MmMWalter2017_SUVsVU.pdf

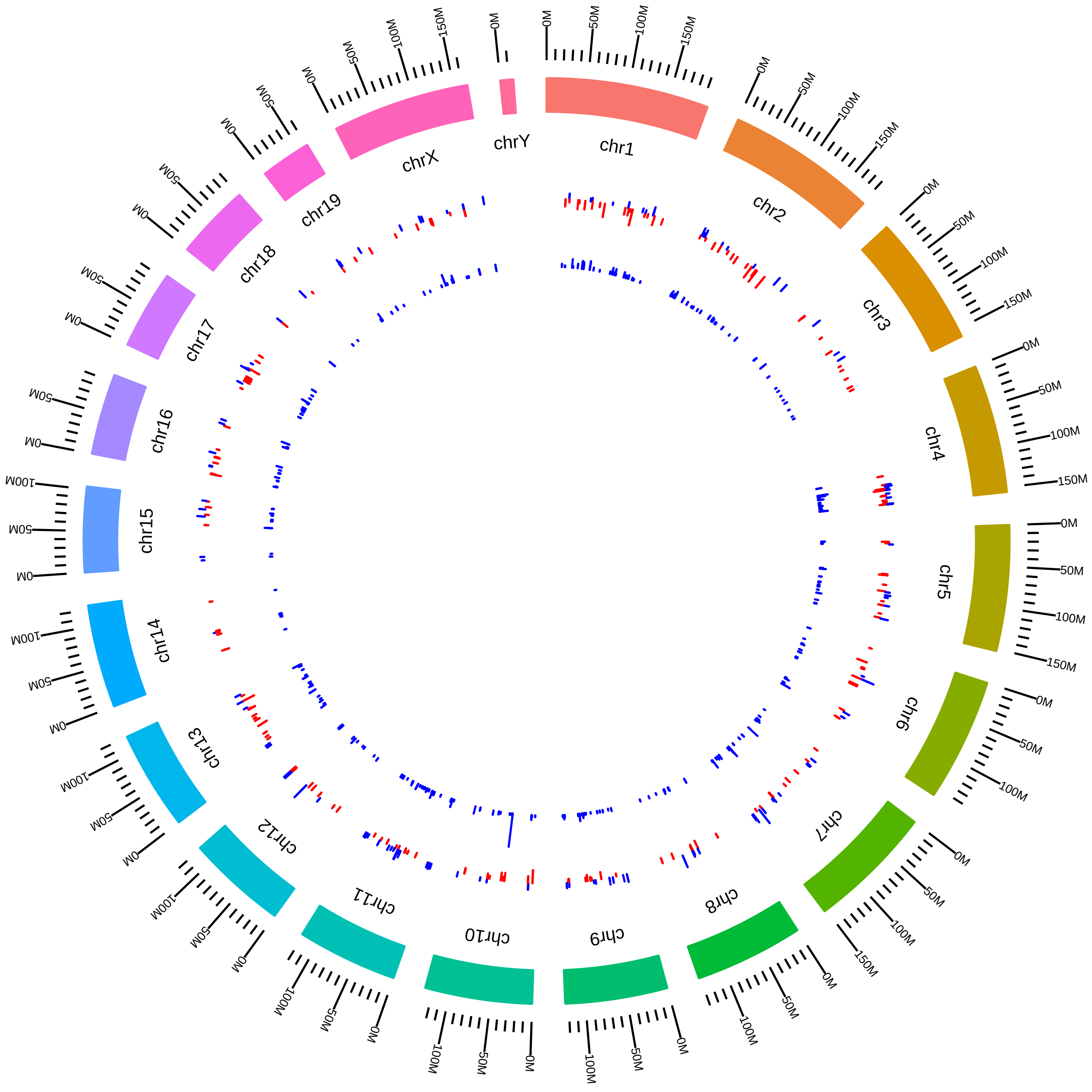

### rsfU2af1MmMWalter2017_SWVsVW.pdf

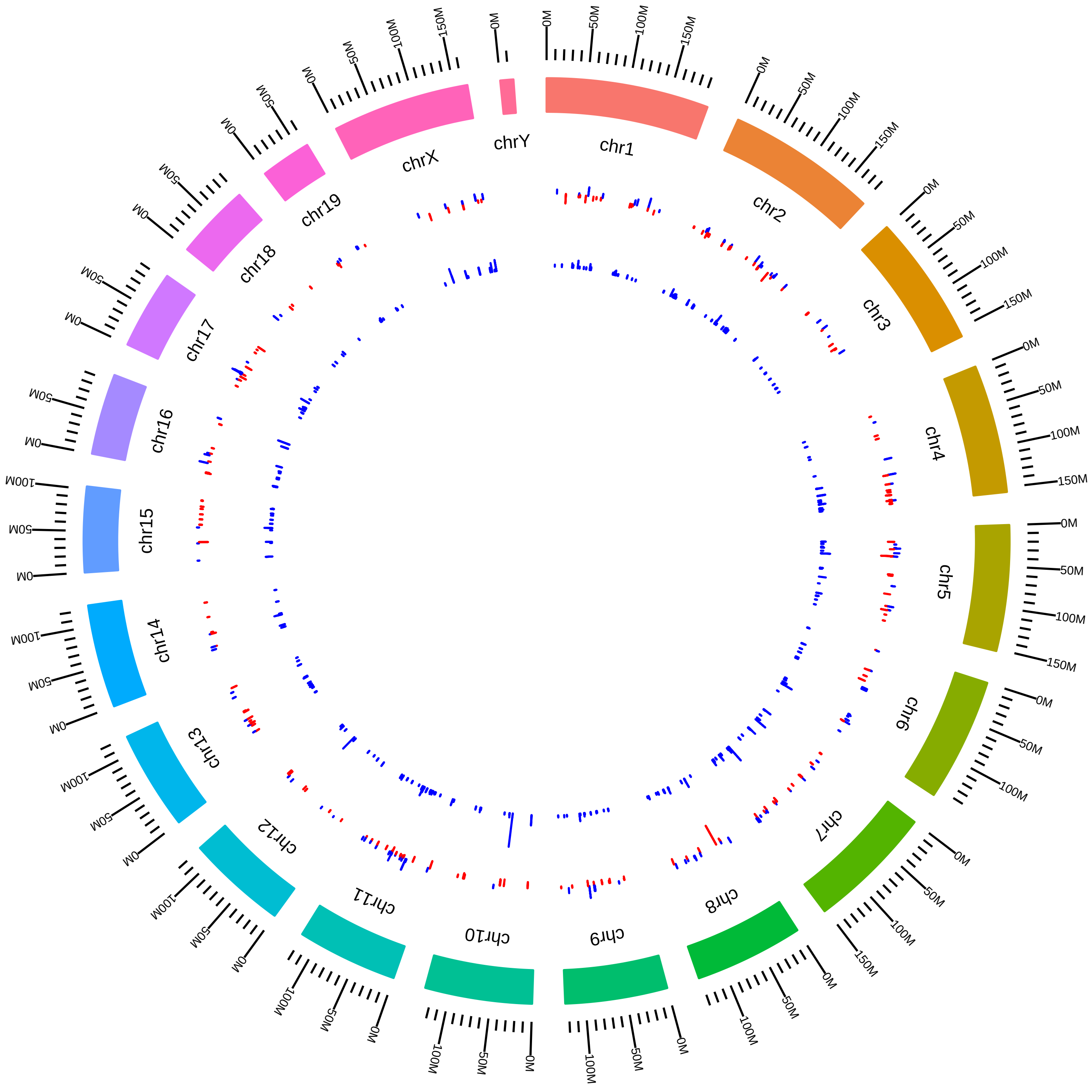

### rsfU2af1MmMWalter2017_VUVsVW.pdf

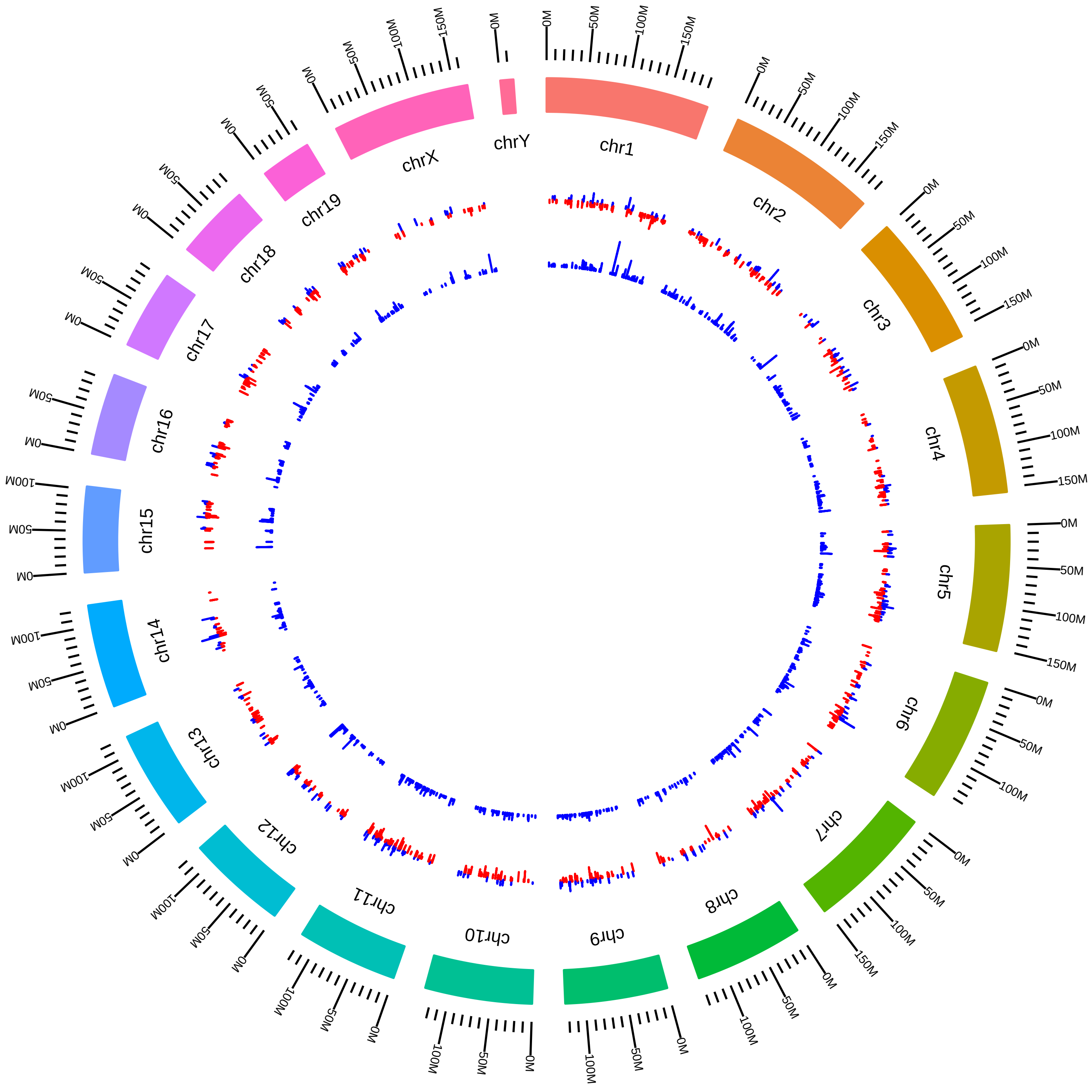

### rsfU2af1MmMWalter_MT_vs_WT.pdf

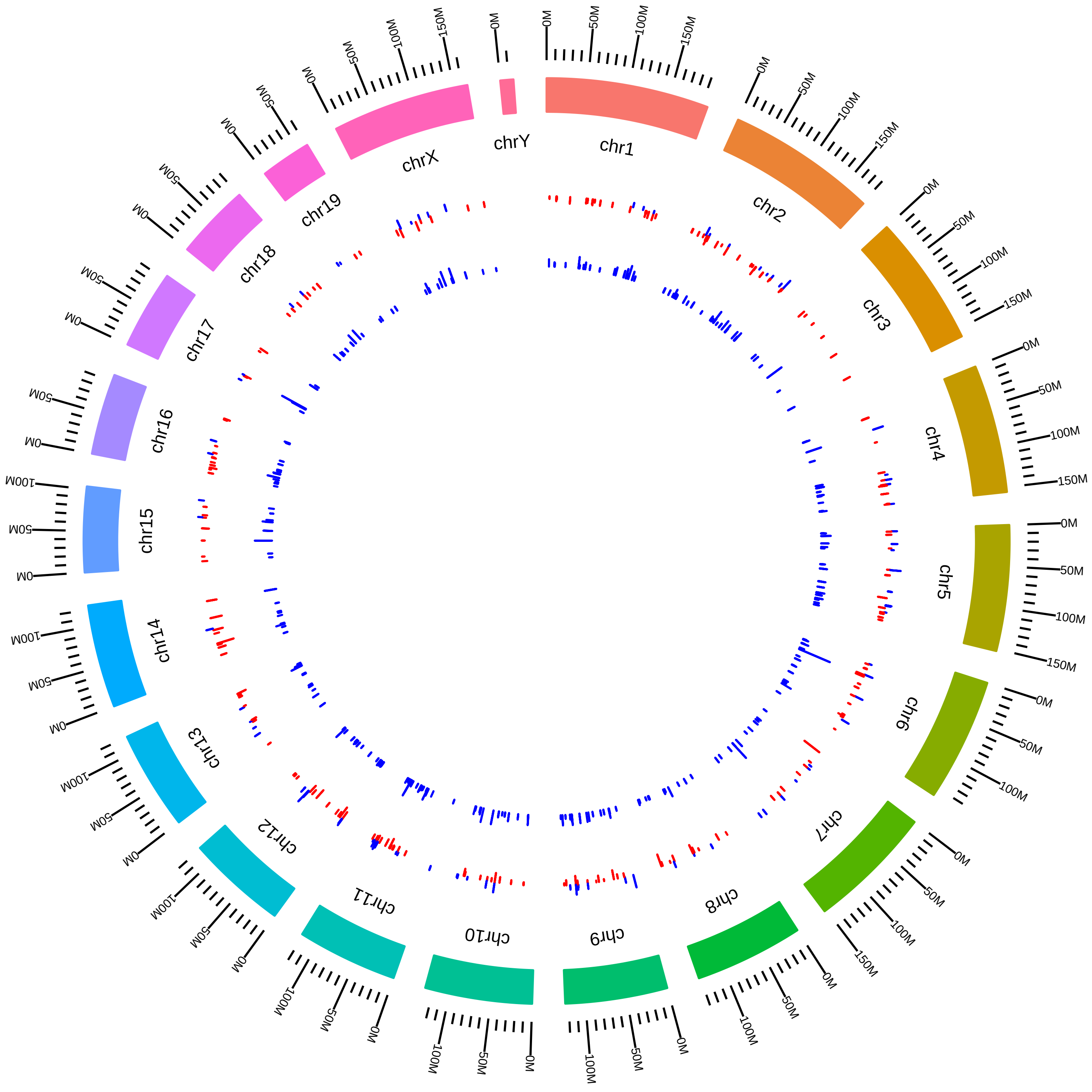
