## Supplementary figures and images for "Integrated analysis of a compendium of RNA-Seq datasets for splicing factors"

### rbpCirbpMmJunYan_C_vs_H.pdf

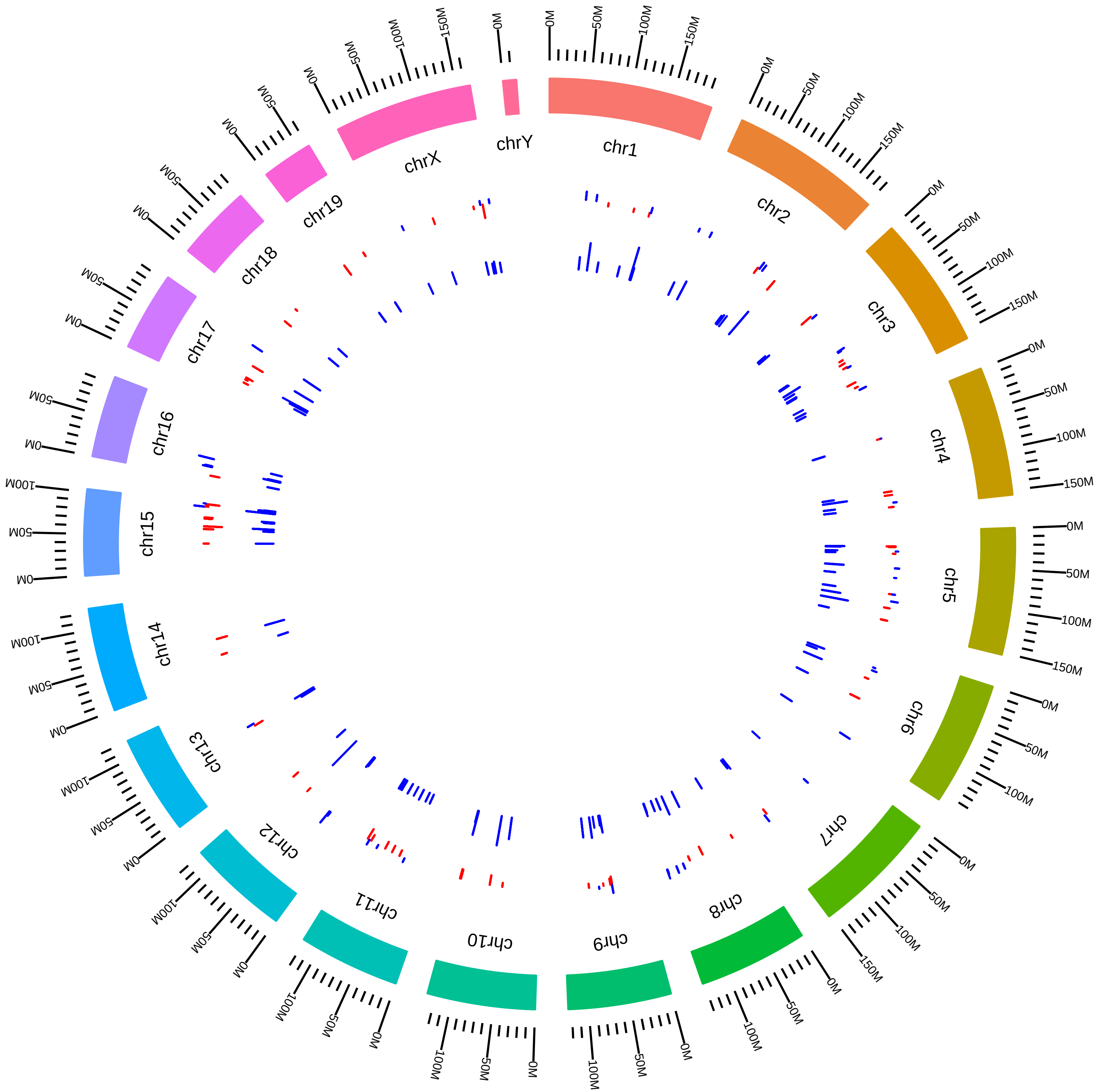

### rbpCirbpMmJunYan_R_vs_H.pdf

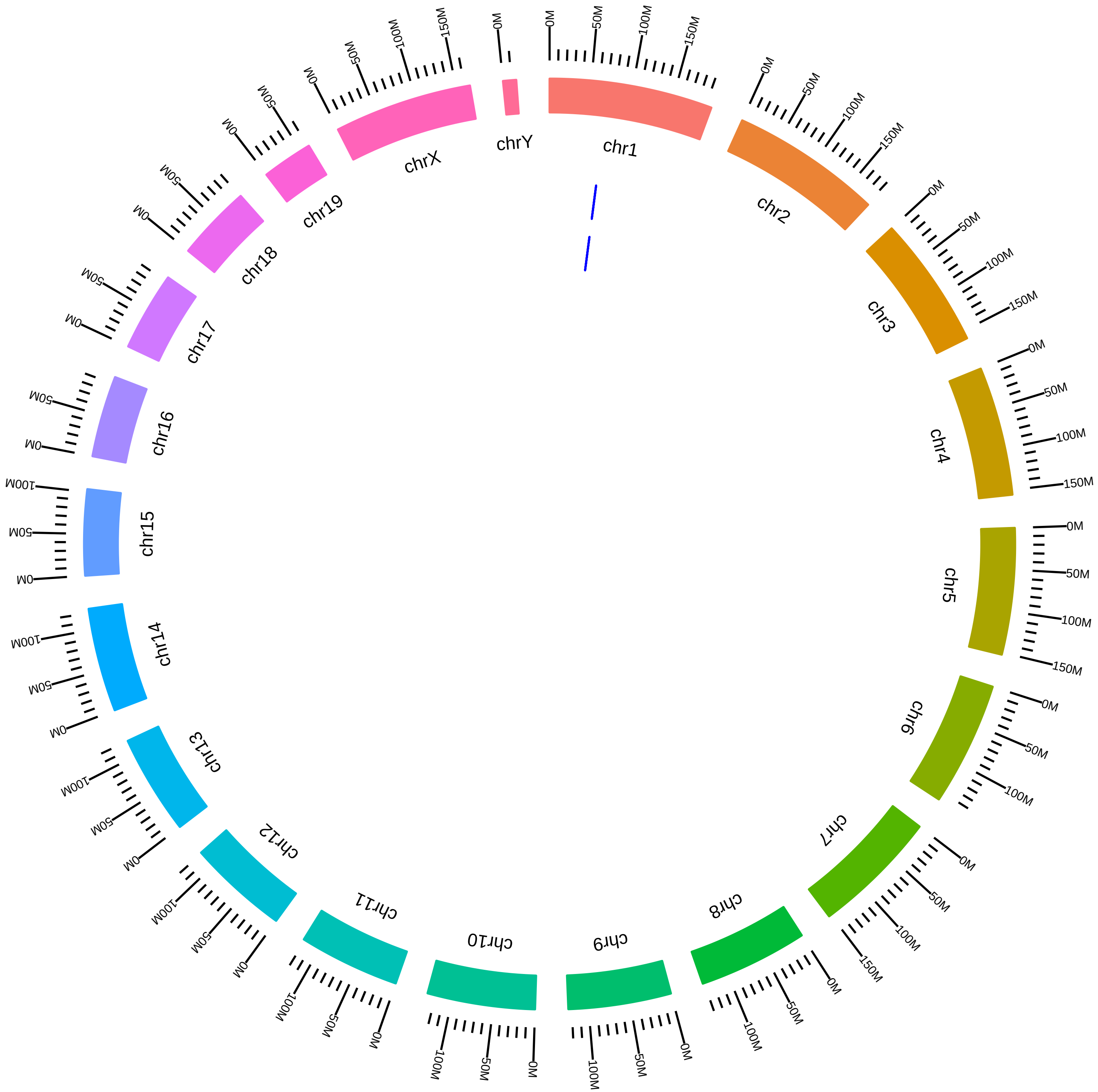

### rsfCd2bp2MmCFreund2015_KVsW.pdf

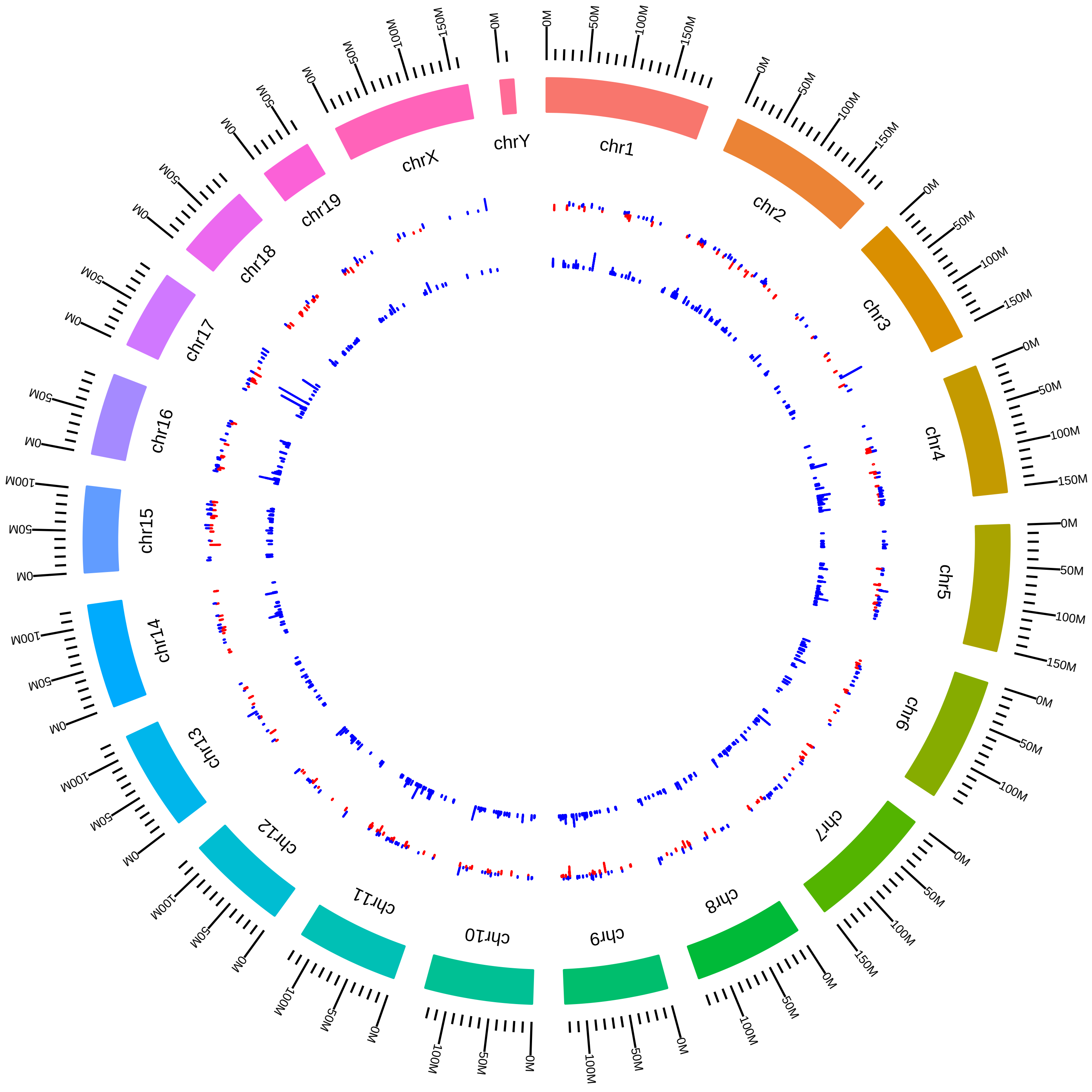

### rsfCelf1MmCBurgeHeart_L_vs_S.pdf

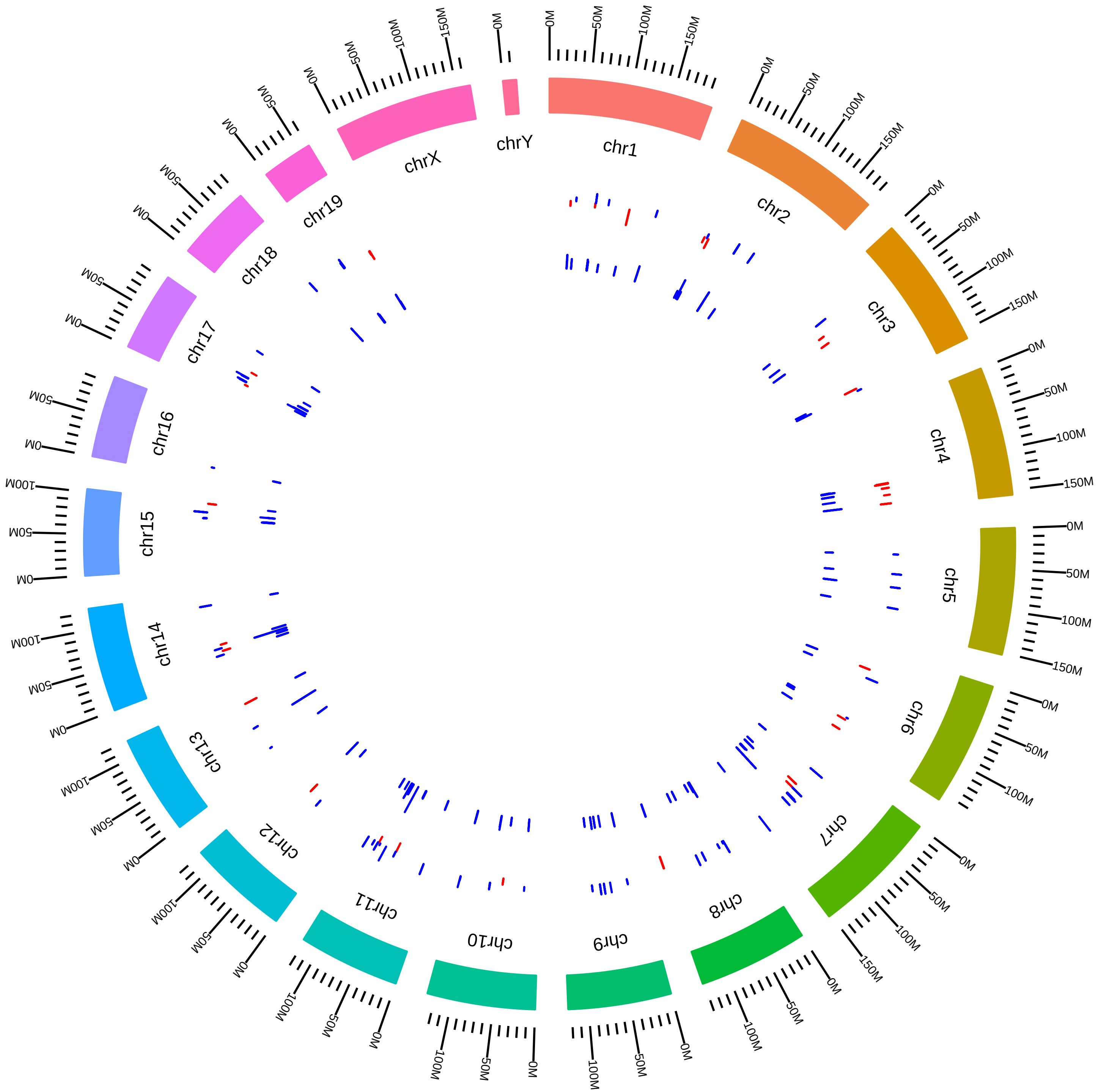

### rsfCelf1MmCBurgeMuscle_L_vs_S.pdf

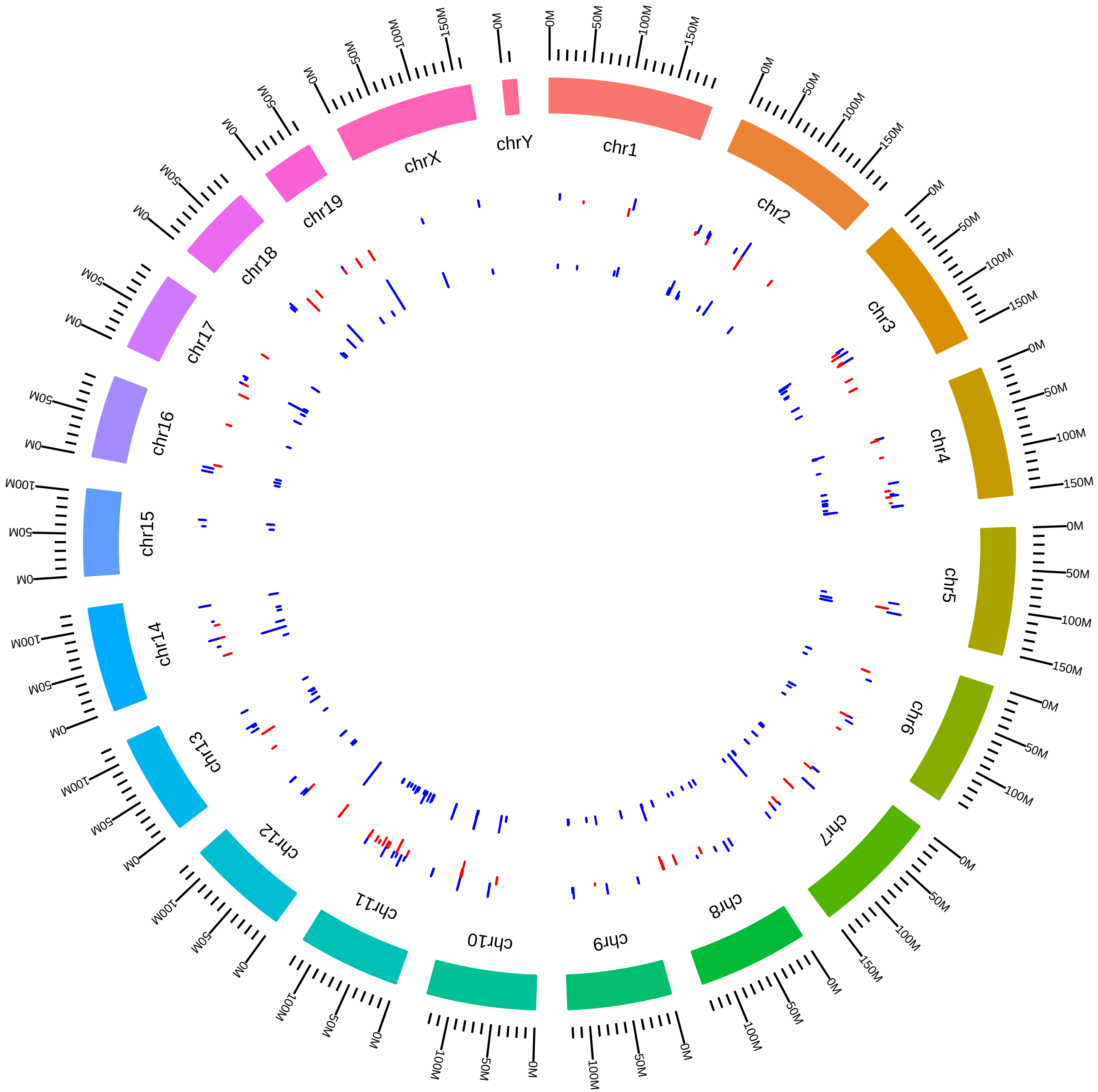

### rsfCelf1MmTCooper_KO_vs_WT.pdf

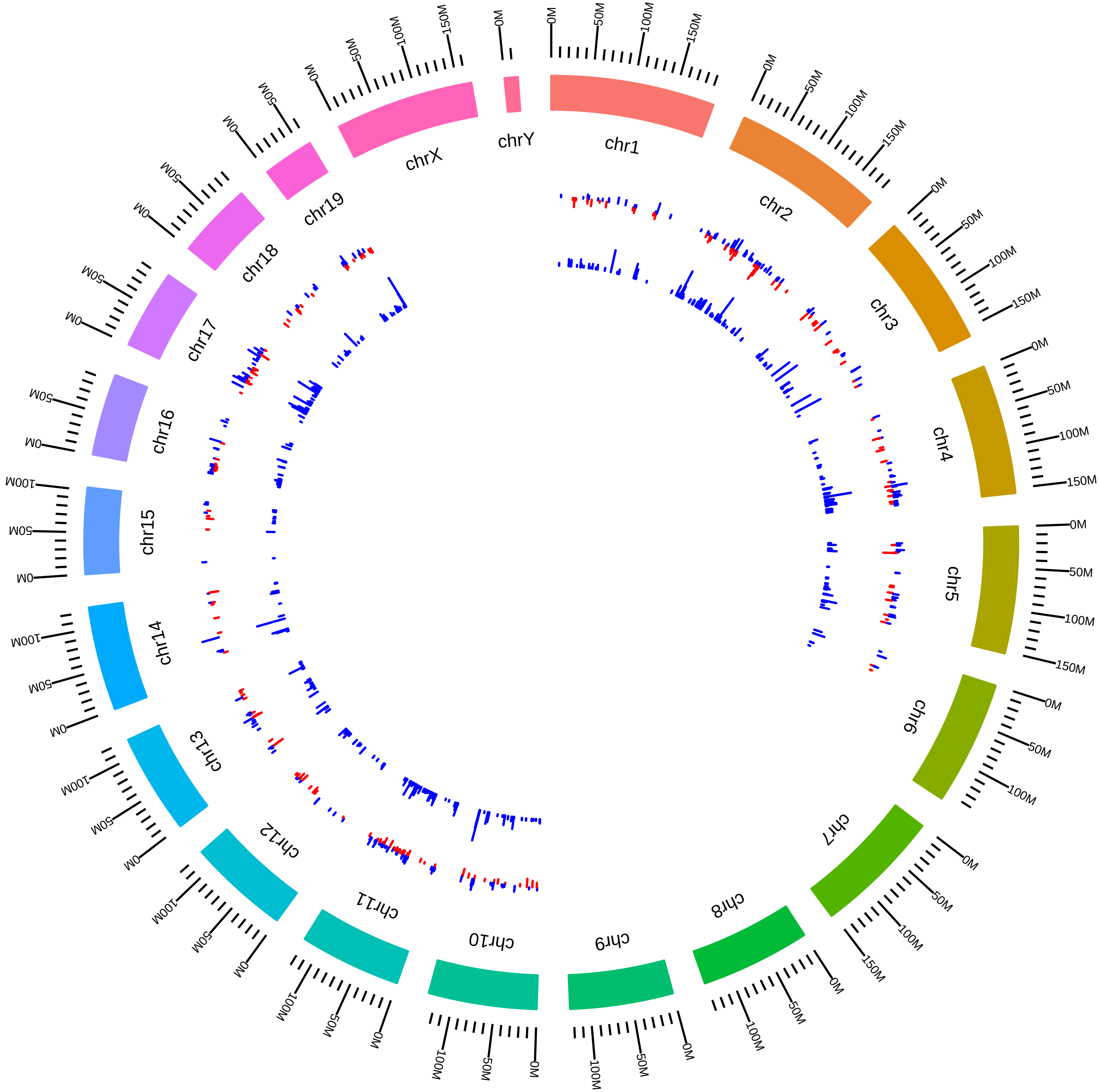

### rsfCelf2MmCBurge_L_vs_S.pdf

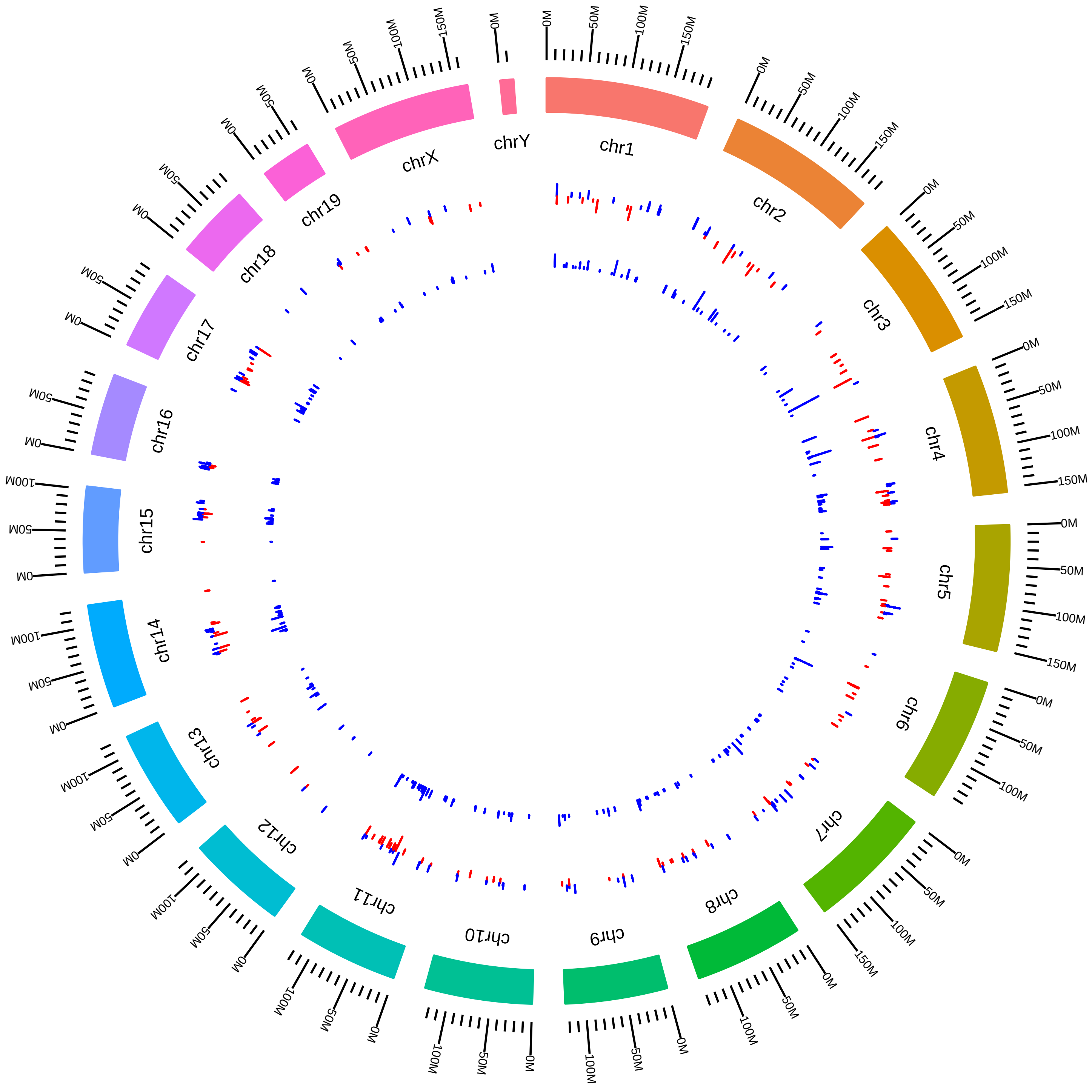

### rsfCirbpRbm3MmJunYan2013_CVsH.pdf

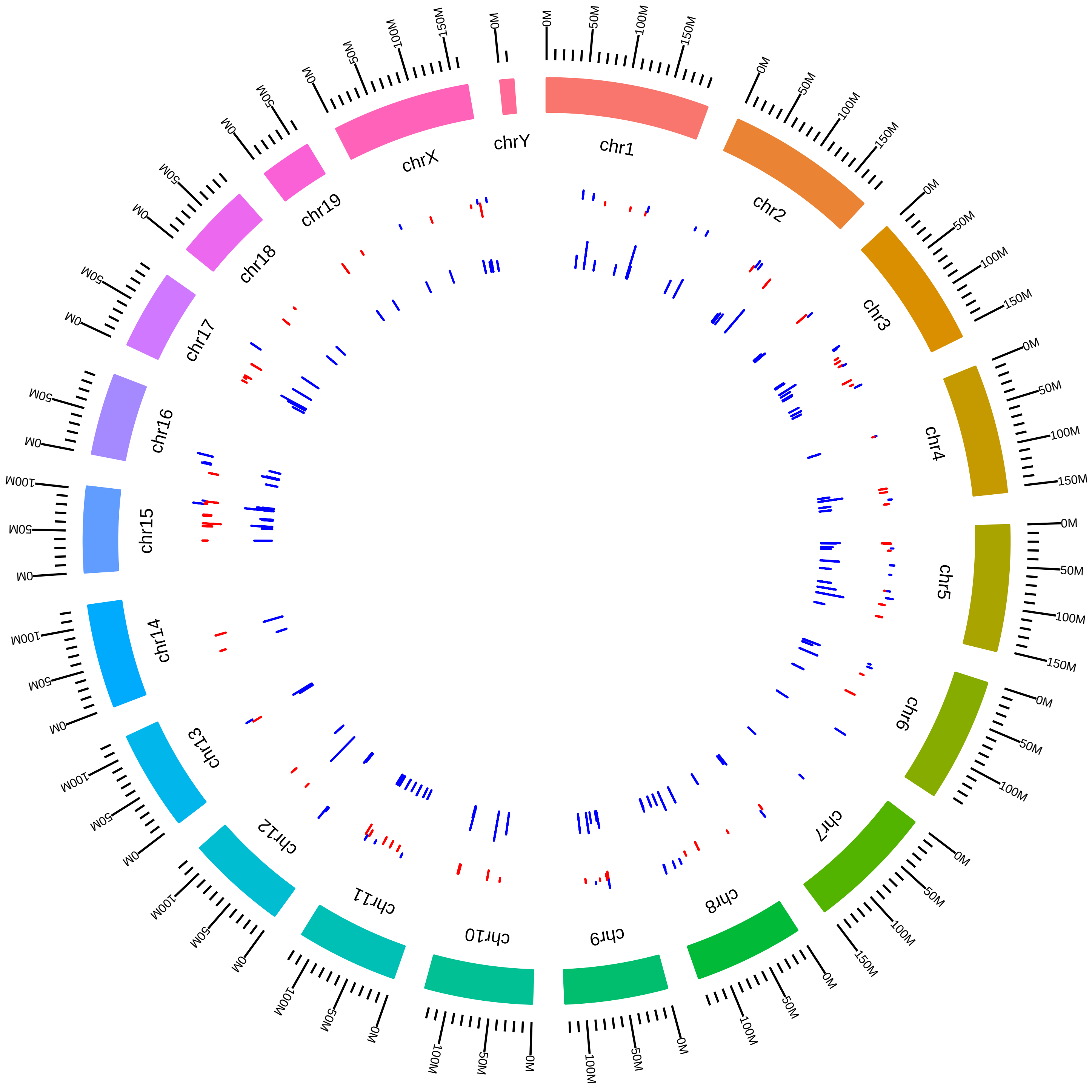

### rsfCirbpRbm3MmJunYan2013_RVsH.pdf

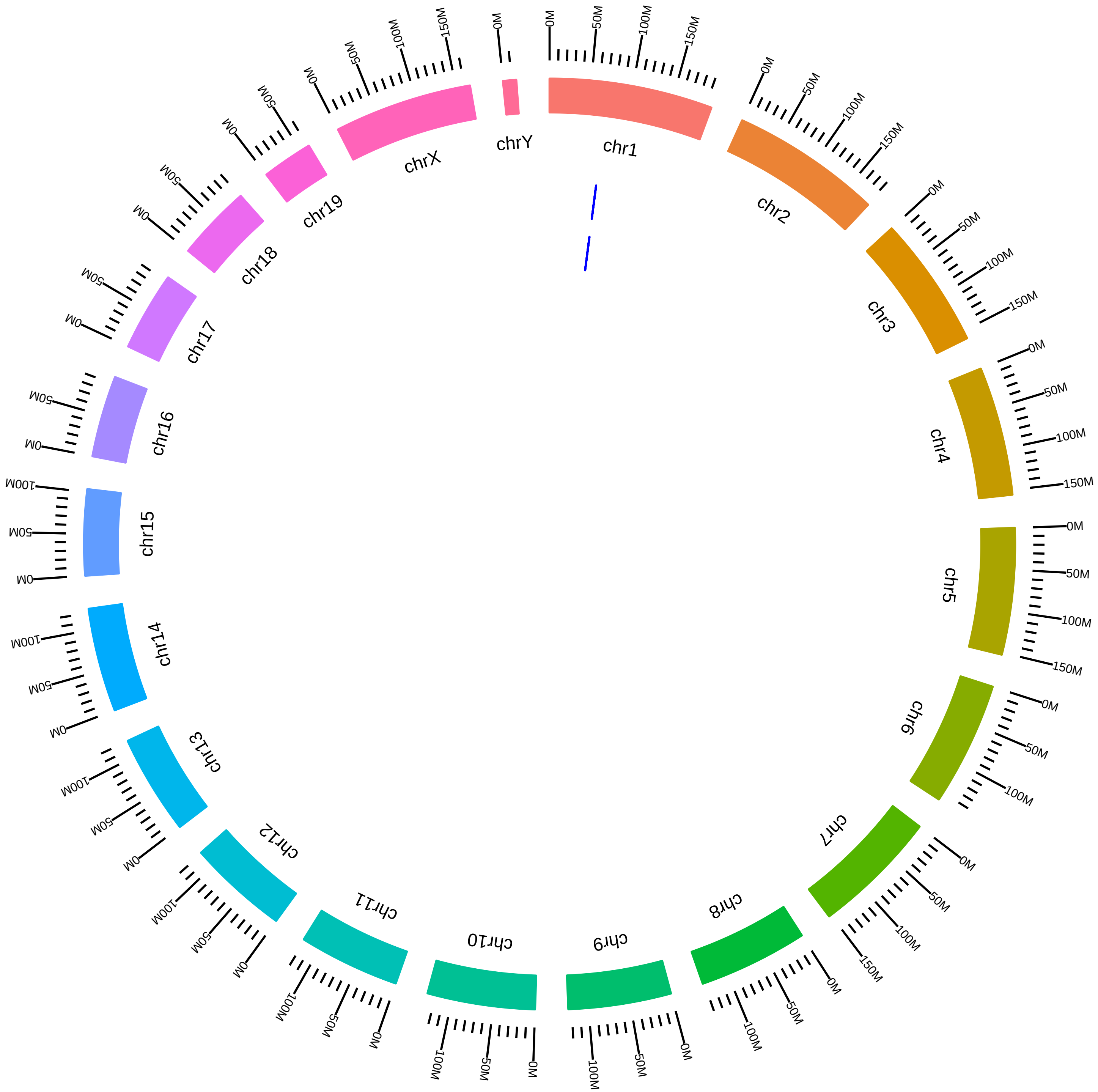

### rsfDdx5MmBStillman_KD_vs_CL.pdf

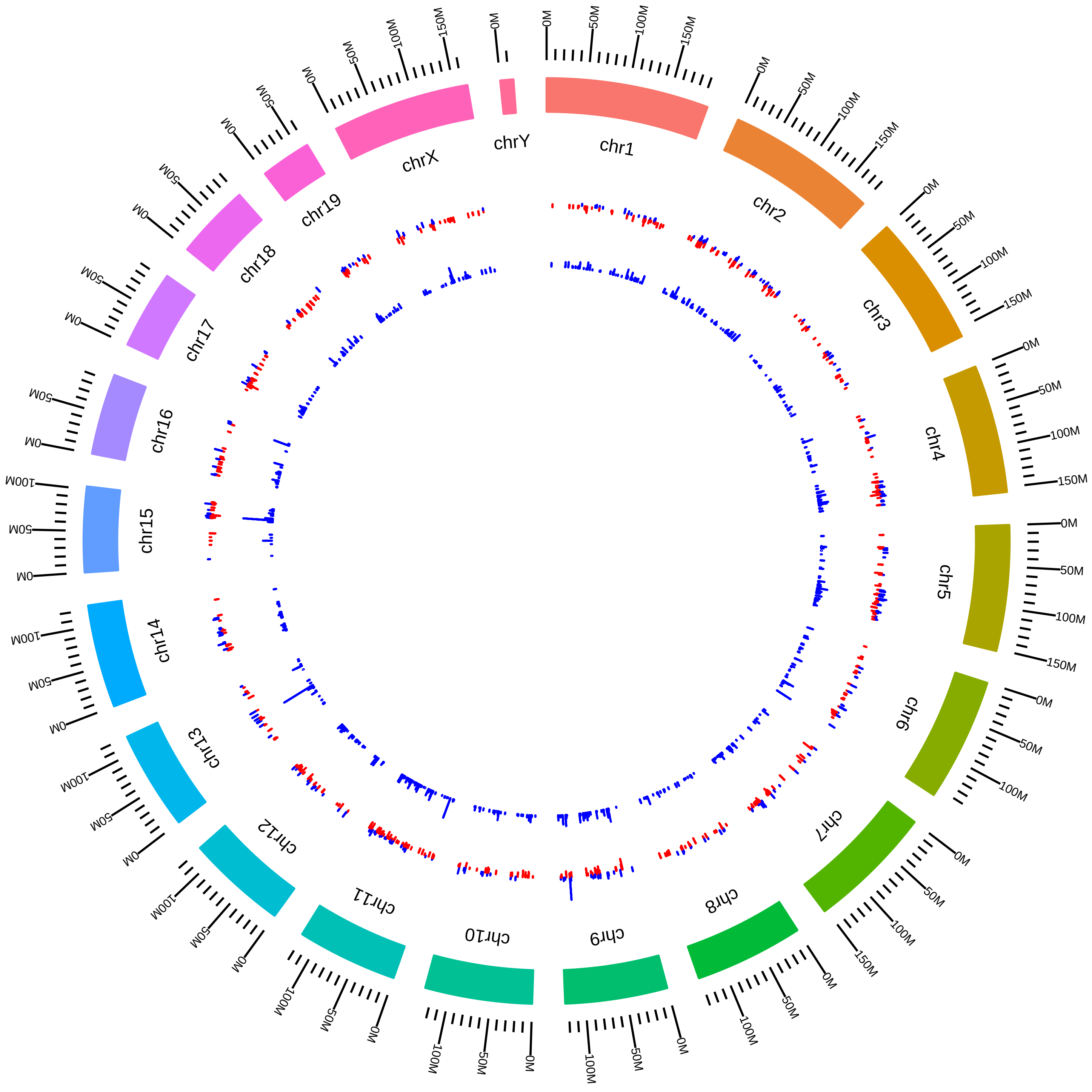

### rsfDdx5MmDRLittman_KO_vs_WT.pdf

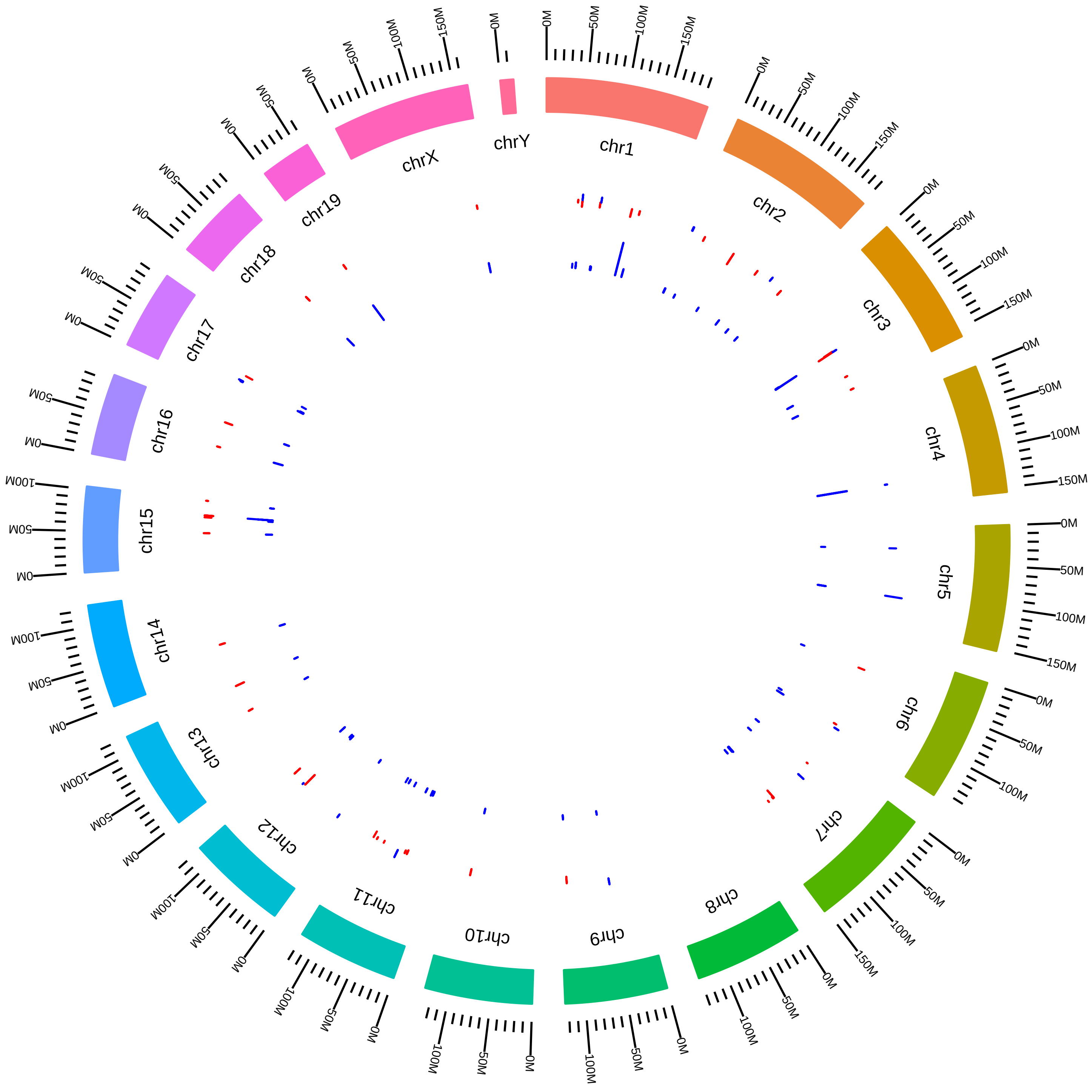

### rsfDdx5MmHongjieYao2017_KVsW.pdf

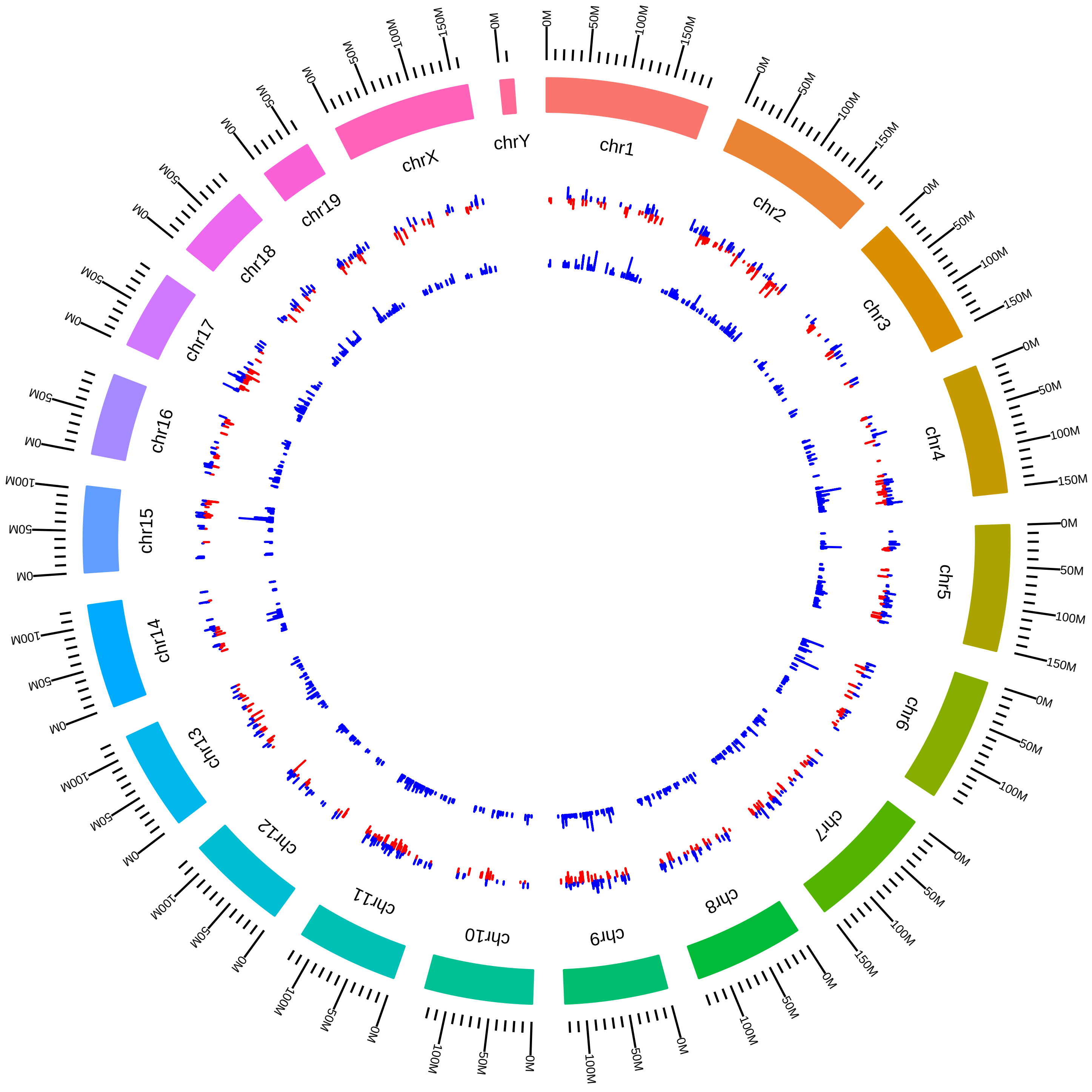

### rsfDyrk1aMmRXavier2015_DVsC.pdf

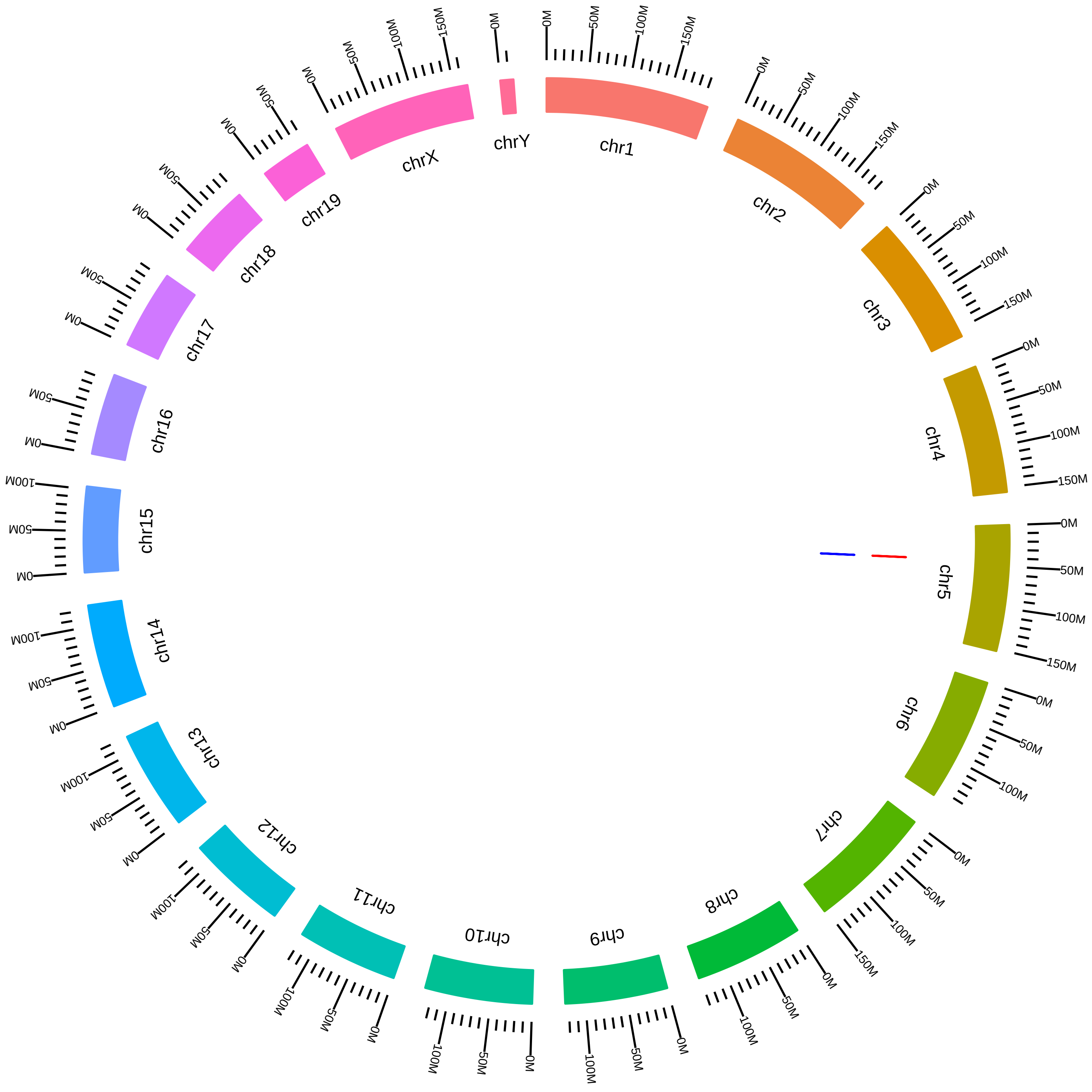

### rsfEif4a3MagohRbm8aMmDSilver2016_EVsC.pdf

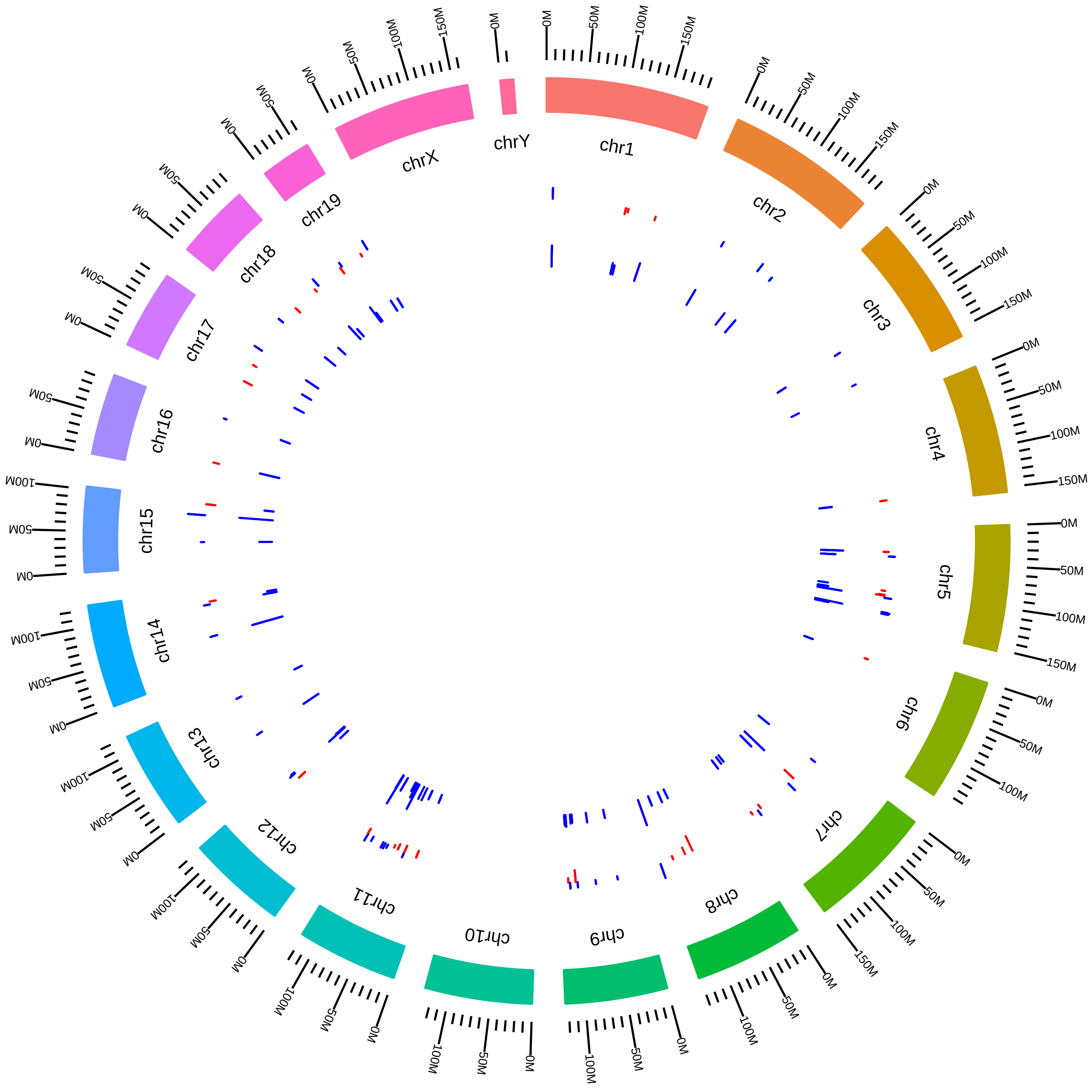

### rsfEif4a3MagohRbm8aMmDSilver2016_MVsC.pdf

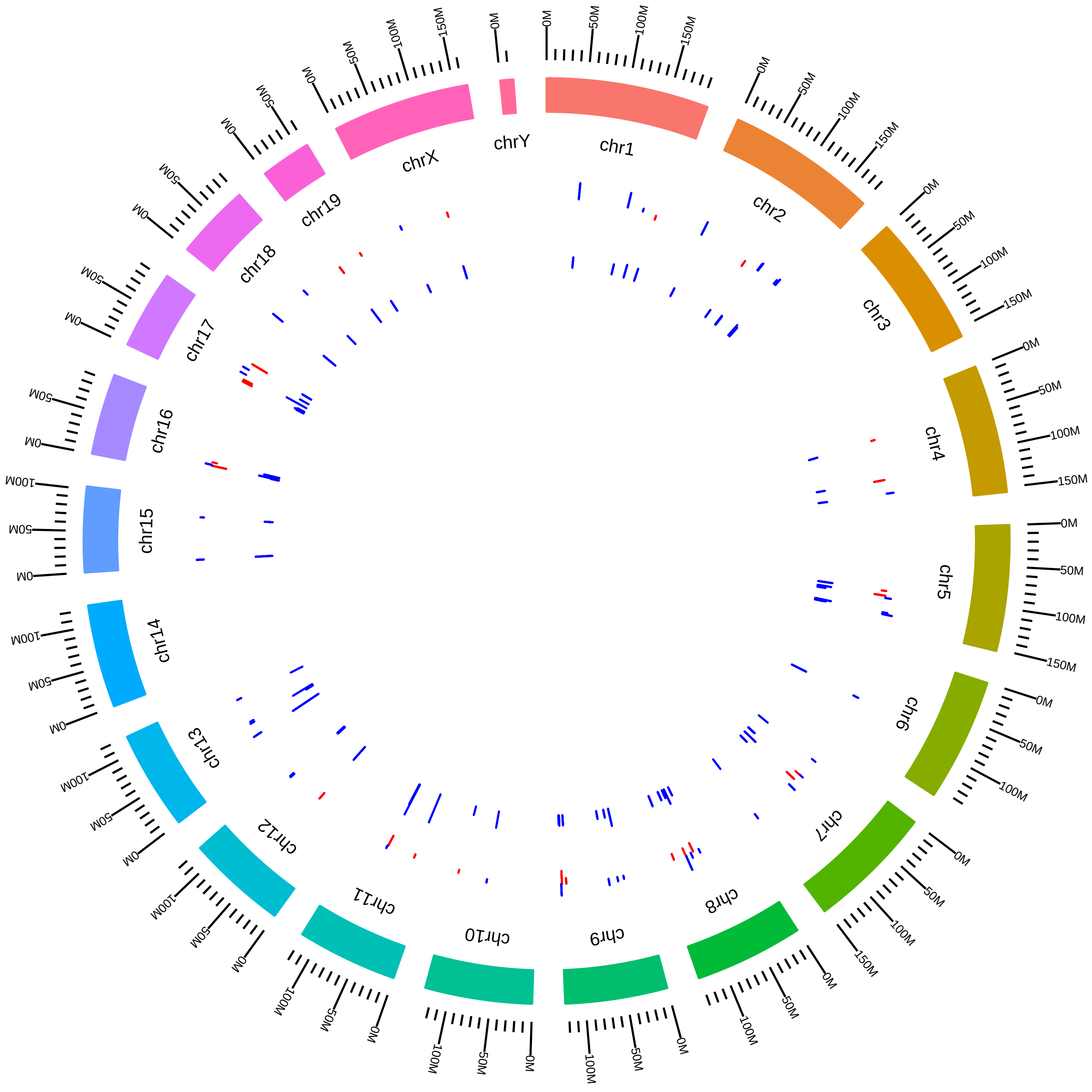

### rsfEif4a3MagohRbm8aMmDSilver2016_RVsC.pdf

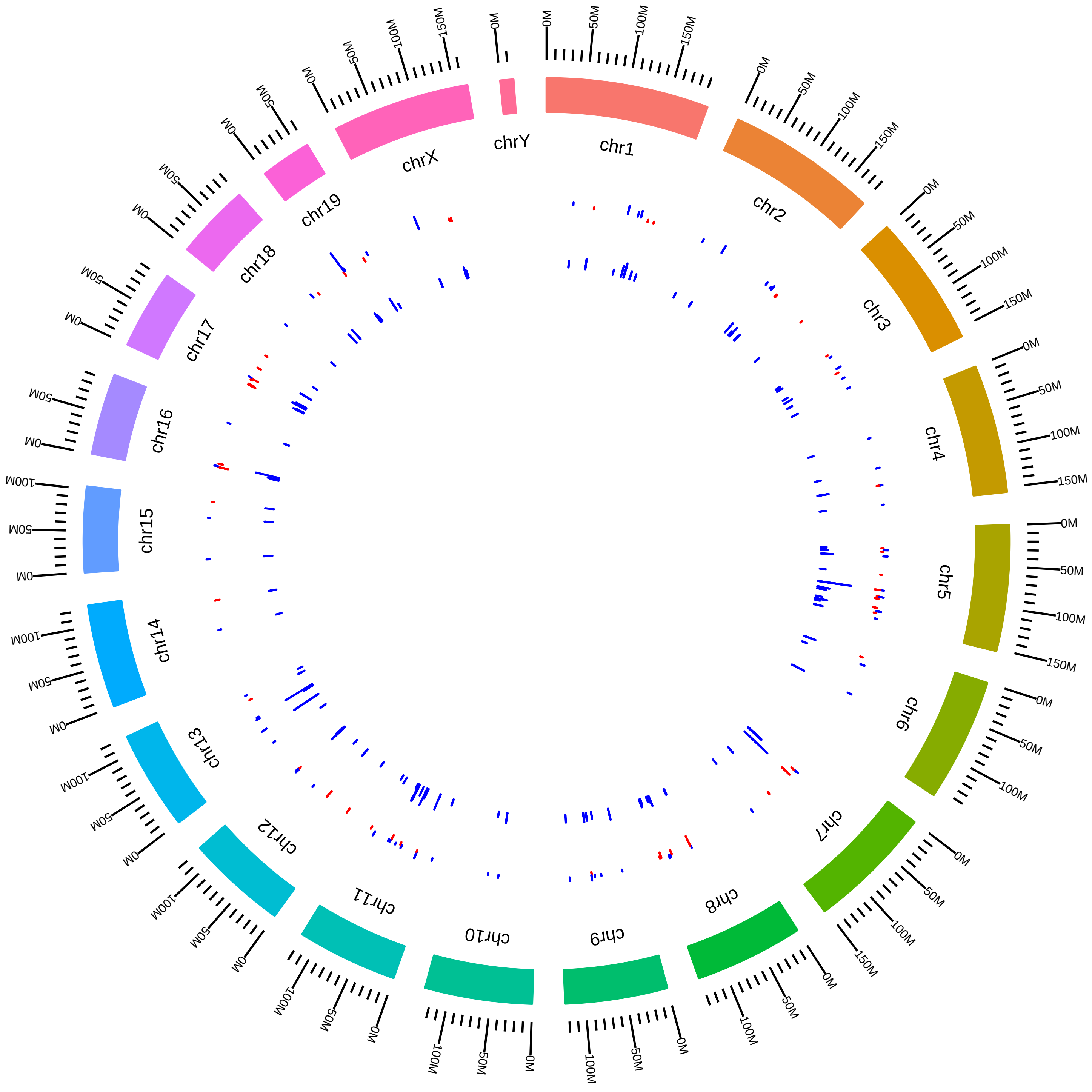

### rsfEsrp1Esrp2MmRCarstens_DKO_vs_WT.pdf

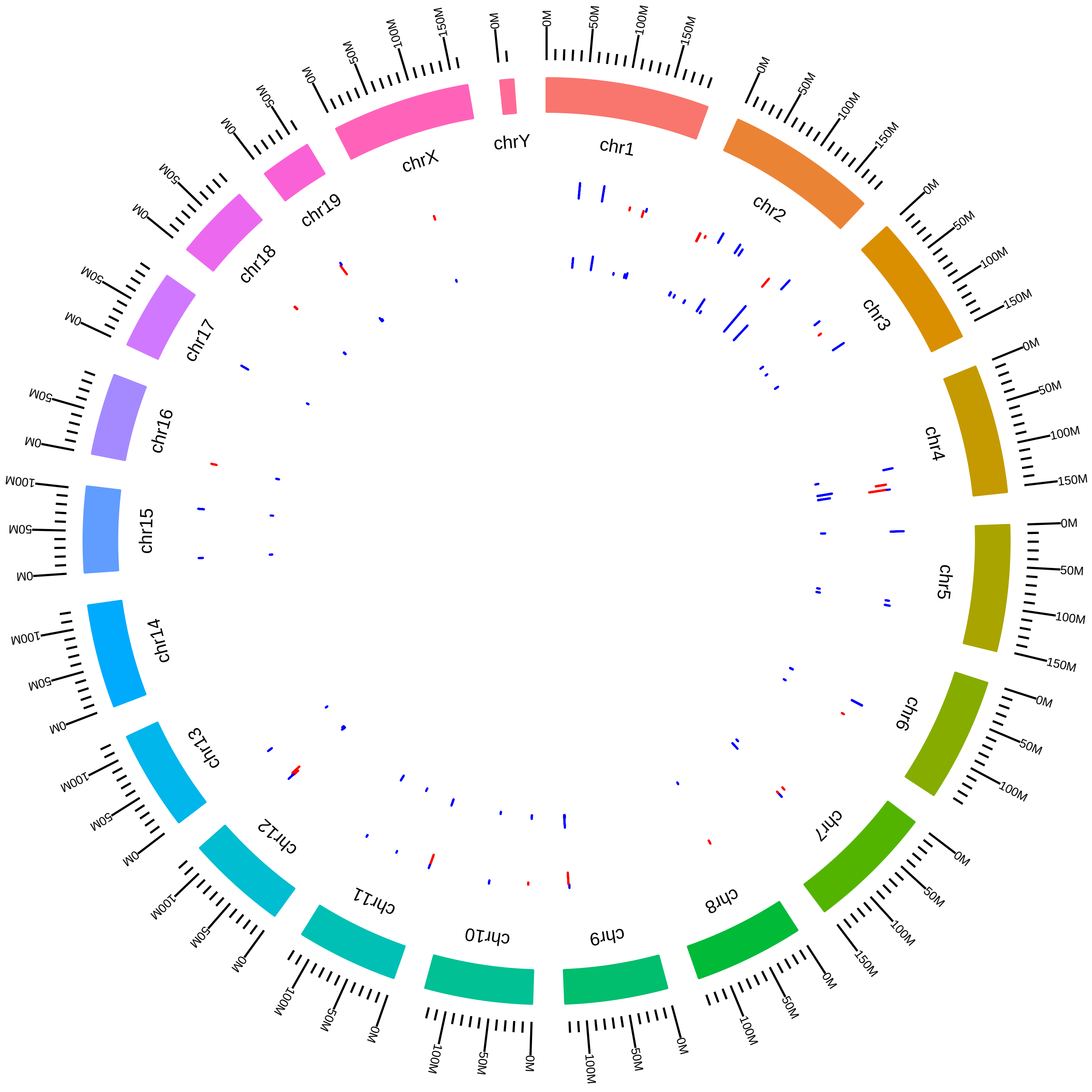

### rsfEsrp1Esrp2MmYiXing2015_DVsH.pdf

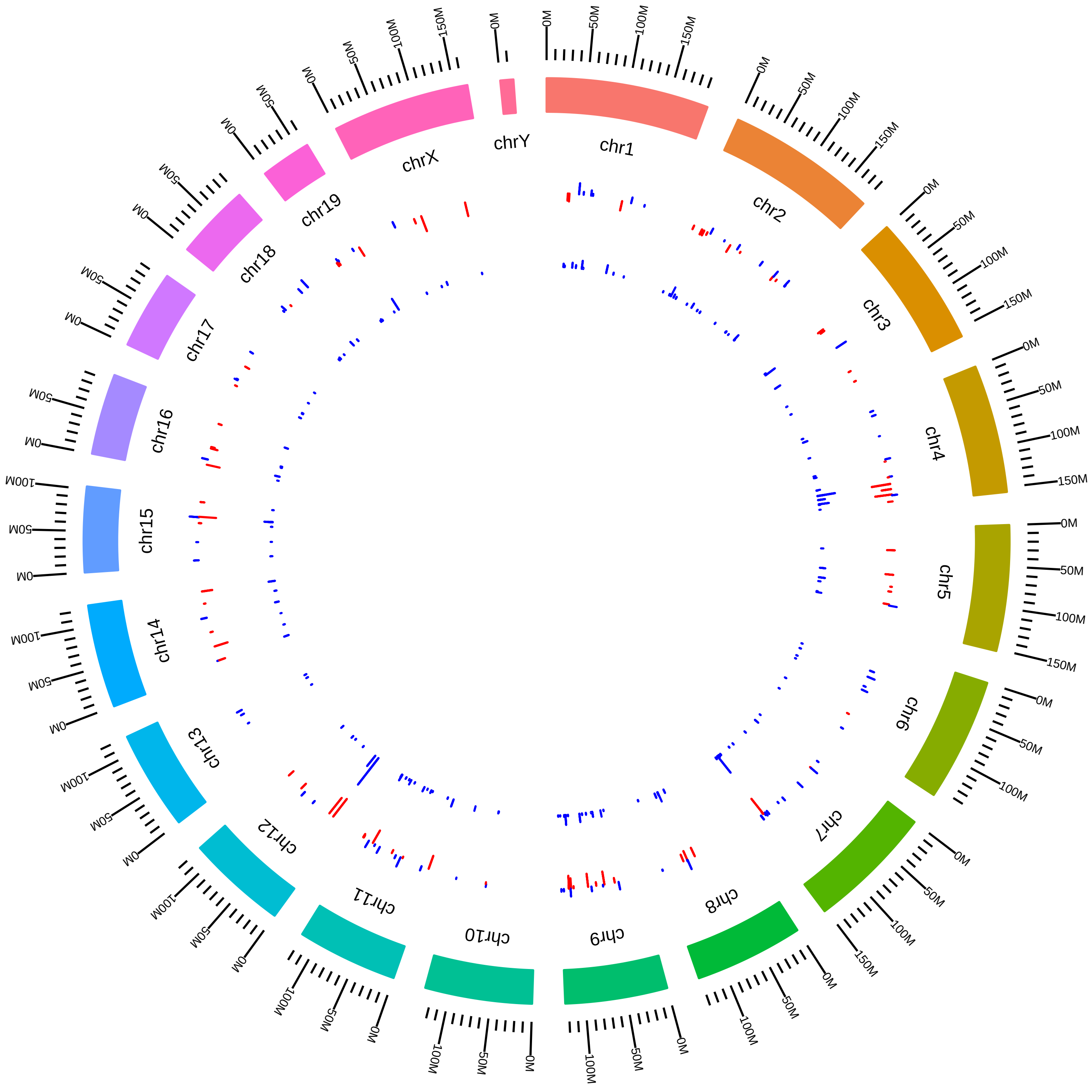

### rsfEsrp1Esrp2MmYiXing2015_DVsK.pdf

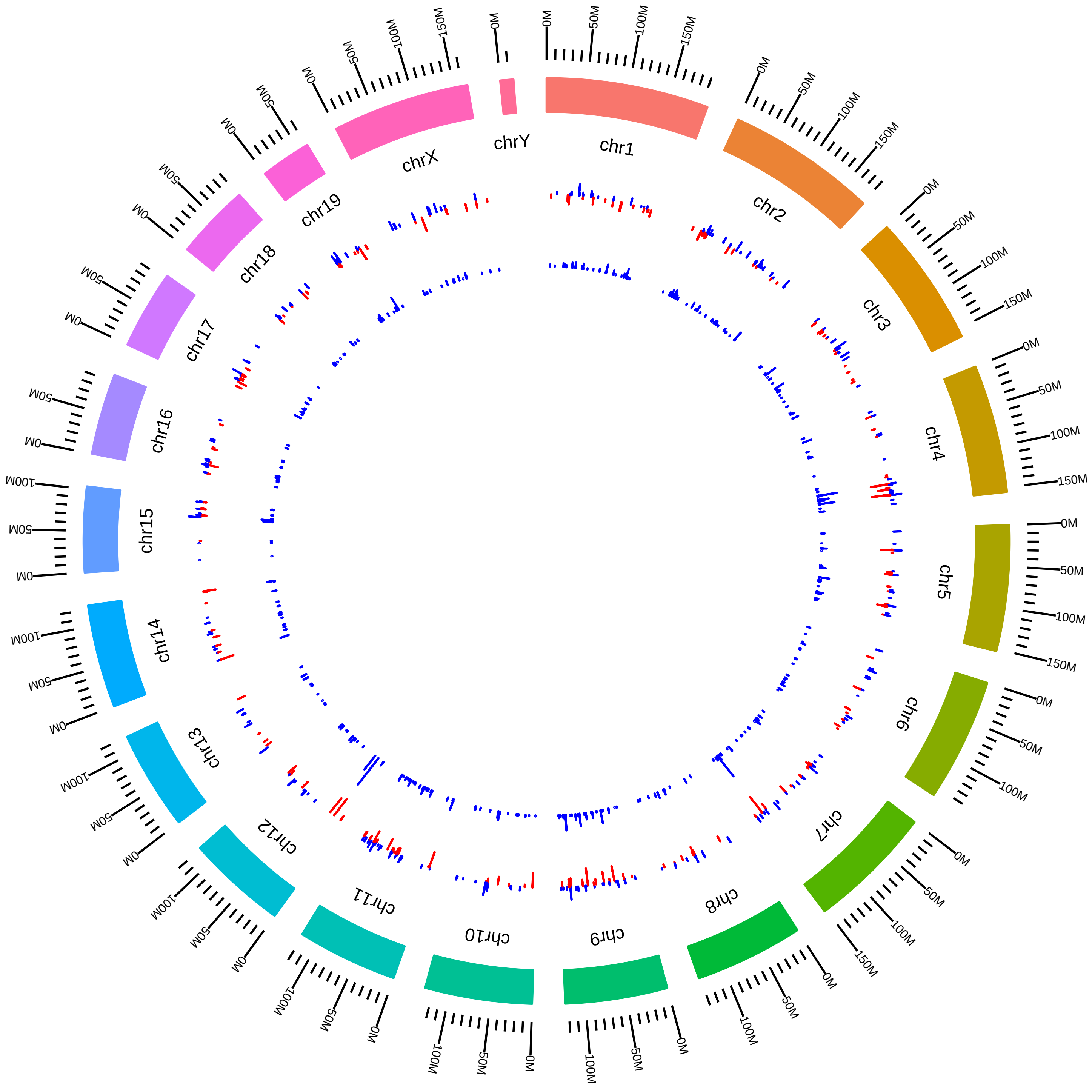

### rsfEsrp1Esrp2MmYiXing2015_DVsW.pdf

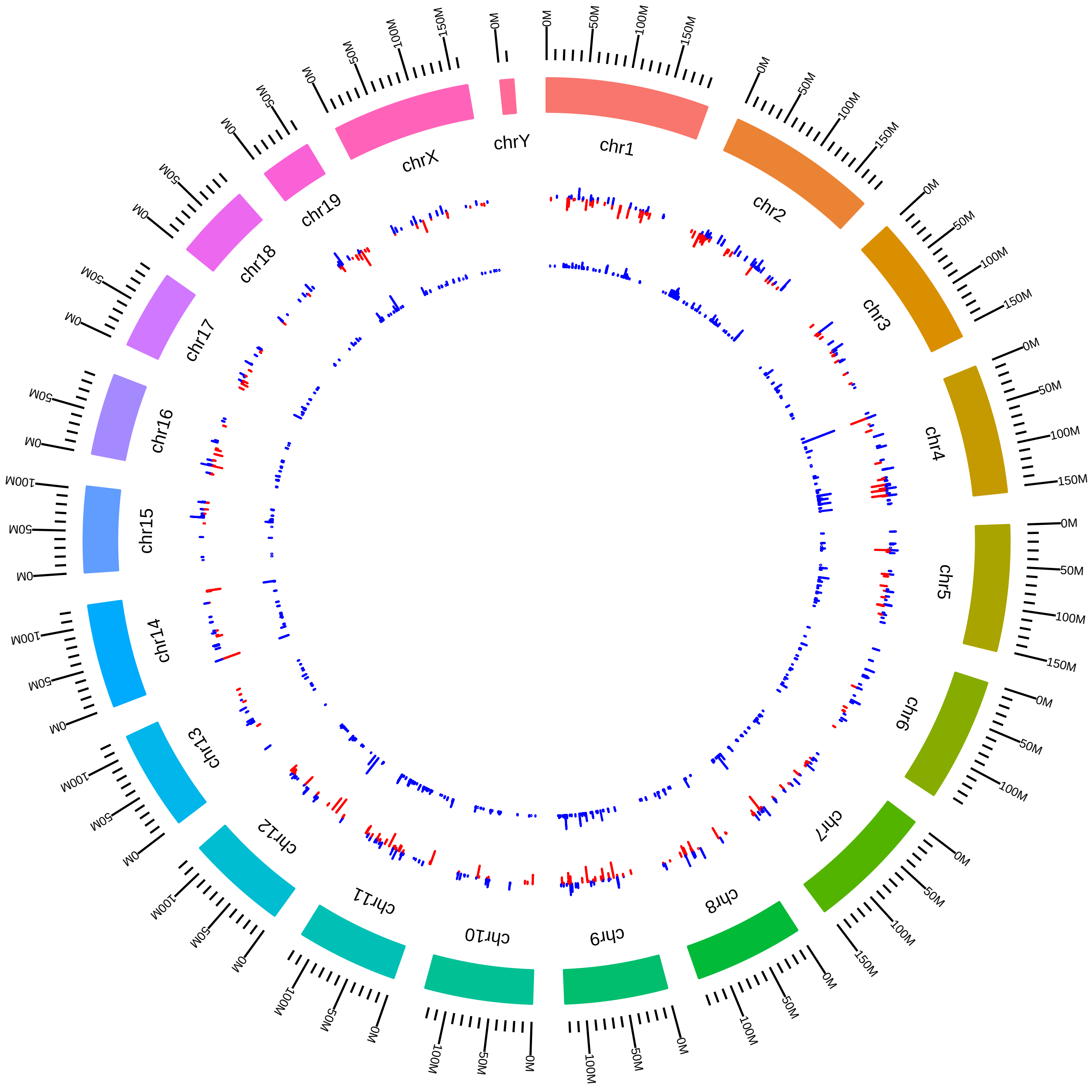

### rsfEsrp1Esrp2MmYiXing2015_HVsK.pdf

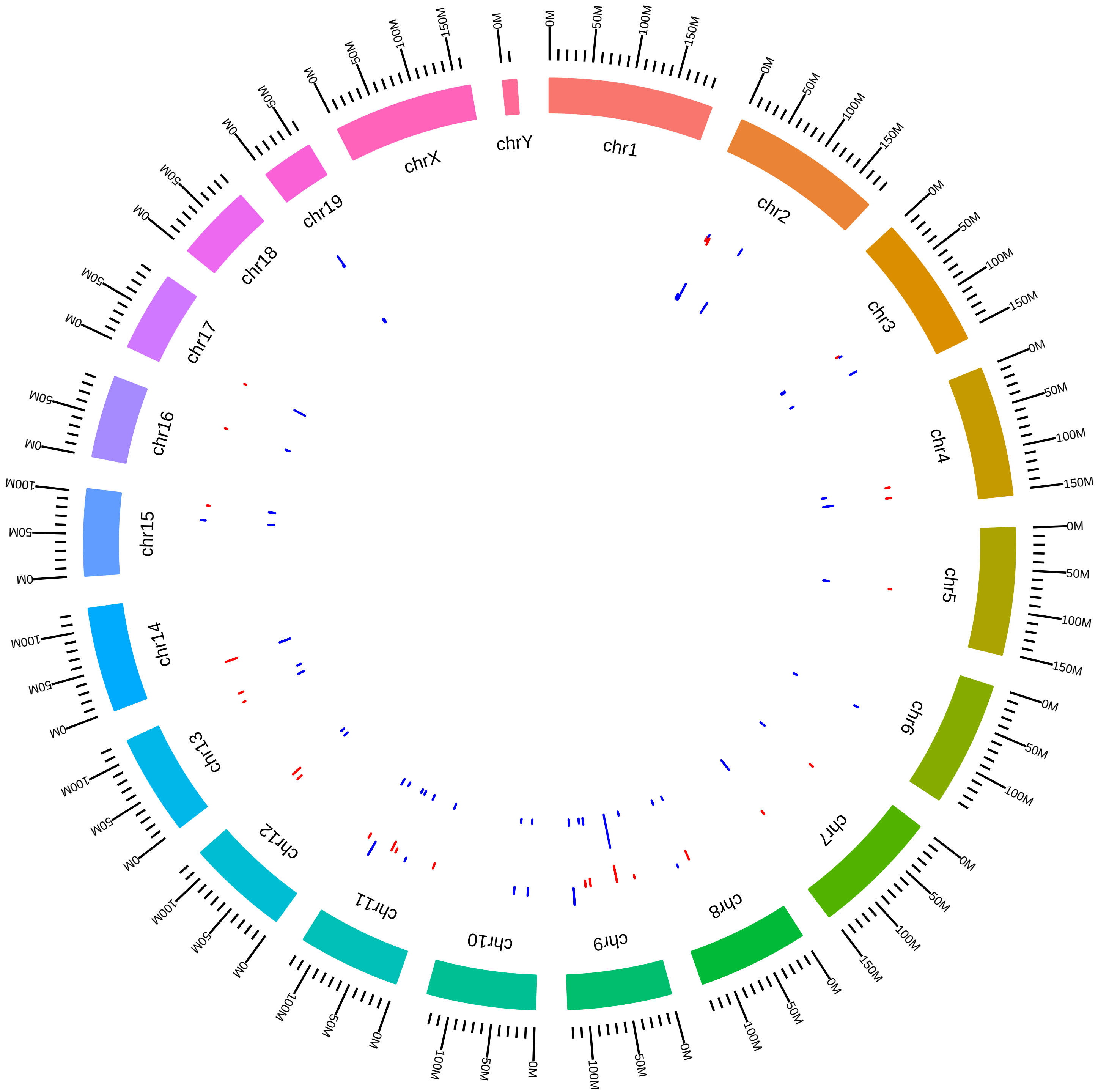

### rsfEsrp1Esrp2MmYiXing2015_HVsW.pdf

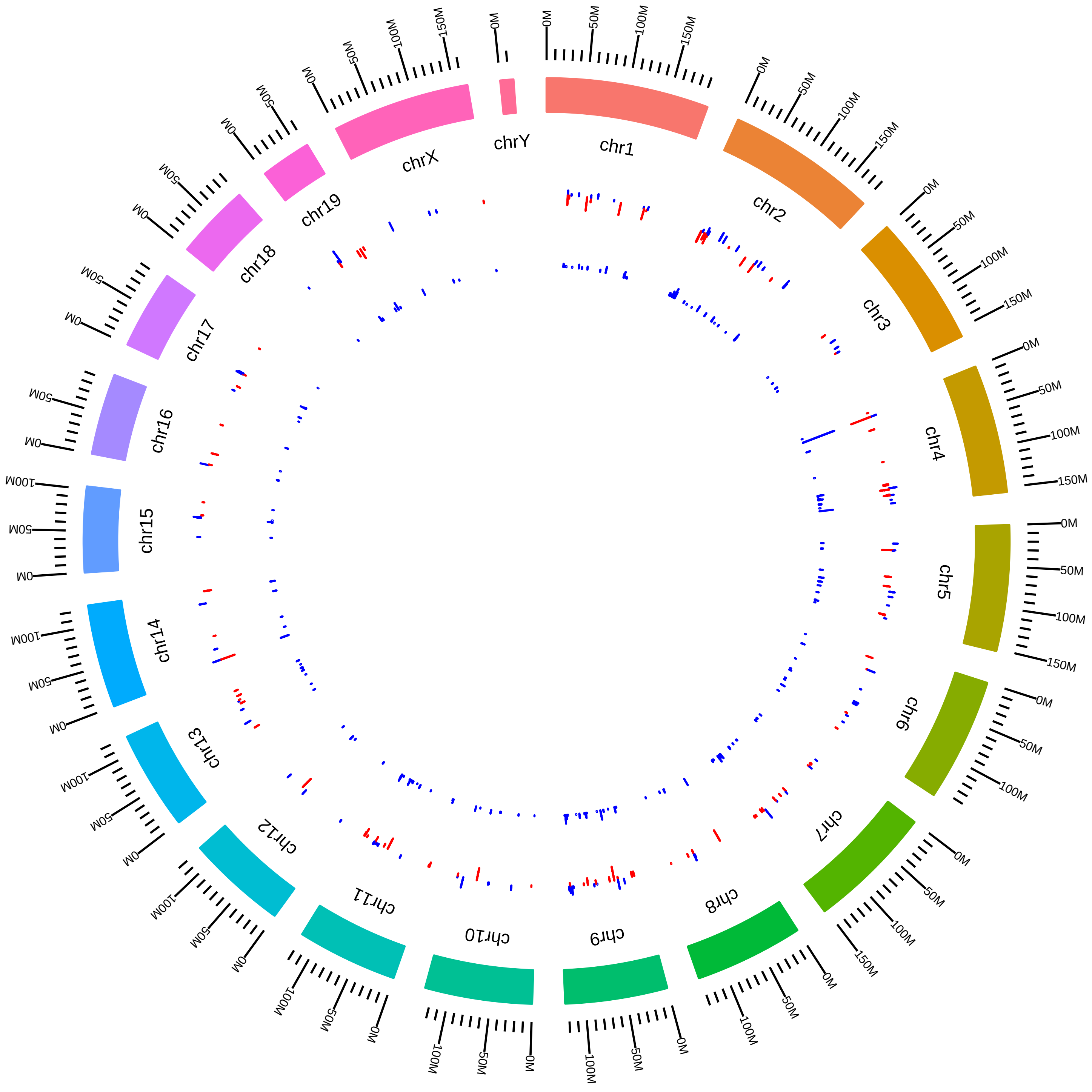

### rsfEsrp1Esrp2MmYiXing2015_KVsW.pdf

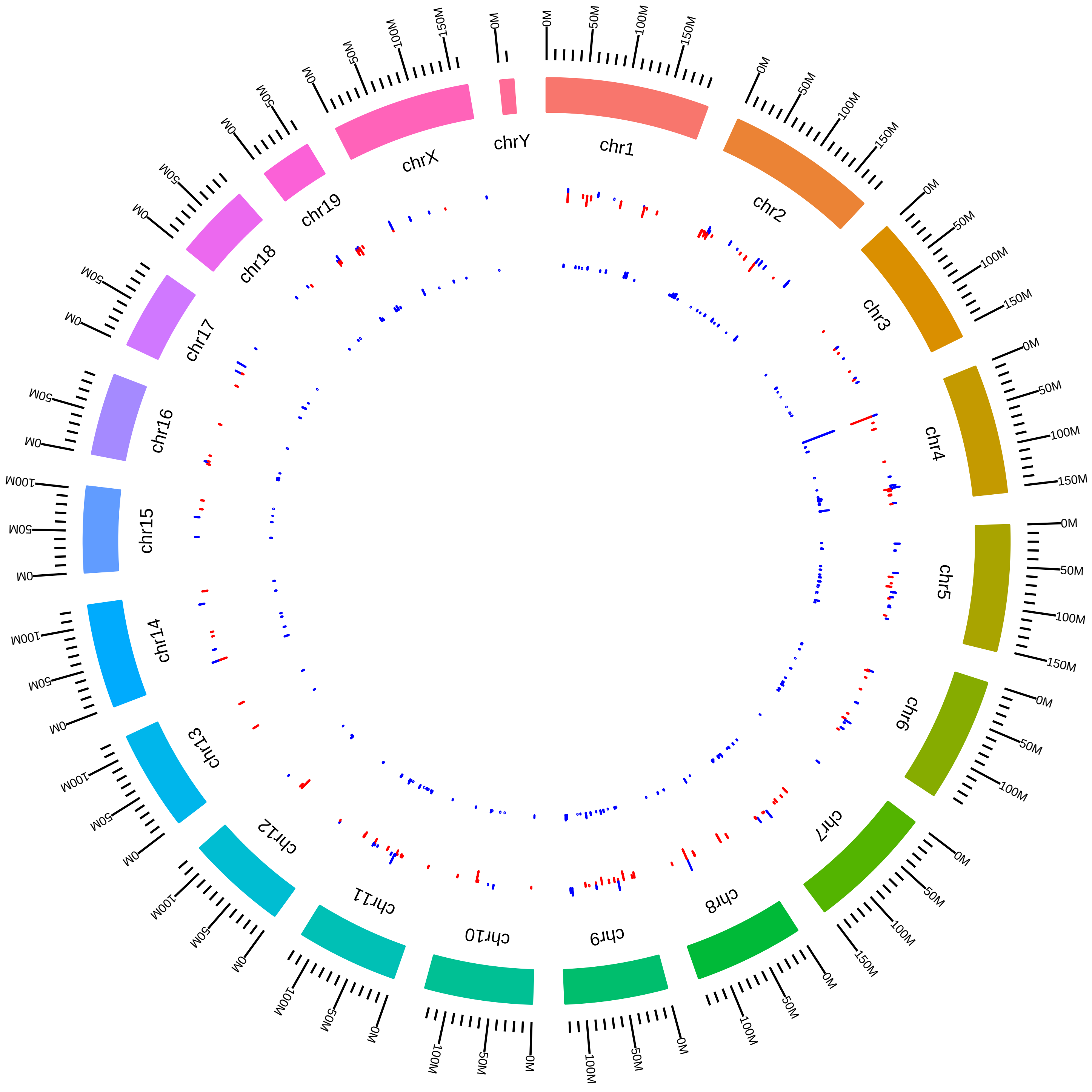

### rsfEsrp1MmRCarstens2016_KVsC.pdf

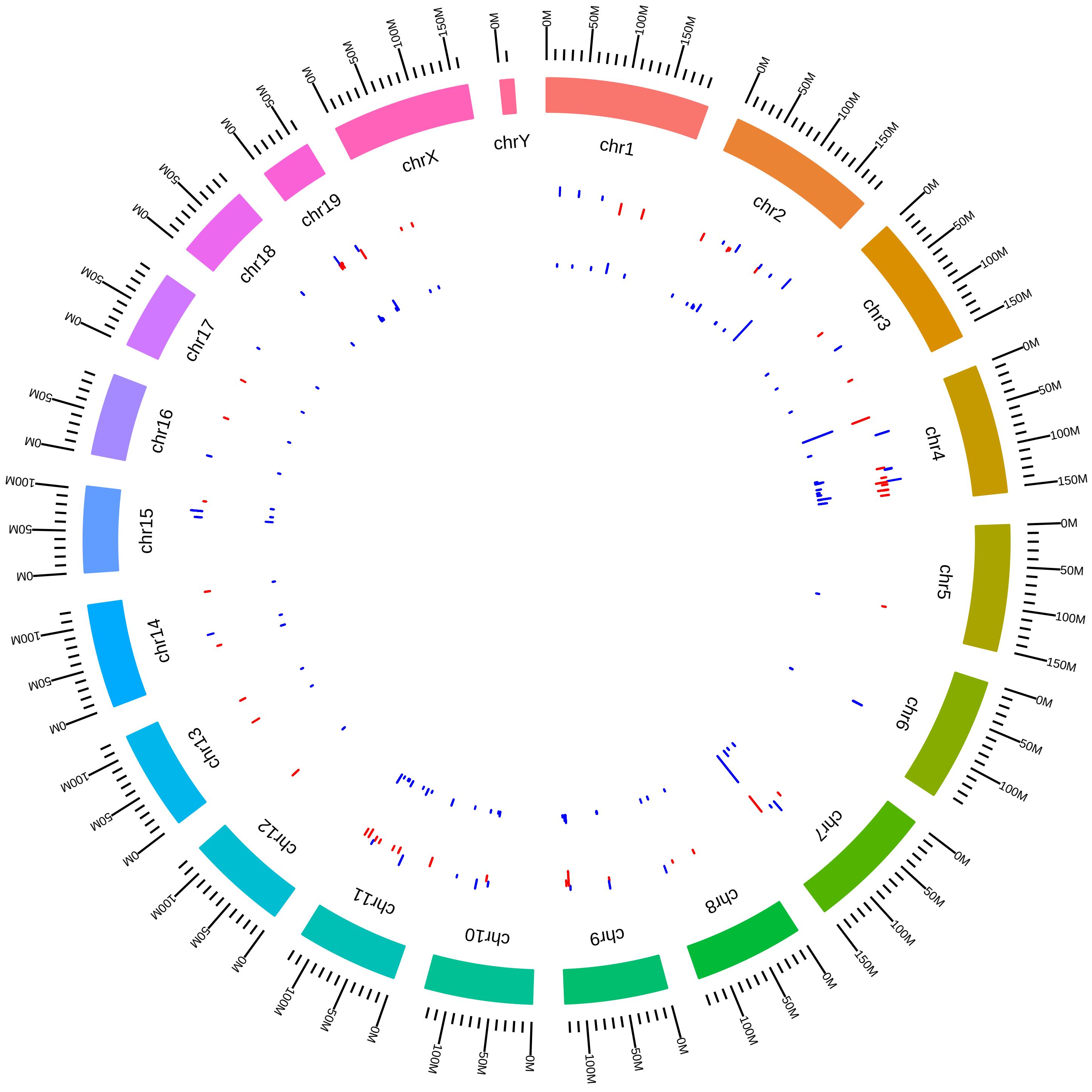

### rsfHnrnpa2b1MmGYeo2016_HVsC.pdf

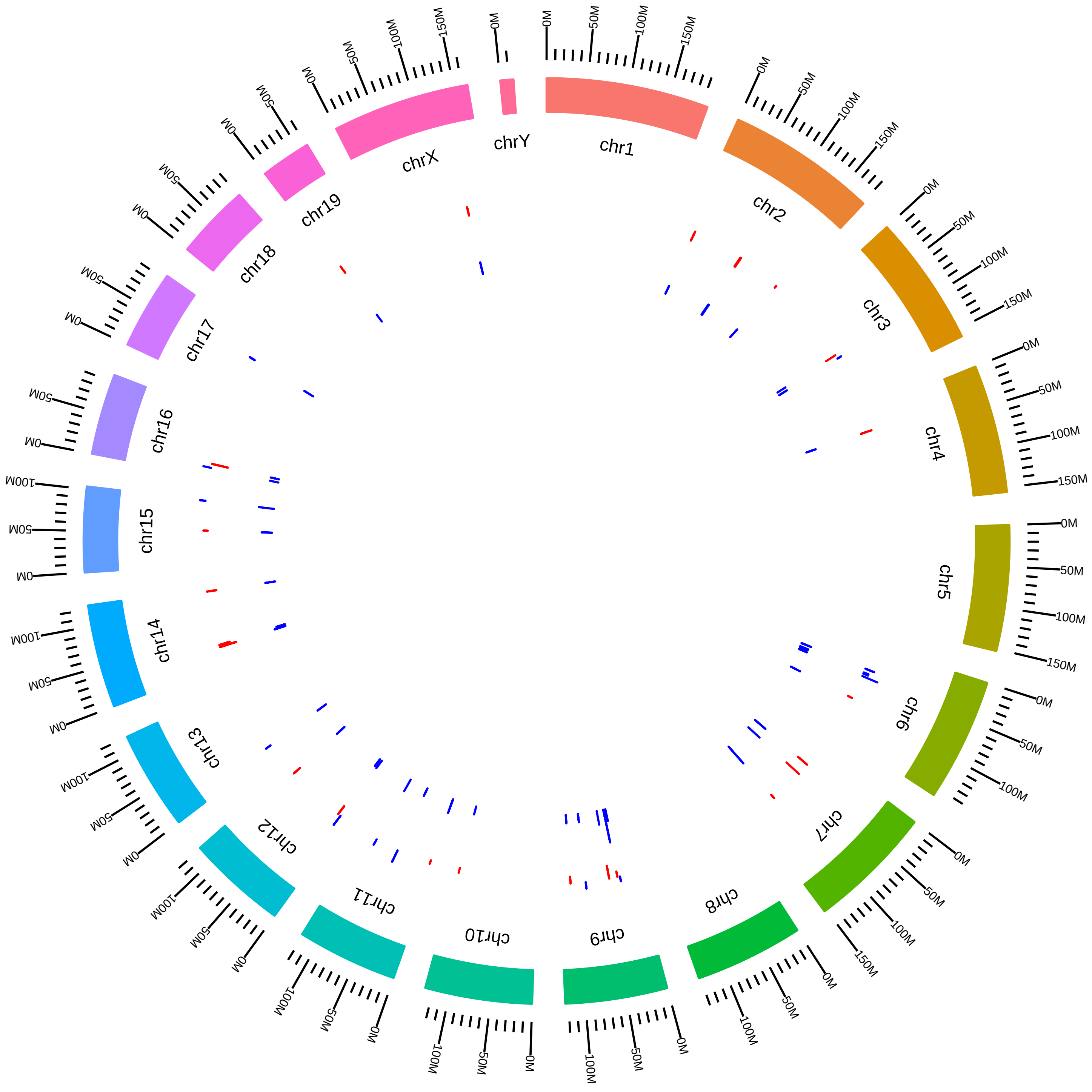

### rsfHnrnph1MmCBryant_FKVsFW.pdf

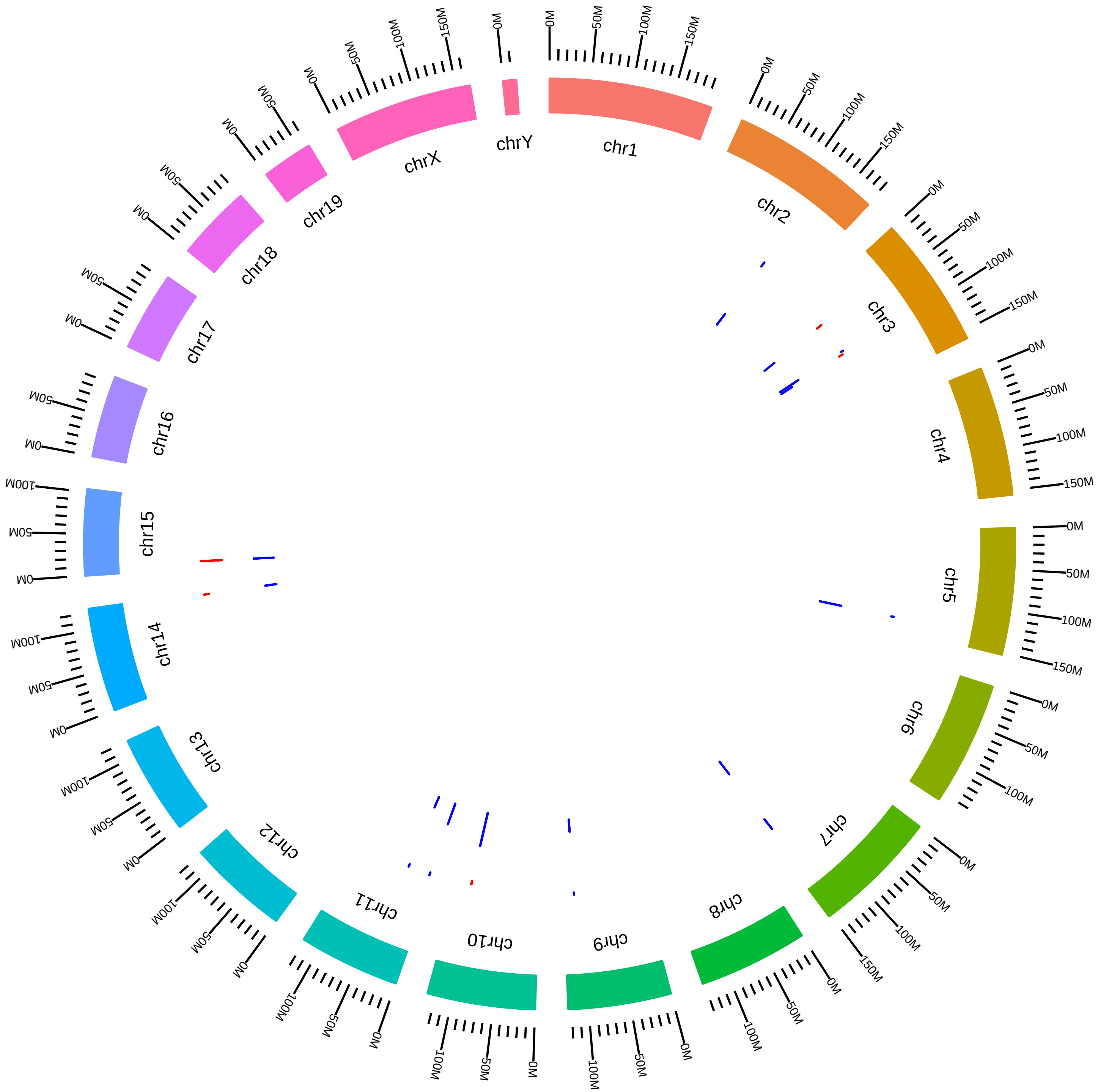

### rsfHnrnph1MmCBryant_KVsW.pdf

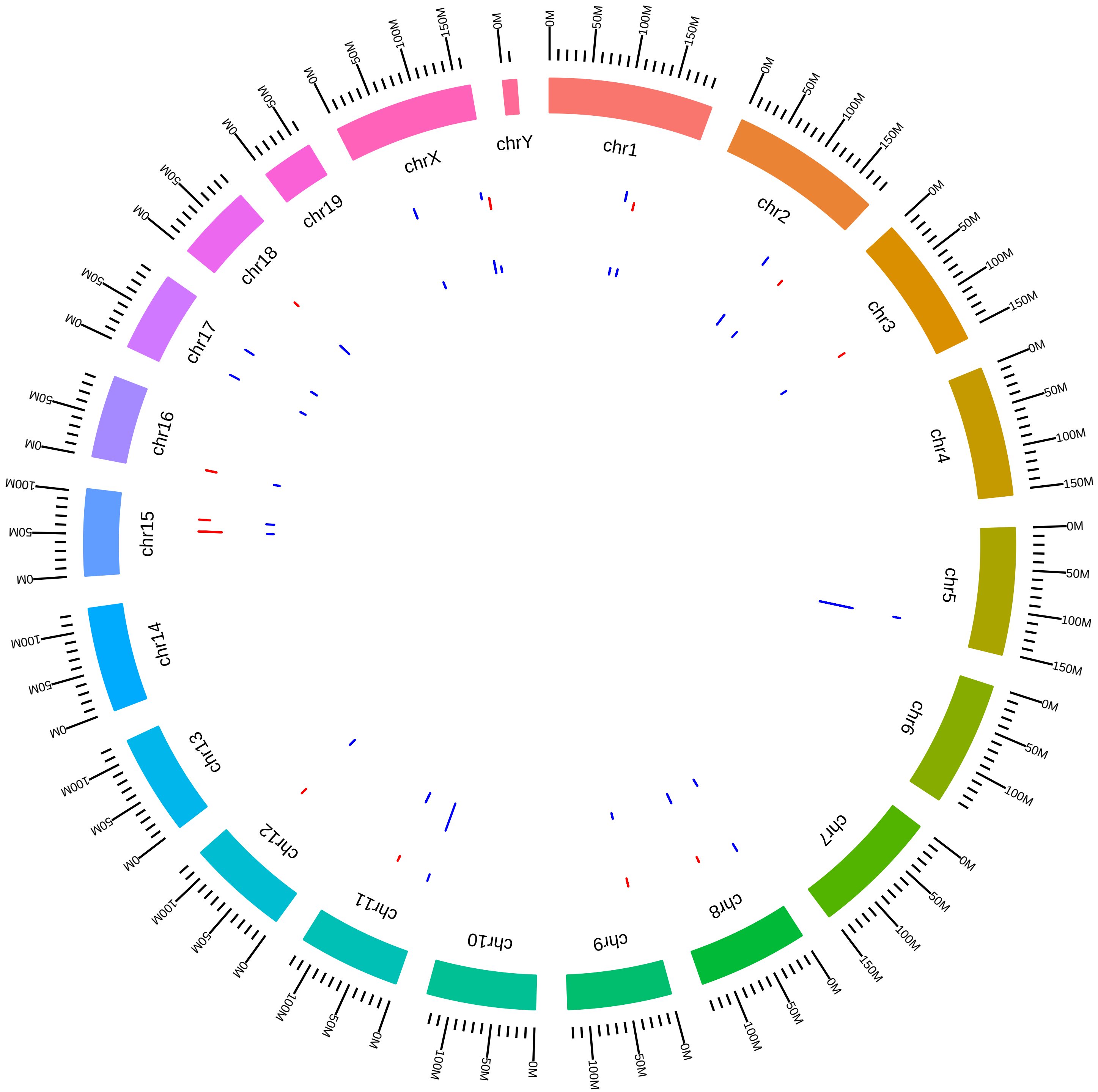

### rsfHnrnph1MmCBryant_MKVsMW.pdf

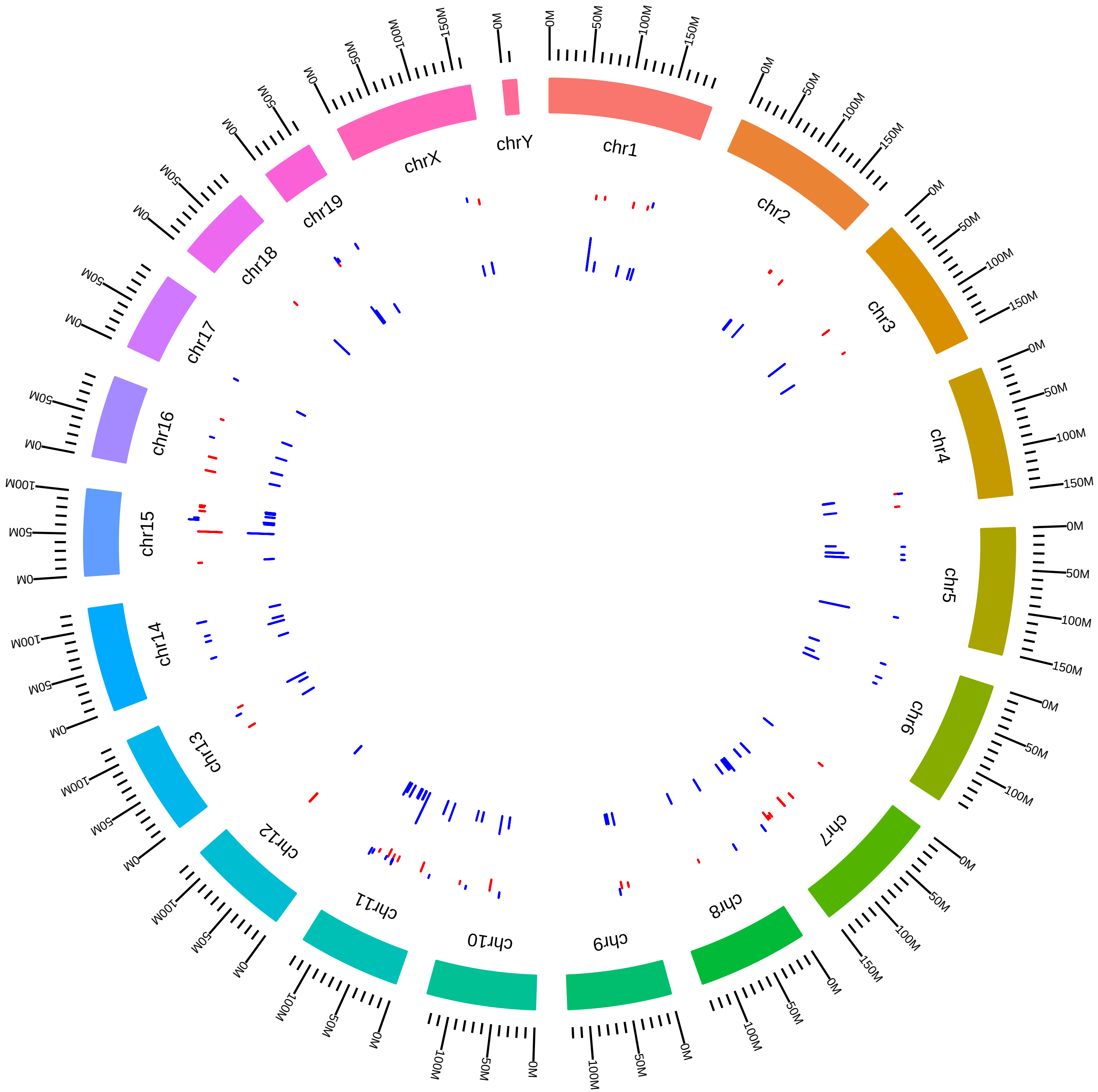

### rsfHnrnpkMmMLorincz2015_DVsC.pdf

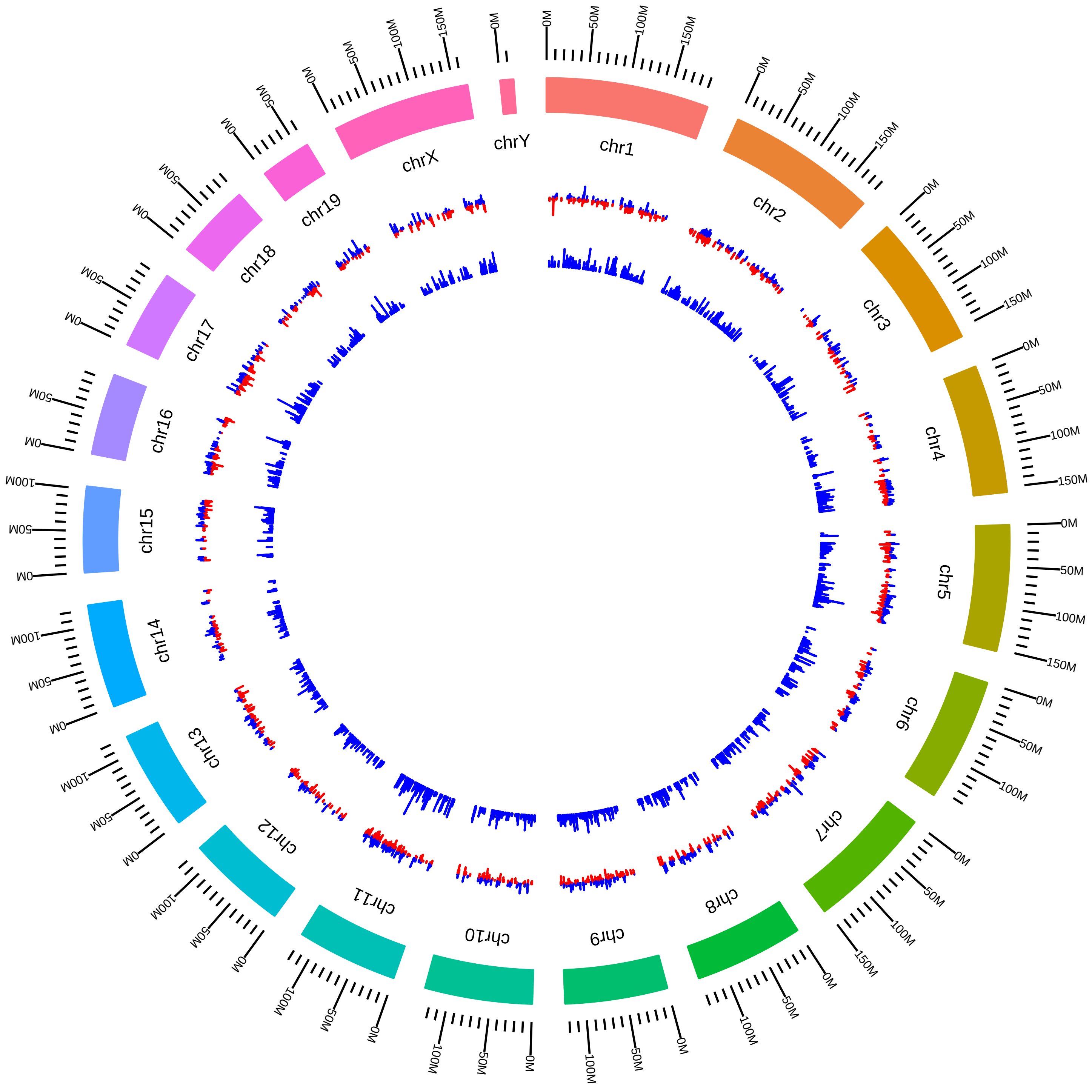

### rsfHnrnplMmTMoroy2016_KVsW.pdf

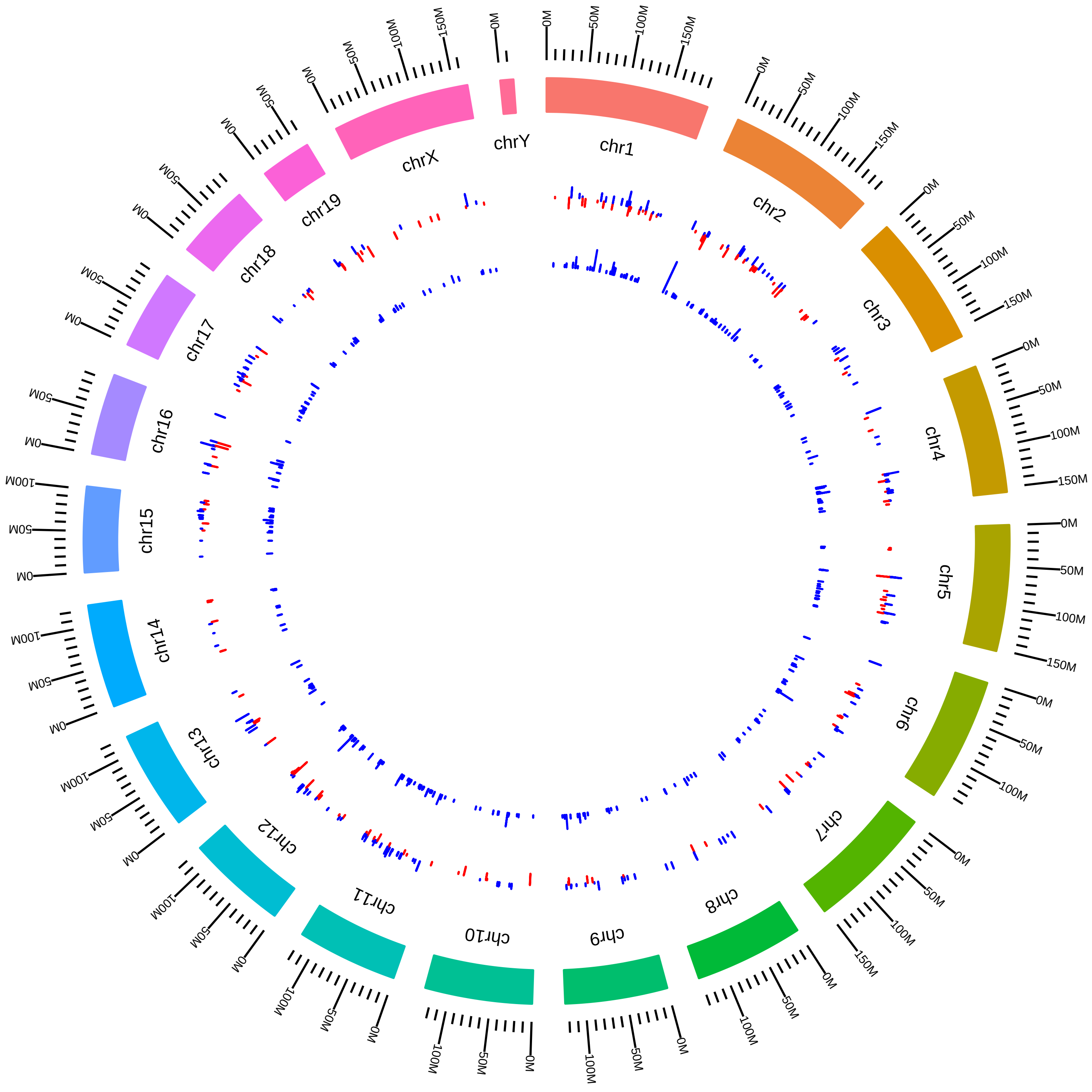

### rsfHnrnpuMmTManiatis_MT_vs_WT.pdf

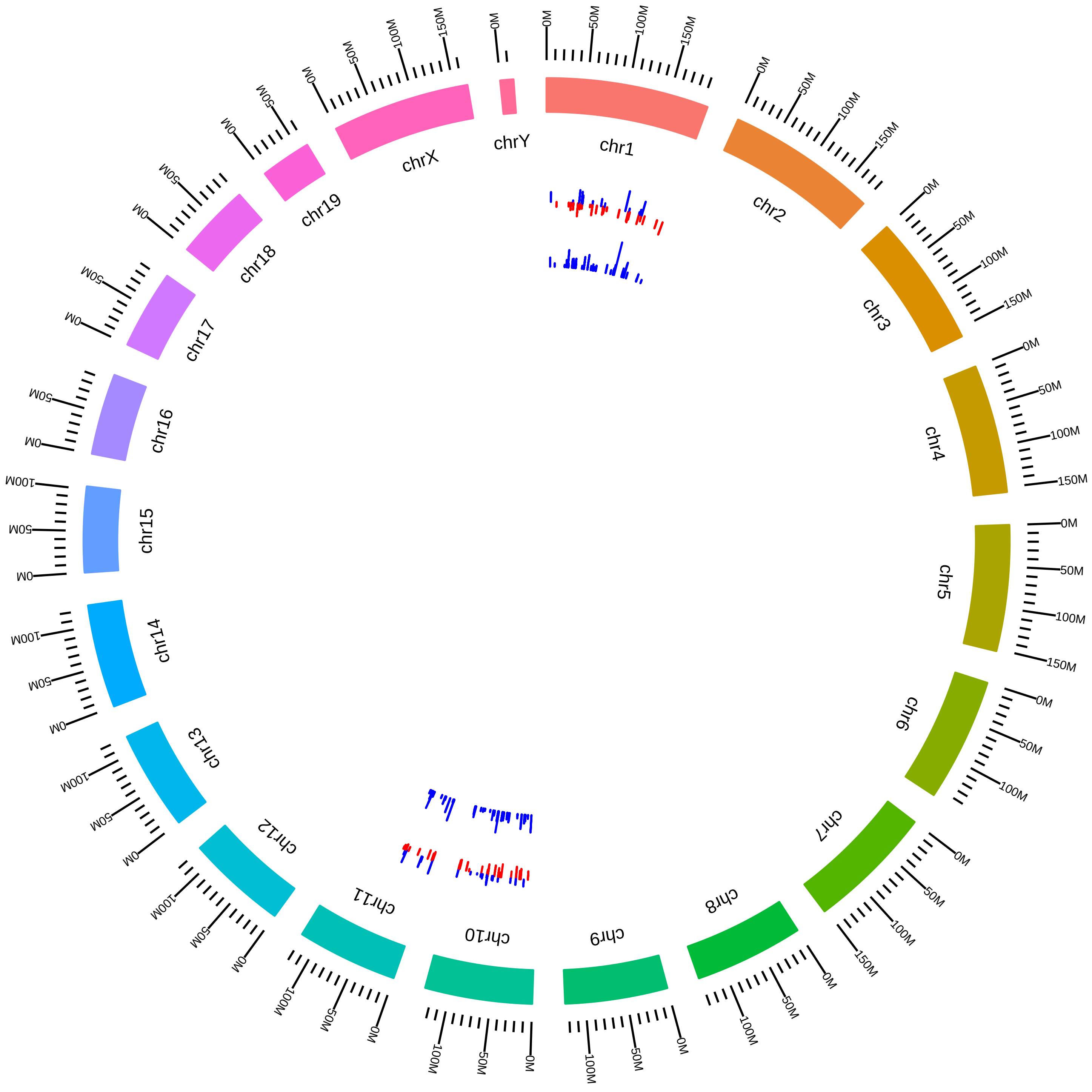

### rsfJmjd6MmFYoshinori2015_KVsW.pdf

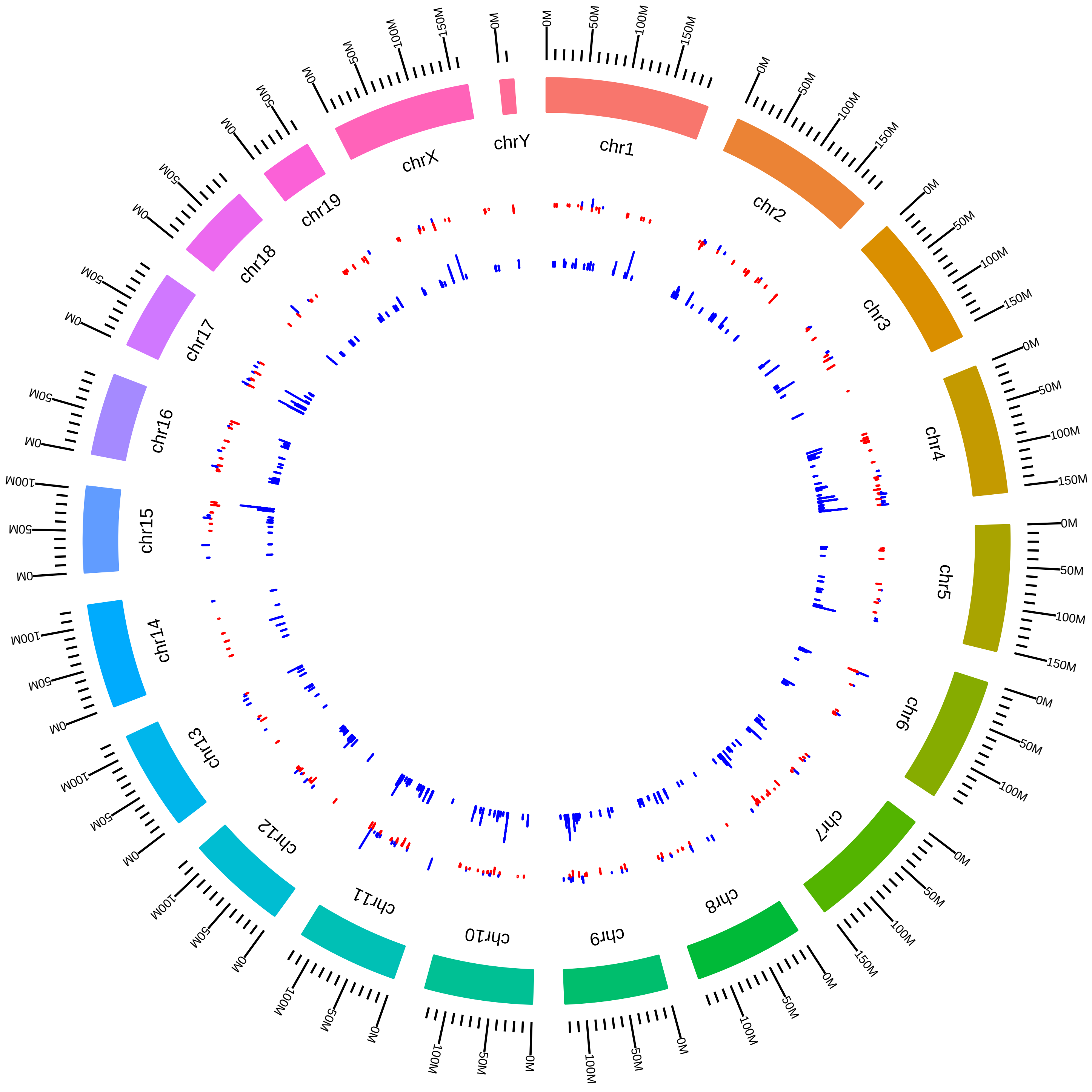

### rsfKhdrbs3MmDElliott2016_KVsW.pdf

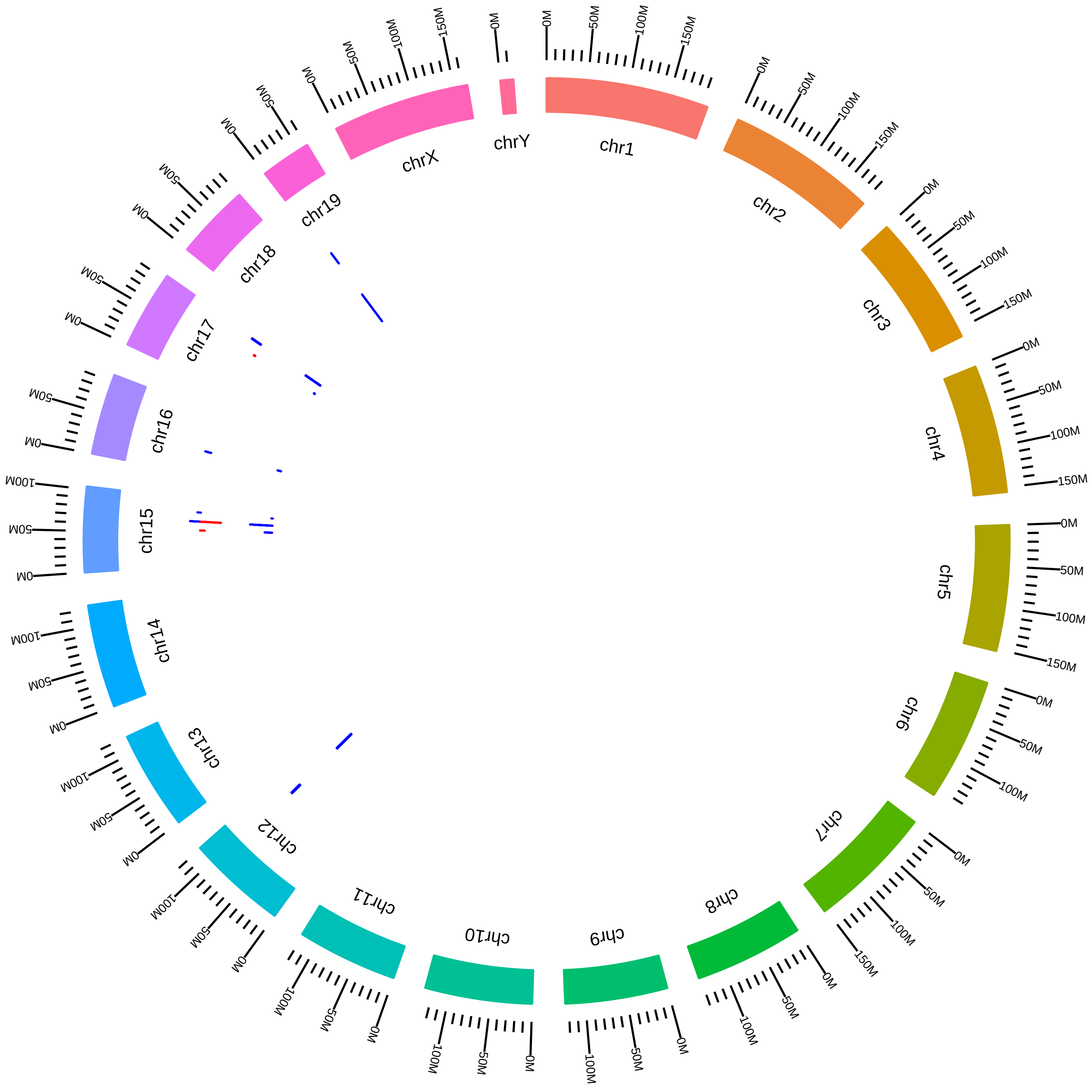

### rsfMagohMutMmDSilver2016_HVsW.pdf

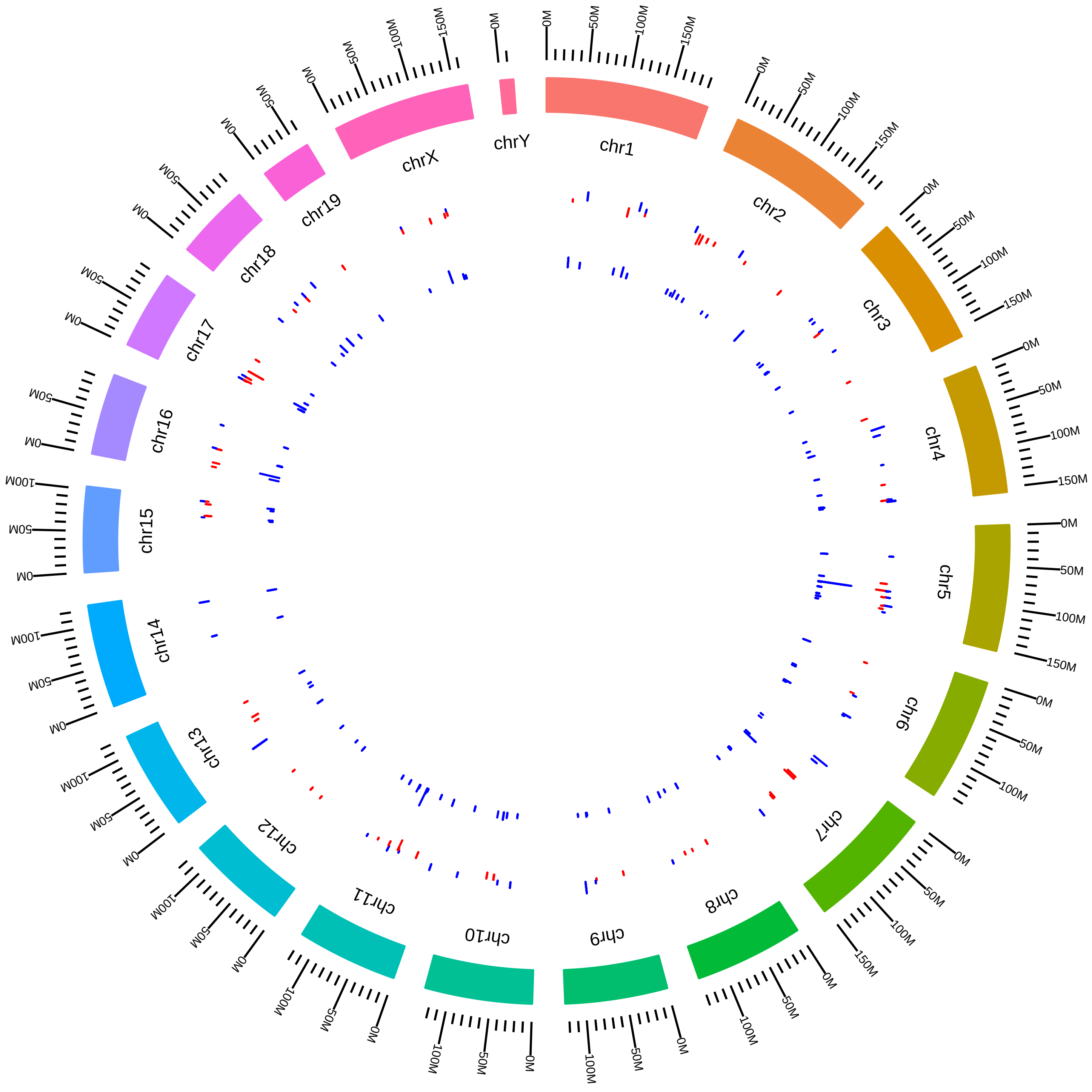

### rsfMalat1MmDLSpectorOrganoid_T_vs_C.pdf

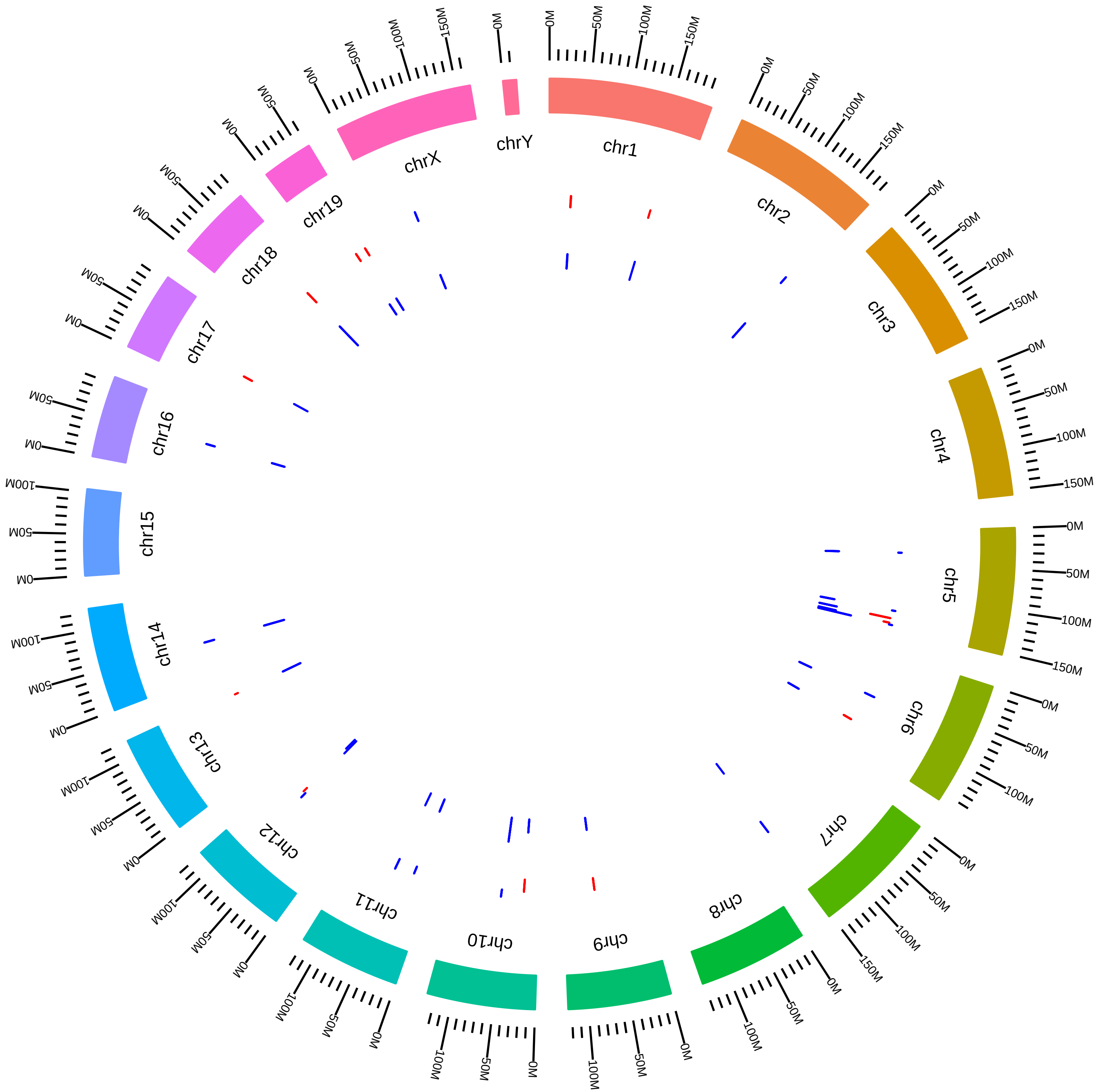

### rsfMalat1MmDLSpectorTumor_T_vs_C.pdf

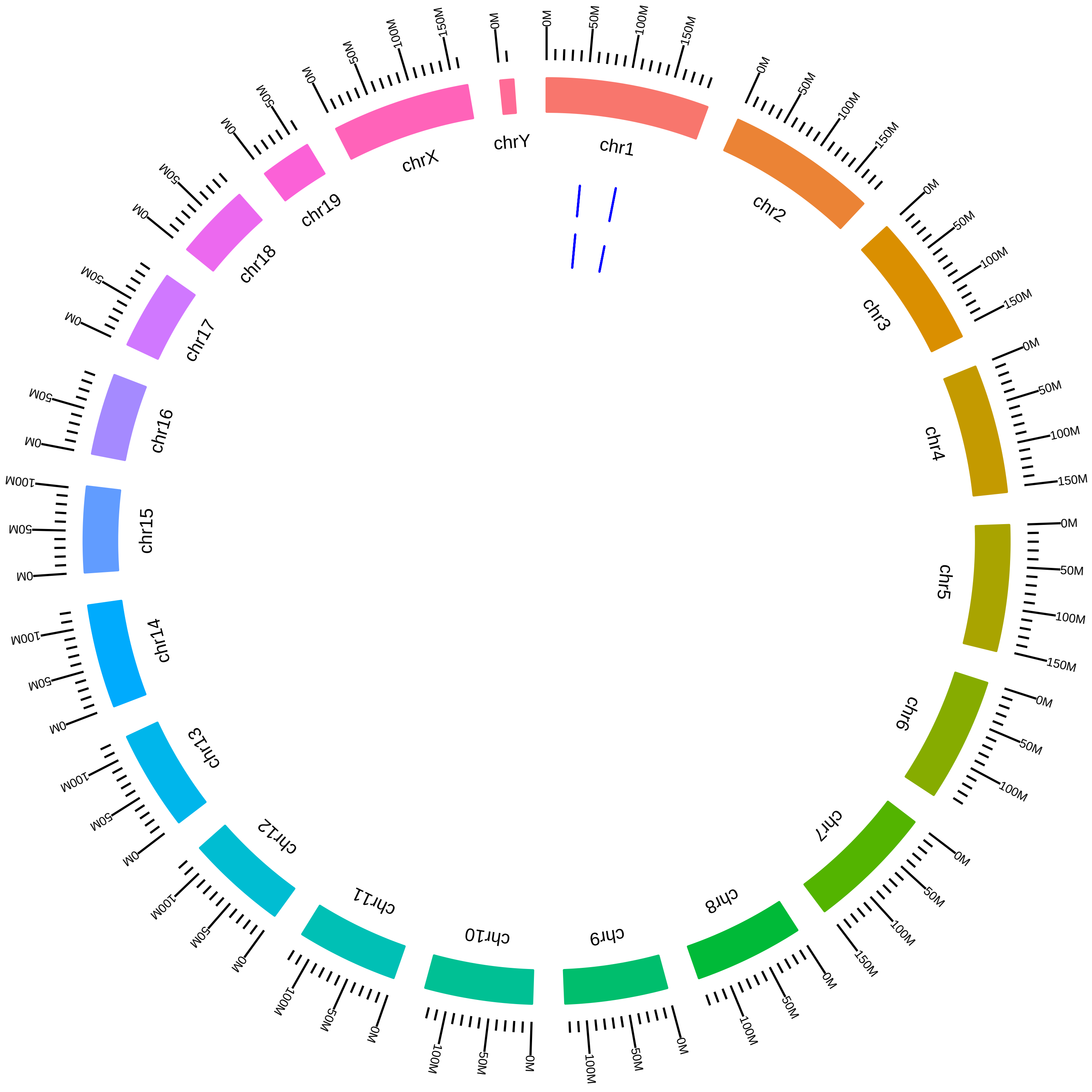

### rsfMbnl1Mbnl2Mbnl3MefMmMSwanson2014_DVsC.pdf

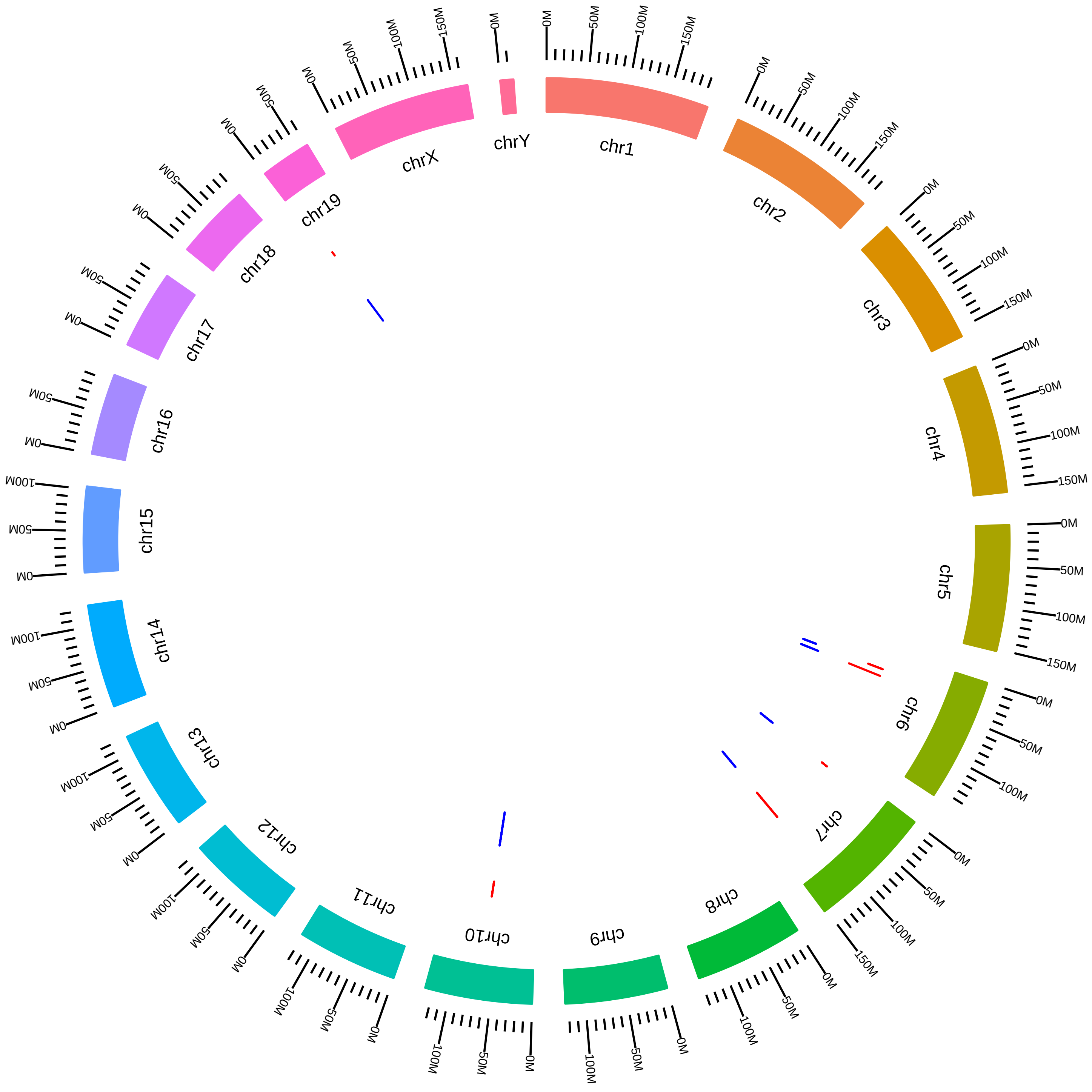

### rsfMbnl1Mbnl2Mbnl3MefMmMSwanson2014_DVsK.pdf

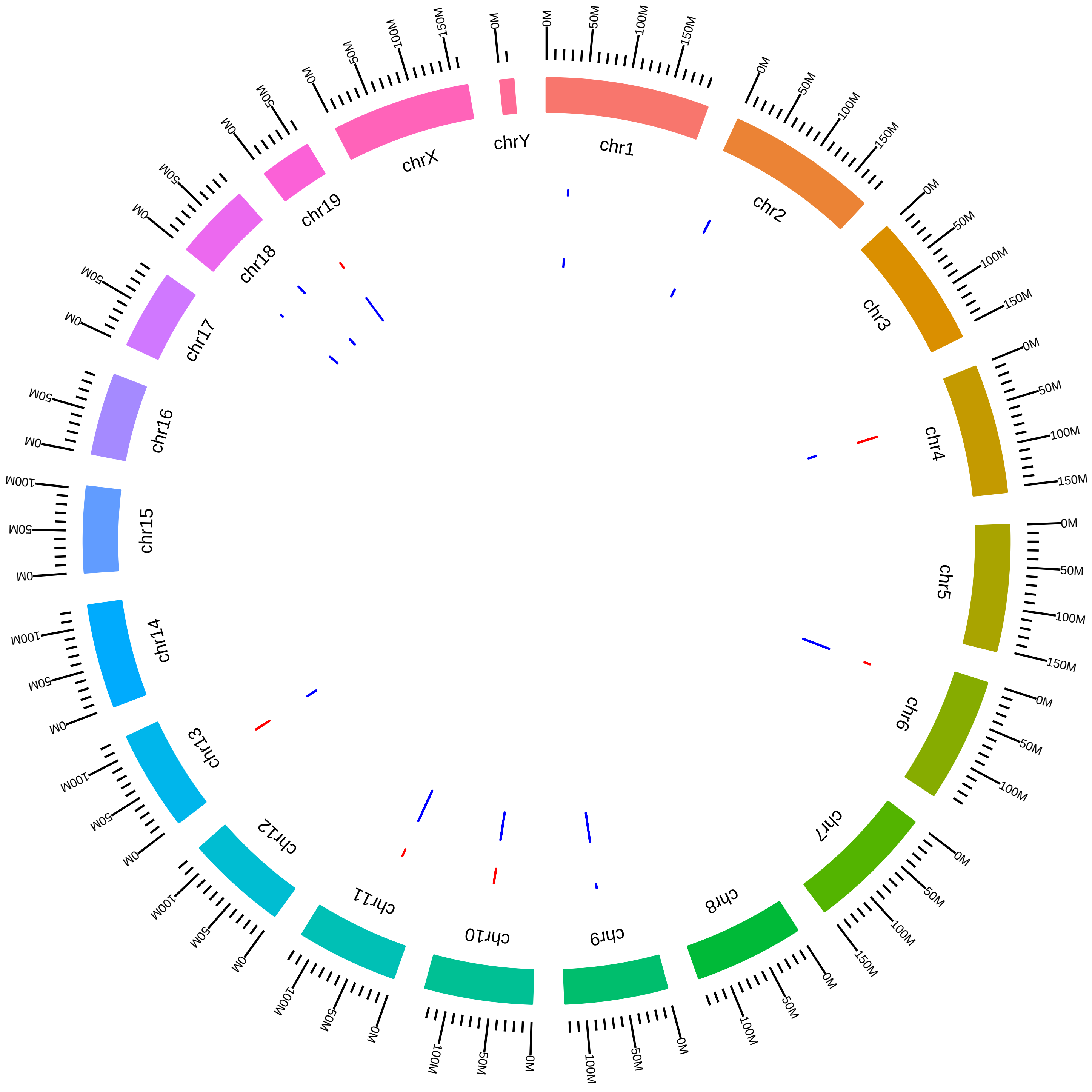

### rsfMbnl1Mbnl2Mbnl3MefMmMSwanson2014_DVsW.pdf

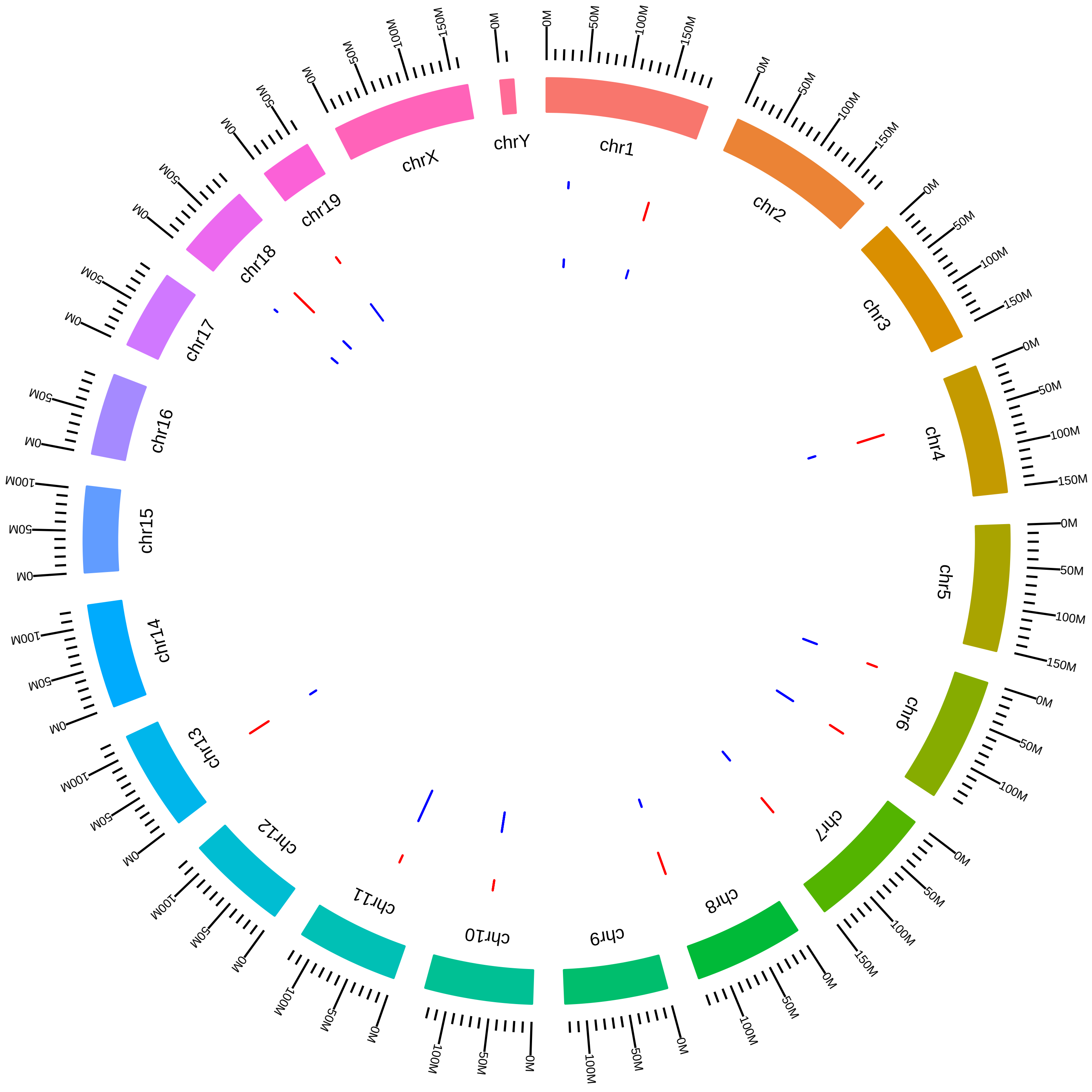

### rsfMbnl1Mbnl2Mbnl3MefMmMSwanson2014_KVsW.pdf

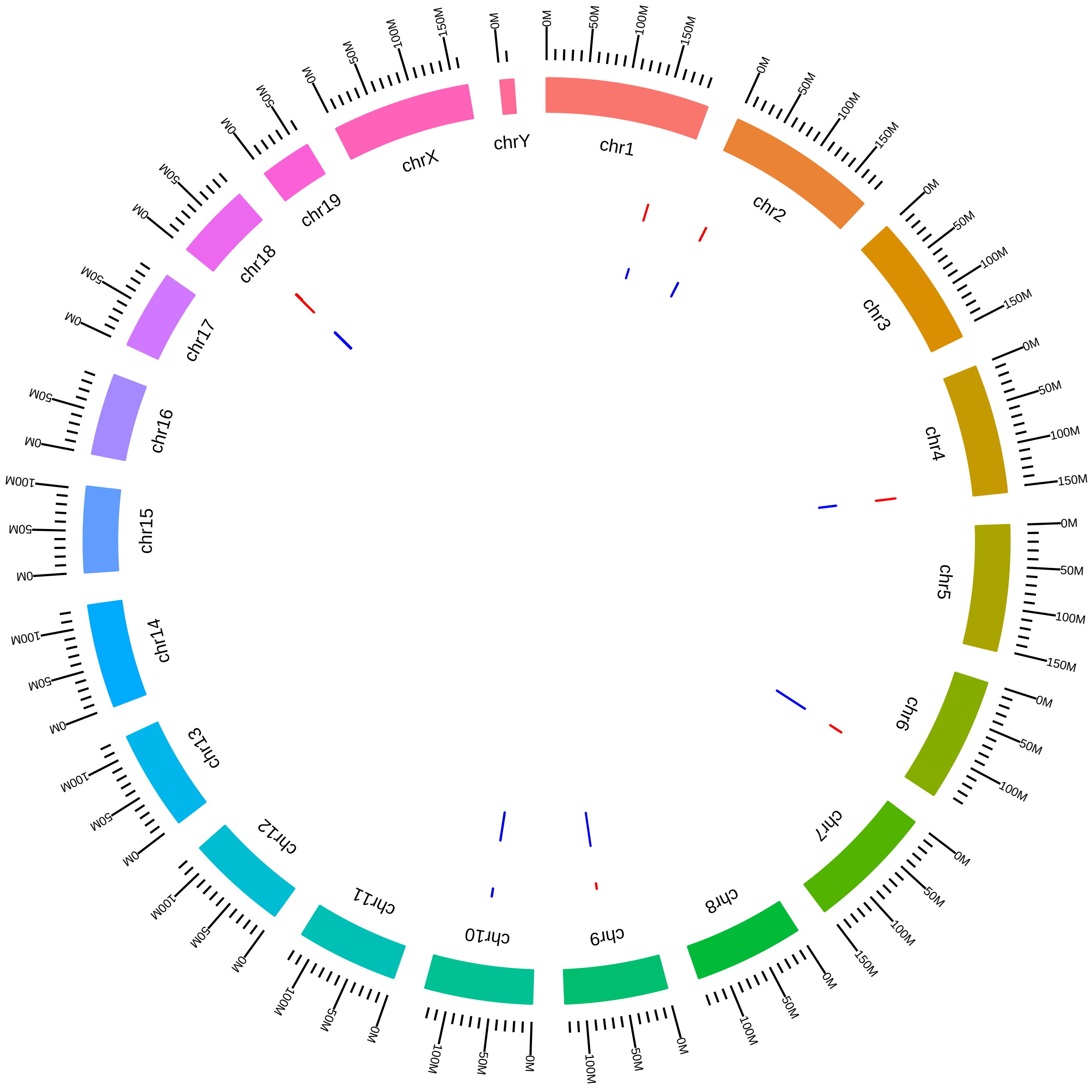

### rsfMbnl1Mbnl2MmCBurgeCADneuriteDKD_DKD_vs_WT.pdf

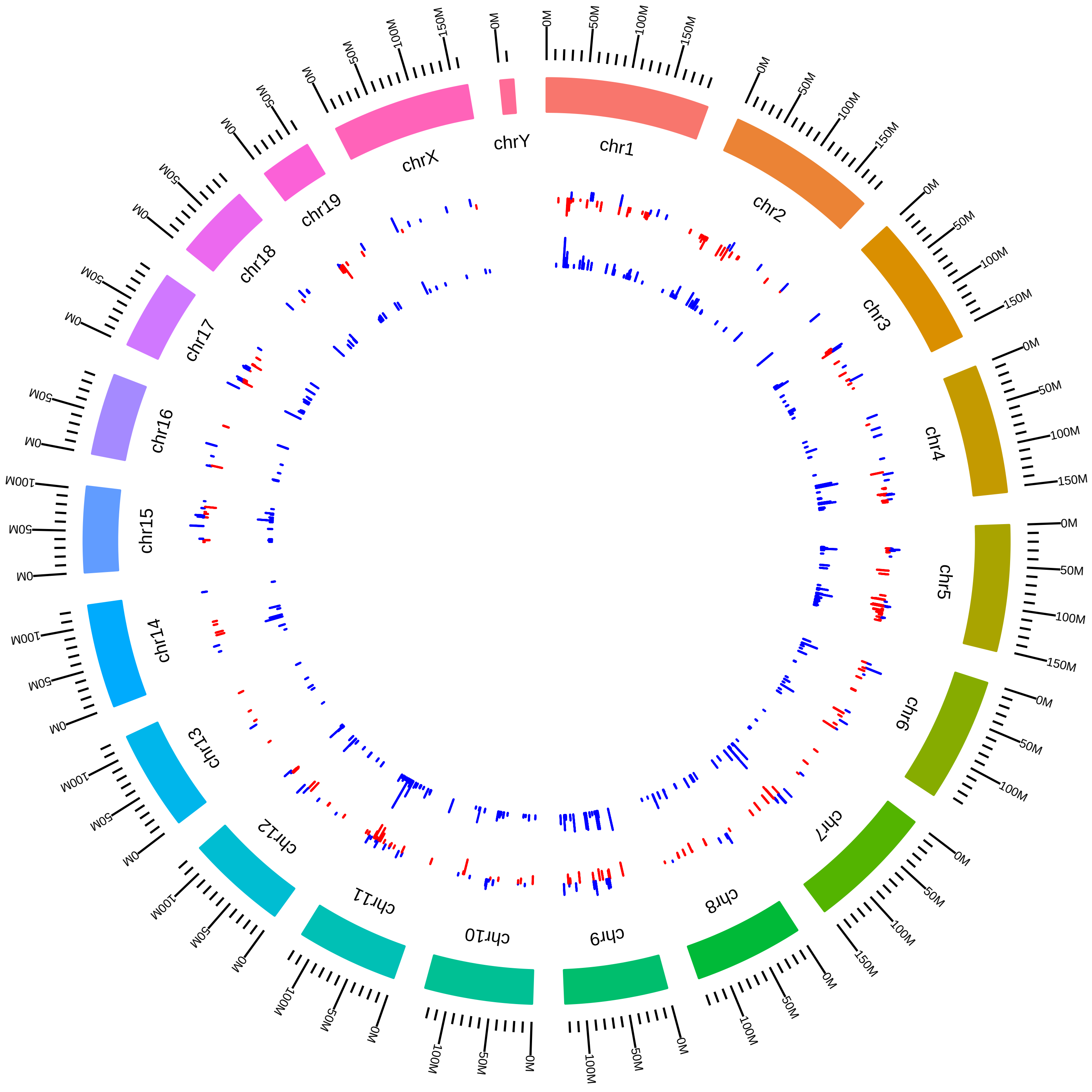

### rsfMbnl1Mbnl2MmCBurgeCADsomaDKD_DKD_vs_WT.pdf

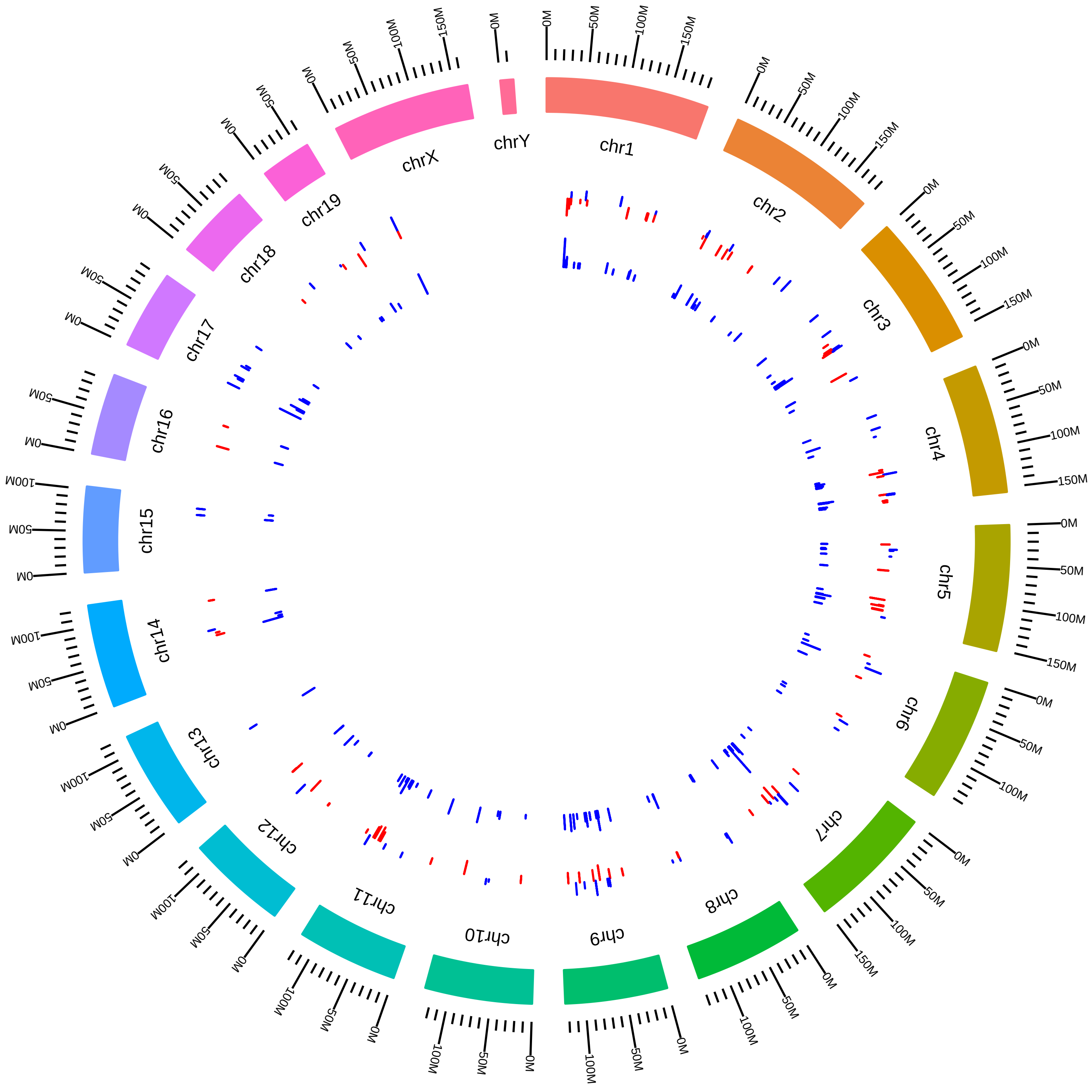

### rsfMbnl1Mbnl2MmCBurgeCortexneuriteDKO_DKO_vs_WT.pdf

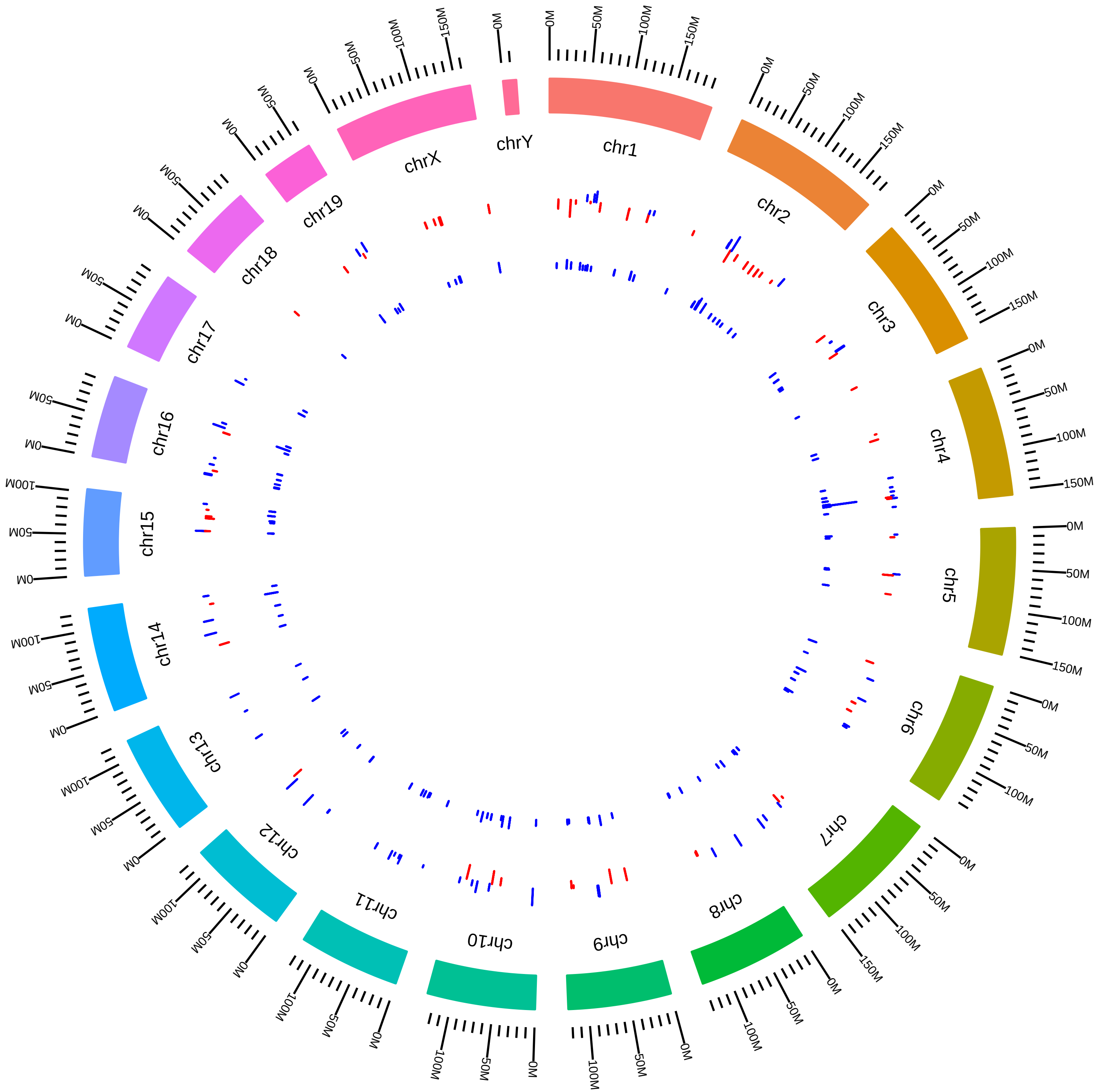

### rsfMbnl1Mbnl2MmCBurgeCortexneuriteKO_Mbnl1KO_vs_WT.pdf

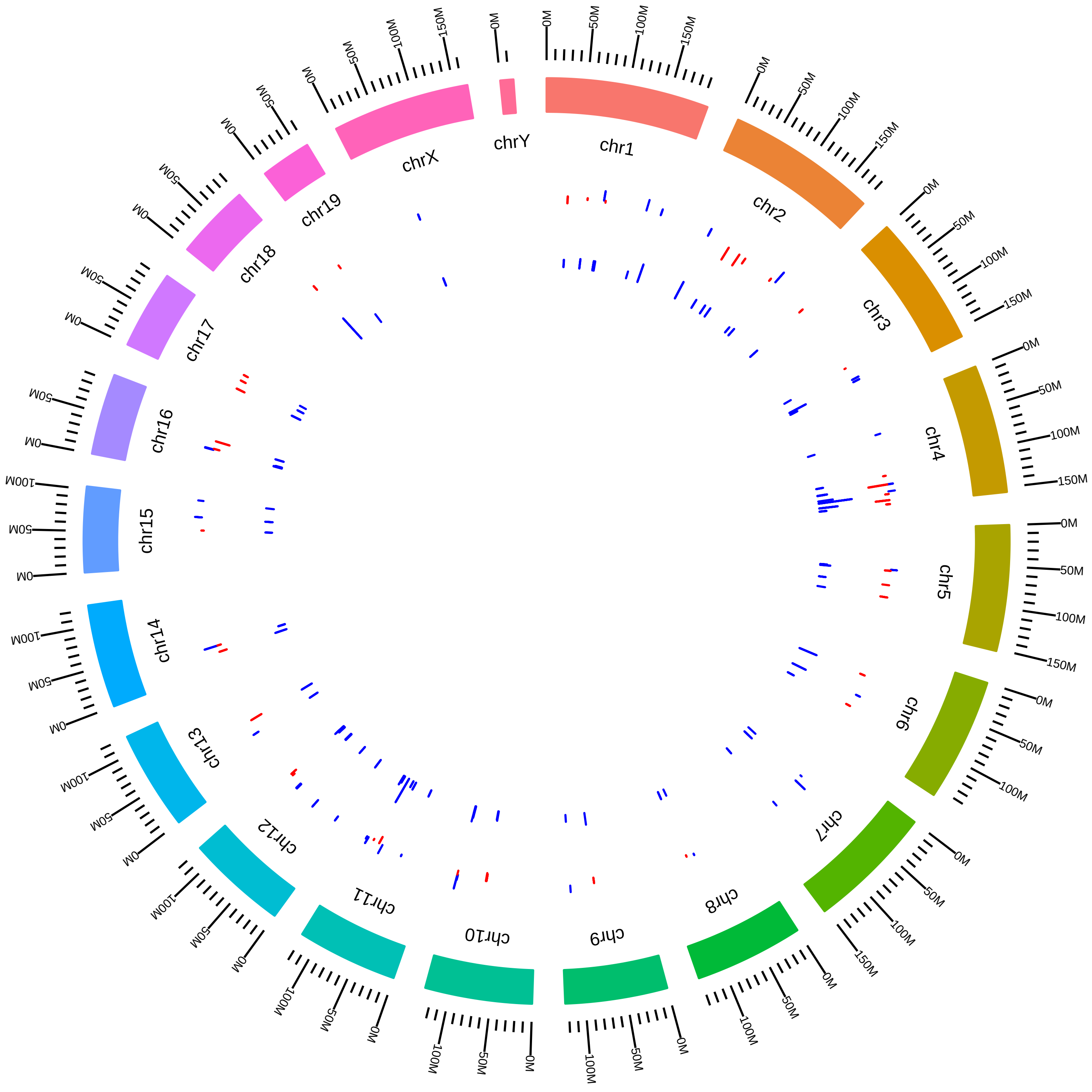

### rsfMbnl1Mbnl2MmCBurgeCortexneuriteKO_Mbnl2KO_vs_WT.pdf

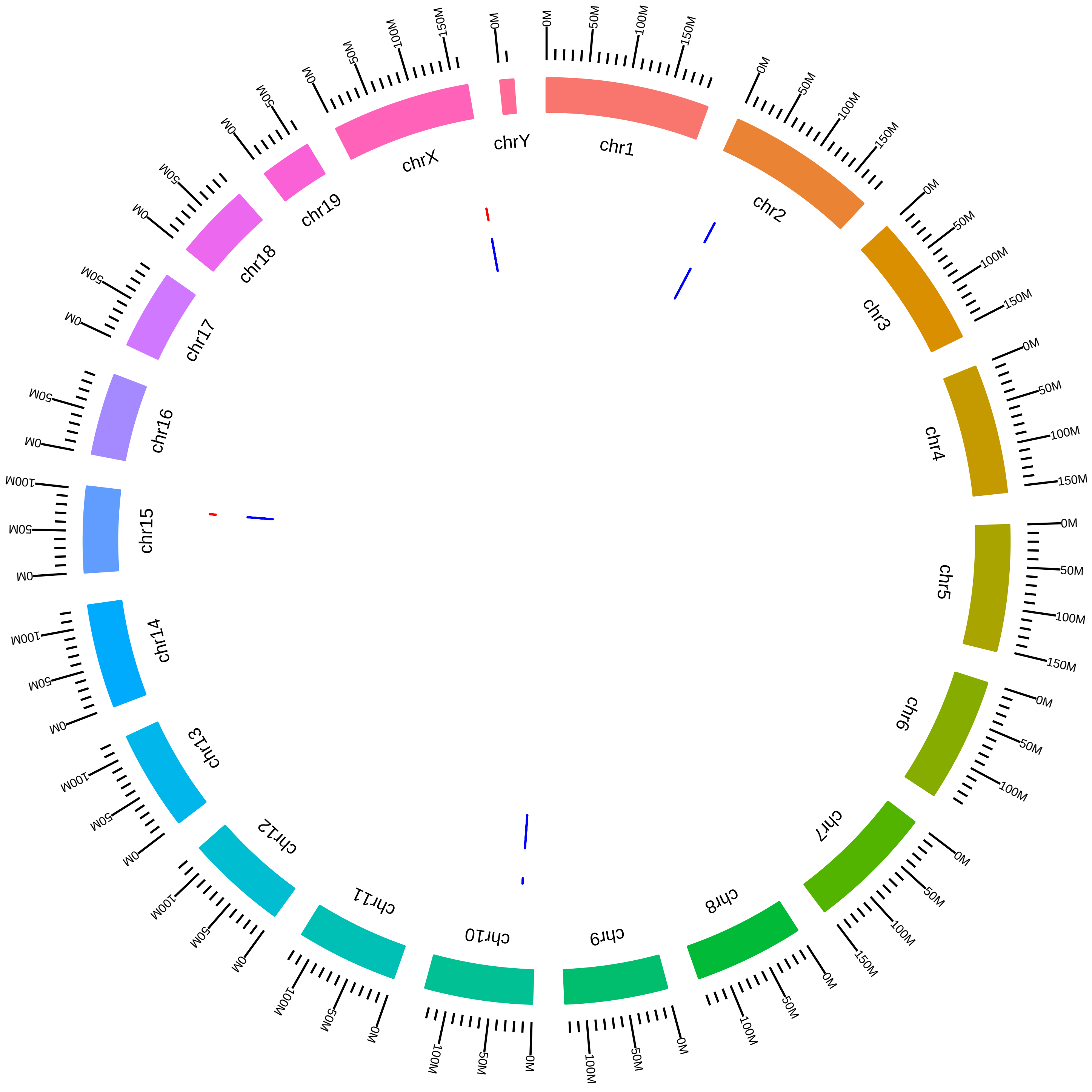

### rsfMbnl1Mbnl2MmCBurgeCortexsomaDKO_DKO_vs_WT.pdf

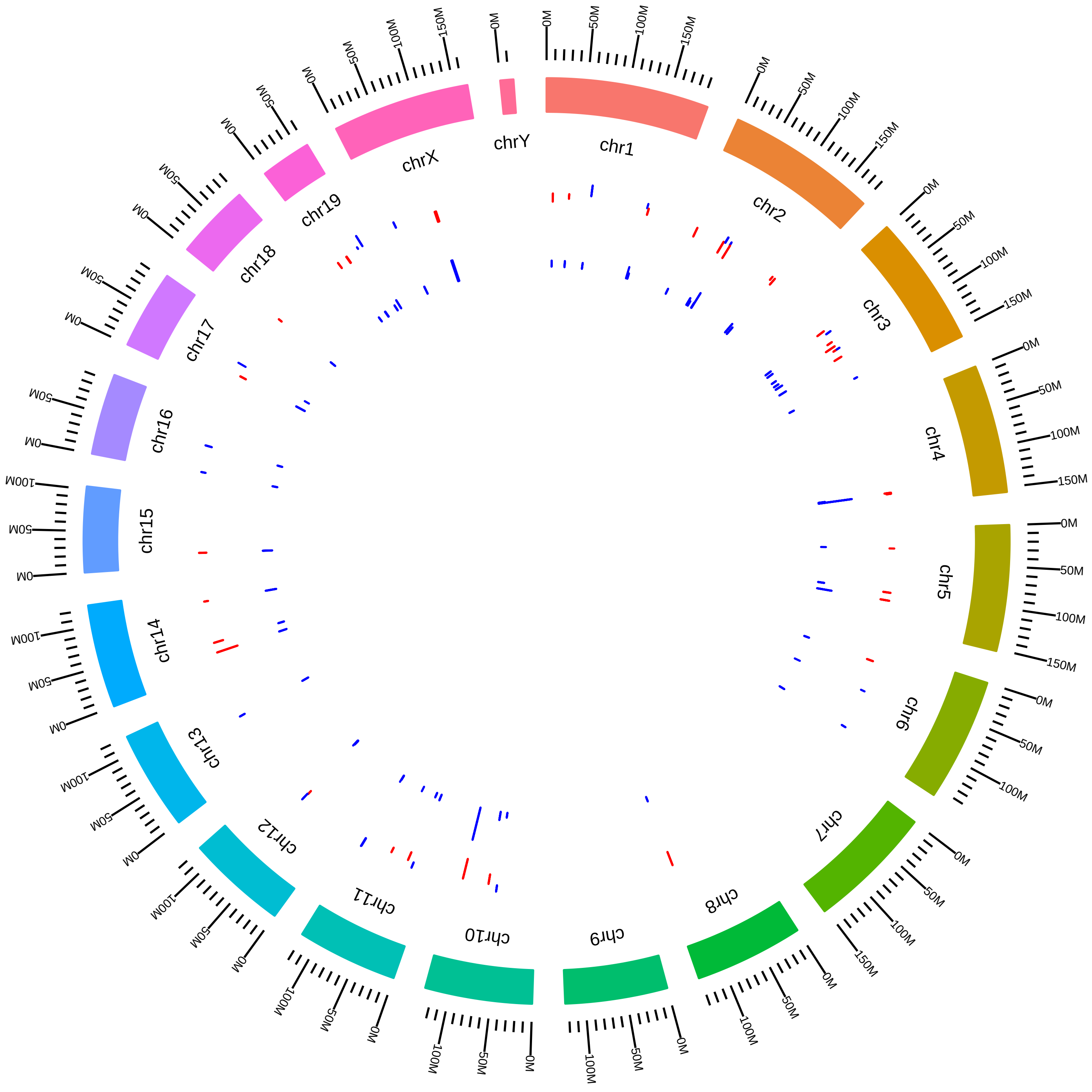

### rsfMbnl1Mbnl2MmCBurgeCortexsomaKO_Mbnl1KO_vs_WT.pdf

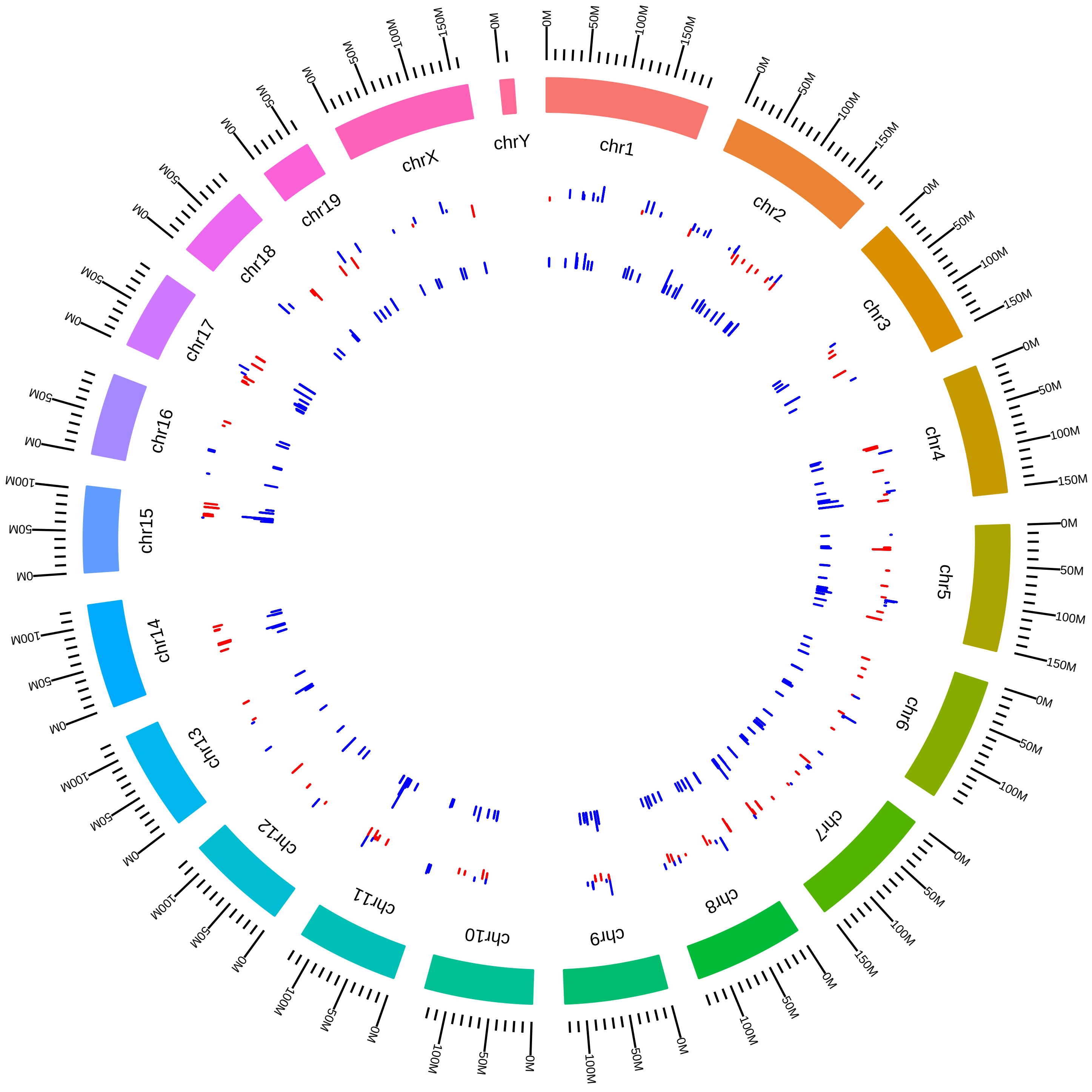

### rsfMbnl1Mbnl2MmCBurgeCortexsomaKO_Mbnl2KO_vs_WT.pdf

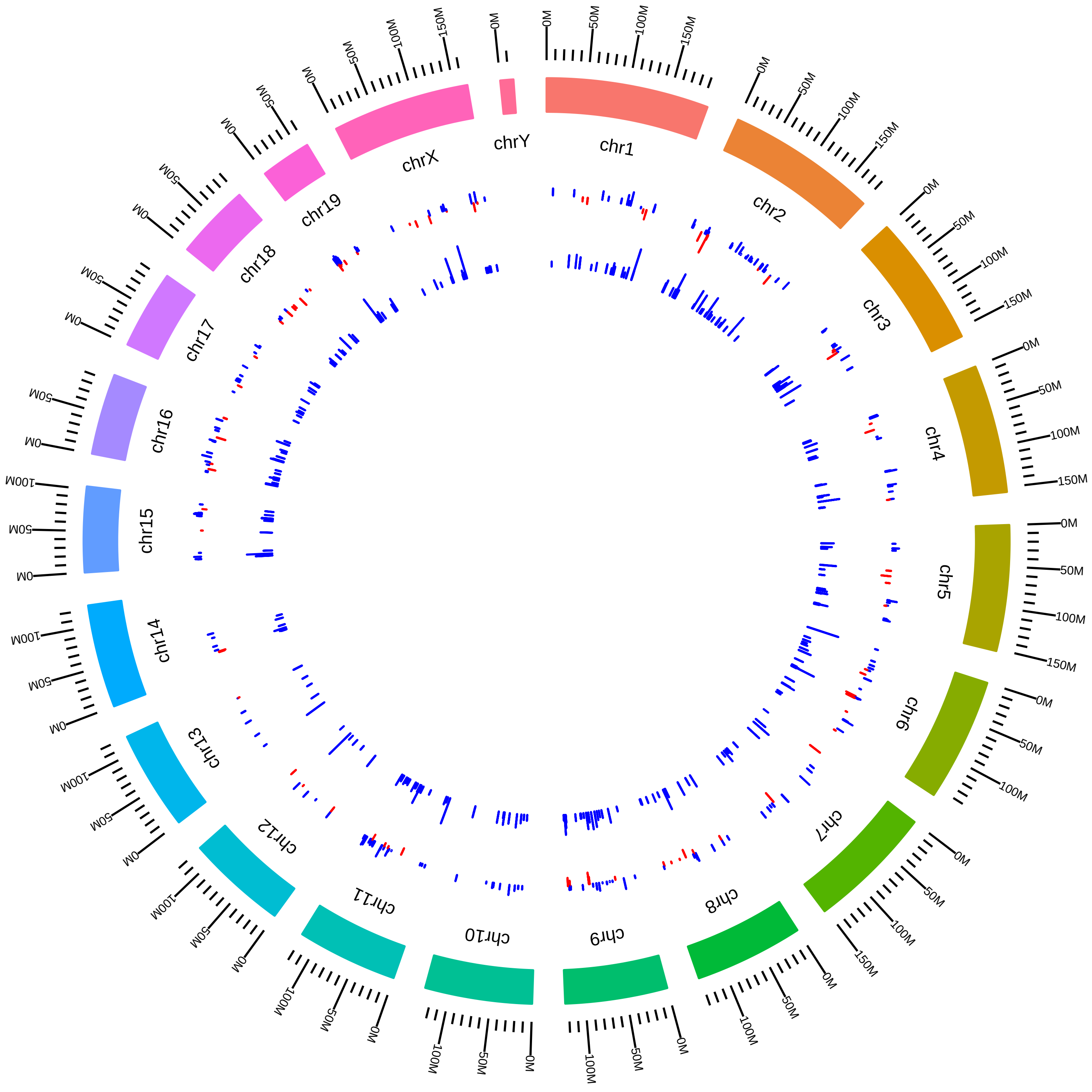

### rsfMbnl1Mbnl2MmEWang_DKO_vs_WT.pdf

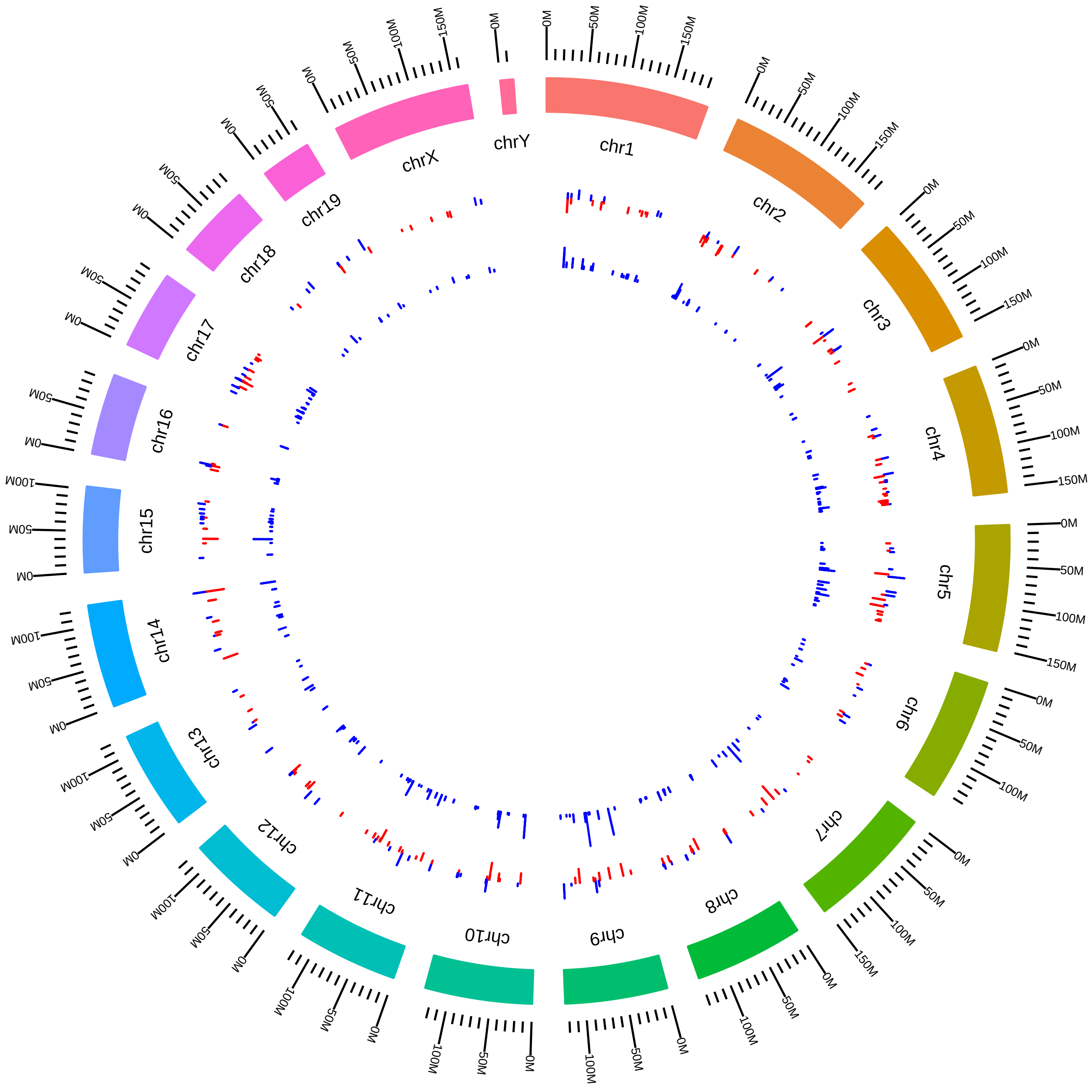

### rsfMbnl1Mbnl2MmMSwanson2015_KVsW.pdf

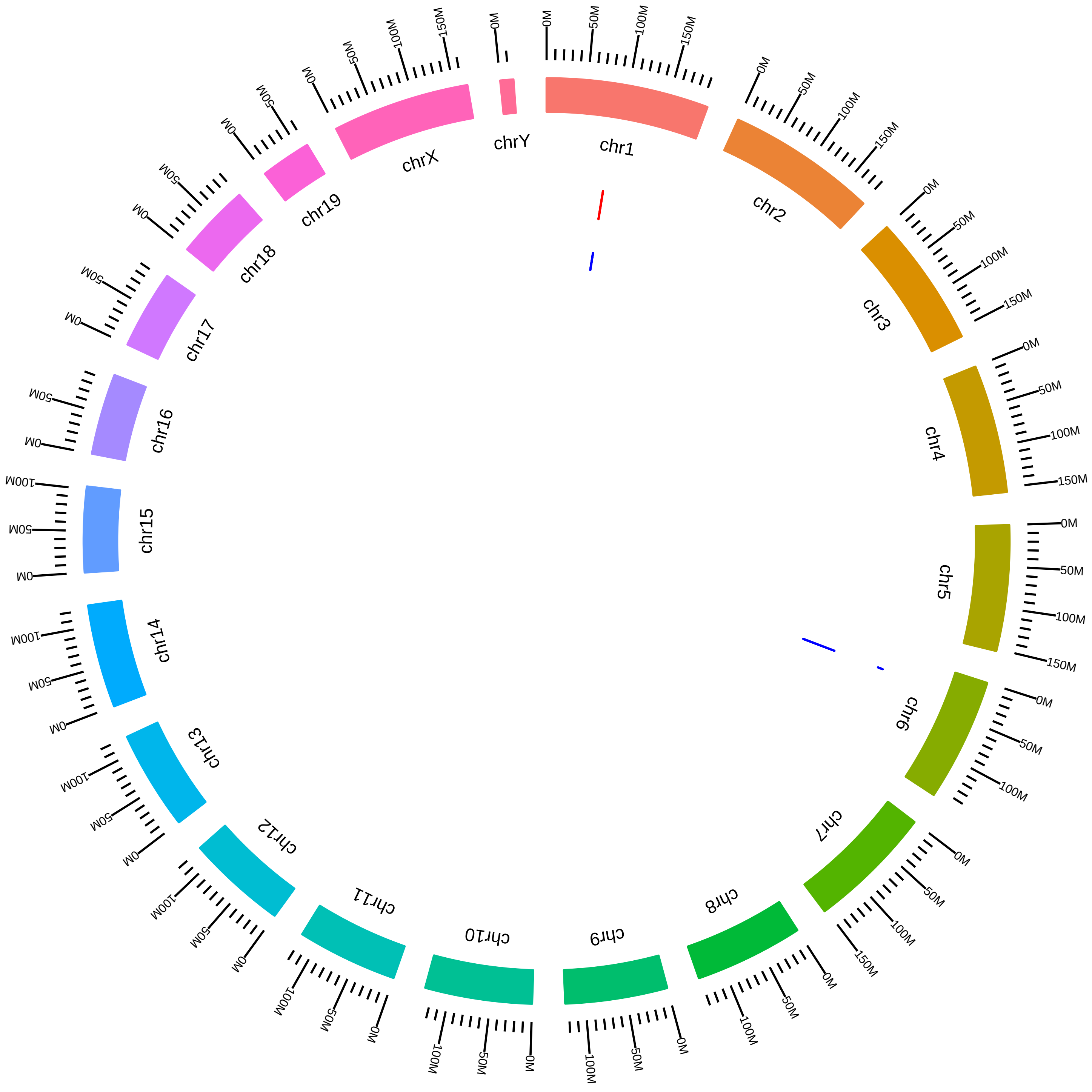

### rsfMbnl1MmCBurgeHeart_KO_vs_WT.pdf

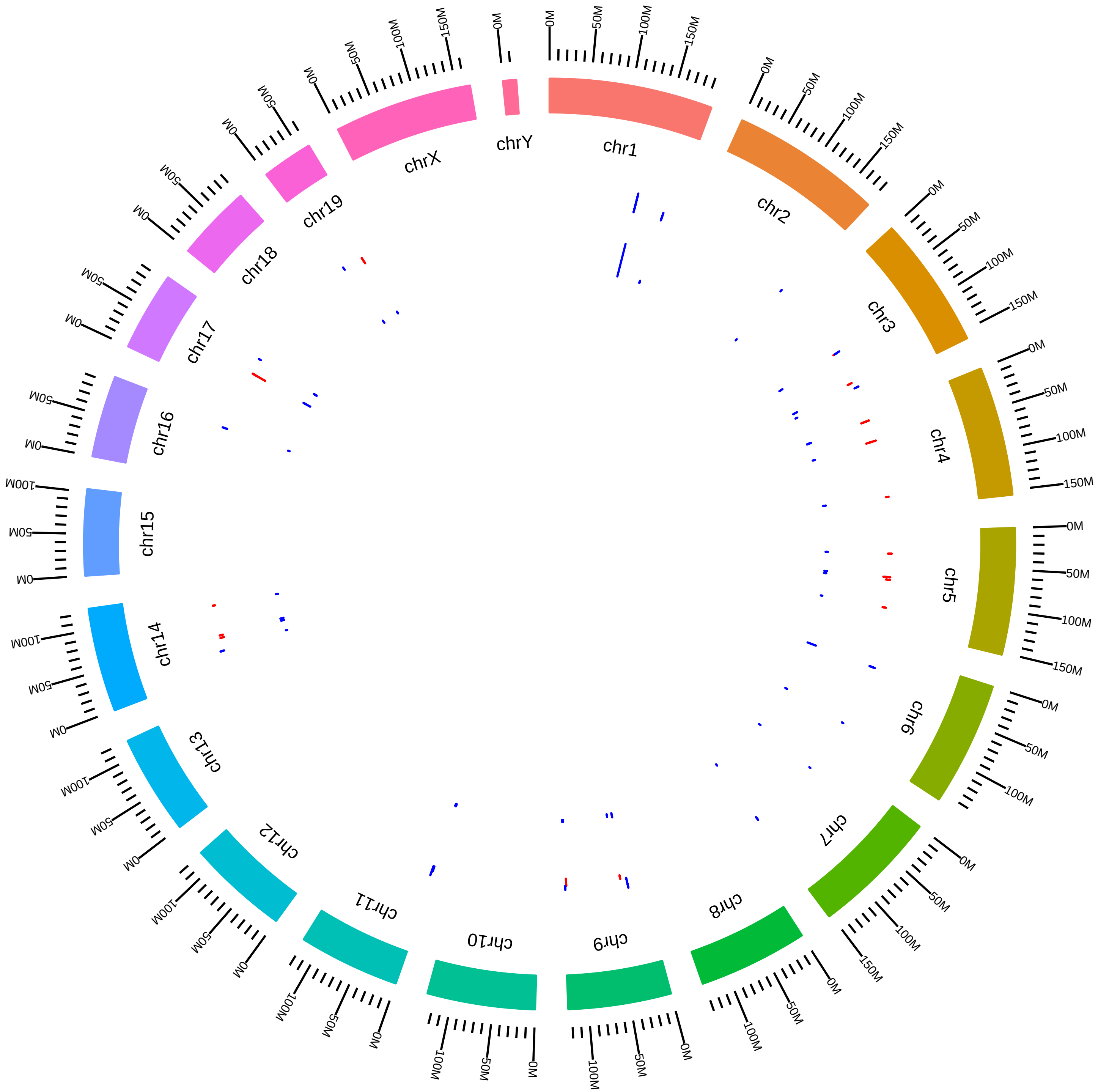

### rsfMbnl1MmCBurgeMuscle_KO_vs_WT.pdf

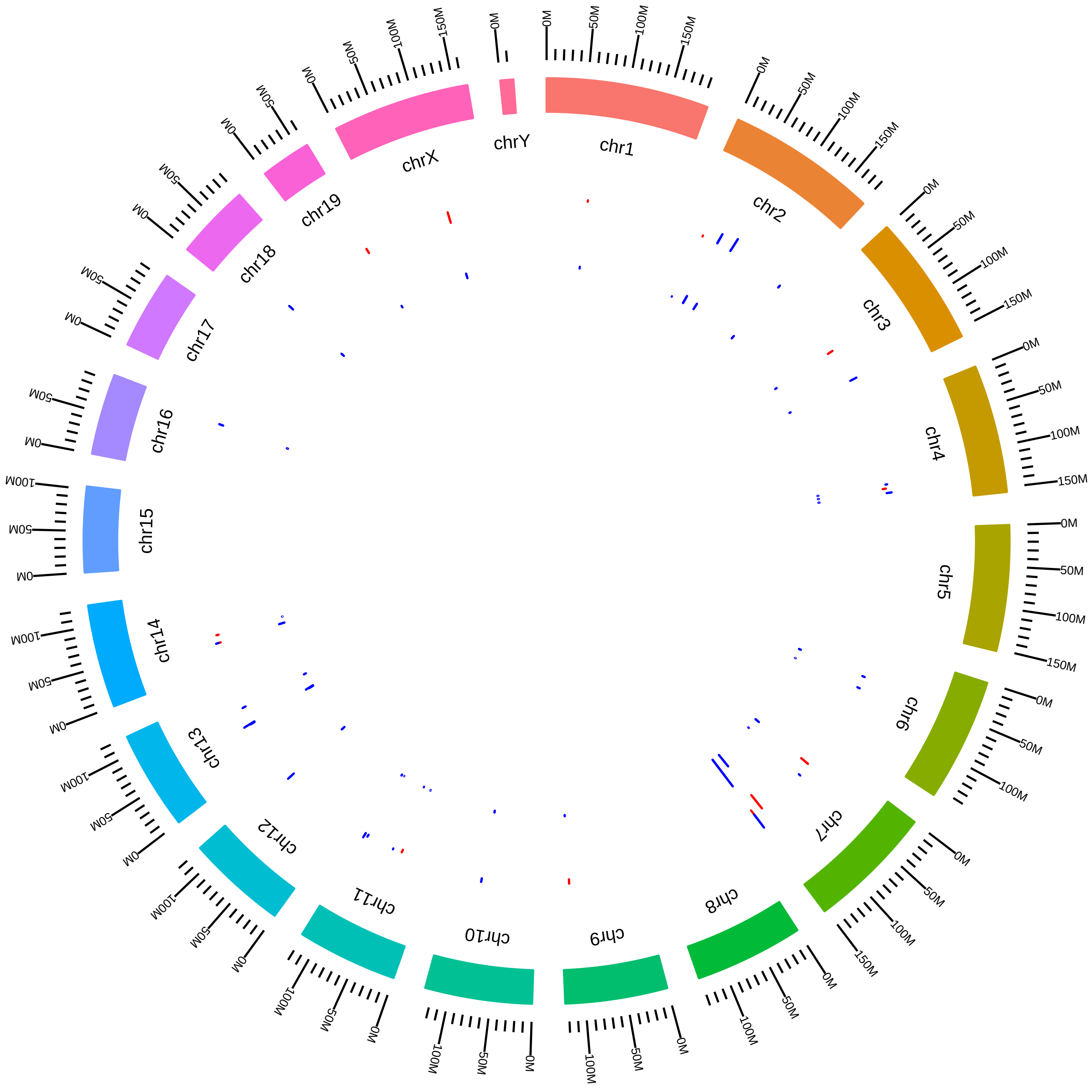

### rsfMbnl2MmMSwanson_KO_vs_WT.pdf

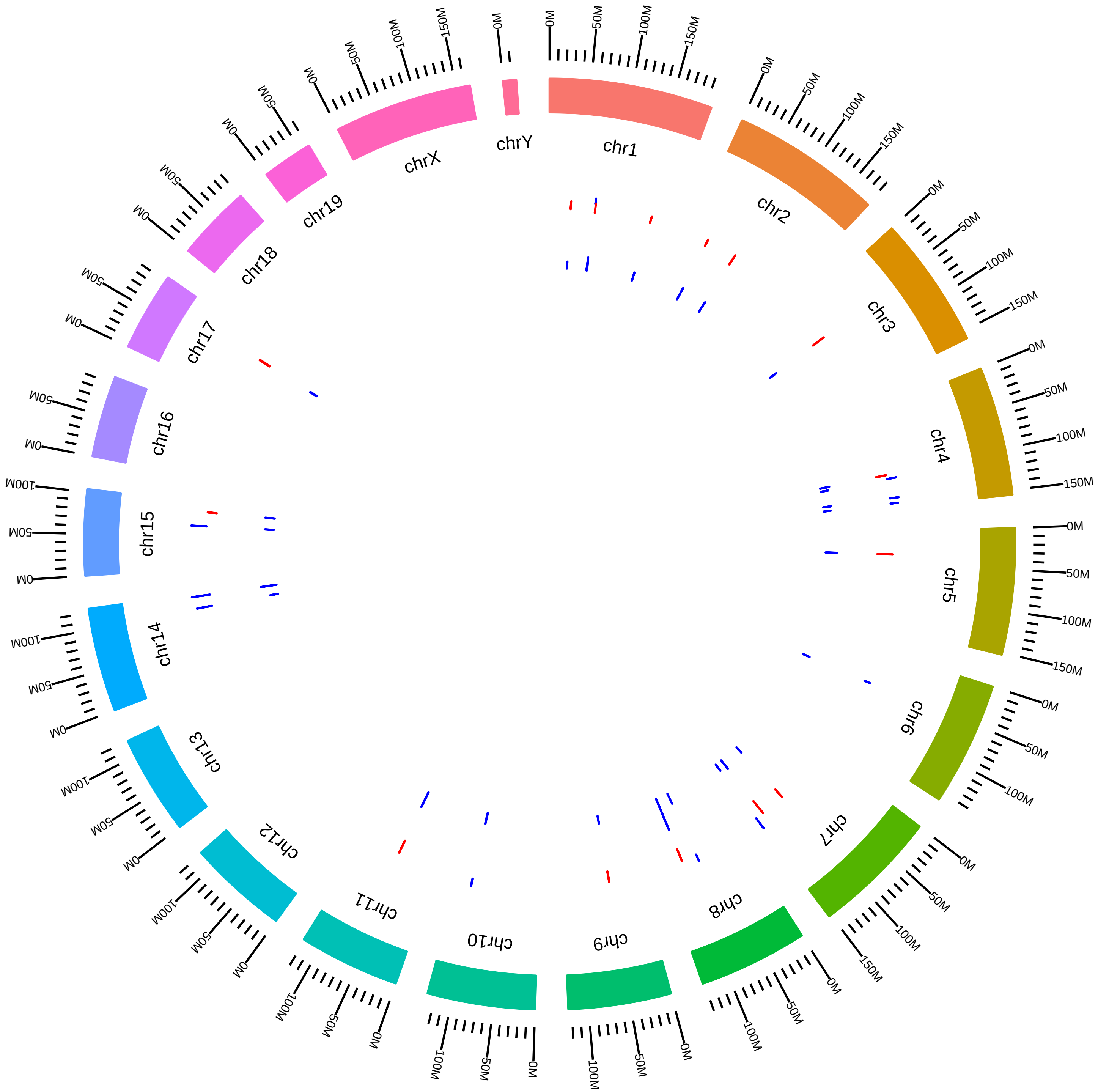

### rsfMettl3MmJHanna2015_BKVsBW.pdf

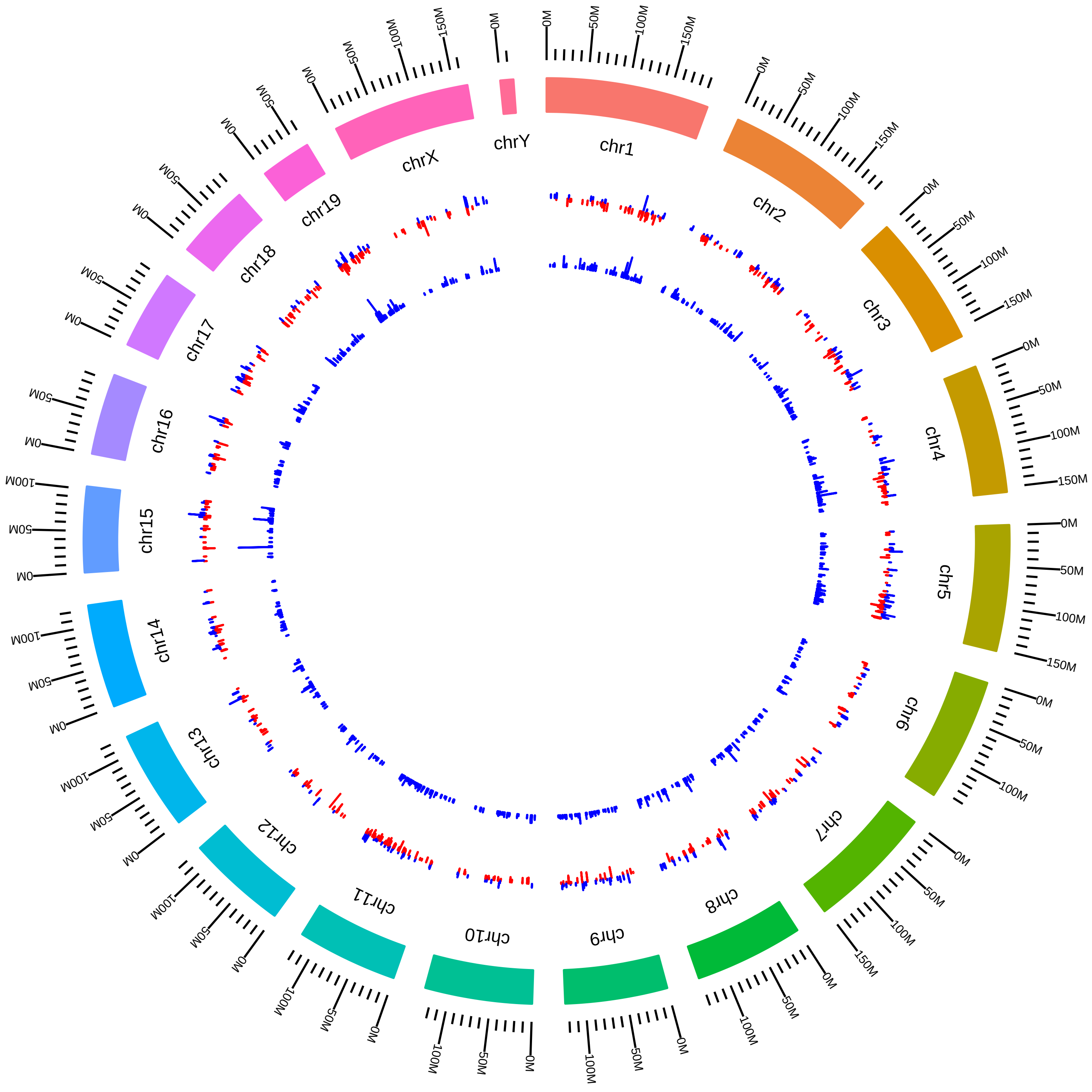

### rsfMettl3MmJHanna2015_SKVsSW.pdf

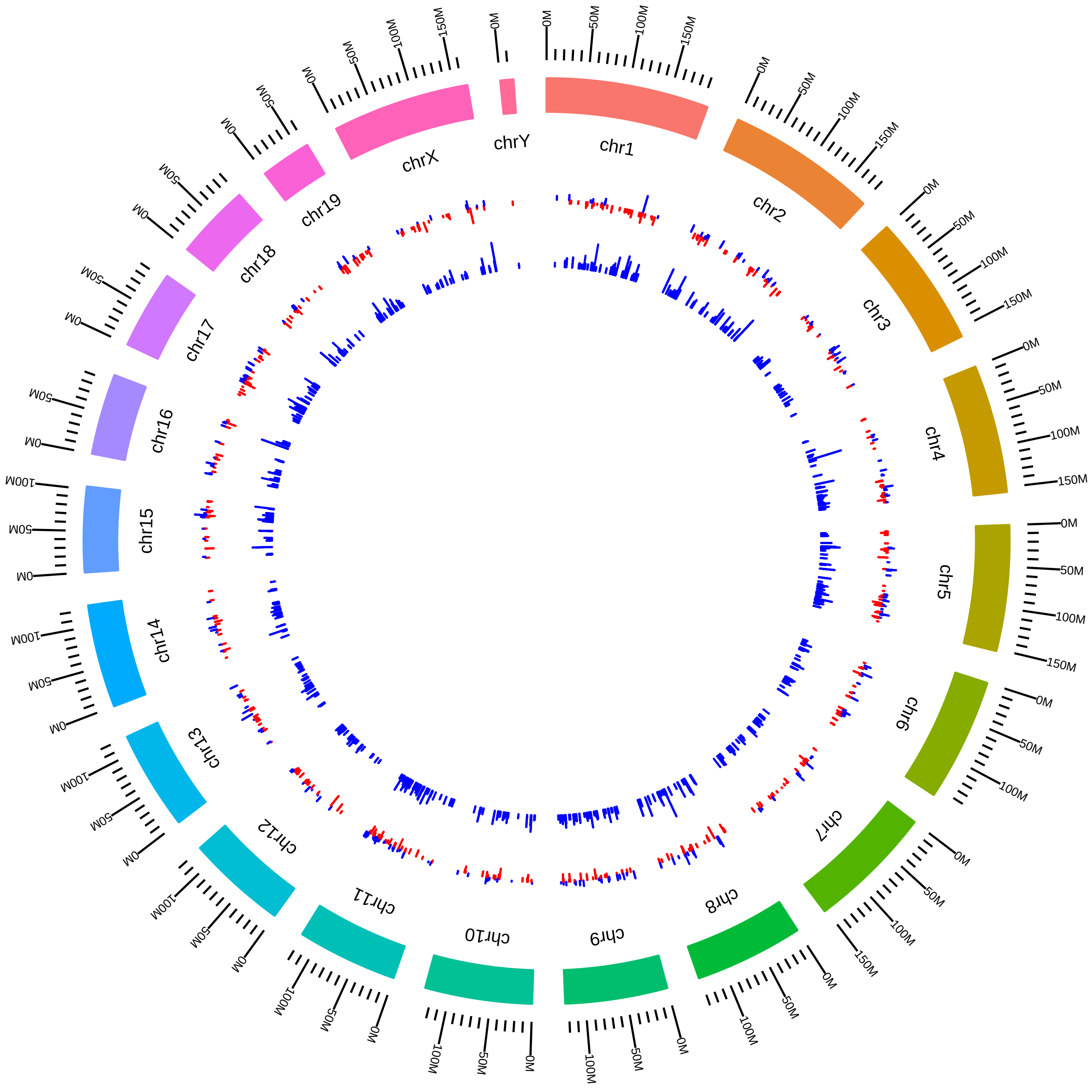

### rsfMettl3MmYunguiYang2014_DVsC.pdf

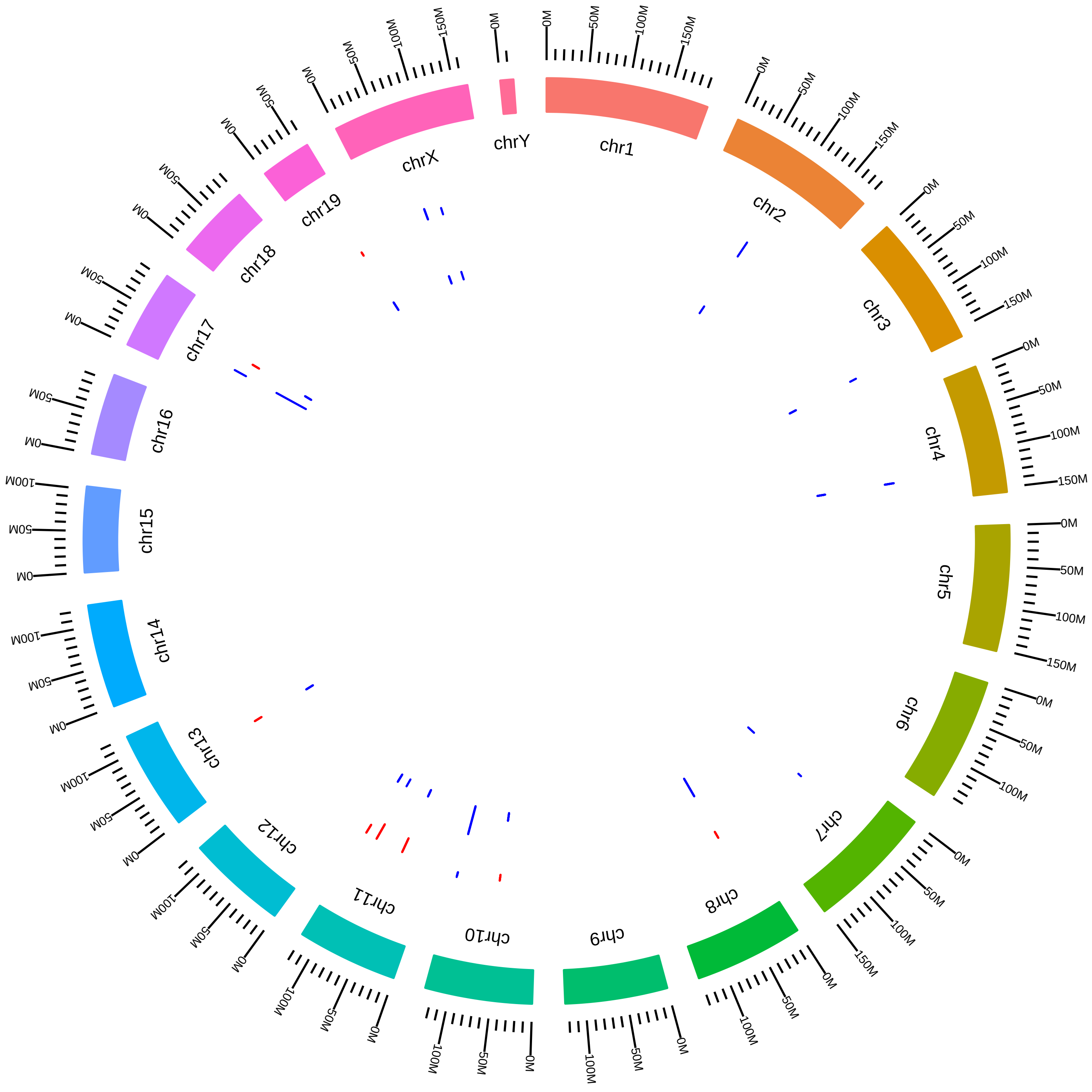

### rsfNonoMm2015SBrown2015_KVsW.pdf

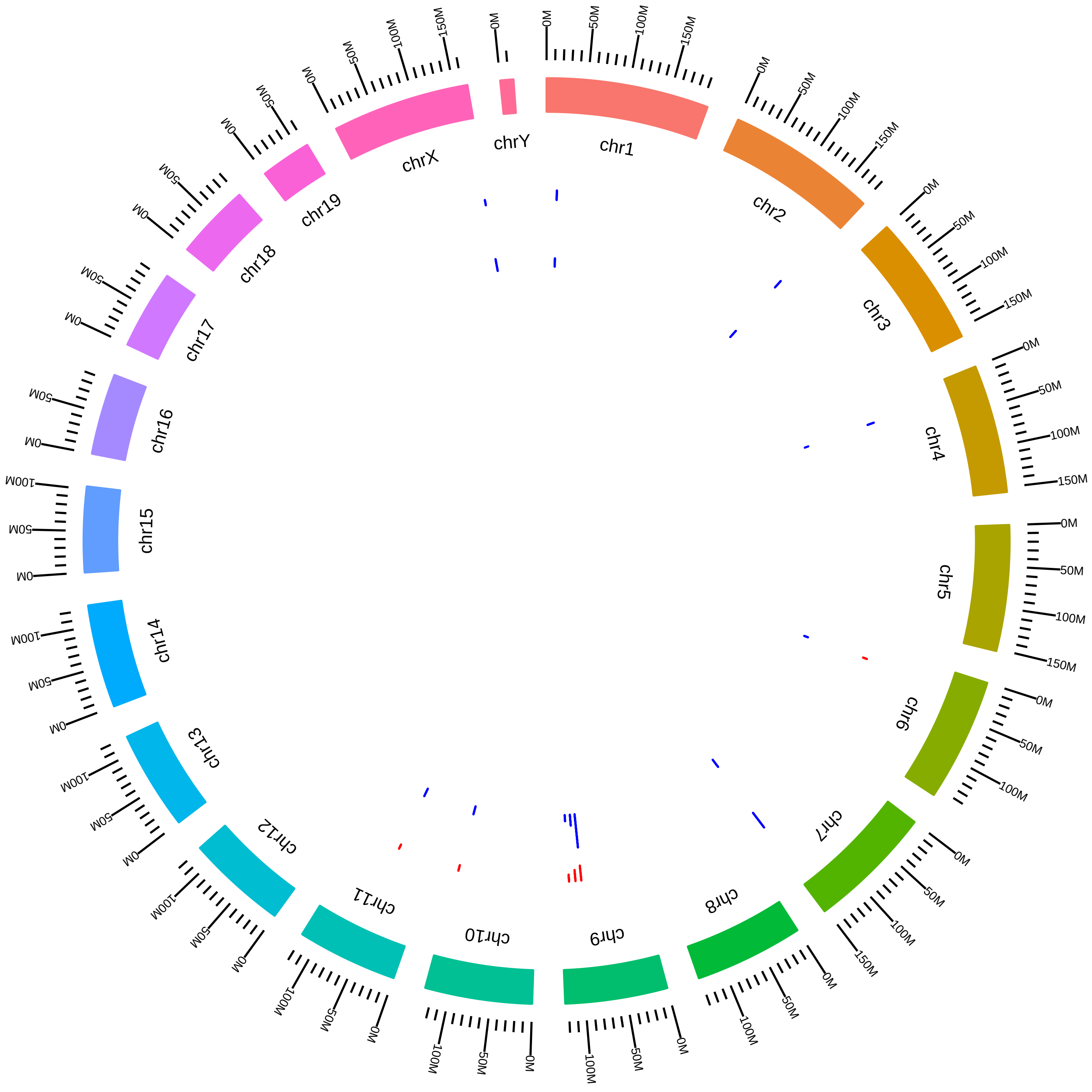

### rsfNova1Nova2MmRDarnellBrainNova1_KO_vs_WT.pdf

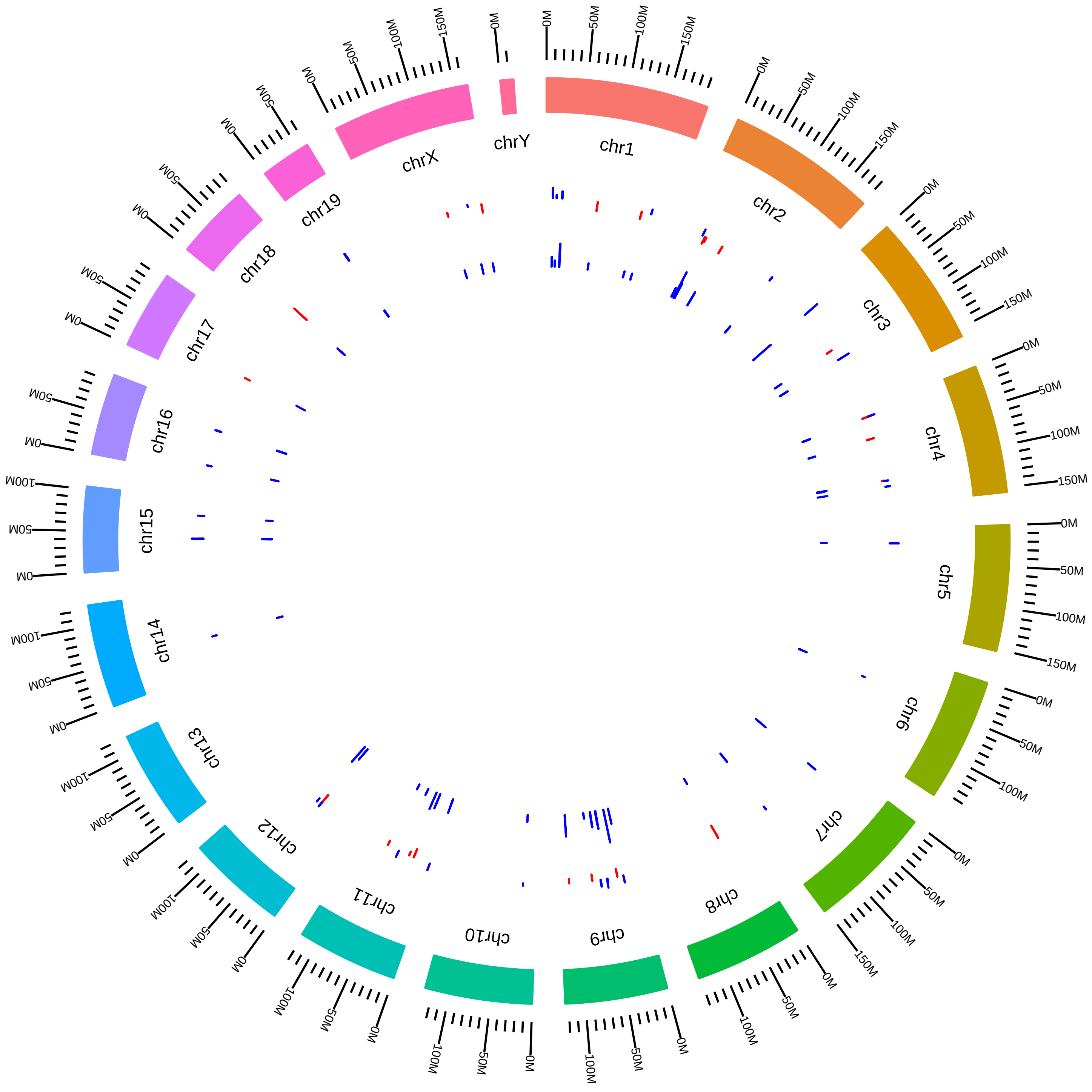

### rsfNova1Nova2MmRDarnellCortexNova1_KO_vs_WT.pdf

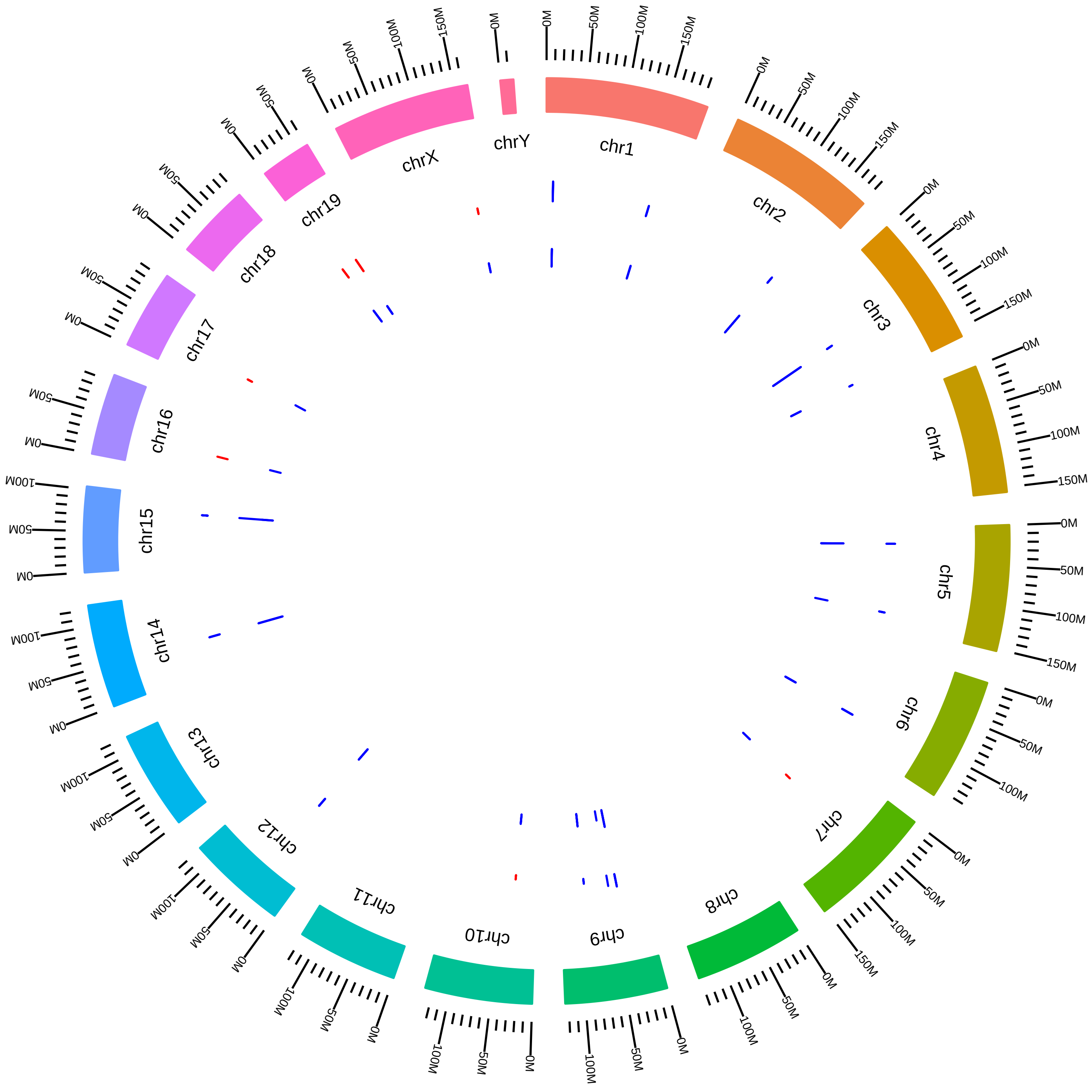

### rsfNova1Nova2MmRDarnellCortexNova2_KO_vs_WT.pdf

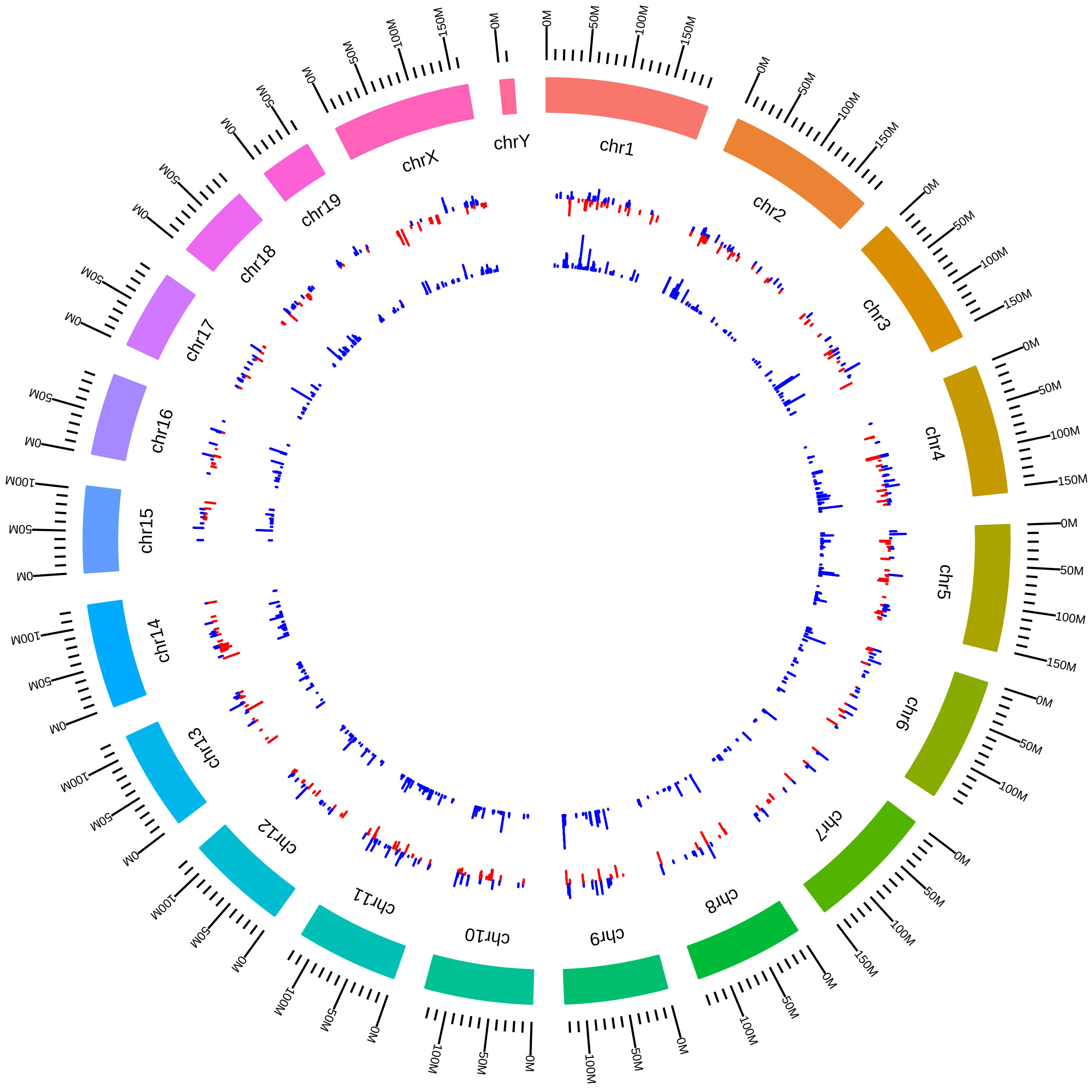

### rsfNova1Nova2MmRDavis_Nova1KO_vs_WT.pdf

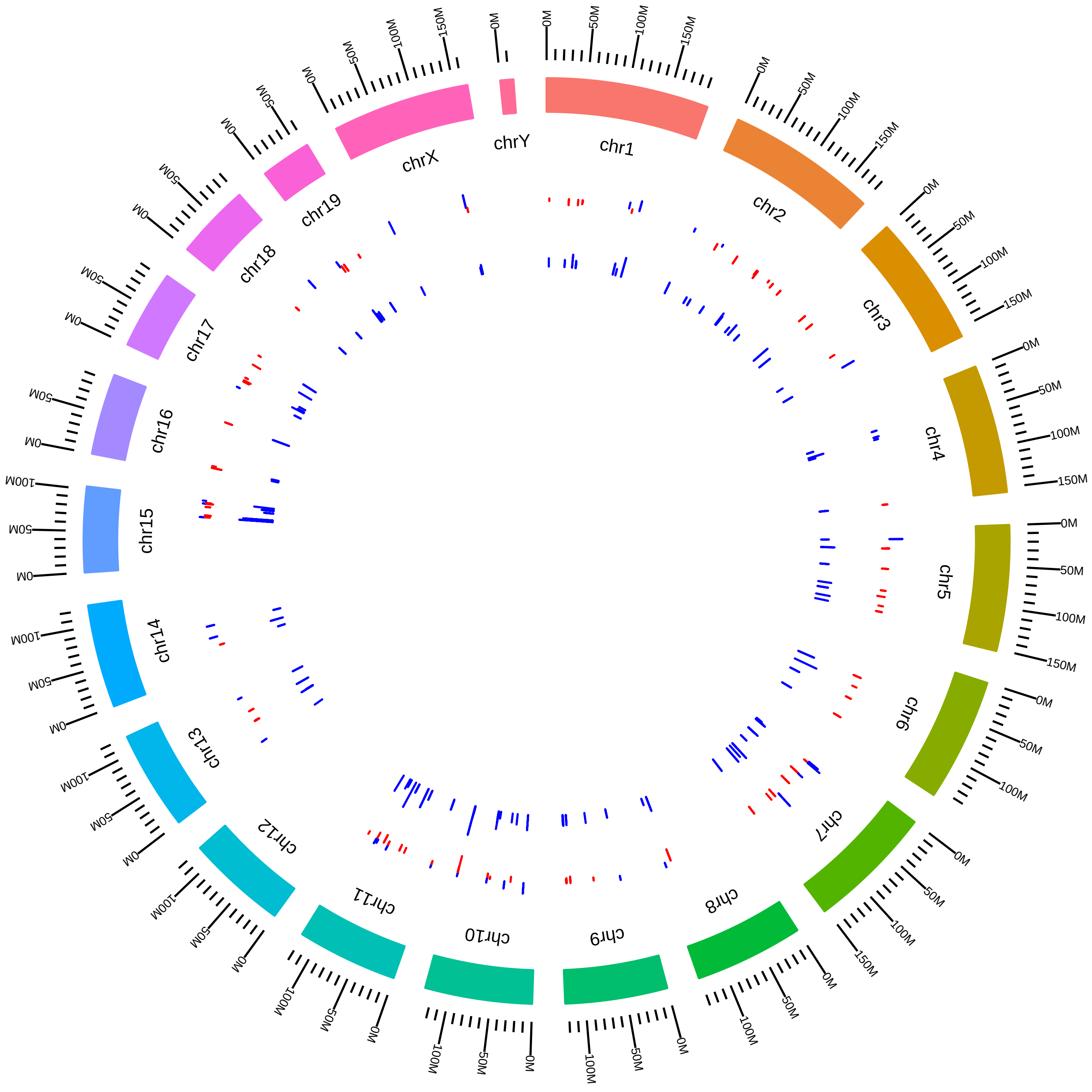

### rsfNova1Nova2MmRDavis_Nova2KO_vs_WT.pdf

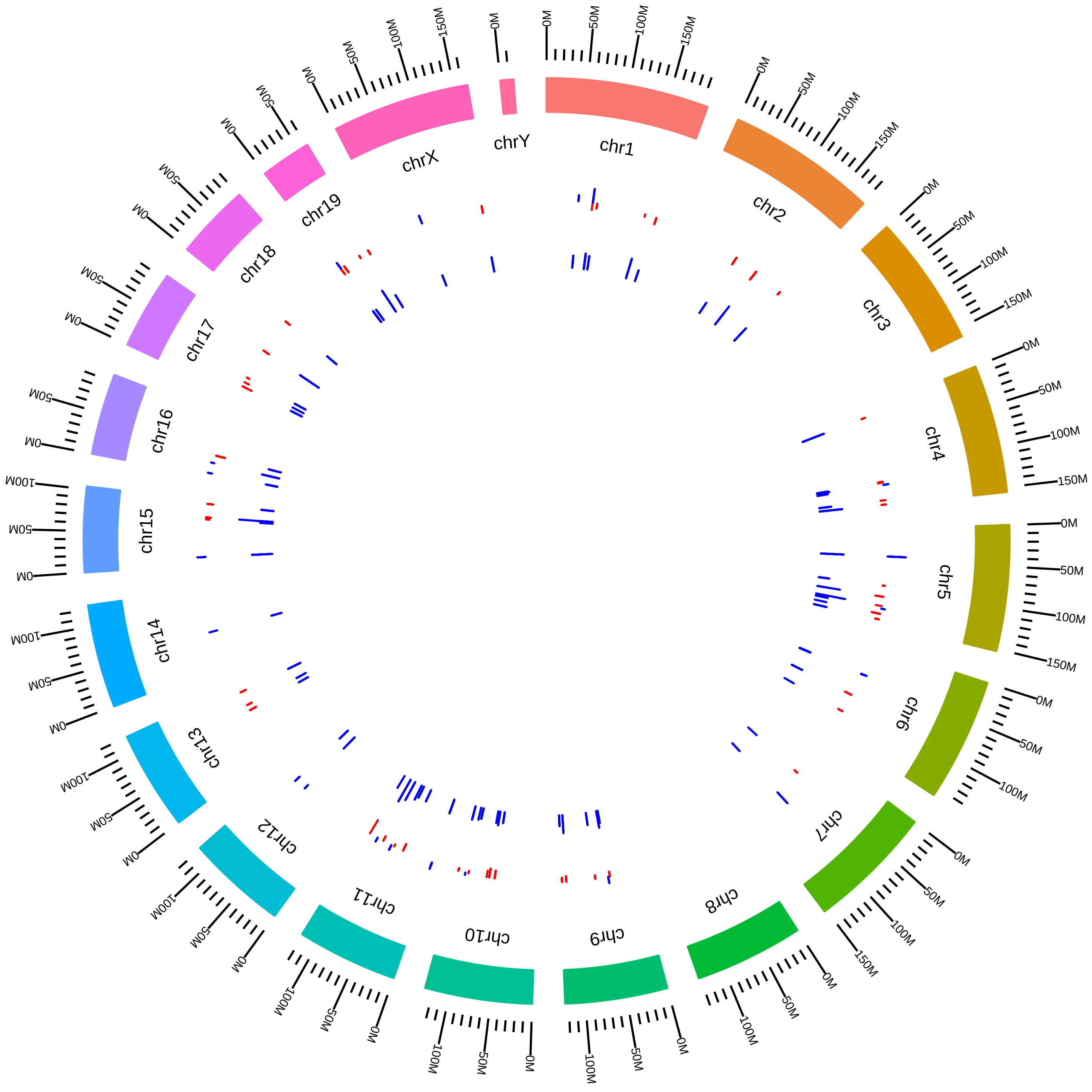

### rsfNova1Nova2MmRDavisIII_DKO_vs_WT.pdf

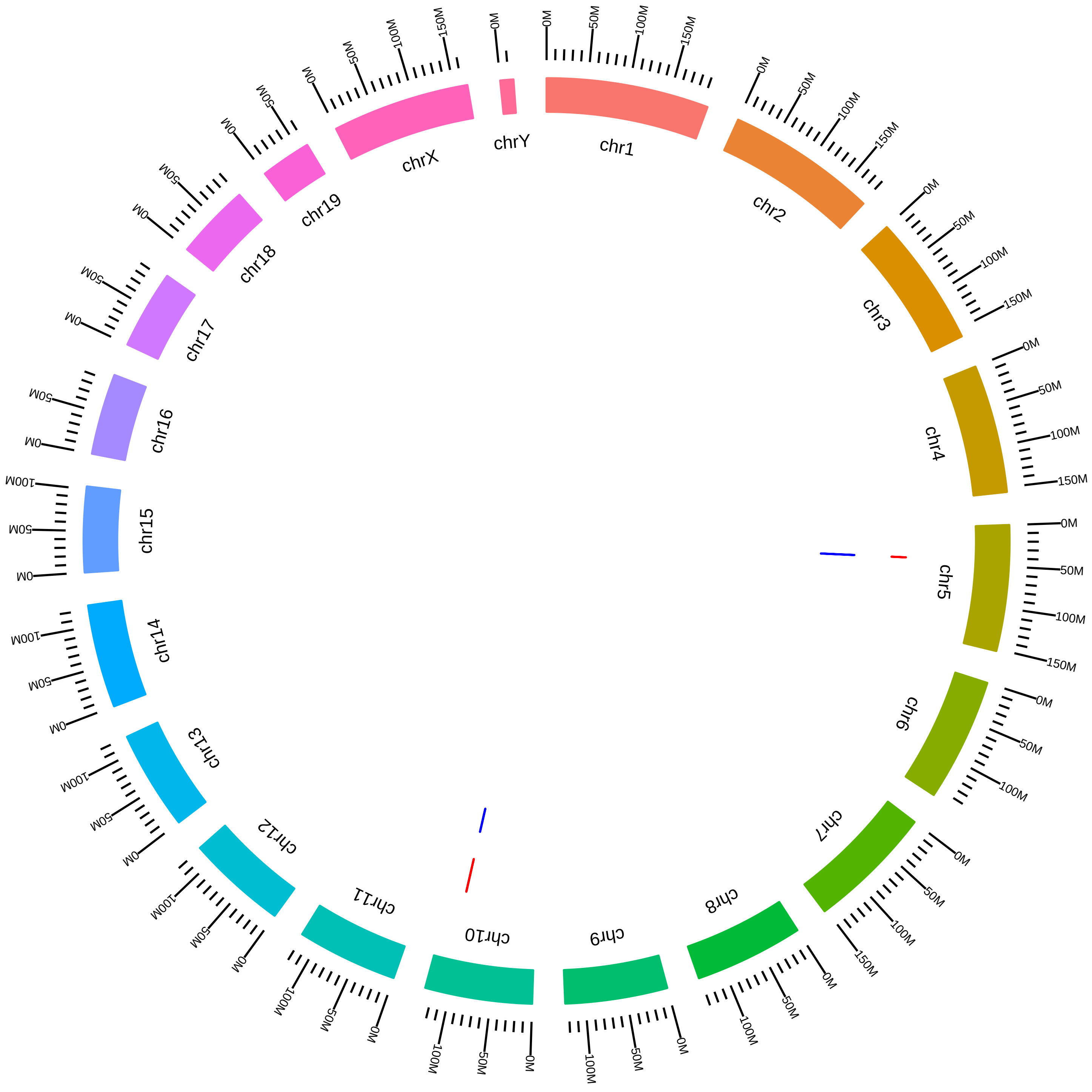

### rsfPhf5aPaf1MmIAifantis2016_AVsC.pdf

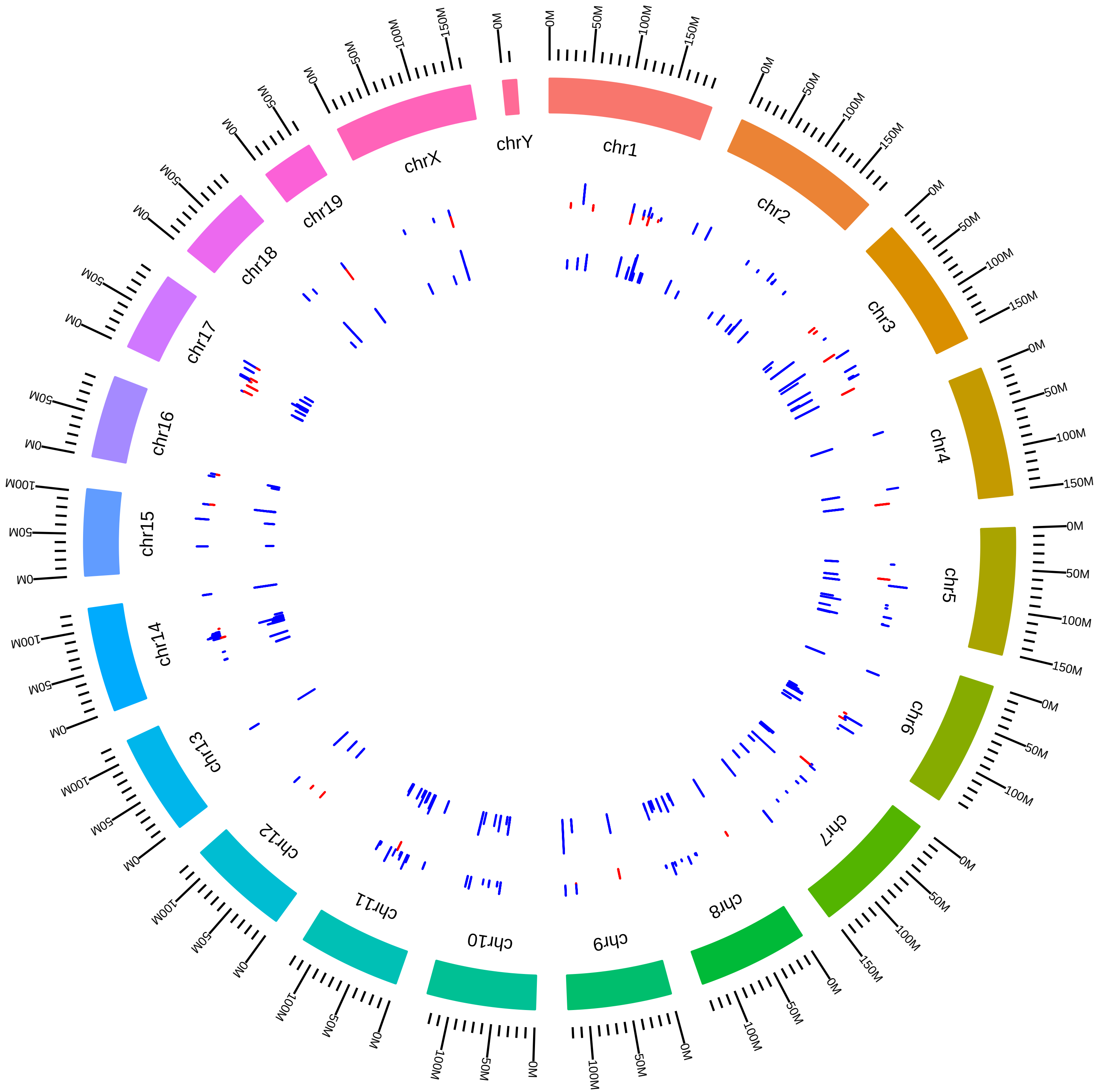

### rsfPhf5aPaf1MmIAifantis2016_HVsC.pdf

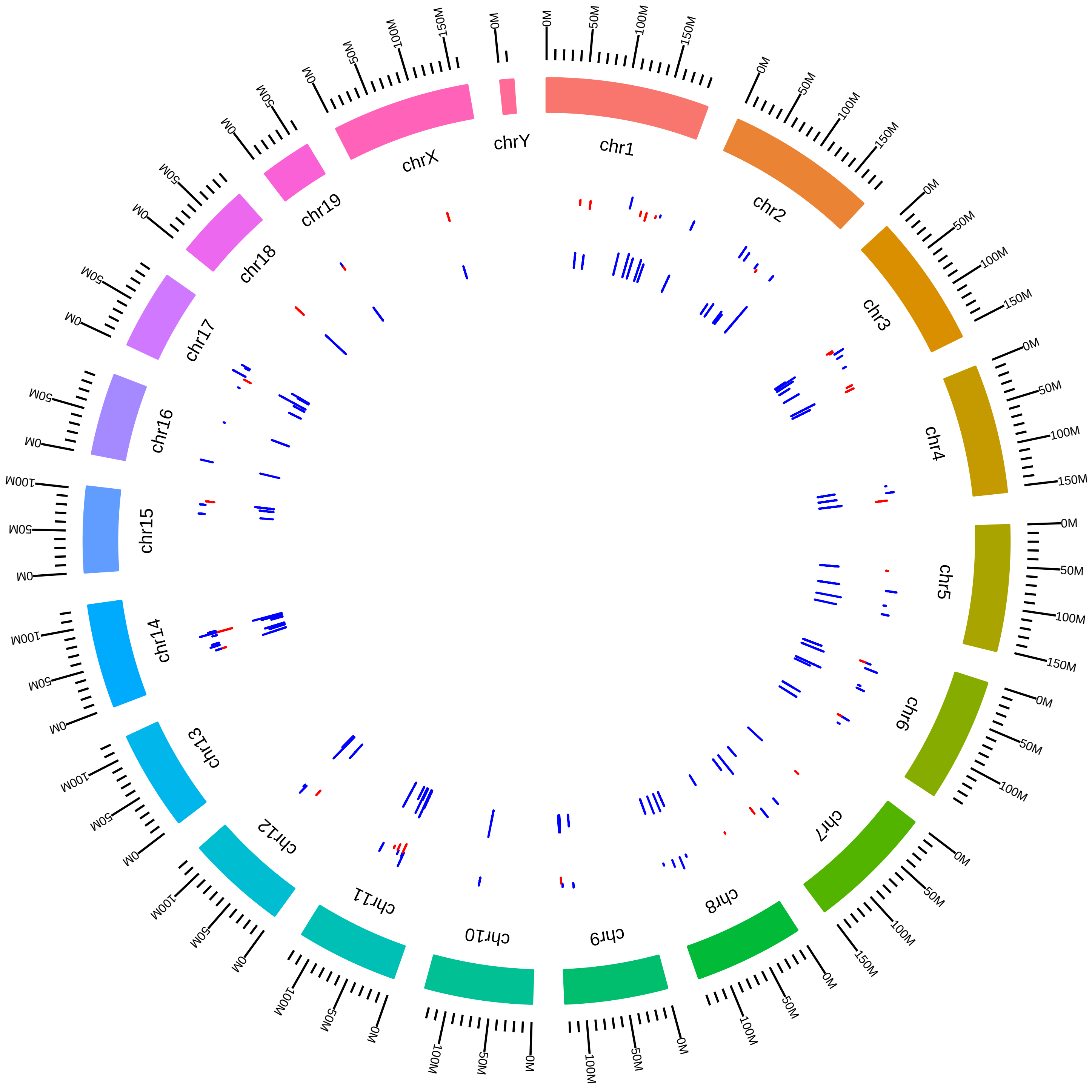

### rsfPqbp1MmPSilver_KD_vs_WT.pdf

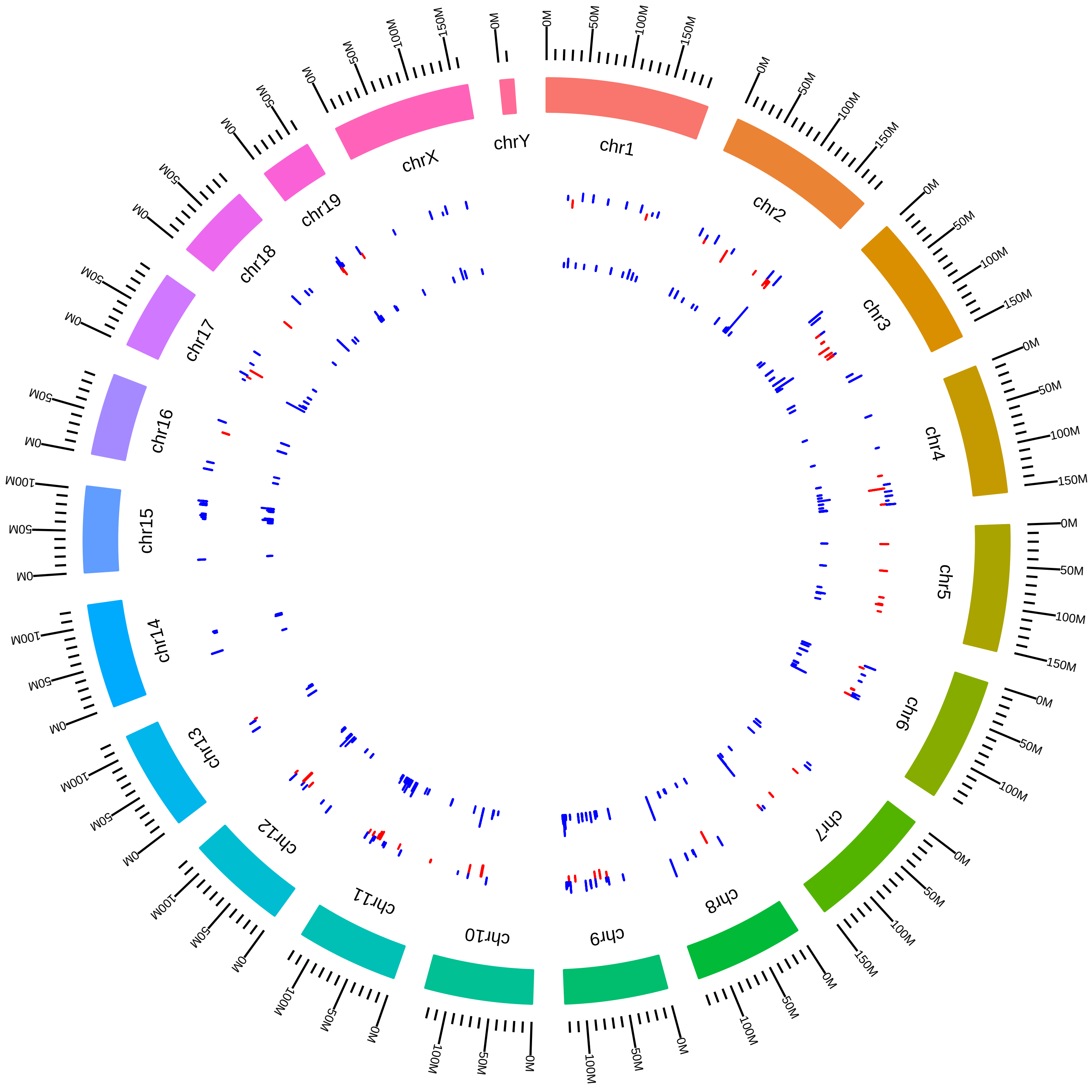

### rsfPrmt5MmATClark_KO_vs_WT.pdf

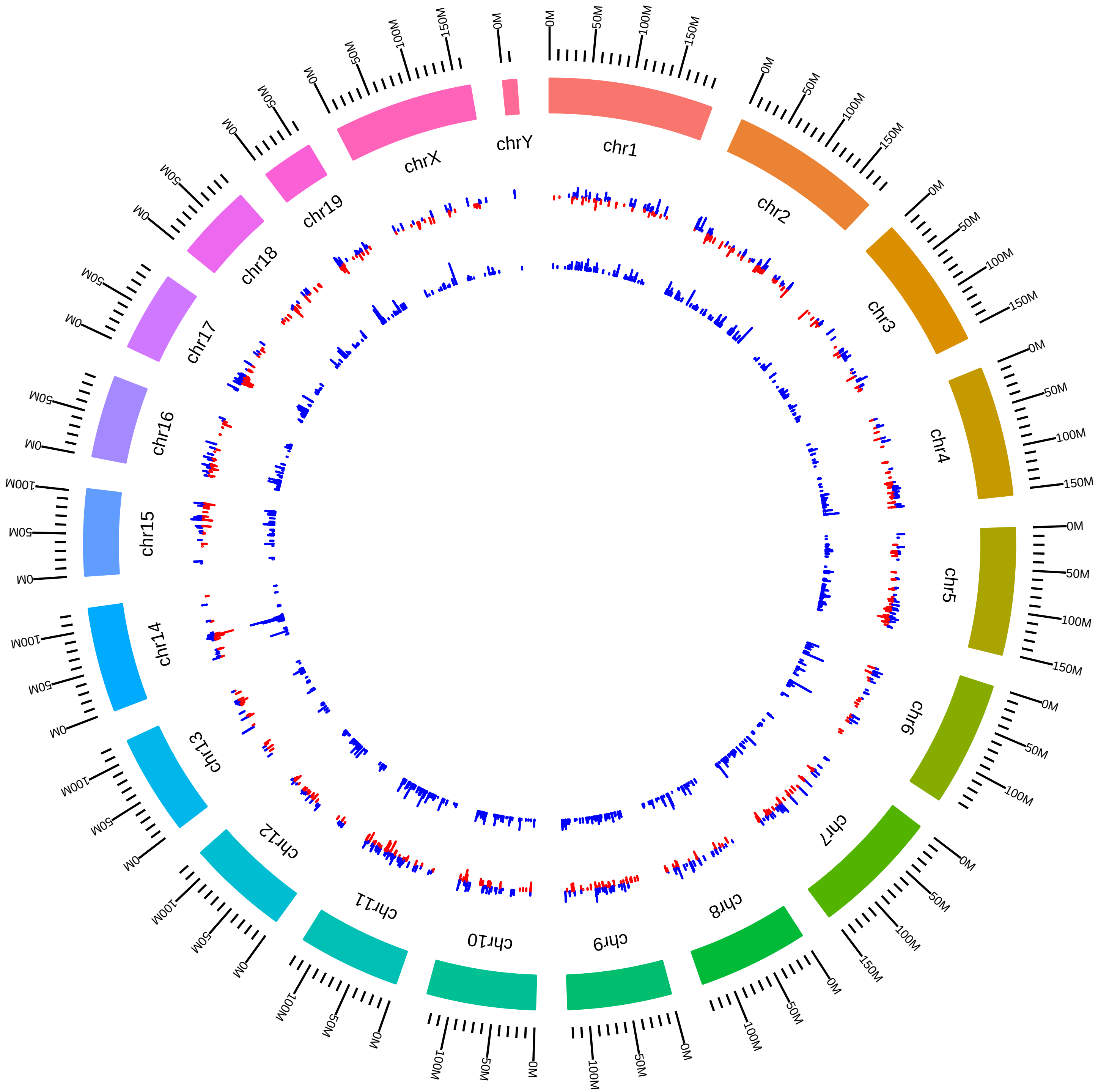

### rsfPrmt5MmEGuccione2015_KVsC.pdf

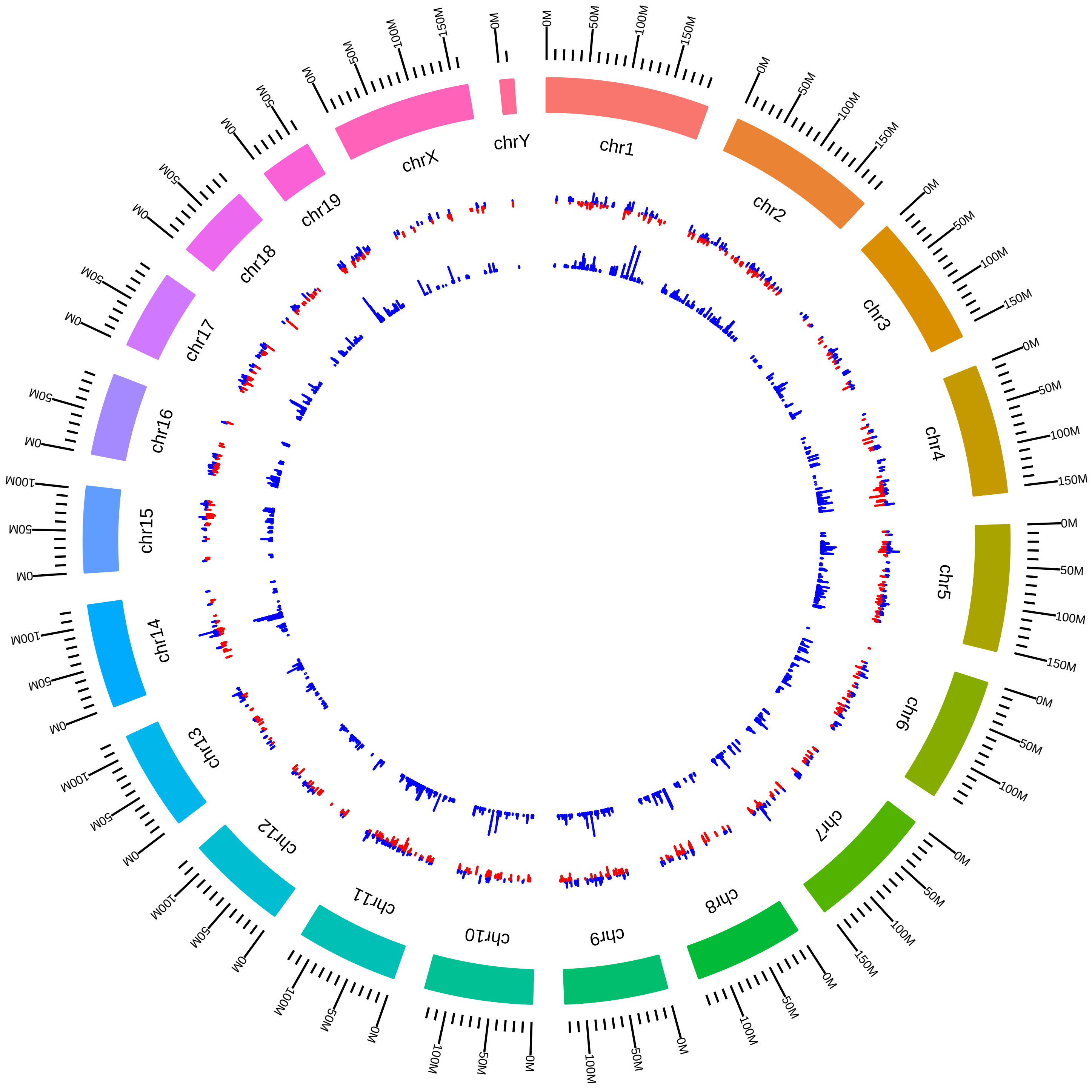

### rsfPrmt5MmEGuccioneCondition_MK_vs_MW.pdf

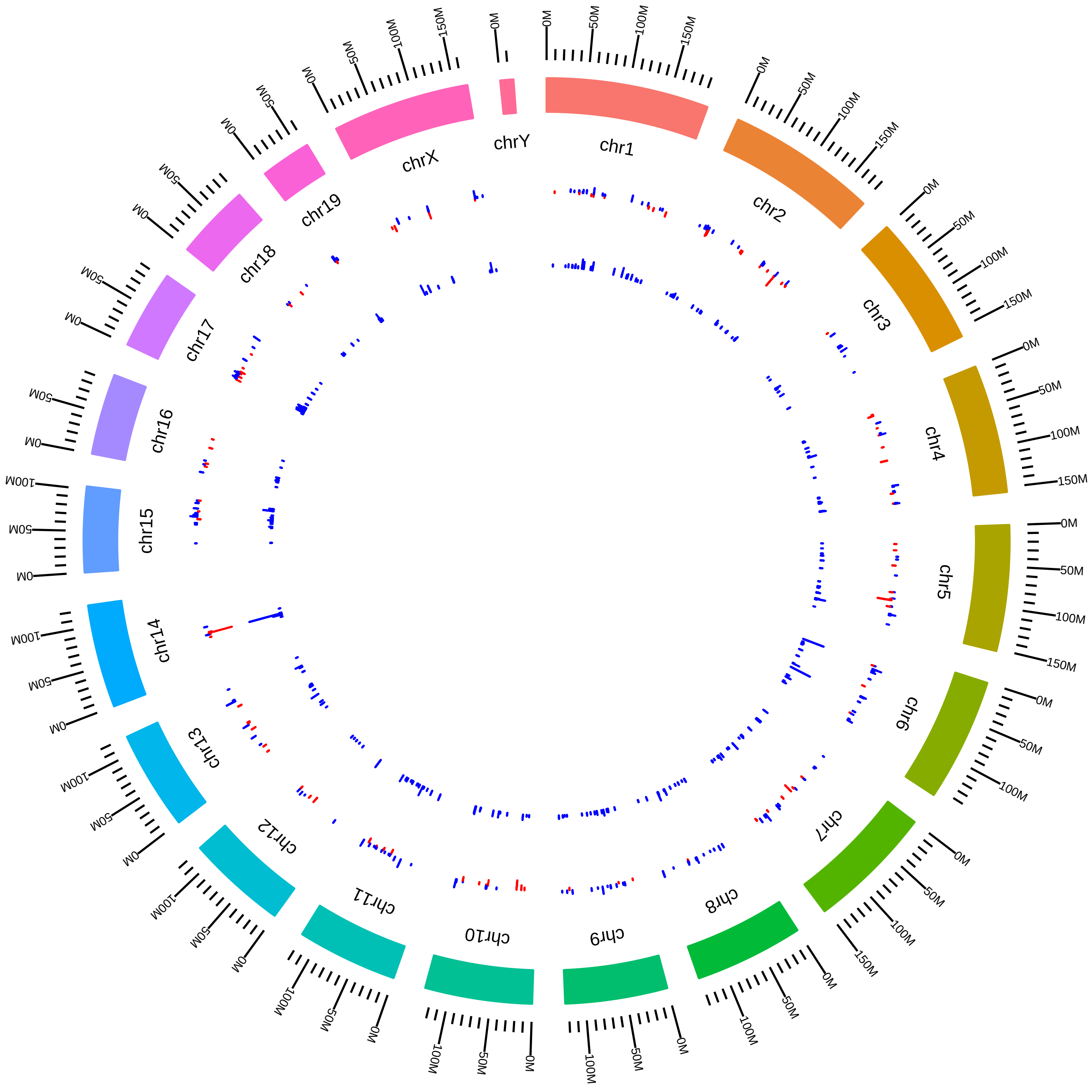

### rsfPrmt5MmEGuccioneCondition_NK_vs_NW.pdf

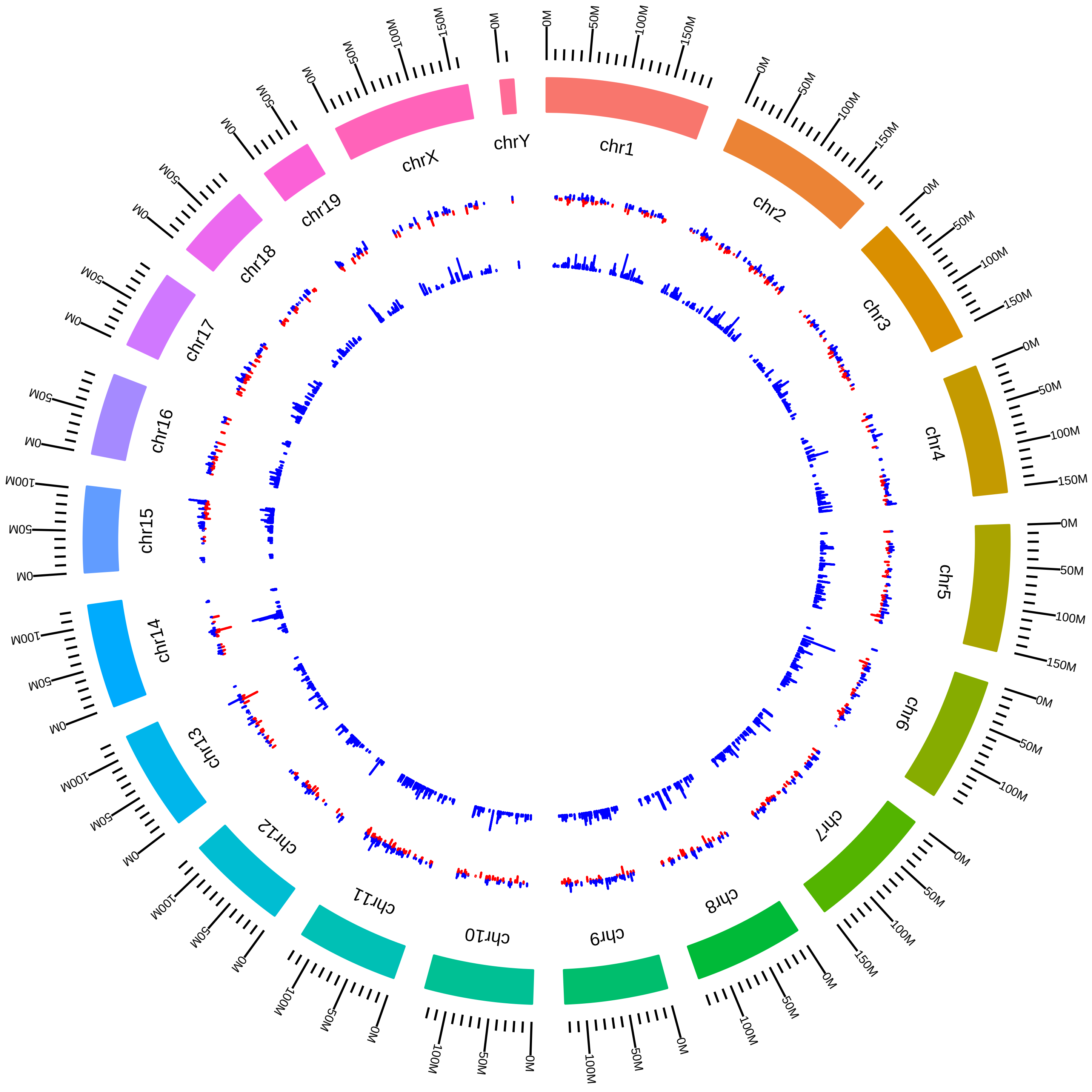

### rsfPrmt5MmJKim2015_KVsW.pdf

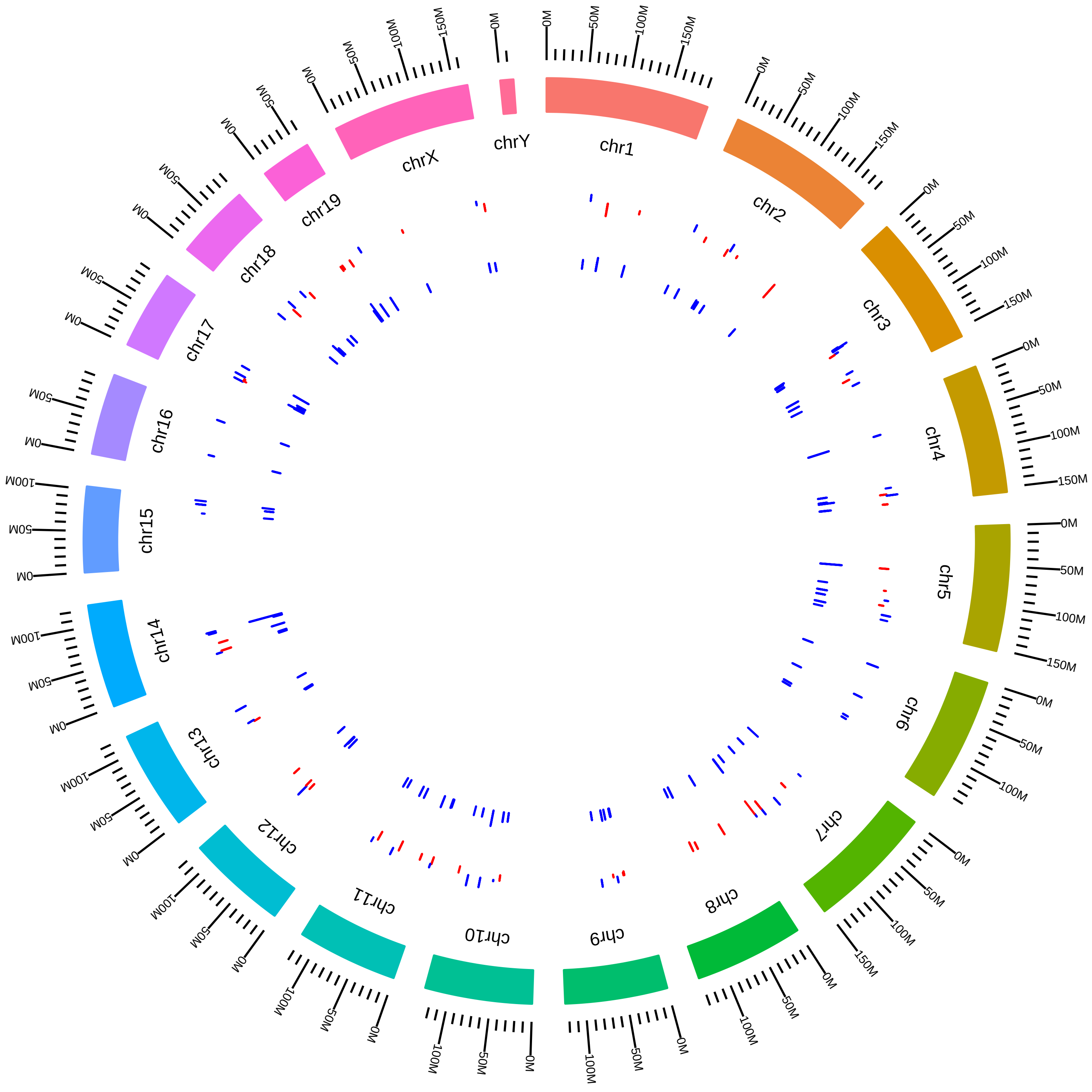

### rsfPrmt5MmSDNimer_KO_vs_WT.pdf

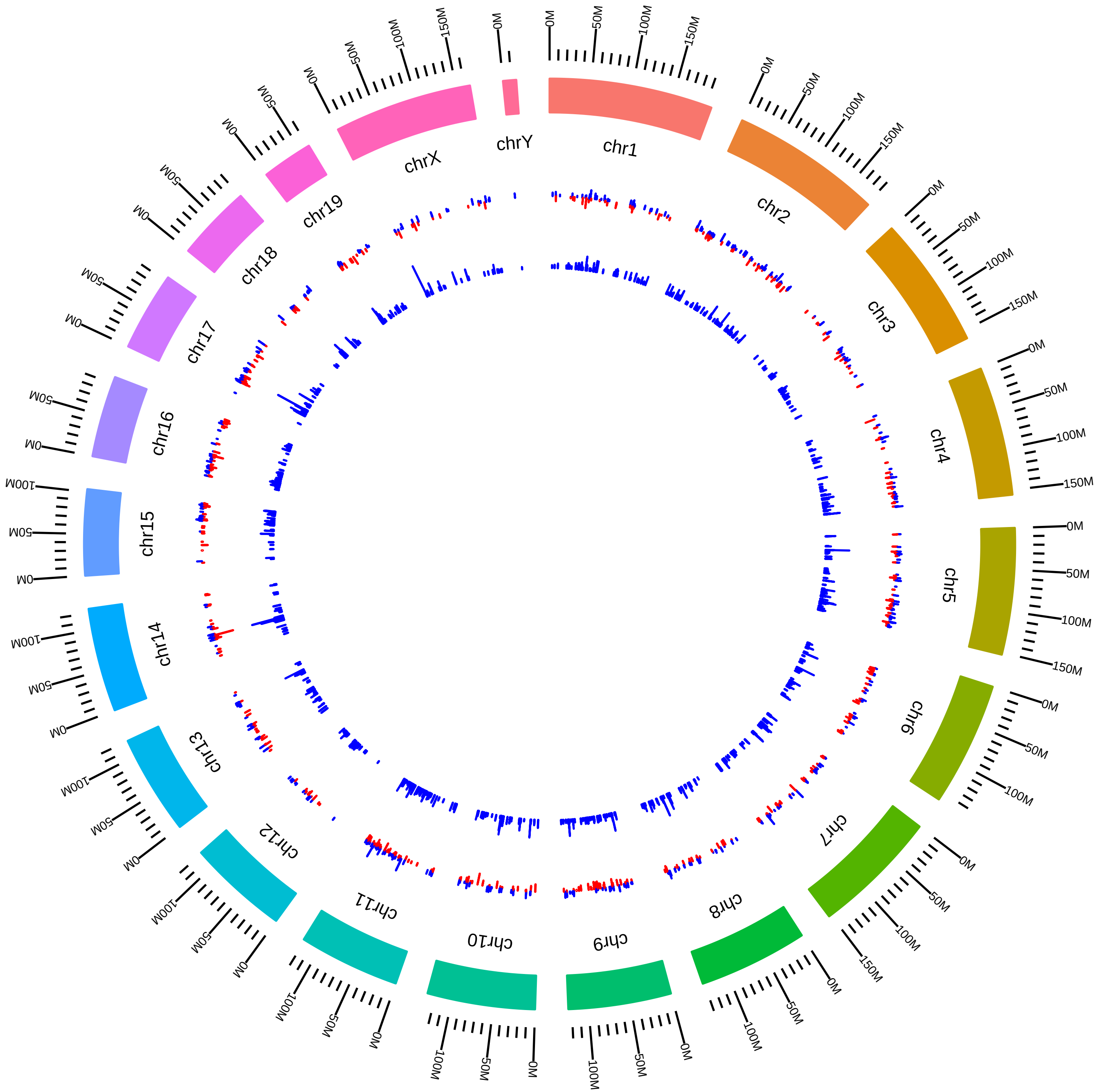

### rsfPrmt5MmSVokes2016_KVsW.pdf

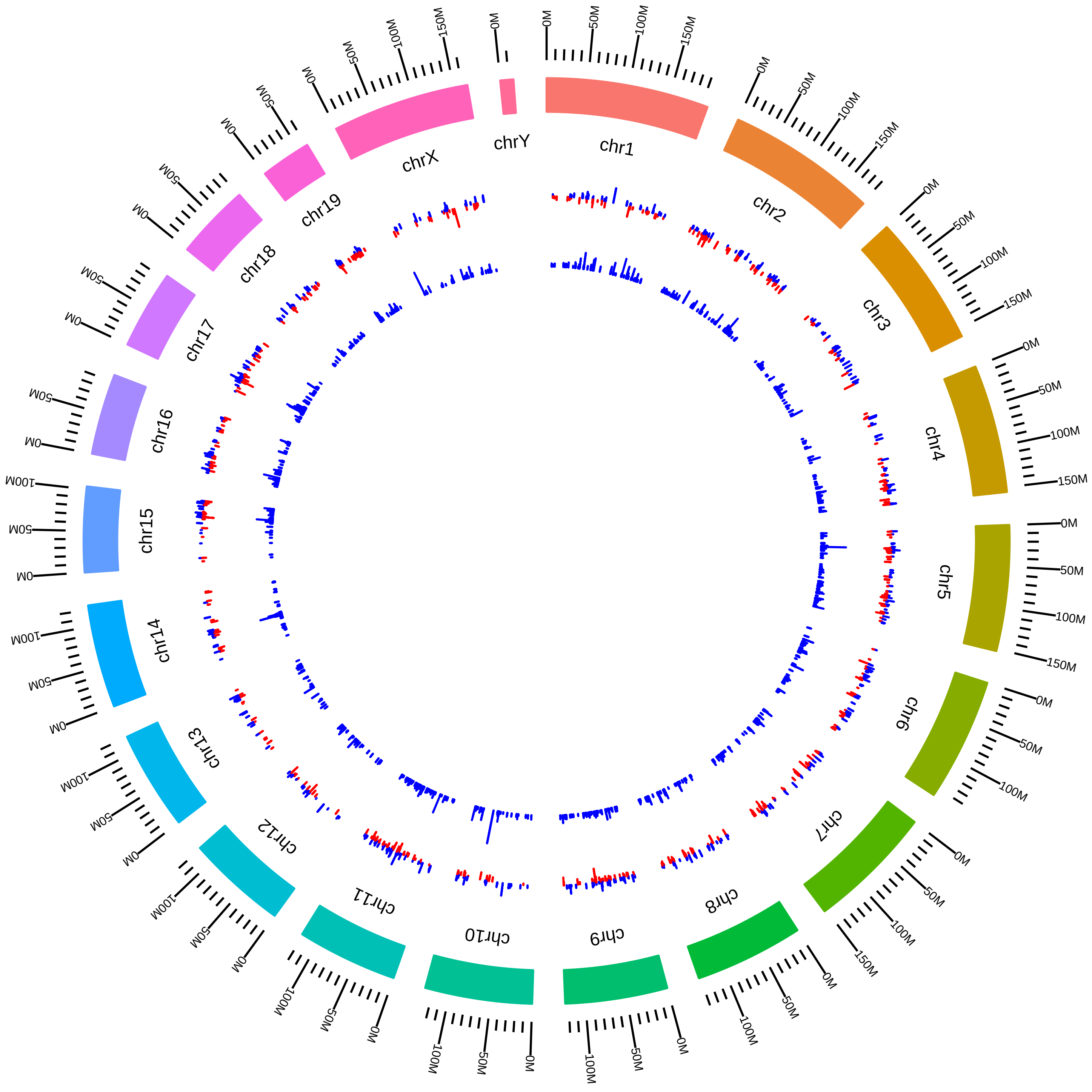

### rsfPrmt7MmSRichardreverse_KO_vs_WT.pdf

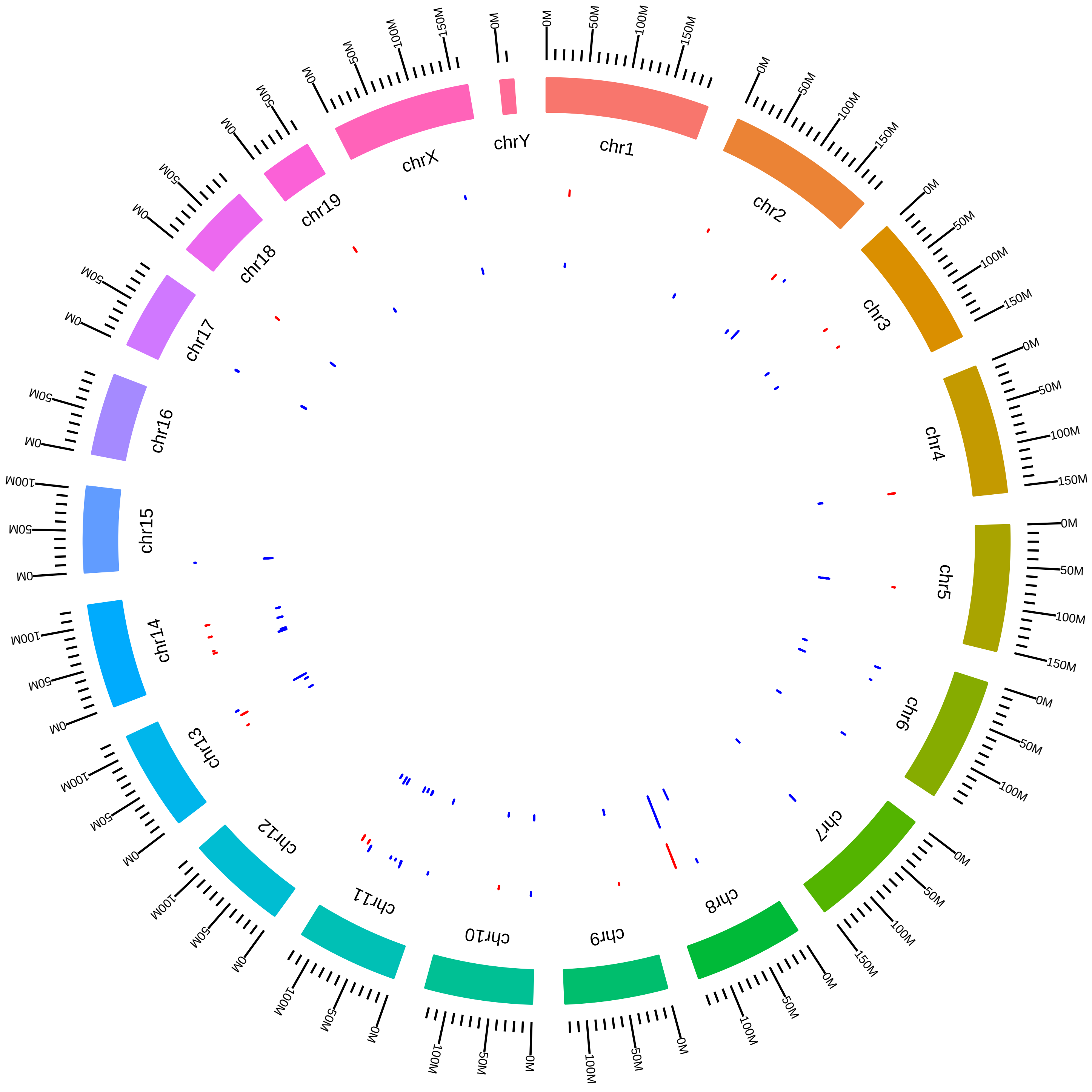

### rsfPtbp1MmDLim_KD_vs_WT.pdf

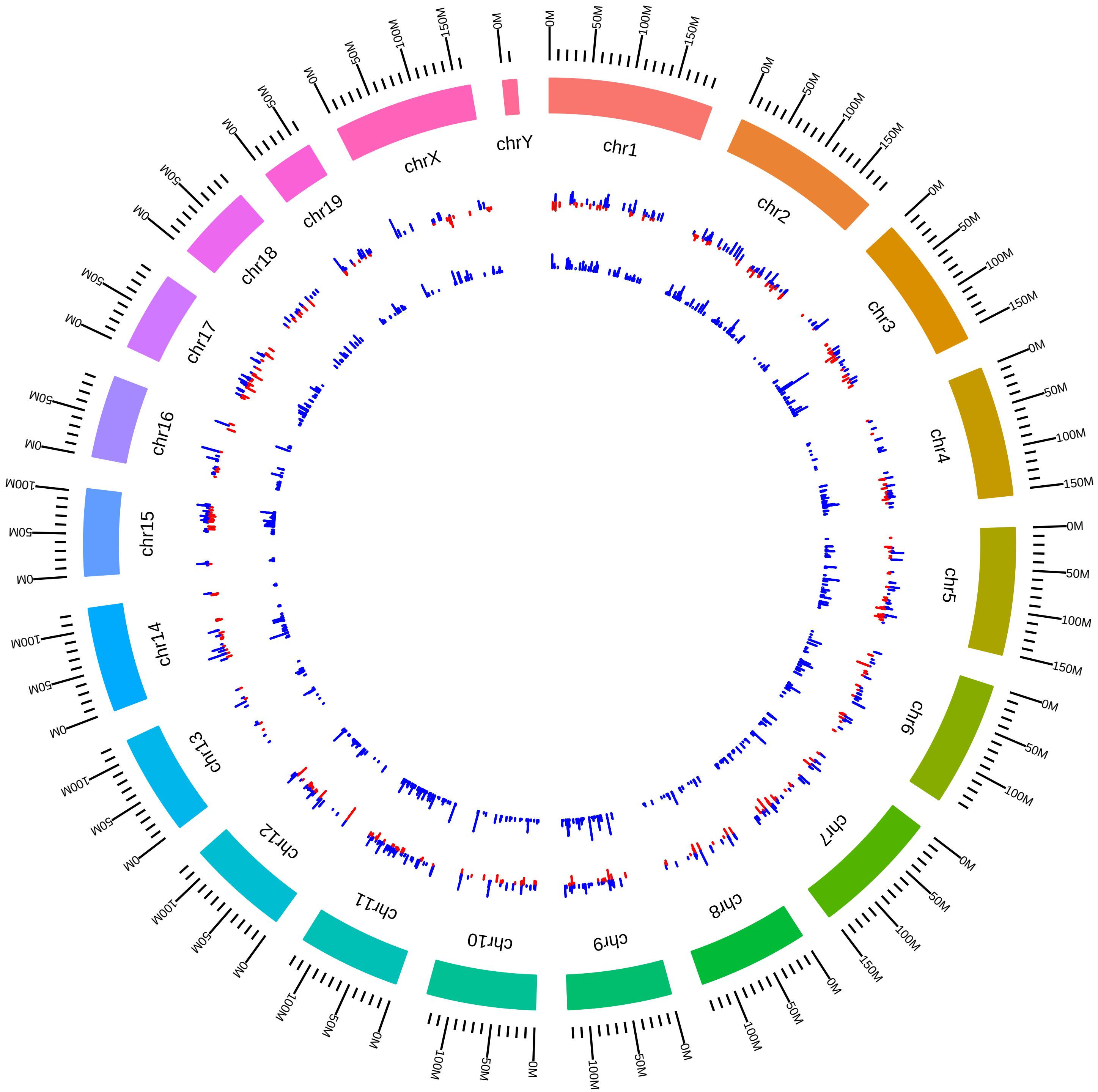

### rsfPtbp1Ptbp2MmDBlackmESC_DKD_vs_WT.pdf

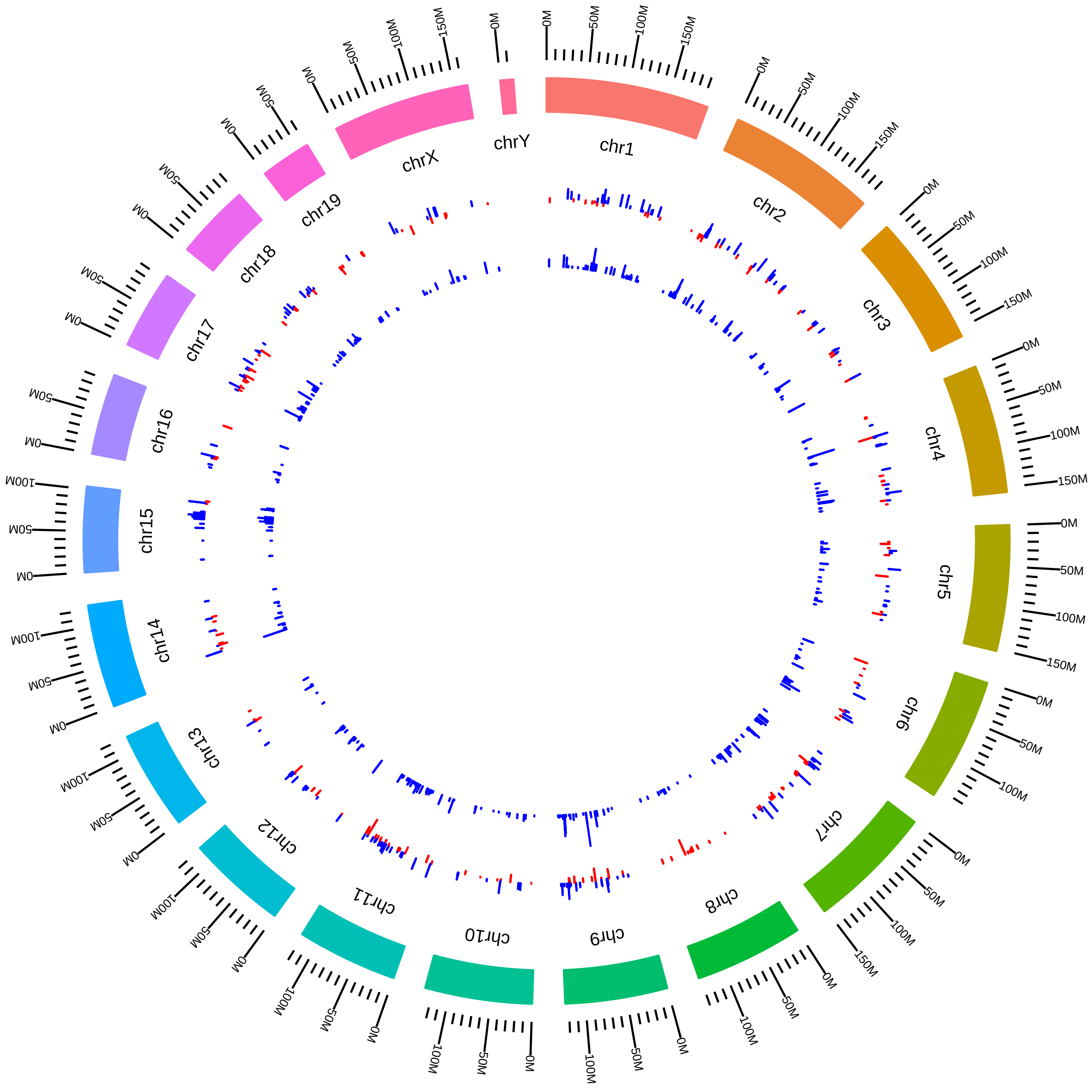

### rsfPtbp1Ptbp2MmDBlackmESC_Ptbp1KD_vs_WT.pdf

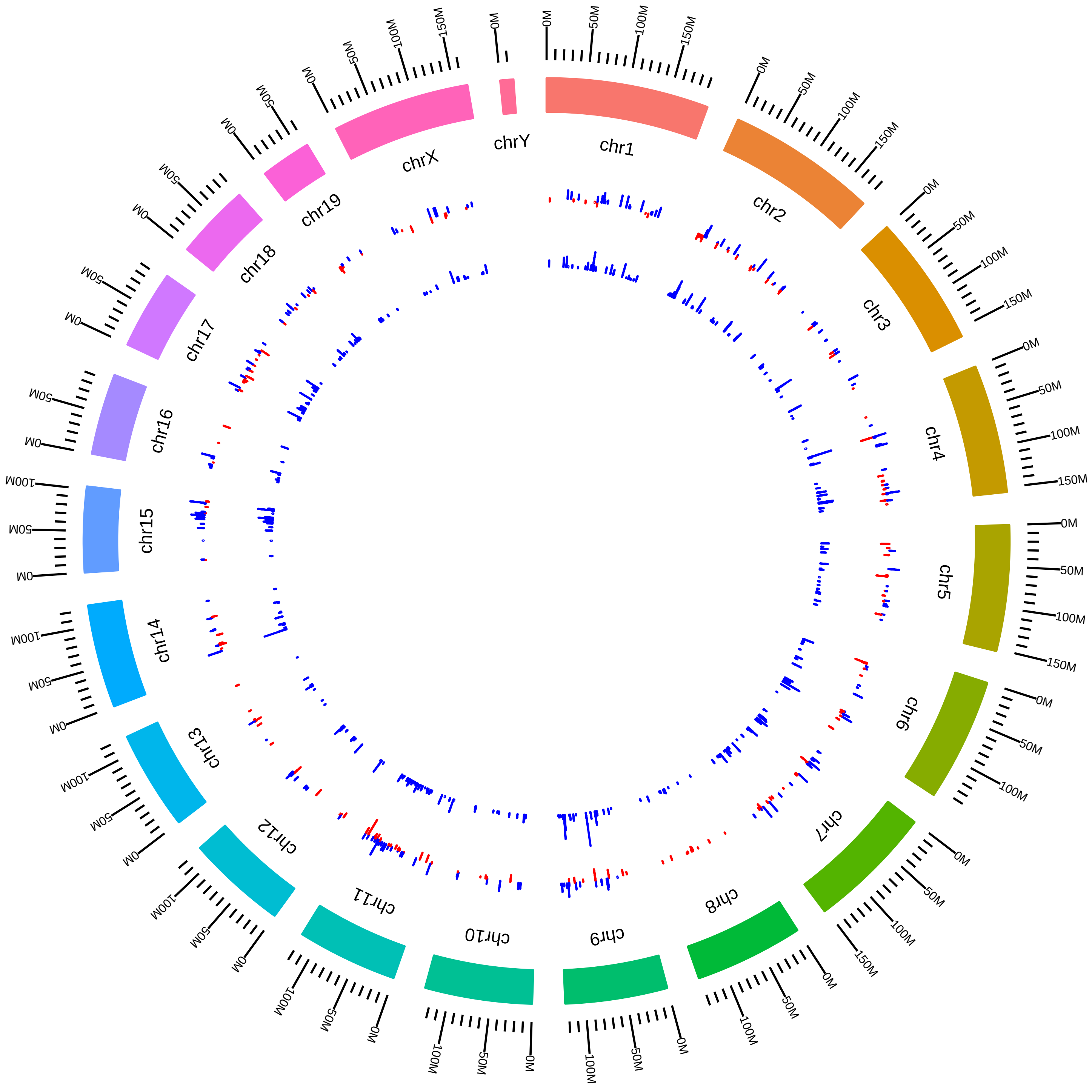

### rsfPtbp1Ptbp2MmDBlackmNPC_DKD_vs_WT.pdf

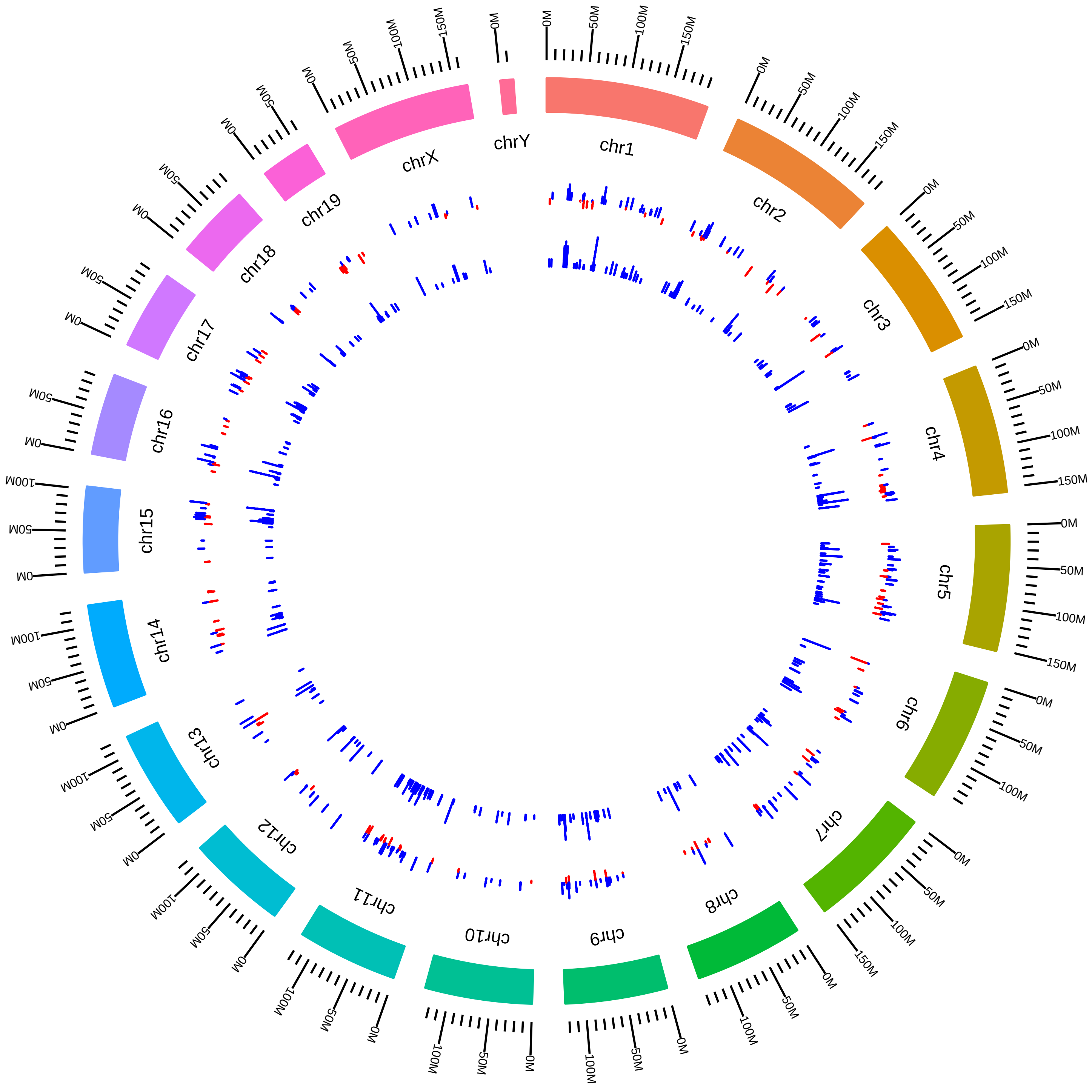

### rsfPtbp1Ptbp2MmDBlackmNPC_Ptbp1KD_vs_WT.pdf

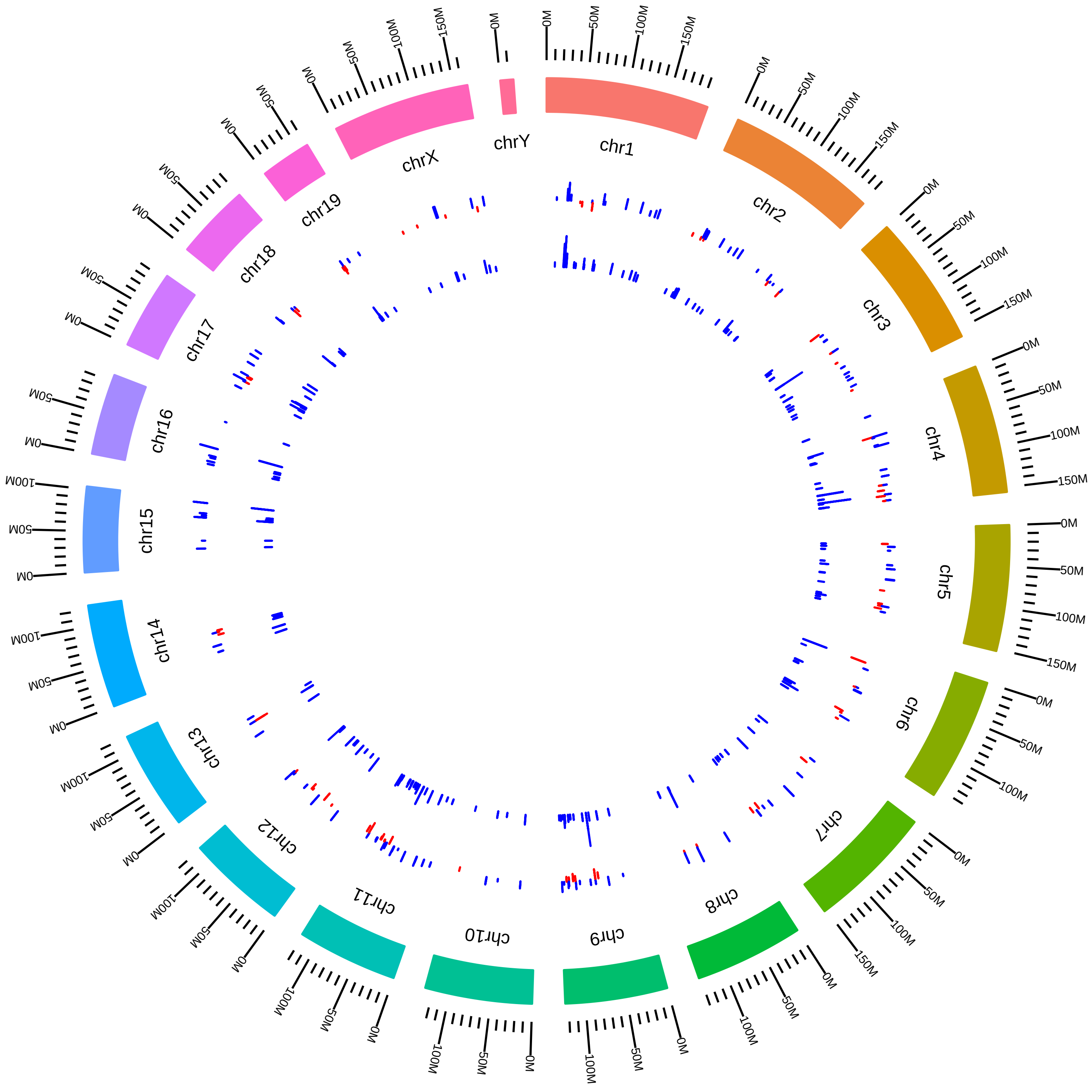

### rsfPtbp1Ptbp2MmDBlackmNPC_Ptbp2KD_vs_WT.pdf

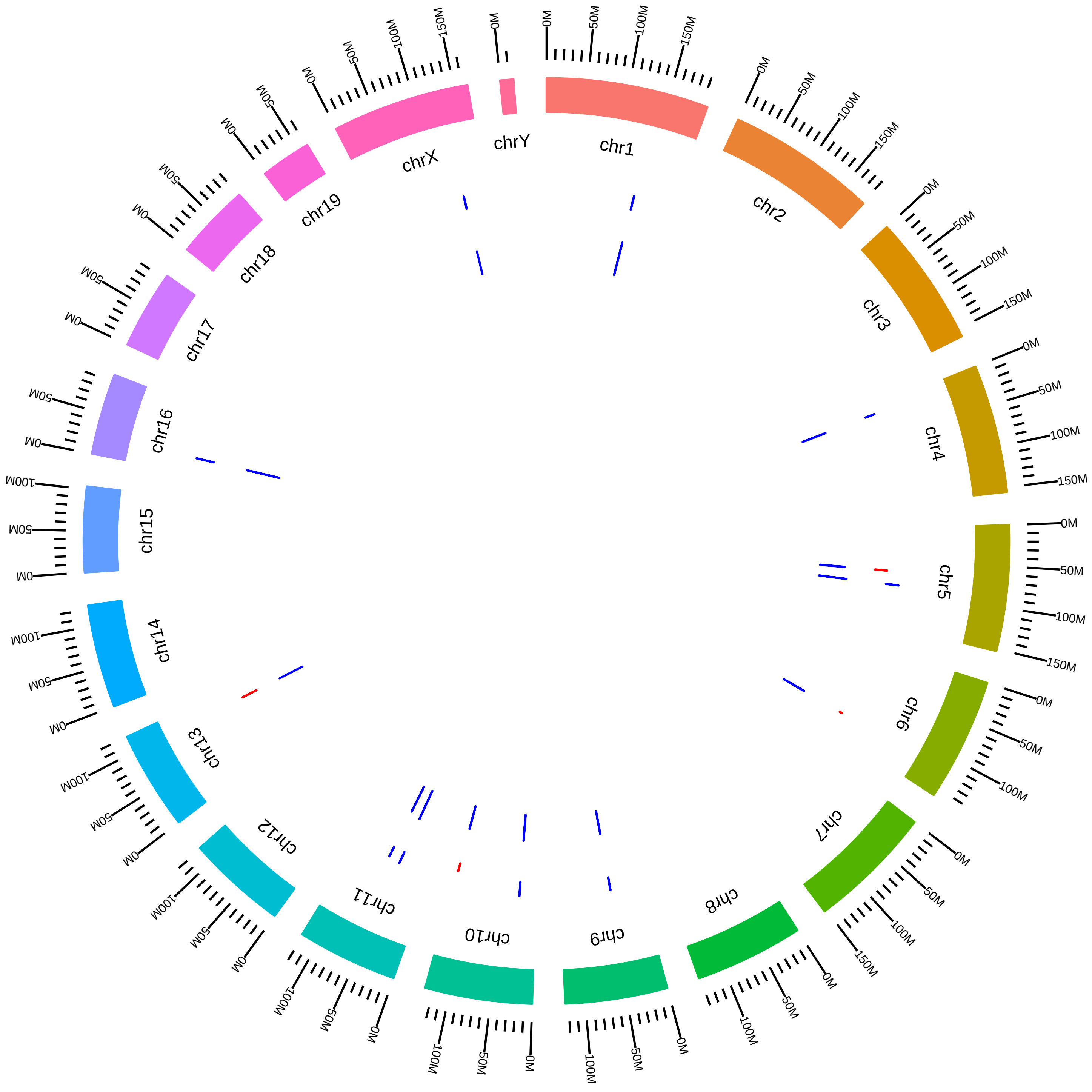

### rsfPtbp2MmDBlack_KO_vs_WT.pdf

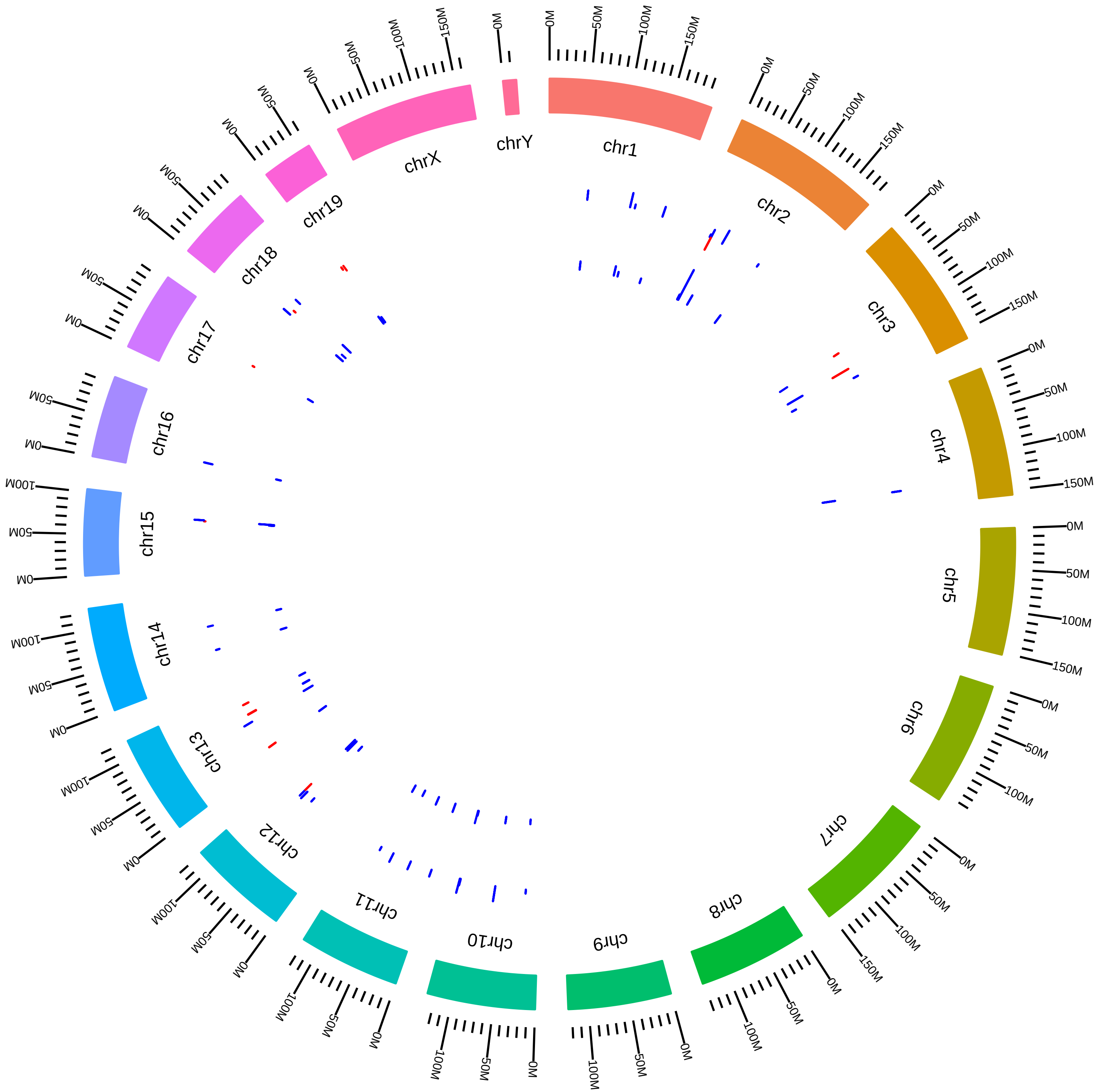

### rsfQkMmJianHu2017_KVsC.pdf

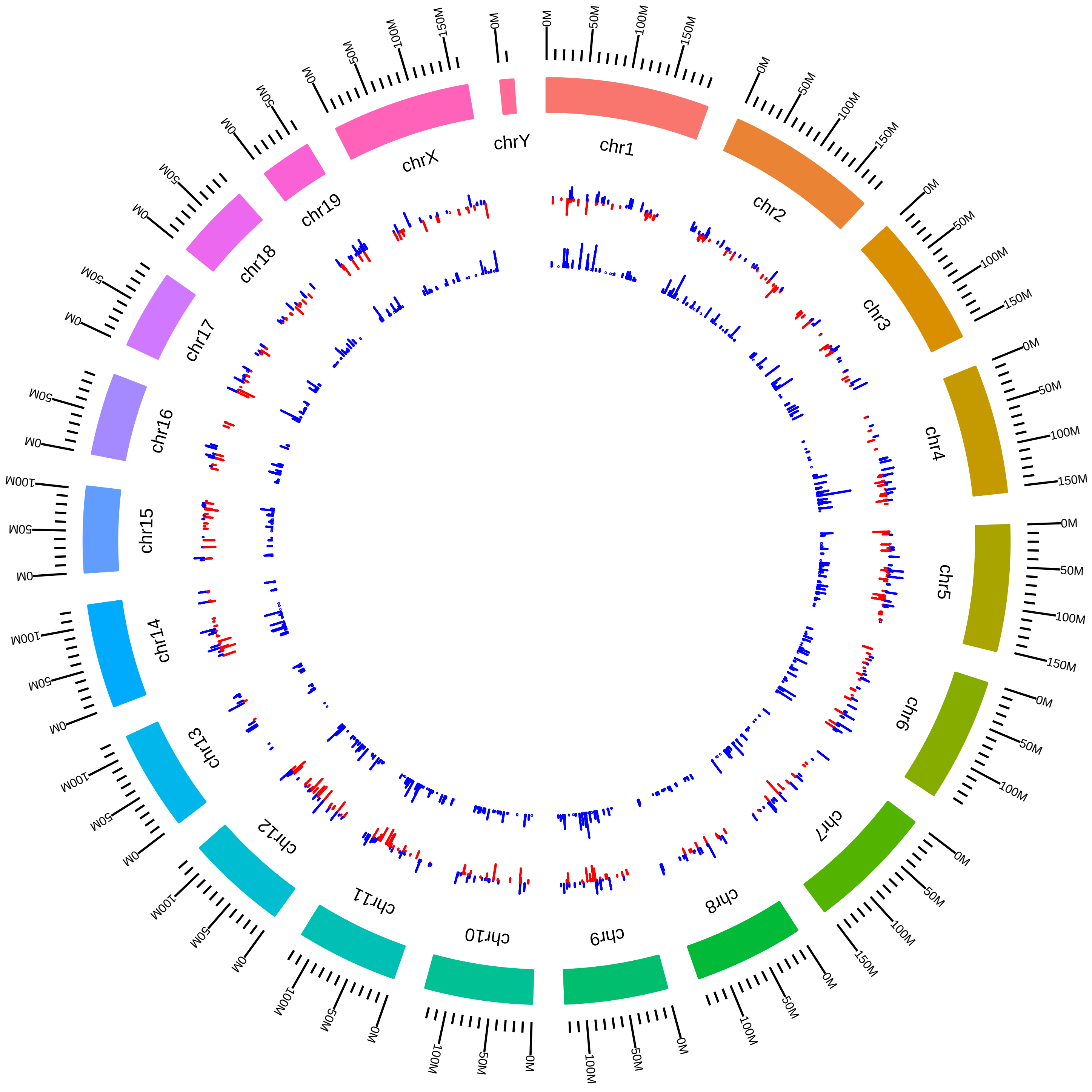

### rsfQkPtenTrp53MmJianHu2017_TVsD.pdf

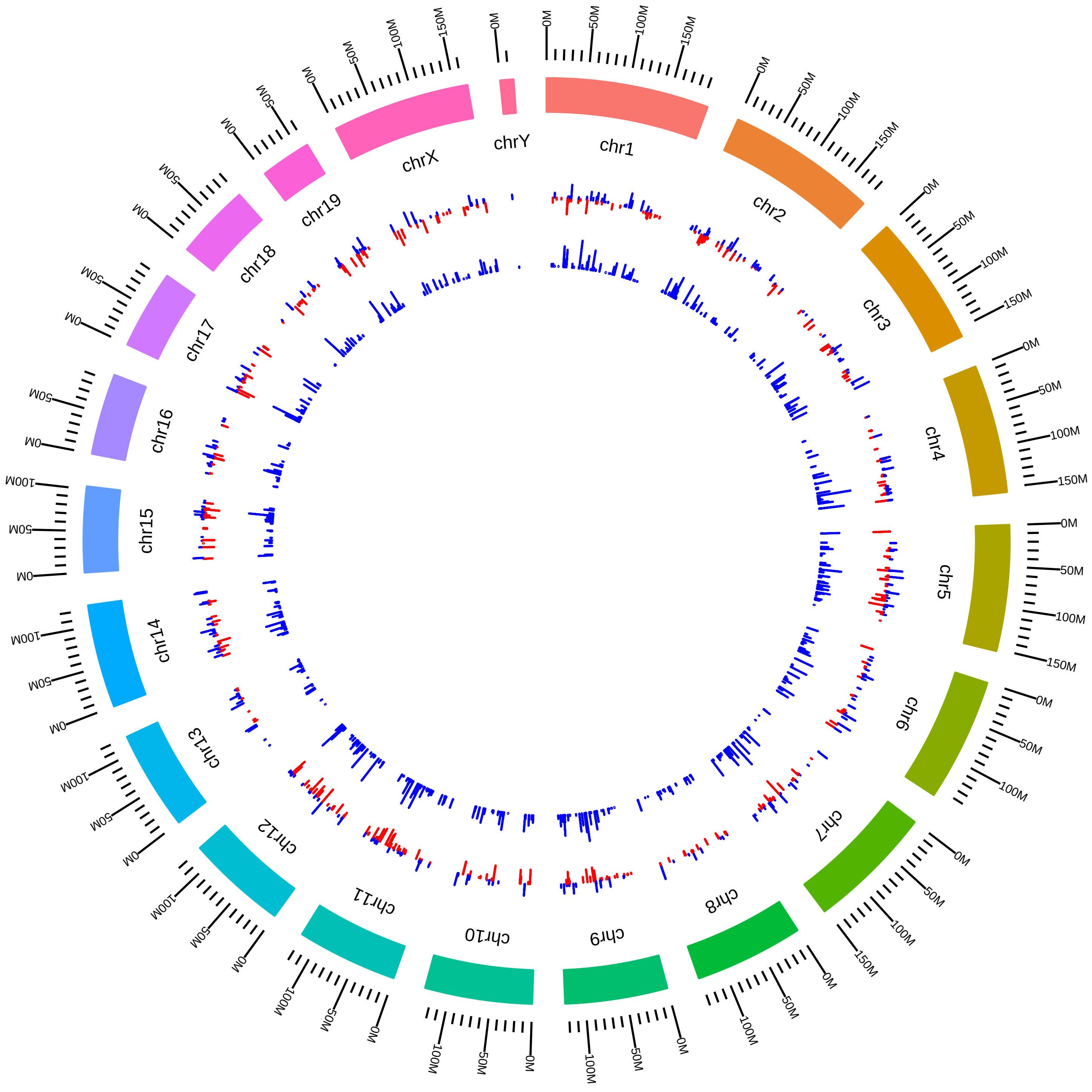

### rsfRbfox1Rbfox2MmTCooperMerge_Rbfox1KD_vs_WT.pdf

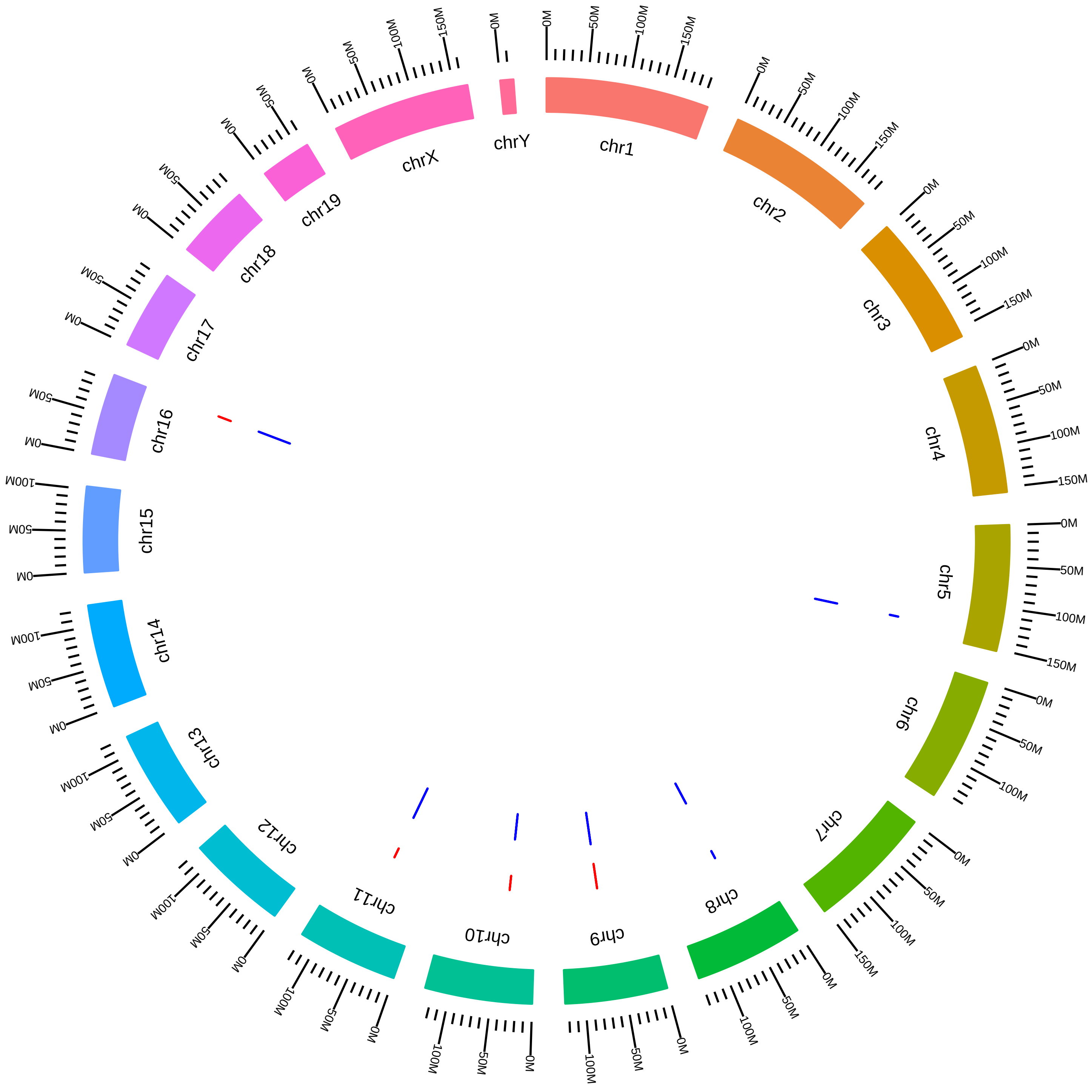

### rsfRbfox1Rbfox2MmTCooperMerge_Rbfox2KD_vs_WT.pdf

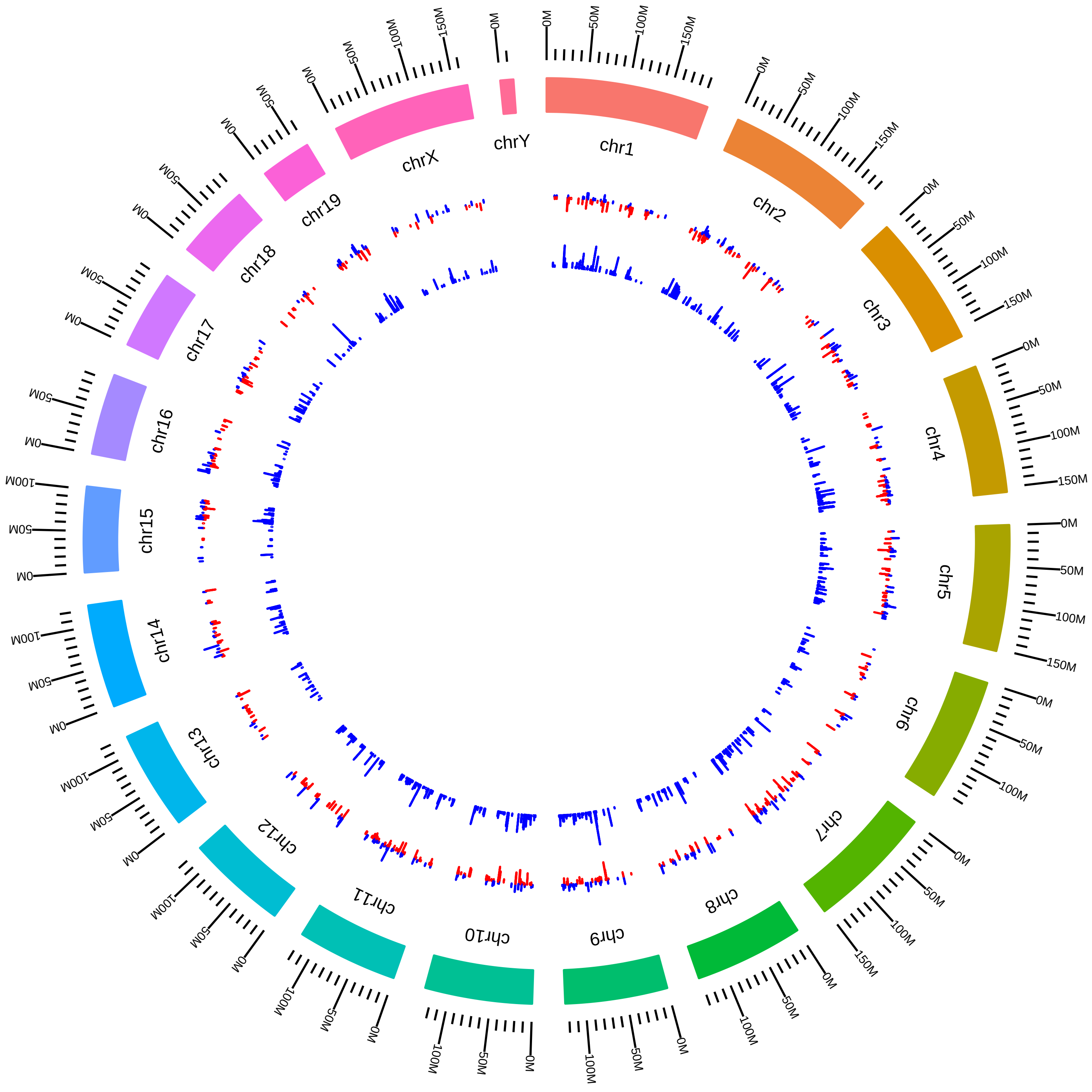

### rsfRbfox1Rbfox3MmKMartin2016_DVsC.pdf

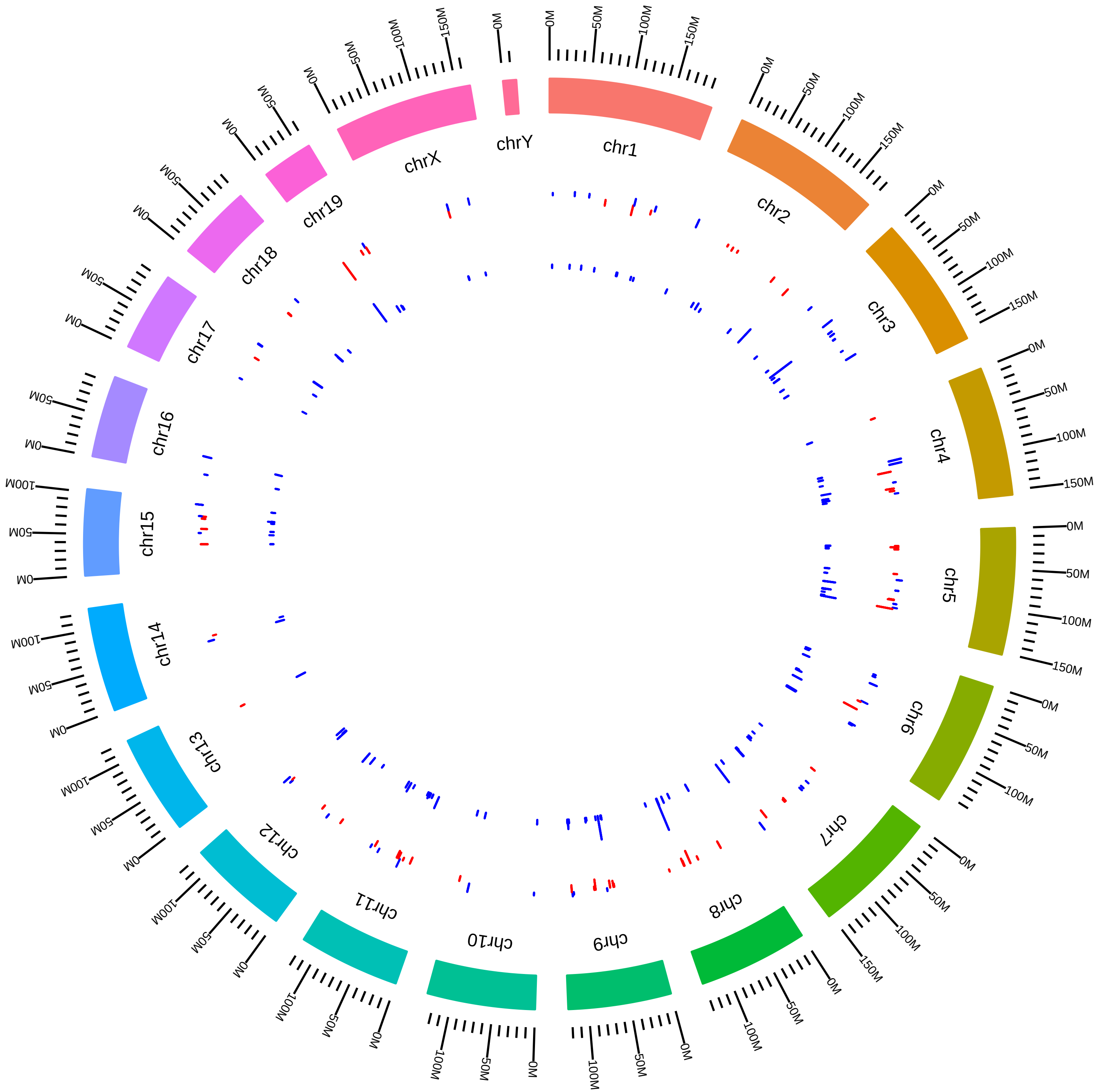

### rsfRbfox1Rbfox3MmKMartin2016_NVsD.pdf

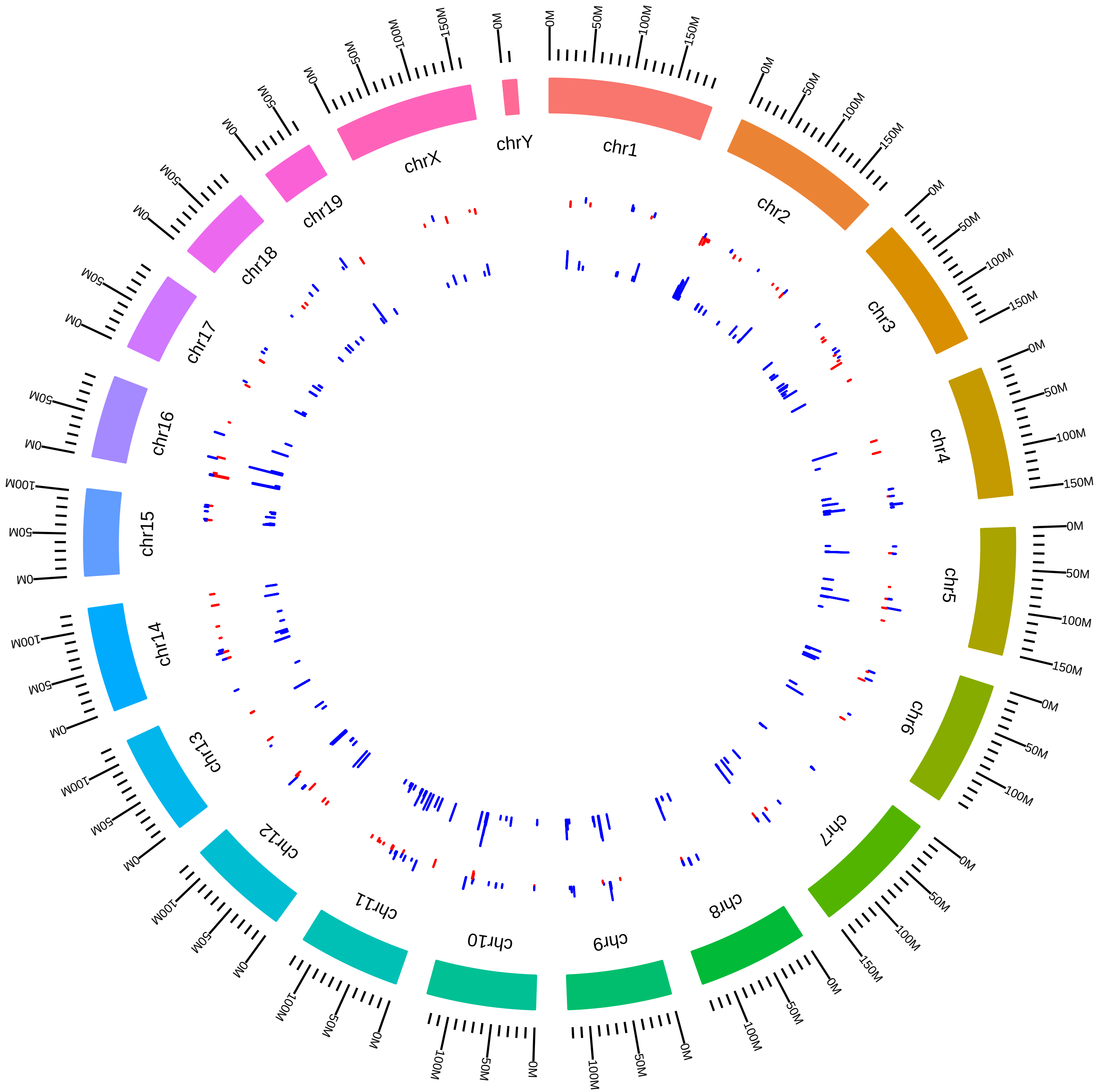

### rsfRbfox1Rbfox3MmKMartin2016_YVsD.pdf

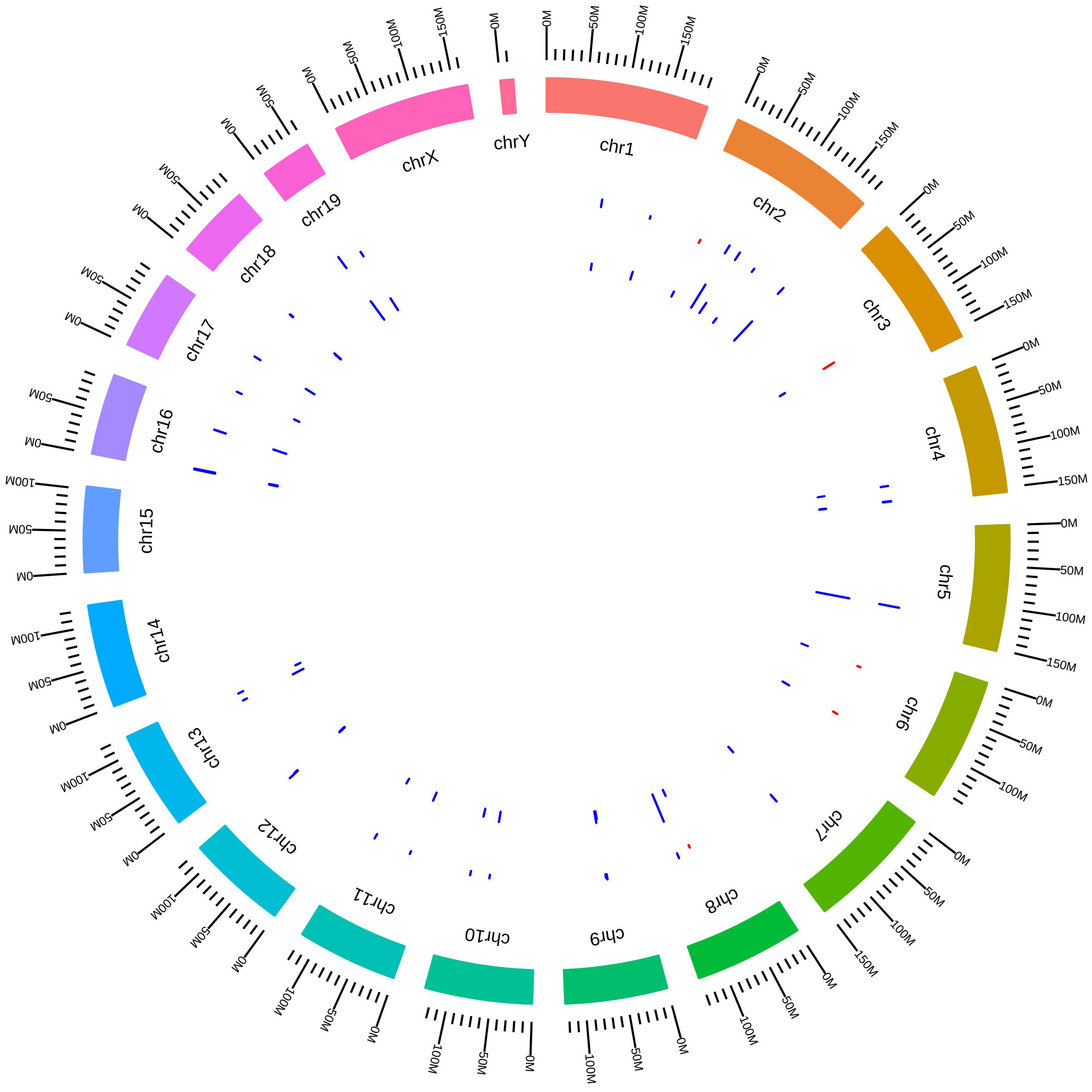

### rsfRbfox2MmPSharpMerge_KD_vs_WT.pdf

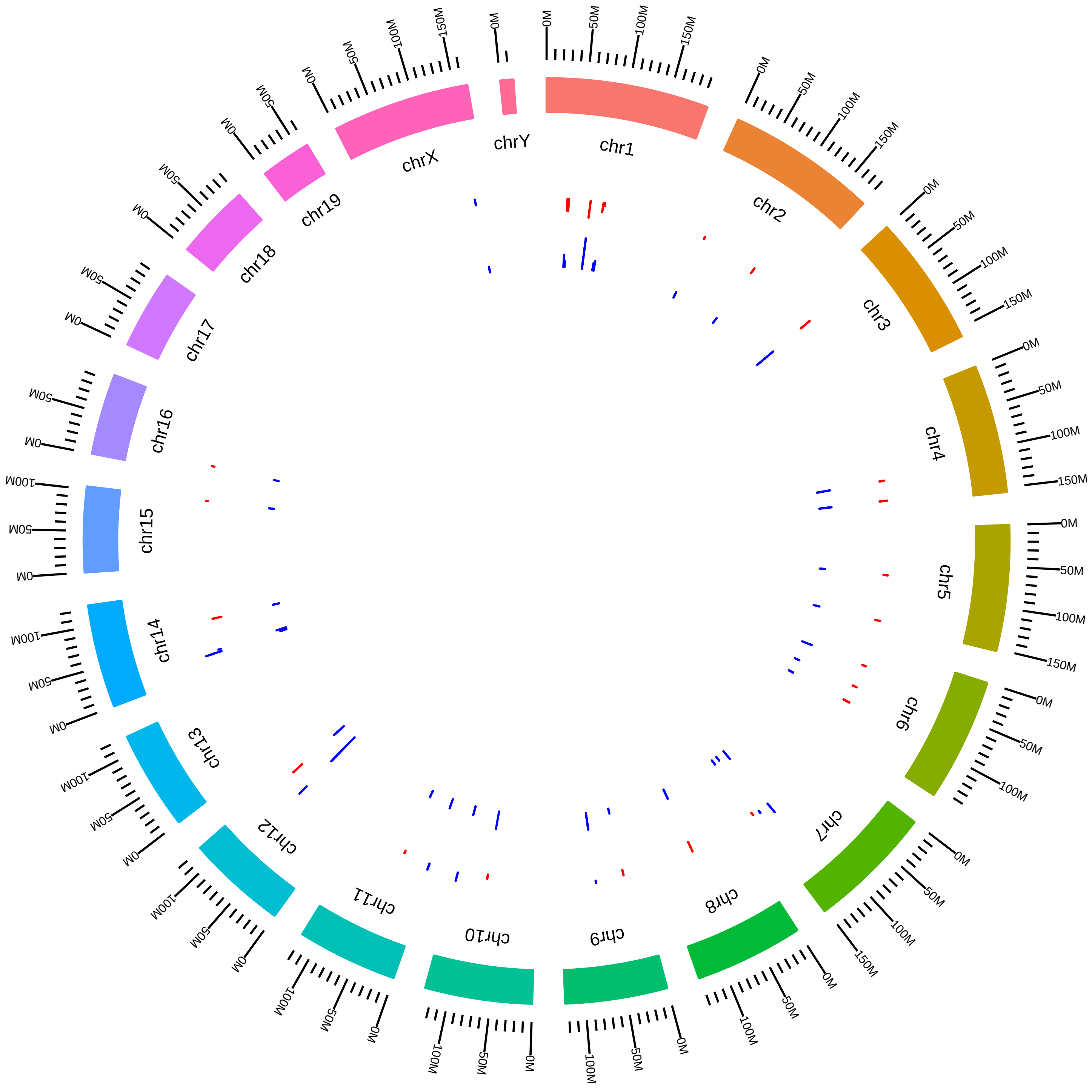

### rsfRbm10ESMmJCaceres2017_RVsW.pdf

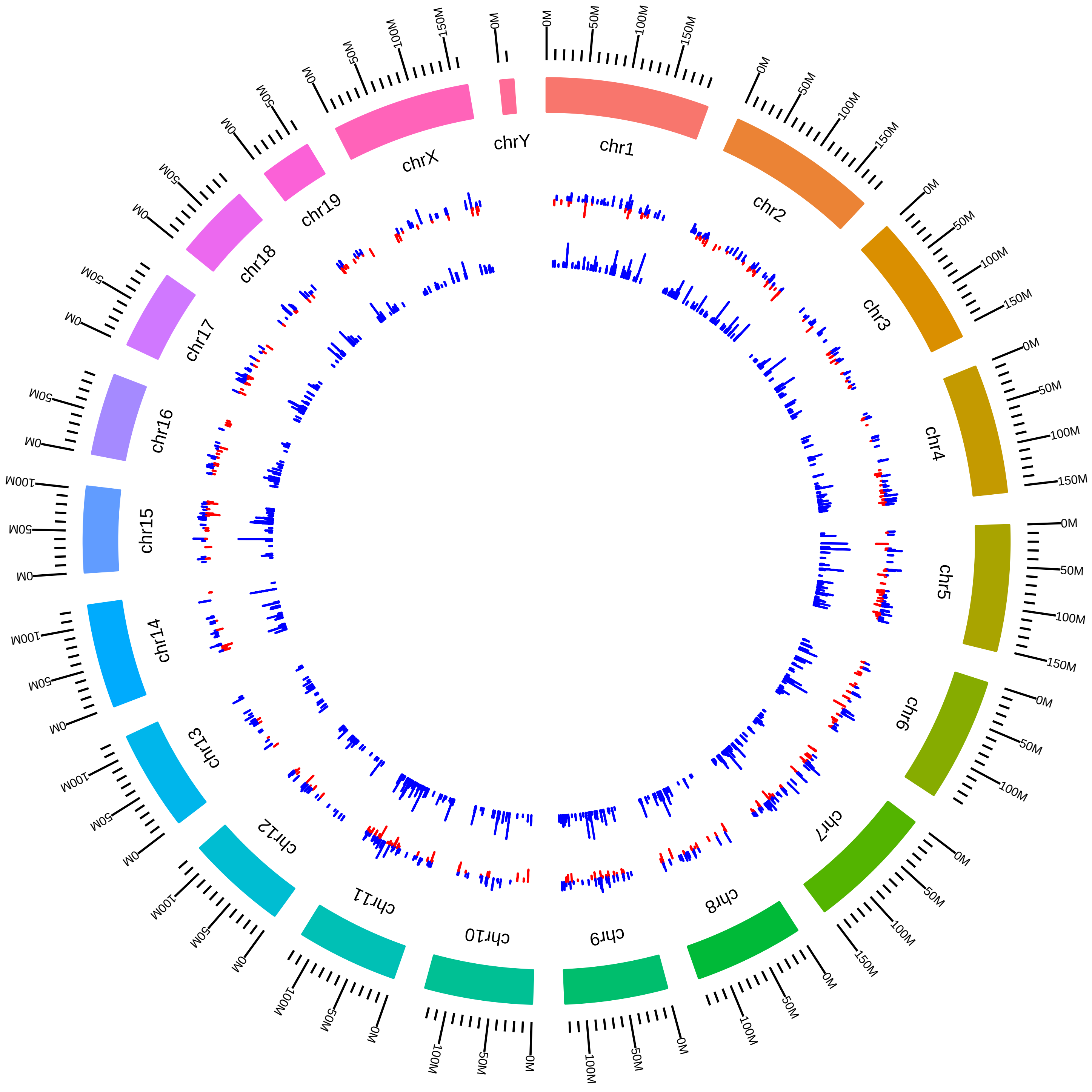

### rsfRbm10MNMmJCaceres2017_RVsW.pdf

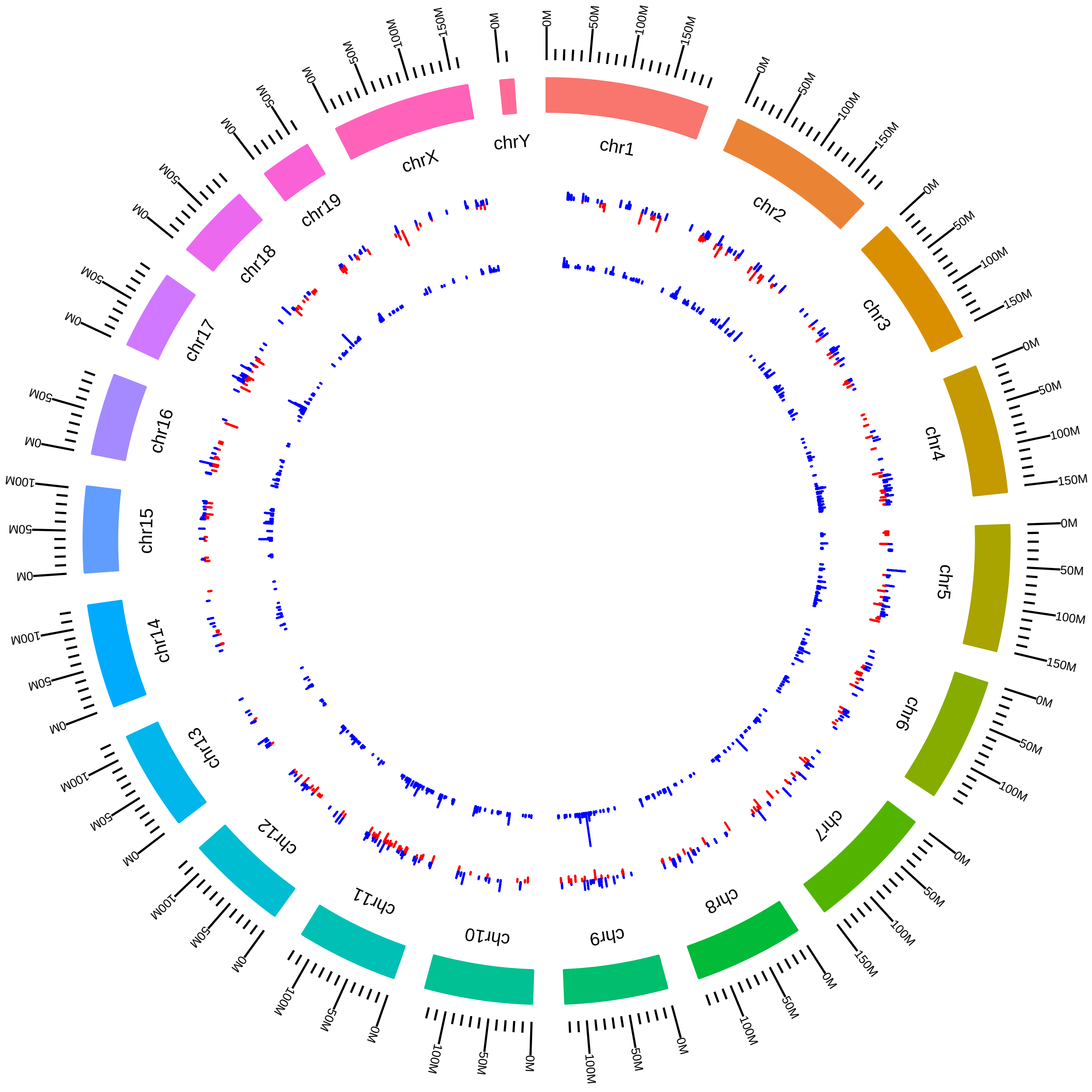

### rsfRbm17MmHZoghbi2016_KVsW.pdf

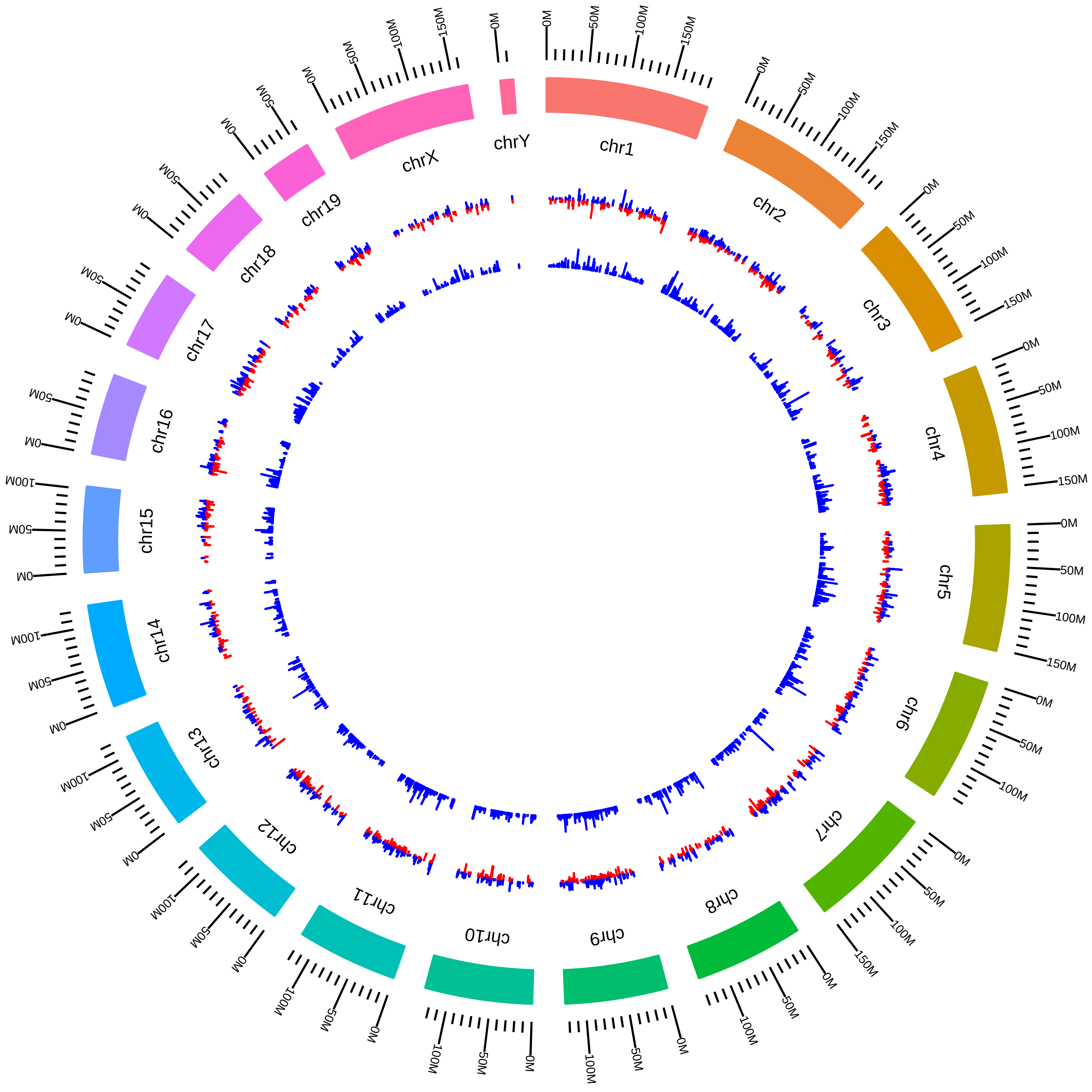

### rsfSf3a1MmSAlper_KO_vs_WT.pdf

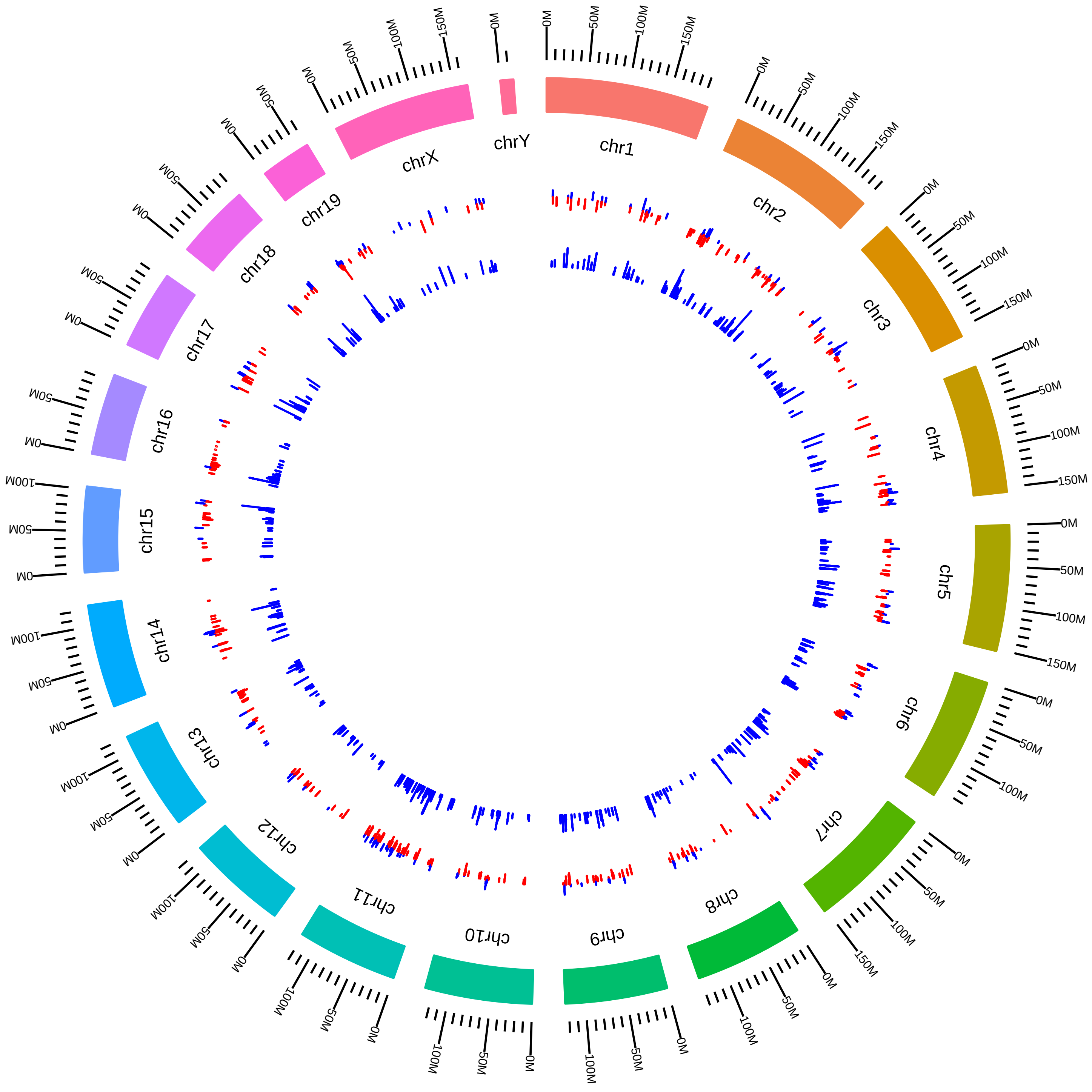

### rsfSf3b1MmBEbert2016_IVsW.pdf

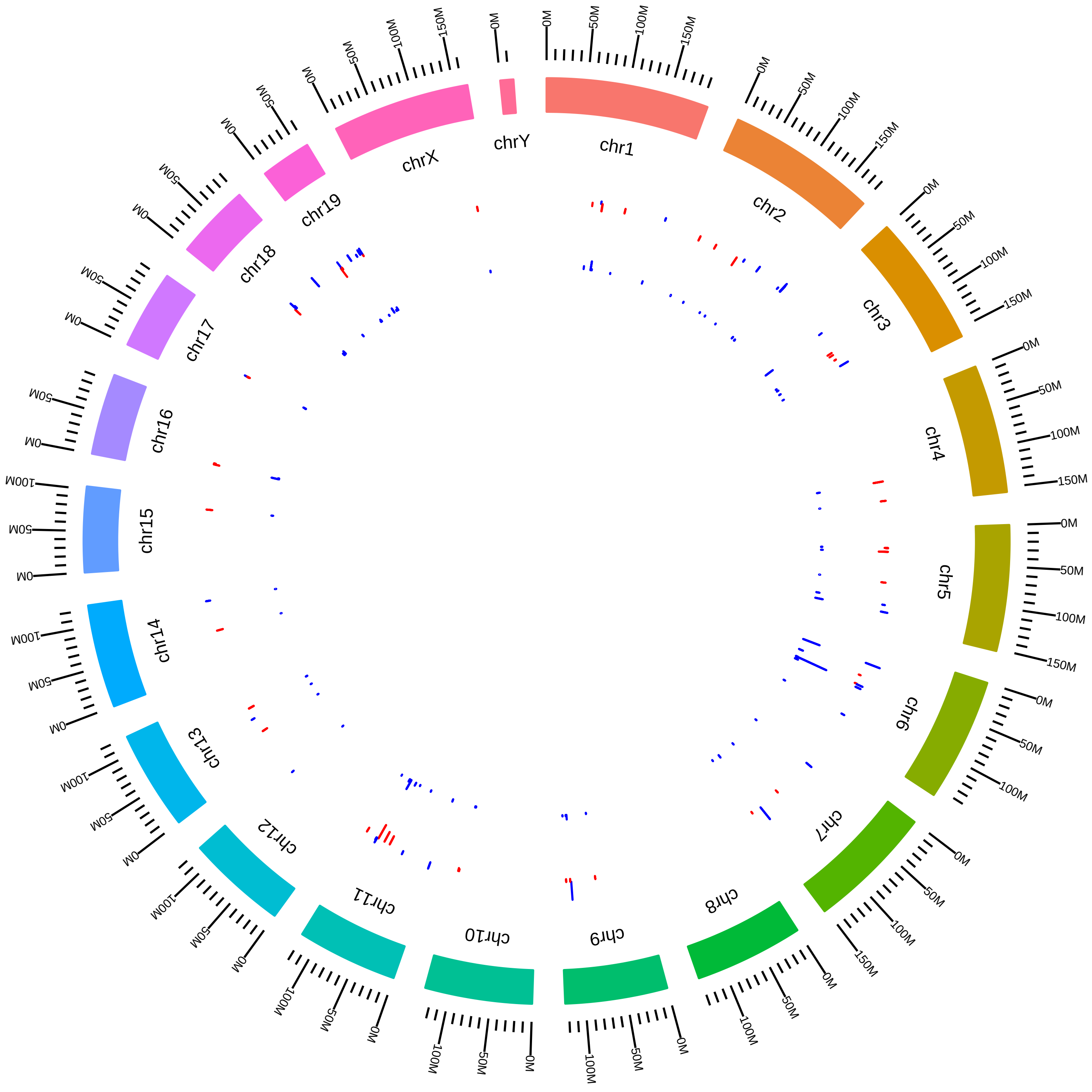

### rsfSf3b1MmSBuonamici2015_SVsW.pdf

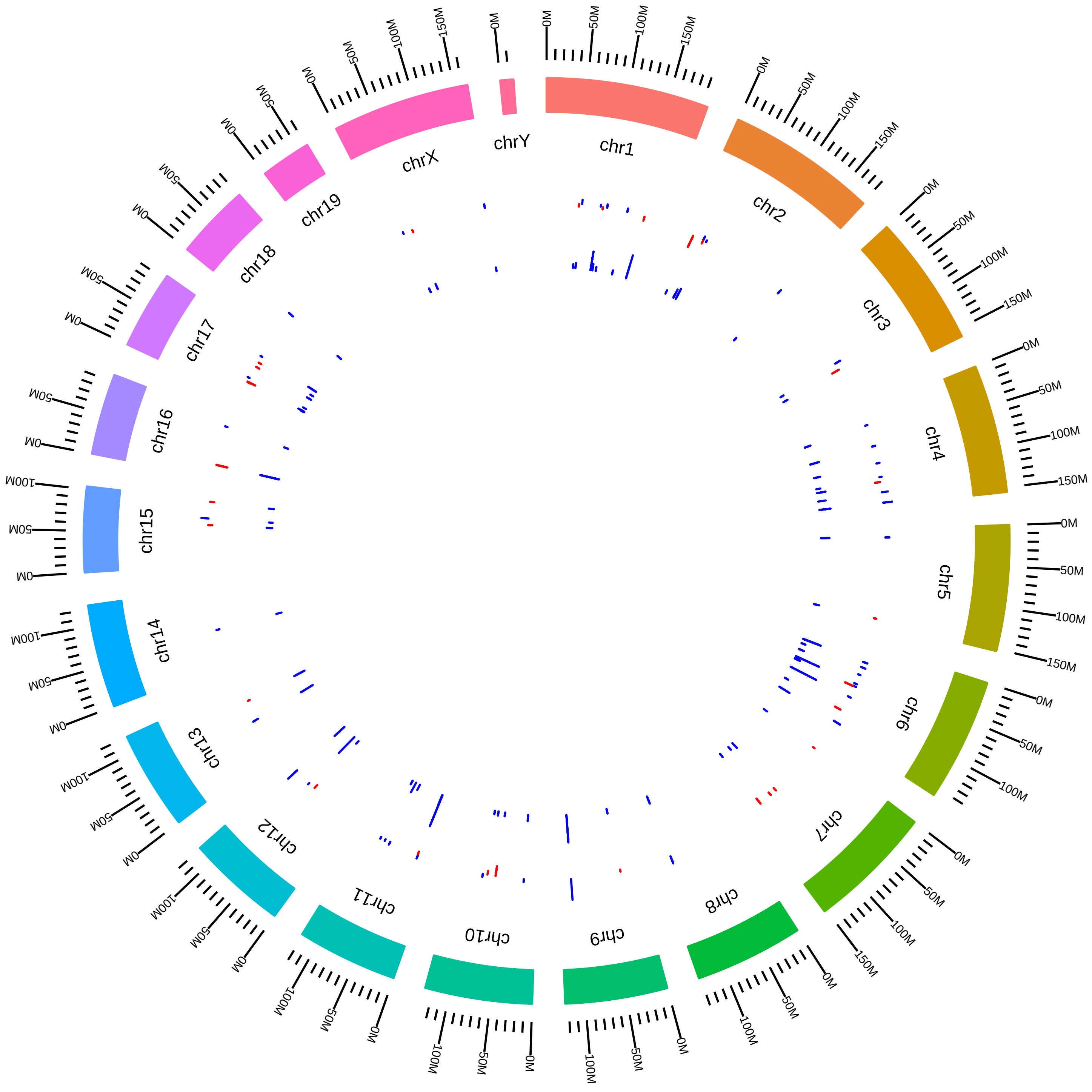

### rsfSf3b1MmWKrek_CFVsNN.pdf

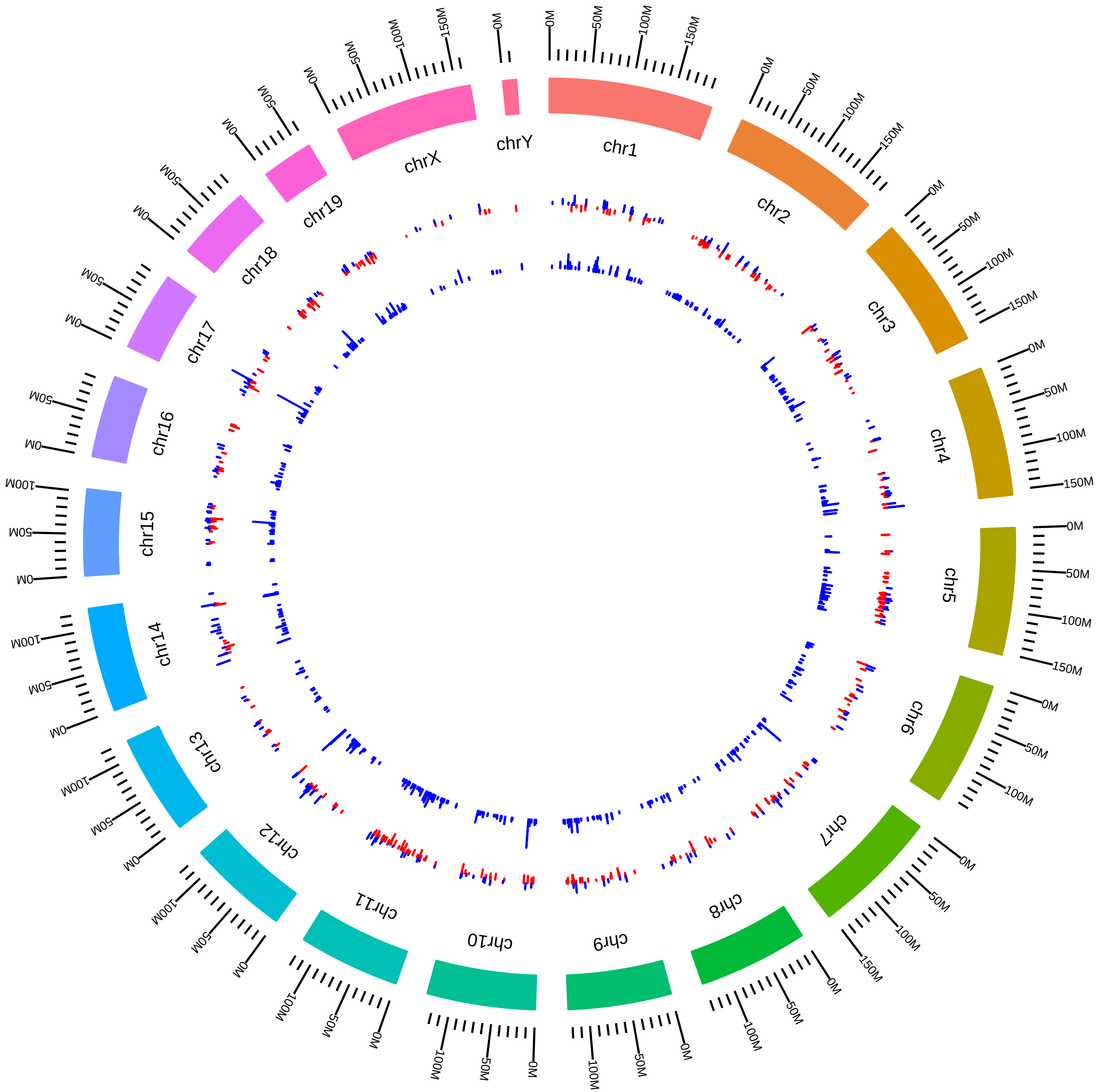

### rsfSmn1MmMButchbach2014_KVsW.pdf

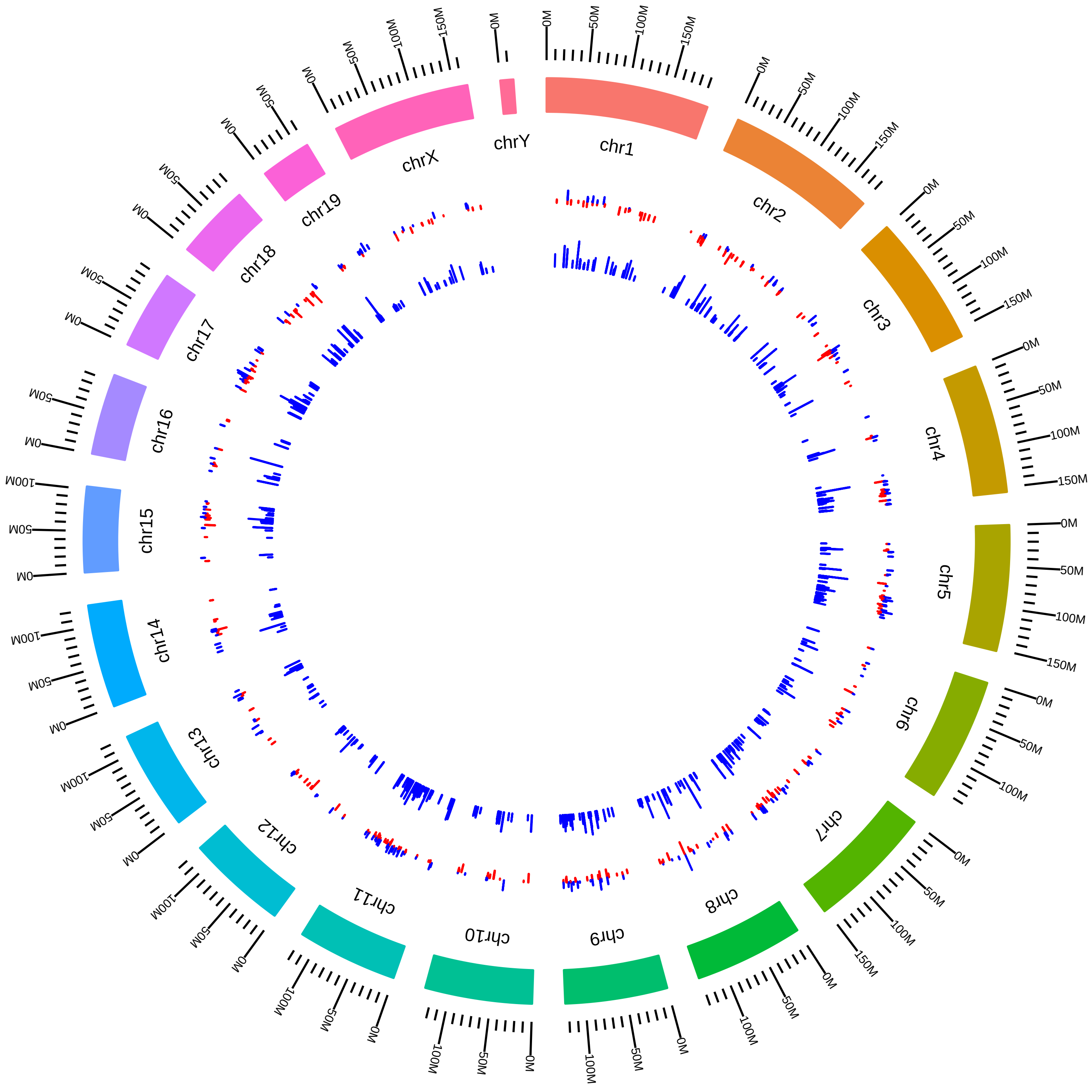

### rsfSrrm4MmBBlencowe2014_DVsC.pdf

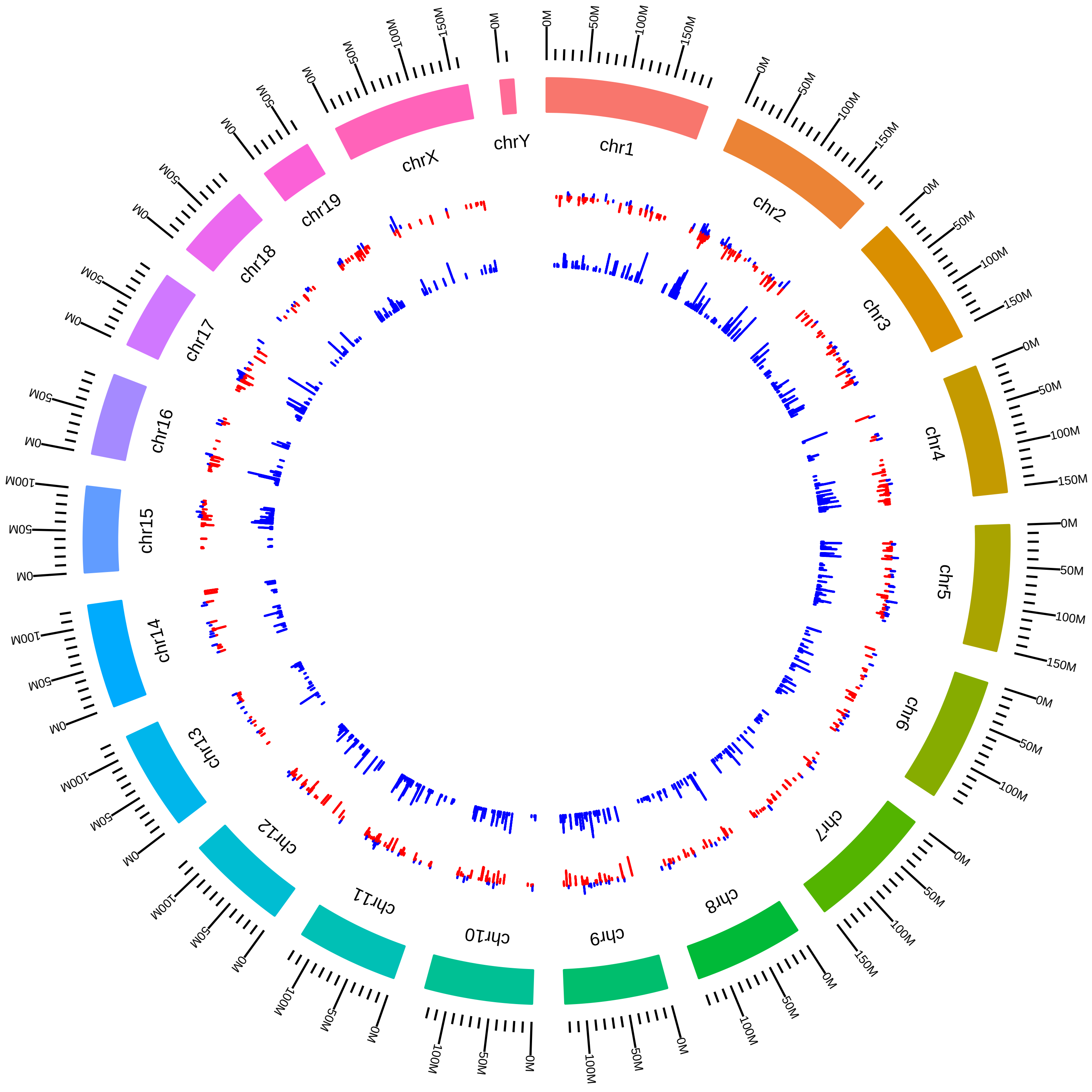

### rsfSrrm4MmBBlencoweCTX_KO_vs_WT.pdf

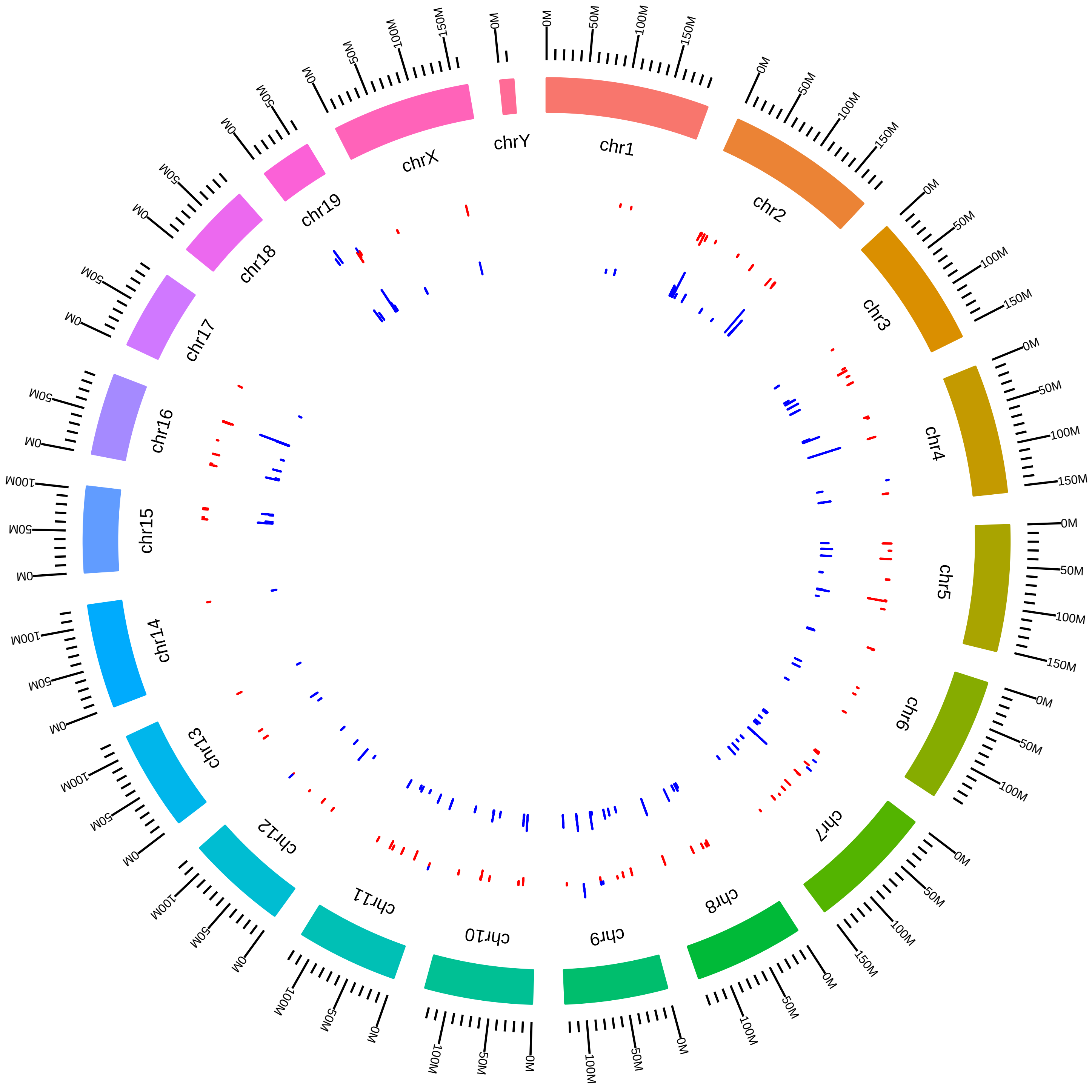

### rsfSrrm4MmBBlencoweHPC_KO_vs_WT.pdf

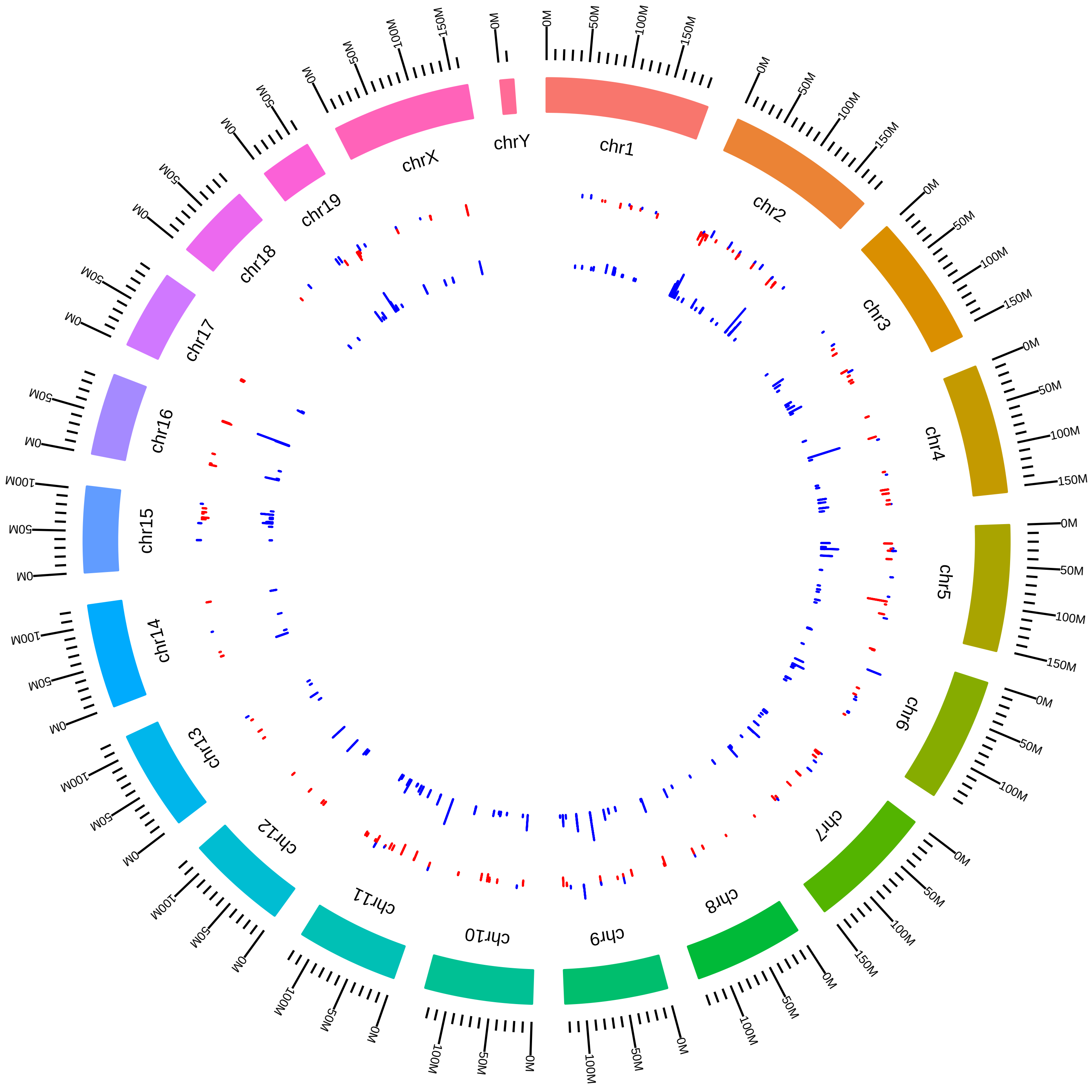
